# *Streptococcus pneumoniae* vaccine serotype persistence following 13-valent pneumococcal conjugate vaccine introduction in Mongolia: investigating changes in epidemiology, immunology and virulence

**DOI:** 10.1101/2025.09.23.678142

**Authors:** Paige Skoko, Sam Manna, Laura K. Boelsen, Jason Hinds, Stephanie W. Lo, Casey L. Pell, Belinda D. Ortika, Rachel A. Higgins, Vicki Bennett-Wood, Hanwen Hou, Odgerel Tundev, Eileen M. Dunne, Cattram D. Nguyen, Fiona M. Russell, E. Kim Mulholland, Tuya Mungun, Paul V. Licciardi, Claire von Mollendorf, Stephen D. Bentley, Catherine Satzke

## Abstract

**Background:** *Streptococcus pneumoniae* is a leading cause of pneumonia globally. Vaccine serotypes can persist despite pneumococcal conjugate vaccine (PCV) introduction. To examine serotype persistence, we leveraged 6,545 nasopharyngeal swabs collected from children hospitalised with pneumonia before and after PCV13 introduction in Mongolia and undertook molecular, epidemiological and experimental analyses.

**Methods:** Serotype, genetic lineage and antimicrobial resistance genes were inferred from DNA microarray. Patients carrying lineages that were predominant pre- and post-PCV introduction were examined for differences in disease severity. We also compared the pre- and post-PCV lineages by bacterial adhesion to hydrocarbon (BATH) and enzyme-linked immunosorbent (ELISA) assays. Capsule gene expression was measured by quantitative reverse transcription polymerase chain reaction (qRT-PCR), capsule thickness by transmission electron microscopy, and virulence using an infant mouse model.

**Findings:** Changes in lineage composition were observed within serotypes 6A, 6B, 14, 19F and 23F over the six-year surveillance period. Children carrying pre-PCV lineages were more likely to have severe pneumonia than those carrying post-PCV lineages (serotypes 6B, 14 and 19F). Pre-PCV lineages were more likely to be multi-drug resistant than post-PCV lineages (serotypes 6A, 6B and 23F). For serotype 6B the post-PCV lineage had higher cell surface hydrophobicity, lower IgG, lower expression of capsule genes and evidence of thinner capsule than the pre-PCV lineage. Notably, the post-PCV 6B lineage was also less virulent in mice than the pre-PCV 6B lineage.

**Interpretation:** Despite persistence of vaccine serotypes in highly vaccinated populations, vaccination may select for lineages with differences in capsule and antibody binding and that are less virulent. When evaluating the true value of vaccination, it is important to consider factors beyond serotype alone.

**Funding:** National Health and Medical Research Council, Murdoch Children’s Research Institute, GAVI, the Vaccine Alliance.

## Introduction

*Streptococcus pneumoniae* (the pneumococcus) is a leading cause of bacterial pneumonia, with high global burden especially in children from low-middle income countries.^1^ The pneumococcus is carried in the human nasopharynx,^2^ and although carriage is generally asymptomatic, it is critical for both transmission and disease.^2,3^

The polysaccharide capsule is the main pneumococcal virulence factor.^3^ There are over 100 capsular serotypes.^4,5^ Pneumococcal conjugate vaccines (PCVs) generate serotype-specific protection for vaccinees. PCVs also decrease carriage of vaccine serotypes, reducing the likelihood of transmission, thereby providing indirect (herd) protection for unvaccinated individuals.^2^ PCVs exert a powerful selective force that can result in significant changes to the pneumococcal population including serotype replacement, whereby vaccine serotypes are subsequently replaced with non-vaccine serotypes in carriage and disease.^6^

While PCV introduction decreases the carriage prevalence of vaccine serotypes, vaccine serotypes can persist in some settings. Some persistent vaccine serotypes are accompanied by changes in genetic lineage composition with important phenotypic differences.^7,8^ For example, post-PCV surveys in Malawi found that serotypes 3 and 23F persisted, with some lineages displaying increased resistance to antimicrobials,^7^ and reduced opsonophagocytic killing.^8^ However, few studies have combined genotypic and phenotypic analyses using pre- and post-PCV data, so the extent and importance of serotype persistence remains unclear.

Previously, we undertook a large surveillance program of children hospitalised with pneumonia in Mongolia.^9^ The 13-valent PCV (PCV13) was introduced into the immunisation schedule with high coverage. Whilst the prevalence of vaccine serotypes declined over time, some vaccine serotypes persisted.^10^ Here, we explore the phenomenon of vaccine serotype persistence following PCV introduction. We combine clinical, molecular, epidemiological and experimental data, to examine persistence of vaccine serotypes in the post-PCV era and its potential public health importance.

## Methods

### Pneumonia surveillance in Mongolia

In Mongolia, phased introduction of PCV13 commenced in 2016, with pneumonia surveillance in children aged 2-59 months carried out from 2015-2021 as described previously.^10^ Children were enrolled if they were admitted to hospital and met specified criteria as described previously.^10^ For vaccination status, ‘undervaccinated’, refers to children <12 months of age who received 0-1 doses of PCV13 or no dose if ≥12 months of age. ‘Vaccinated’ is defined as receipt of ≥2 doses of PCV13 if <12 months of age or ≥1 dose of PCV13 administered if ≥12 months of age.^10^

Nasopharyngeal swabs from consenting participants were collected and stored in accordance with WHO guidelines.^11^ As described previously, pneumococcal detection and quantification were conducted by *lytA* quantitative PCR on DNA extracted from nasopharyngeal swabs.^10,12^ *lytA*-positive samples were cultured on Horse Blood Agar (HBA) plates supplemented with 5 µg/ml gentamicin (Thermo Fisher Scientific). DNA extracted from the resultant growth underwent molecular serotyping by microarray (Senti-SPv1·5, BUGS Bioscience).^10^

### Identification of pneumococcal lineages and antimicrobial resistance genes from nasopharyngeal swabs

Microarray data were used to infer genetic lineage defined by Global Pneumococcal Sequence Cluster (GPSC, PopPunk GPS database v4) and detect the presence of antimicrobial resistance (AMR) genes.^13^ Pre-PCV lineages were defined as lineages that dominated before vaccine introduction and in the early years where vaccine coverage in our study participants was ≤20%. Otherwise, lineages were defined as post-PCV. To examine population structure, dendrograms were generated from array-CGH analysis of genomic profiles based on gene presence or absence, excluding samples with low intensity QC flags. Multi-drug resistance was defined as detection of ≥ 3 antimicrobial resistance genes in a sample.^13^

### Bacterial adhesion to hydrocarbon assay

A bacterial adhesion to hydrocarbon (BATH) assay was used to assess cell surface hydrophobicity.^14^ Overnight bacterial growth on HBA plates (37°C, 5% CO_2_) was harvested and suspended in 1 ml Phosphate-buffered saline (PBS) to an optical density at 600 nm (OD_600_) of 0·2. Viable counts were performed (Time 0). Subsequently, 200 µl of hexadecane (Sigma-Aldrich) was added to the bacterial suspension, vortexed and incubated (37°C, 5% CO_2_) for 30 min. Following incubation, the lower aqueous phase was removed, and pneumococci enumerated by viable count (Time 30). The percent hydrocarbon adherence was calculated as: [(Time 0 – Time 30)/Time 0] x 100.

### Immunological assays

Enzyme Linked Immunosorbent Assay (ELISA) was used to measure antibody binding to pneumococci. ELISAs were performed as previously described^15^ and as detailed in the Supplementary material. Optical density was measured using a microplate reader (BioTek) at OD_450_ (630nm reference filter) and serotype-specific IgG concentrations reported as mean ELISA Units/ml (EU/ml).

### Serotype 6B capsule operon and gene comparison

To examine genomes for differences in the capsule operon we undertook whole-genome sequencing from pneumococcal isolates as previously described^11^ and as detailed in the Supplementary material.

### Quantitative reverse transcriptase polymerase chain reaction (qRT-PCR)

Extraction of RNA from pneumococci was performed using the RNeasy Mini Kit (Qiagen) according to the manufacturer’s instructions with the incorporation of an additional lysis step as previously published^16^ and as detailed in the Supplementary material. We used qRT-PCR to measure levels of gene expression of capsule genes *wzg, wzy* and *rmlB*. Extracted RNA was used for first strand cDNA synthesis using the iScript cDNA synthesis kit (Biorad) as described previously^16^ and as detailed in the Supplementary material.

### Capsule thickness

For transmission electron microscopy, overnight bacterial growth on HBA plates was suspended in 1 ml PBS to OD_600_ of 0·4. Bacteria were fixed and prepared for microscopy^17^ as detailed in the Supplementary material.

### Mouse model of pneumococcal disease

Five-day old C57BL/6 mice were given 2×10^3^ colony forming units (CFU) of pneumococci intranasally (without anaesthesia) and monitored for up to 15 days post-infection. Mice exhibiting one or more signs of disease (e.g. weight loss, lethargy, lack of response to provocation, hunched posture, ruffled fur or balance issues) were euthanised upon reaching our pre-defined humane intervention criteria. Nasopharyngeal, lung, blood, brain and middle ear tissues were harvested, and each placed in 1·5 ml RPMI 1640 medium (Sigma-Aldrich). Blood was also collected and placed in 0·5 ml microvette lithium heparin-coated tubes (Sarstedt). To enumerate pneumococci, tissue homogenates were serially diluted, plated on HBA plates supplemented with 5 µg/ml gentamicin and incubated overnight (37 °C, 5% CO_2_).

### Data analysis

Analysis was undertaken and figures generated in GraphPad Prism (version 9·4·2), R (version 4·1·2) and R package ‘ggplot2’ (version 3·3·5). Dendrograms were generated using R package ‘ggtree’. For each of the vaccine serotypes tested in experimental assays, four isolates were randomly selected from each lineage using the Rand() and Index() formulas in Microsoft Excel (version 16·39). Differences in proportions were compared using Fisher’s exact test. Unless otherwise stated, bacterial density data were log_10_ transformed, expressed as median and inter-quartile range (IQR) and groups compared using the Mann-Whitney test. Time to event data were compared using Kaplan-Meier survival curve and Mantel-Cox test.

### Ethical approval

The pneumonia surveillance program was approved by the Mongolian Ministry of Health National Ethics Committee for Health Research and the Royal Children’s Hospital Human Research Ethics (HREC 33203). Written and informed consent was obtained from all parents/caregivers of children enrolled in the study. The study has animal ethics approval from the Murdoch Children’s Research Institute (MCRI) Animal Ethics Committee (A945 and A976). All mouse experiments were conducted in accordance with the Australian Code of Practice for the Care and Use of Animal for Scientific Purposes.^18^

## Results

### Lineage composition of persistent vaccine serotypes following vaccine introduction

Of 15,411 nasopharyngeal samples collected from children hospitalised with pneumonia between 2015-2021 in Mongolia, 6,545 were screened for pneumococci and 3,056 (47%) were found to contain pneumococci.^10^ Serotyping data was available for 2,410 samples, with 50% (1216/2410) containing vaccine serotypes, 59% (1412/2410) containing non-vaccine serotypes, and 9% (218/2410) containing multiple serotypes.

Vaccine serotype carriage among hospitalised children declined over the six-year sampling period from 451/706 (64%) pre-PCV to 659/1695 (39%) post-PCV (p<0·0001).^10^ For most vaccine serotypes, prevalence declined over time (Figure 1A). Interestingly, the proportion of pneumococcal-positive samples containing 6B in 2015 declined from 21/288 (7%) to 11/409 (3%) in 2018 (p=0.005), but then increased in prevalence in 2020 (18/254 (7%) v 2018, p=0.01).

**Figure 1.**
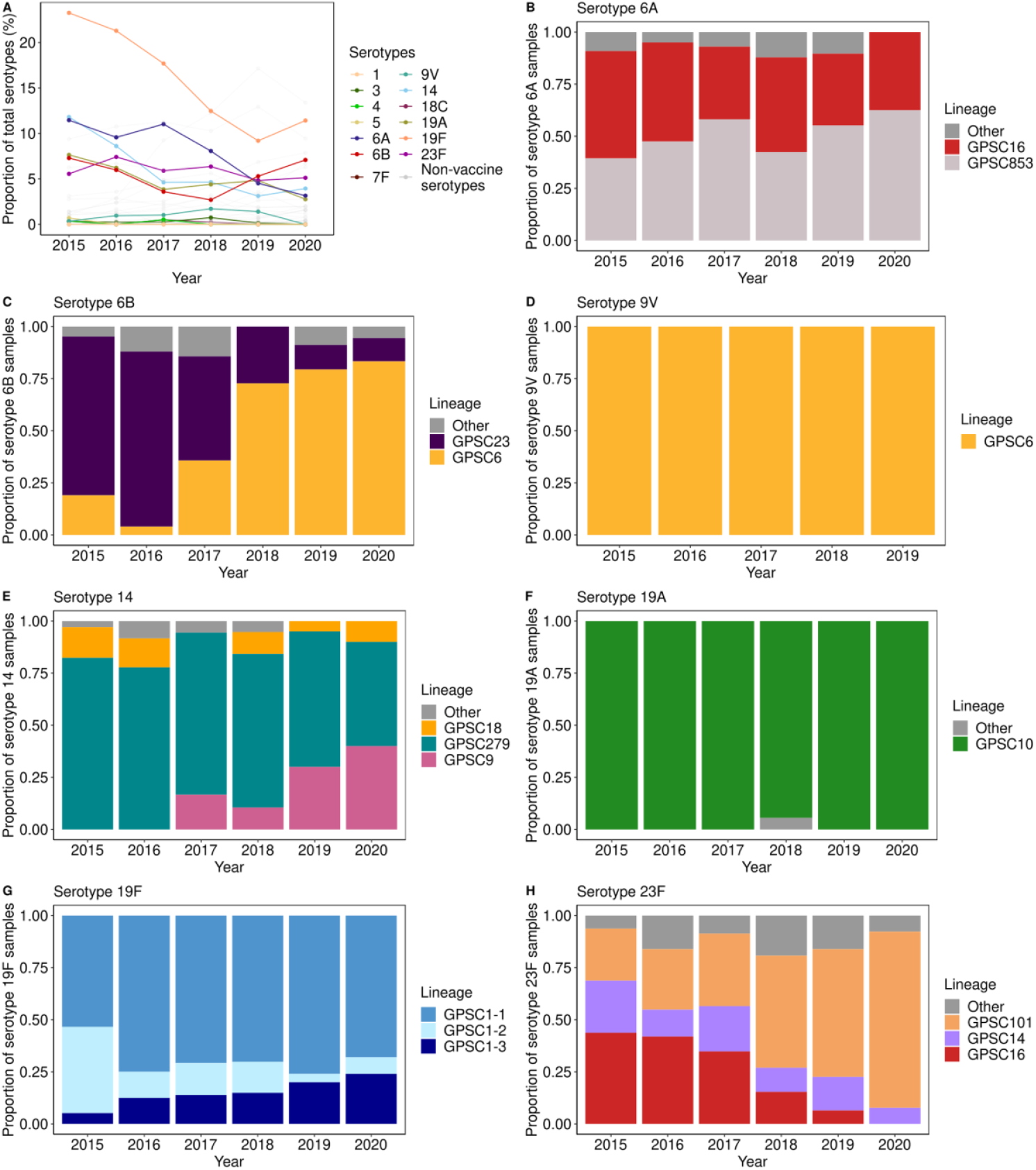
Vaccine serotypes and lineages circulating in Mongolia between 2015 and 2020. Pneumococcal serotypes as a proportion of all children carrying pneumococci (A). Lineage composition for persistent vaccine serotypes 6A, 6B, 9V, 14, 19A, 19F and 23F (panels B-H, respectively) by year. Vaccine serotypes 1, 3, 4, 5, 7F and 18C were uncommon (n=0, n=7, n=3, n=2, n=0 and n=4 samples, respectively) and were not examined for lineage composition. For samples with multiple serotype calls, the most abundant serotype was used to infer the GPSC. When a lineage was detected in fewer than 10 samples for a given serotype, it was termed ‘other’. Bars are coloured by lineage (Global Pneumococcal Sequence Cluster or sub-cluster).

We determined lineage composition changes over time within each vaccine serotype by inferring lineage from microarray analysis (Figure 1B-E). Within serotype 6A, lineage GPSC16 reduced from 17/33 (52%) in 2015 to 3/8 (38%) in 2020 (p=0·70), and lineage GPSC853 increased from 13/33 (39%) to 5/8 (63%) over the same time period (p=0·72) (Figure 1B). For serotype 6B, in 2015, GPSC23 was the predominant lineage (16/22, 73%), but this decreased in 2020 (2/18 (11%); p<0·01) (Figure 1C). In contrast, the GPSC6 lineage was relatively uncommon in 2015 (4/22, 18%) but increased to 15/18 (83%) in 2020 (p<0·01). Serotype 9V was comprised of one GPSC (GPSC6) over the six-year sampling period (Figure 1D). Within serotype 14 GPSC279 was detected in 28/34 (82%) samples in 2015 but reduced to 5/10 (50%) in 2020 (p=0·09) (Figure 1E). In contrast, GPSC9 was not detected in 2015 nor 2016 (0/34 (0%) and 0/36 (0%), respectively) but emerged in 2017, and by 2020 was detected in 4/10 (40%) samples containing serotype 14 (p<0·01). Serotypes 19A and 19F were each almost completely comprised of a single GPSC (GPSC10 and GPSC1, respectively) (Figure 1F and Supplementary Figure S1, respectively). Within serotype 23F, GPSC16 decreased over time (7/16 (44%) in 2015 v 0/13 (0%) in 2020, p=0·02) and was replaced by GPSC101 (4/16 (25%) in 2015 v 11/13 (85%) in 2020, p<0·01) (Figure 1H).

A dendrogram of samples containing vaccine serotypes was generated to further examine the relationship between serotype, lineage and year of sample collection (Figure 2 and Supplementary Figure S2). Within serotypes 6A, 6B, 9V, 14, 19A and 23F, we observed clustering by lineage. For serotype 19F, we observed three distinct sub-lineages within GPSC1 (GPSC1-1, GPSC1-2 and GPSC1-3) (Figure 2). GPSC1-2 declined over time from 24/58 (41%) in 2015 to 2/25 (8%) in 2020 (p=0·002), and GPSC1-3 increased from 3/58 (5%) in 2015 to 6/25 (24%) in 2020 (p=0·019) (Figure 1G).

**Figure 2.**
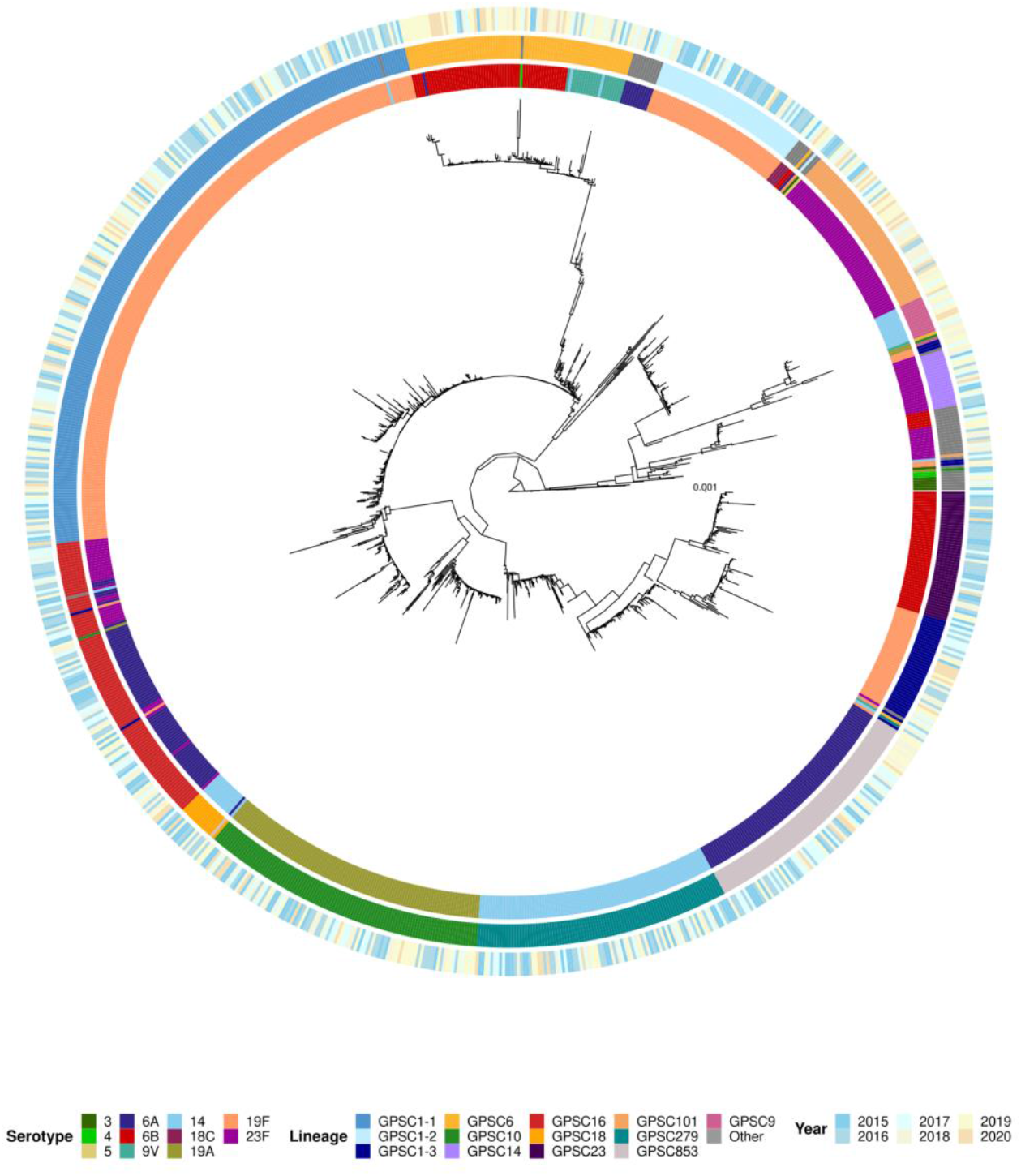
Dendrogram of pneumococcal vaccine serotypes circulating in Mongolia from 2015 to 2020. The dendrograms are based on array-CGH analysis of the genome backbone component of microarray data,^13^ scale bar represents the divergence between samples. Dendrograms are overlayed with serotype (inner ring), lineage (based on Global Pneumococcal Sequence Cluster or sub-cluster; middle ring) and year (outer ring).

### Colonisation density, disease severity and vaccine status of children with pneumonia

High pneumococcal density in the nasopharynx is associated with disease severity and transmission.^3^ For the vaccine serotypes that displayed lineage replacement following vaccine introduction, we compared the median nasopharyngeal density of the predominant pre-PCV and post-PCV lineages, finding they were similar for 6B (p=0·32), 14 (p=0·29), 19F (p=0·48) and 23F (p=0·77) (Supplementary Figure S3). The density of 6A pre-PCV lineage GPSC16 was slightly lower than post-PCV lineage GPSC853 (an absolute difference of 0·5 log_10_ genome equivalents/ml; p<0·01) (Supplementary Figure S3).

We assessed the proportion of children with different markers of disease severity carrying pre-PCV or post-PCV lineages. The proportion of hypoxic children were similar for the pre- and post-PCV lineages of serotypes 6A (p=0·59), 6B (p=0·46), 14 (p=1·00) and 19F (p=0·46) (Table 1). For serotype 23F, the proportion of hypoxic children was higher for those carrying the pre-PCV lineage compared with the post-PCV lineage (p=0·03). Compared with the post-PCV lineage, more carriers of the pre-PCV lineage were hospitalised with primary endpoint pneumonia for serotypes 6B (p=0·02), 14 (p=0·01) and 19F (p=0·03). For serotype 6A and 23F, the proportion of children hospitalised with primary endpoint pneumonia were similar for the pre- and post-PCV lineages (p=0·17 and p=0·27, respectively) (Table 1).

**Table 1.** Disease severity for Mongolian children hospitalised with pneumonia and carrying a vaccine serotype where there was evidence of lineage replacement following PCV introduction.

|  | Serotype 6A |  |  | Serotype 6B |  |  | Serotype 14 |  |  | Serotype 19F |  |  | Serotype 23F |  |  |
| --- | --- | --- | --- | --- | --- | --- | --- | --- | --- | --- | --- | --- | --- | --- | --- |
|  | GPSC16<br>n/N (%) | GPSC853<br>n/N (%) | p-value<sup>c</sup> | GPSC23<br>n/N (%) | GPSC6<br>n/N (%) | p-value<sup>c</sup> | GPSC279<br>n/N (%) | GPSC9<br>n/N (%) | p-value<sup>c</sup> | GPSC1-2<br>n/N (%) | GPSC1-3<br>n/N (%) | p-value<sup>c</sup> | GPSC16<br>n/N (%) | GPSC101<br>n/N (%) | p-value<sup>c</sup> |
| Hypoxic<sup>a</sup> | 22/77<br>(29) | 21/88<br>(24) | 0.59 | 11/47<br>(23) | 10/60<br>(17) | 0.47 | 14/92<br>(15) | 2/15<br>(13) | 1.00 | 12/49<br>(25) | 8/45<br>(18) | 0.46 | 11/31<br>(36) | 9/63<br>(14) | 0.03 |
| Primary<br>endpoint<br>pneumonia<sup>b</sup> | 29/69<br>(42) | 25/82<br>(31) | 0.17 | 24/50<br>(48) | 14/58<br>(24) | 0.02 | 49/92<br>(53) | 2/14<br>(14) | 0.01 | 29/51<br>(57) | 15/45<br>(33) | 0.03 | 12/29<br>(41) | 18/64<br>(28) | 0.24 |

<sup>a</sup>Hypoxic defined as an oxygen saturation of <90,<sup>10</sup> <sup>b</sup>WHO defined primary endpoint pneumonia.<sup>11</sup> <sup>c</sup>Disease severity data were compared using Fisher's exact test. For each serotype, the lineage that was common pre-PCV is listed first, followed by the lineage that was most common post-PCV. Hypoxic and primary endpoint pneumonia data were not available for every child.

For fully vaccinated children, a lower proportion carried pre-PCV lineages for serotypes 6B (p<0·01), 14 (p=0·03), 19F (p=0·06) and 23F (p=0·01) than post-PCV lineages. No difference was found for serotype 6A (p=0·45) (Supplementary Table S1).

### Antimicrobial resistance in pre-PCV and post-PCV lineages

For the selected serotypes and samples, we compared the number of antimicrobial resistance genes detected in pre-PCV and post-PCV lineages. At least one antimicrobial resistance gene was detected in 98% (416/424) of samples assessed. The proportion of pneumococci carrying the *ermB* gene (confers resistance to macrolides) was higher in the pre-PCV lineages than in the post-PCV lineages for serotypes 14 (p<0·0001) and 23F (p<0·0001), but no differences were detected for 6B and 19F (Supplementary Table S2). The proportion of pneumococci carrying the *mefA* gene (confers resistance to macrolides) was higher in the pre-PCV than the post-PCV lineage for serotype 6A (p<0·0001), but higher in the post-PCV lineage for serotype 23F (p=0·001). The proportion of pneumococci carrying the *cat* gene (confers resistance to chloramphenicol) was higher in the pre-PCV lineages than in the post-PCV lineages for serotypes 6A (p<0·0001), 6B (p<0·0001), and 23F (p<0·0001), but there were no differences for serotypes 14 (p=0.06), and 19F (p=1·00). The proportion of pneumococci that were multi-drug resistant (defined as ≥3 resistance genes detected) was higher in the pre-PCV lineages than in the post-PCV lineages for serotype 6A (p<0·0001), 6B (p<0·0001) and 23F (p<0·0001), however, there was no difference found for serotypes 14 (p=0·06) and 19F (p=0·20) (Supplementary Table S2).

### Differences in capsule of pre-PCV and post-PCV lineages

Given PCV targets the pneumococcal polysaccharide capsule, we performed several assays to examine the capsule. First, we looked at changes in cell surface hydrophobicity, using the BATH assay, which is reported to be negatively correlated with capsule amount.^14^ We tested control strains (Supplementary Figure S4) and four randomly selected isolates from each pre-PCV and post-PCV lineage for serotypes 6A, 6B, 14, 19F and 23F. Although randomly selected, the isolates represent the diversity within each lineage (Supplementary Figure S2). Levels of hydrophobicity were similar for isolates representing pre-PCV or post-PCV lineages of serotypes 6A (p=0·08), 14 (p=0·99), 19F (p=0·29) and 23F (p=0·63) (Figures 3A i, iii-v, respectively). For serotype 6B, the pre-PCV lineage (GPSC23) was less hydrophobic than the post-PCV lineage (GPSC6) (p=0·02) (Figure 3A ii). We next compared the ability of pre-PCV and post-PCV lineages to bind capsule antibodies from sera derived from PCV13-vaccinated children^15^ (Figure 3B). For serotype 6A, there was some evidence of a difference in antibody binding between pre-PCV lineage GPSC853 (mean 26 EU/ml [95% CI, 16 – 36]) and post-PCV lineage GPSC16 (22 EU/ml [15 – 29]; p=0·05) (Figure 3B i). Similarly, for serotype 23F, pre-PCV lineage GPSC16 showed lower antibody binding (28 EU/ml [14 – 42]) compared with post-PCV lineage GPSC101 (31 EU/ml [15 – 48]; p=0·05) (Figure 3B v). For serotype 6B, the pre-PCV lineage GPSC23 had higher antibody binding (84 EU/ml [47 – 121]) than the post-PCV lineage GPSC6 (63 EU/ml [36 – 90]; p=0·03) (Figure 3B ii).

**Figure 3.**
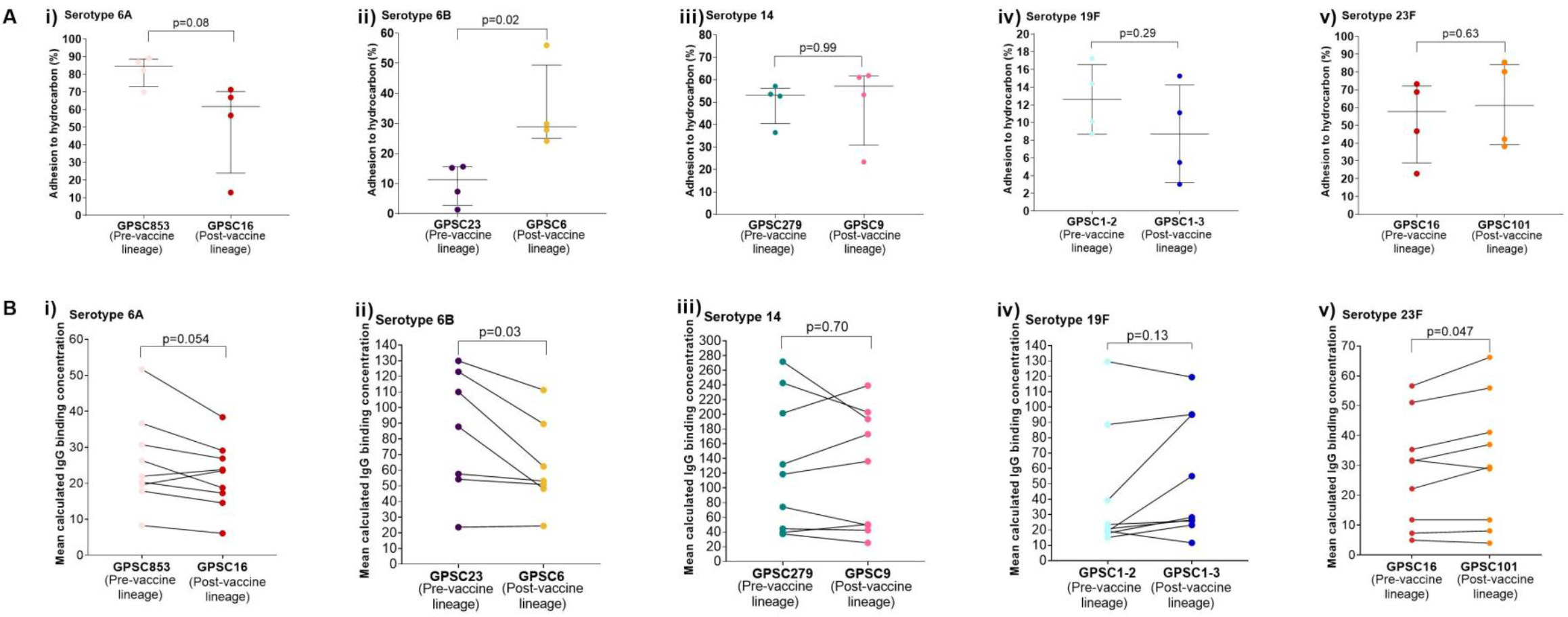
Adhesion to hydrocarbon of pre-PCV and post-PCV lineages using Bacterial Adhesion to Hydrocarbon (BATH) assay (A) and antibody binding of pre-PCV and post-PCV lineages using ELISAs (B). The BATH assay was used to measure amount of binding of serotype 6A, 6B, 14, 19F and 23F polysaccharide capsule to hexadecane hydrocarbon. There were four isolates of each lineage tested. Each individual dot point represents one isolate. Error bars represent the median and interquartile range. Percent adhesion to hydrocarbon was calculated for serotypes 6A, 6B, 14, 19F and 23F (Panel A i-v, respectively) and differences were assessed using a paired t-test. Amount of antibody binding to serotype 6A, 6B, 14, 19F and 23F (Panel B i-v, respectively). Four isolates of each lineage were tested. Data from each individual serum sample is connected by a solid line. Sera was derived from seven (for 6B) and nine (for 6A, 14, 19F and 23F) PCV13 vaccinated children with high titre antibodies (>8 μg/ml) to all serotypes tested. Each individual dot point is the mean of the four isolates tested against each serum. Differences were assessed using a paired t-test.

*Differences in capsule thickness and virulence for serotype 6B pre-PCV and post-PCV lineages* The post-PCV lineages of serotype 6B showed clear differences in cell surface hydrophobicity and antibody binding, consistent with lower capsule production. However, the capsule loci and regulatory sequences were all identical between the pre and post-vaccine lineages. The exception was one GPSC6 isolate which had 88 single nucleotide polymorphisms over two genes (*wciO* and *wciP*) (Supplementary figure S5). Next, we examined if there were any differences in expression of capsule genes between the two lineages. We performed qRT-PCR targeting three genes across the capsule locus (*wzg, wzy* and *rmlB*). We observed lower expression of capsule genes in the post-PCV lineage GPSC6 relative to our pre-PCV lineage GPSC23 (-1·46, -1·39 and -1·18-fold change in *wzg, wzy* and *rmlB*, respectively) (Figure 4A). Moreover, we observed differences in the capsule thickness of pre-PCV and post-PCV lineages using transmission electron microscopy to measure capsule thickness (Figure 4B-D). At the middle of the cell, the capsule was thicker for the pre-PCV lineage (median of 44 nm [IQR 26 – 52]) compared with the post-PCV lineage (28 nm [19 – 43]) (p<0·01) (Figure 4D). There was a similar trend at the poles (pre-PCV, 85 nm [57 – 97]; post-PCV, 70 nm [52 – 93]), but this did not reach statistical significance (p=0·38). No differences were observed at the septum (pre-PCV, 19 nm [13 – 27]; post-PCV, 20 nm [13 – 25]) (Figure 4D). We also observed phenotypic differences in the capsule appearance (Figure 4D) between lineages with some displaying a bubble-like appearance. The proportion of cells with this appearance was lower in the pre-PCV lineage (16/74, 21·6%) compared with the post-PCV lineage (59/81, 72·8%) (p<0·0001).

**Figure 4.**
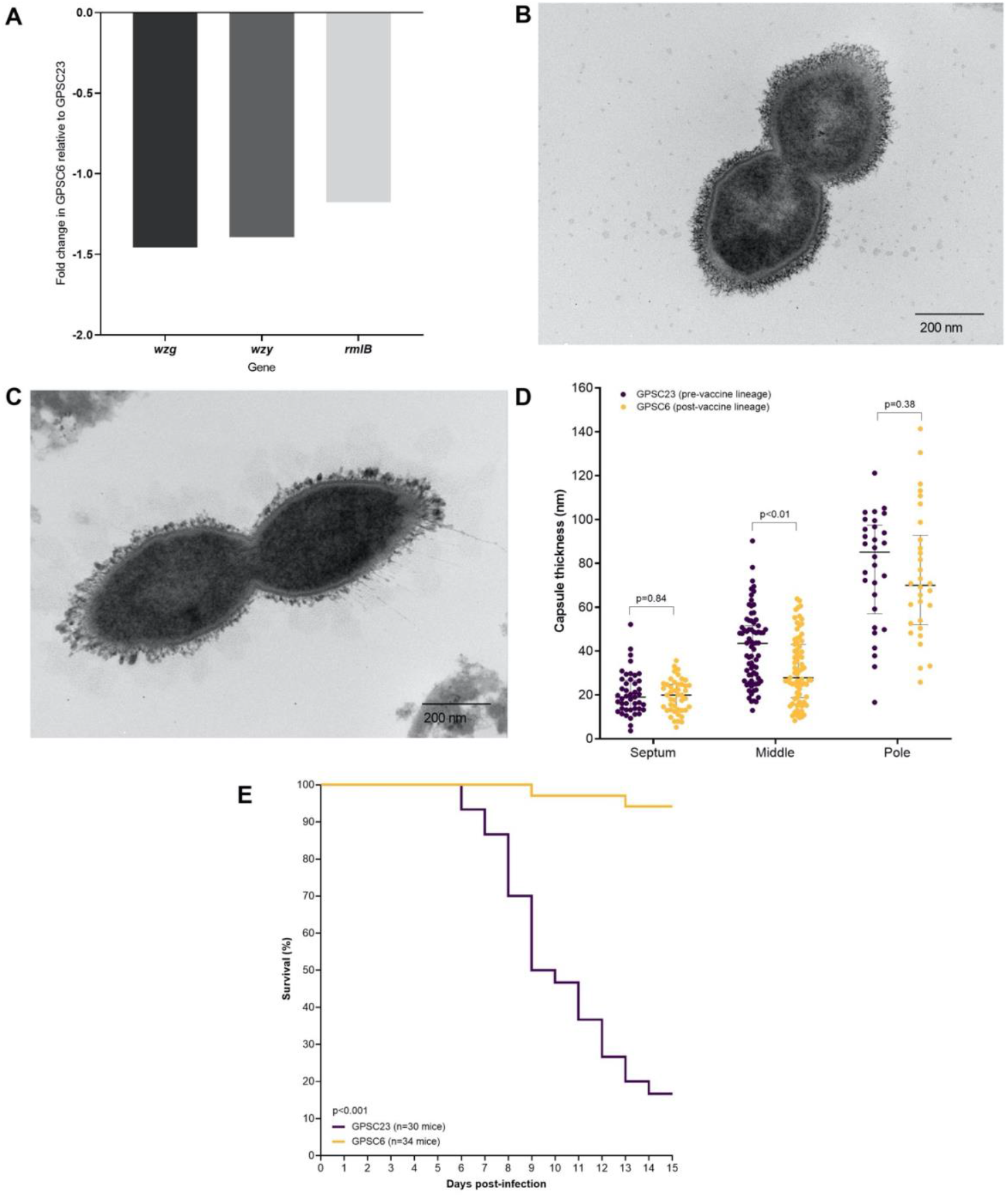
Capsule thickness and virulence for serotype 6B pre-PCV and post-PCV lineages. Expression of capsule genes *wzg, wzy* and *rmlB* were measured using quantitative reverse transcription polymerase chain reaction (qRT-PCR) (A). Data were normalised to the *gyrA* gene using the 2^-ΔΔCt^ method. Transmission electron microscopy was used to visualise capsule thickness with a representative serotype 6B pre-PCV lineage (GPSC23, B) and post-PCV lineage (GPSC6, C) shown. Capsule thickness (in nm) (D) was measured using Image J software. Measurements were taken at the septum, middle and pole of cells. Capsule thickness was compared using Mann-Whitney test with p-values shown. A p-value less than 0.05 was considered statistically significant. Individual dots represent an individual measurement, error bars represent the IQR. Murine model of invasive pneumococcal disease was used to examine differences in virulence between serotype 6B pre-(GPSC23) and post-(GPSC6) PCV lineages and presented as a survival curve (E). Survival was compared using Kaplan-Meier survival curve and Log-rank test.

The polysaccharide capsule is the main pneumococcal virulence factor.^3^ Given capsule is an important virulence factor, we observed differences in capsule between the serotype 6B pre-PCV and post-PCV lineages, we next examined their virulence using a murine model of disease. All mice had an initial 5-day asymptomatic period. After that, GPSC23 was more likely to transition to disease with 83% (25/30) of mice reaching our pre-defined humane intervention criteria compared with 6% (2/34) of mice infected with GPSC6 (p<0·001) (Figure 5E). Results were consistent for individual isolates of the same lineage (Supplementary Figure S6A). Where present, pneumococci were recovered from the lungs, blood, middle ear and brain of symptomatic mice (Supplementary Figure S6C).

## Discussion

In some settings, vaccine serotypes can persist in the population despite high vaccine coverage.^19^ This persistence can generate uncertainty in the value of PCV programs especially for governments and other vaccine policy decision-makers. In this study, we found that whilst certain vaccine serotypes persisted in Mongolia, there were substantial shifts in the pneumococcal lineages within most of these serotypes. Lineages that were more common post-PCV tended to carry fewer antimicrobial resistance genes and were found in carriers with lower clinical severity markers. Of note, the post-PCV lineage of serotype 6B also had lower expression of capsule genes, a thinner capsule, was more able to evade antibody binding, and had reduced virulence compared with the pre-PCV lineage of 6B in a murine model.

Few studies have examined lineage changes within persistent vaccine serotypes, particularly from low- and middle-income countries.^10,20-21^ Consistent with a study from Malawi,^20^ we identified changes to the lineage composition of serotypes 14 and 23F. Interestingly, we also noted an expansion of GPSC9 within serotype 14, however the lineages within serotype 23F that increased post-PCV were different in Mongolia and Malawi. Within serotype 6B, we found that lineage GPSC6 emerged in the post-PCV era. To our knowledge, GPSC6 has not been detected in 6B previously.^22^ These results illustrate that there can be both similarities and differences within and across settings, including in the specific lineages that emerge post-PCV introduction. This reflects the importance of implementing effective local surveillance to ensure such differences are captured and monitored over time.

The reasons underpinning vaccine serotype persistence are likely complex and may vary by serotype and setting. Our study, together with existing literature, emphasises the complexity of the pneumococcal population dynamics following vaccine introduction. For example, a comprehensive study on the pneumococcal population four years post-PCV13 introduction in Malawi found that whilst there were shifts in the metabolic types (MTs) present in several non-vaccine serotypes and that emergent MTs had higher levels of antimicrobial resistance, the patterns for other phenotypes such as growth, invasion and murine colonisation were less consistent.^7^

The selective pressure of PCV could contribute to the emergence of post-PCV lineages with enhanced vaccine evasion and ultimately resulting in persistence of vaccine serotypes post-PCV. This hypothesis is consistent with the observation that the post-PCV lineage of serotype 6B had reduced antibody binding, lower expression of capsule genes and thinner capsule. Moreover, evidence from Malawi that serotype 23F had non-synonymous genetic changes in the capsular polymerase gene *wzy*,^20^ and common post-PCV lineages of serotype 3 carriage isolates were less susceptible to opsonophagocytosis killing than lineages that were common pre-PCV, also support this hypothesis.

Moreover, the selection for less virulent and less multi-drug resistant lineages post-PCV adds confidence in current vaccine programmes. If children are more likely to carry less virulent strains of vaccine serotype pneumococci, this would likely translate to fewer cases of severe pneumonia. Such findings would be consistent with the broader literature on vaccine escape, for example the emergence of non-typeable (i.e. unencapsulated) *Haemophilus influenzae* following vaccine introduction protecting against type B capsule.^23^ However, in our study, selection of less virulent and less multi-drug resistant lineages varied by serotype, and continued surveillance is crucial to capture and monitor changes over time.

Our study has limitations. Due to reductions in pneumonia incidence (likely from PCV introduction and/or COVID non-pharmaceutical interventions) there are few data from 2021 so year was excluded from the analysis.^9^ In line with other studies, we primarily used results from microarray analysis to characterise antimicrobial resistance, and infer lineage.^10,13^ The selected genes serve to monitor overall antimicrobial resistance trends, but resistance to other clinically important antimicrobials (including penicillin and fluroquinolones) was not specifically explored and phenotypic antimicrobial susceptibility testing was not performed. As this is an observational study, we were unable to assess the specific contribution vaccine introduction has played in lineage selection. Our study has several important strengths. We utilised samples from a large and long-term vaccine impact program in children with pneumonia. There are very few pneumonia surveillance programs of this scale, particularly in low- and middle-income countries and in Asia generally.^10^ By analysing microarray data to gain insight into the pneumococcal population structure, we demonstrate the utility of such data for the multitude of studies already completed globally. Importantly, we coupled epidemiological, clinical, genomic and experimental data to provide comprehensive insights into vaccine serotype persistence.

Our data indicate that while vaccine serotypes may persist, there is potential for the selection of less virulent lineages following vaccine introduction. As this would likely translate into reductions in disease, our data challenge current approaches for measuring vaccine impact by demonstrating the importance of exploring beyond serotype when estimating the true value of vaccination. Critically, our data help provide decision-makers assurance of the continued value of PCV introduction despite vaccine serotype persistence.

## Contributors

PS, LKB, SM and CS conceptualised the study and devised the analysis plan. PS, LKB, SM, VBW, JH, SWL, CLP, BDO, RAH, HH, OT, TM, and SB acquired the data. PS, LKB, SM, JH, CLP, BDO, RAH, and CvM performed the data analysis. CvM, CDN and CS devised the statistical analysis plan, CDN provided statistical oversight and PS, LKB and SM applied formal statistical and computational techniques to analyse the experimental data. PS, LKB, SM, CLP, BDO and CS accessed and verified the laboratory data. LB, SM, EMD, CvM, FMR, EKM, TM, PVL, SDB, CS acquired the funding. PS, LKB and SM prepared, created and presented the data visually. PS, LKB, SM and CS wrote the first draft of the manuscript. All authors were involved in data interpretation and review of the final manuscript. All authors had full access to all the data in the study and had final responsibility for the decision to submit for publication.

## Data sharing statement

Deidentified group data can be made available for sharing through application to the corresponding author. This application must include the relevant proposal detailing the intended use of the data and relevant ethical approval for this proposal, and it requires a signed data sharing agreement. Whole genome sequences of capsule loci used in this study were deposited in the GenBank database under the identifiers PX311056-PX311063.

## Acknowledgments

We would like to acknowledge the Ministry of Health in Mongolia and the WHO country office for their support for the vaccine impact program. We thank all study participants and their families. We acknowledge the contributions of staff from Songinokhairkan District Hospital, Sükhbaatar District hospital, Bayanzürkh District Hospital, Chingeltei District Hospital, the Tertiary Maternal and Child Health Hospital, and the National Centre for Communicable Diseases (Mongolia). We thank Jeff Weiser for helpful discussions. From the Murdoch Children’s Research Institute (Australia) we thank the Disease Model Unit for services and support provided for mouse studies. We thank Meghan McKinnon and Cong Ho (Royal Children’s Hospital, Melbourne) for their support of the transmission electron microscopy. We also thank Steve Petrovski (La Trobe University, Bundoora) for the provision of reagents for the BATH assay.

