## Supplementary material for "*Streptococcus pneumoniae* vaccine serotype persistence following 13-valent pneumococcal conjugate vaccine introduction in Mongolia: investigating changes in epidemiology, immunology and virulence"

***Immunological assays***

Enzyme Linked Immunosorbent Assays (ELISAs) were used to measure antibody binding to pneumococci. ELISAs were performed by coating high-binding plates (Maxisorp immune plate, Nunc) with a bacterial culture. Pneumococcal cultures were incubated in Todd Hewitt Broth containing 0·5% yeast extract and grown to an optical density at 600 nm ($OD_{600}$) of 0·85 at 37°C in 5% $CO_2$. Serum and control samples were diluted 1:100 in absorption buffer which is phosphate-buffered saline (PBS) containing 10% (v/v) foetal calf serum (PBS/FCS), cell-wall polysaccharide (CPS; 5 µg/ml) and 22F capsular polysaccharide (5 µg/ml) to remove non-specific antibodies. For this study, we used sera of high antibody concentration previously collected from vaccinated children aged 5-7 years in Fiji, 28 days after immunisation with a PCV13 booster.[1] Plates and serum samples were then incubated overnight at 4°C before the ELISA was performed as previously described.[1] Briefly, plates were blocked with phosphate-buffered saline (PBS/FCS) and incubated at 37°C, 5% $CO_2$ for 1 h. Sera samples were diluted further at 1:1,000, 1:5,000 and 1:10,000 in absorption buffer, added to the wells and incubated (37°C, 5% $CO_2$ for 2 h). This was followed by a wash step with PBS containing 0·05% (v/v) Tween. Horseradish peroxidase-conjugated goat anti-human IgG (Bio-Rad) was added and incubated (37°C, 5% $CO_2$ for 2 h) before adding 3·3',5·5'-tetramehylbenzidine (TMB) substrate solution to develop the reaction. The reaction was stopped with 1 M phosphoric acid. Optical density was measured at 450nm (630nm reference filter) using a microplate reader (BioTek). Serotype-specific IgG concentrations were derived from the 007SP reference standard values and expressed as mean ELISA Units per ml (EU/ml).

***Serotype 6B capsule operon and gene comparison***

To examine genomes for differences in the capsule operon we undertook whole-genome sequencing. Pneumococcal-positive samples serotyped as 6B using microarray were cultured on selective agar and pneumococci were isolated for presumptive identification as described previously.[2] Pneumococci were collected from plates using 1 ml of PBS. DNA was extracted from bacterial supensions using the QIAcube HT with the QIAamp 96 DNA QIAcube HT Kit (Qiagen). Extracted DNA from pneumococcal isolates were sequenced using 150 bp paired-end sequencing on the NovaSeq 6000 (Illumina) platform. Quality control was undertaken using Fastp (galaxy version 0·23·2).[3] Genomes were assembled using Shovill (version 1·1·0),[4] with the following parameters: sequences were subsampled to a depth of 100 with an estimated genome size of 2·1M, a minimum contig length of 300 and a minimum contig coverage of 5 x. Assembly quality was evaluated using Quast (version 5·0·2).[5] We used Snippy (version 4·6·0)[6] to compare single nucleotide polymorphisms (SNPs) in the capsule loci of serotype 6B and Clustal Omega (version 1·2·2) to calculate percent identity matrix and align sequences.

***RNA extraction***

Extraction of RNA from pneumococci was performed using the RNeasy Mini Kit (Qiagen) according to the manufacturer's instructions with the incorporation of an additional lysis step as previously published.[7] In brief, pneumococcal cultures were incubated in Todd Hewitt Broth containing 0·5% yeast extract and grown to $OD_{600}$ of 0·2 before storage in RNAprotect Bacteria Reagent (Qiagen). Bacterial pellets were resuspended in 200 µl of lysis buffer containing 600 µg/ml lysozyme (Sigma-Aldrich), 15 µg/ml Mutanolysin (Sigma-Aldrich), 1U/µg SUPERase In RNase Inhibitor (Ambion), 400 µg/ml Proteinase K (Qiagen) in TE buffer, and incubated at 37 °C for 15 min. The remainder of the extraction was performed as per the RNeasy Mini Kit manufacturer's instructions. DNase treatment was performed by adding final concentration of 150 Kunitz U/ml DNase I in buffer RDD (Qiagen) and incubating for 15 min prior to washing and elution in 30 µl RNase-free water. The integrity and concentration of RNA was measured using Tape Station. RNA was stored at -80 °C until use.

***Quantitative reverse transcriptase polymerase chain reaction (qRT-PCR)***

To measure levels of capsule gene expression from extracted RNA, qRT-PCR was used. Extracted RNA underwent a second DNAse treatment step by adding 10 µl of treatment containing 100 ng/µl RNA, 10 X buffer (Promega), 1 U/µl RQ1 DNase (Promega) in RNase-free water. DNase-treated RNA was used for first strand cDNA synthesis using the iScript cDNA synthesis kit (BioRad) as described previously.[7] Briefly, for each sample, 100 ng RNA was added and incubated in a thermocycler under the following conditions: 5 min at 25°C, 30 min at 42°C and 5 min at 85°C.

cDNA was used for qRT-PCR and run in duplicate reactions using the GoTaq® qPCR Master Mix (Promega) as per the manufacturer's instructions. Each reaction contained 2 µl of cDNA (or nuclease free water for the no template control) and 0·2 µM of forward and reverse primers (Supplementary Table S3). Reactions were run on the Mx3005P qPCR system using the following protocol: 2 min at 95 °C followed by 40 cycles of 15 sec at 95 °C and 1 min at 60 °C. Non-specific amplification was assessed by performing a dissociation curve at the end of the run (1 min at 95 °C, 30 sec at 60 °C and then 30 sec at 95 °C). Analysis of each gene for changes in expression of the post-vaccine lineage, relative to the pre-vaccine lineage was performed using the $2^{-\Delta\Delta Ct}$ relative quantification method. Data were normalised to the *gyrA* gene.

***Transmission Electron Microscopy***

Overnight bacterial growth on HBA plates (37ºC, 5% $CO_2$) was suspended in 1 ml PBS to an $OD_{600}$ of 0·4. Bacteria was pelleted at 600 RCF for 5 min and supernatant discarded before resuspension in 1 ml of fixative comprised of 75 mM L-lysine (Sigma-Aldrich), in 0·075% (w/v) ruthenium red (Sigma-Aldrich), 2% (v/v) paraformaldehyde (w/v) (Sigma-Aldrich), and 2·5% (w/v) glutaraldehyde in PBS for 20 min at room temperature. The fixative was removed, and pellets were resuspended in fixative without lysine for 2 h at room temperature. Bacterial pellets were washed three times in PBS and fixed in 1% $OsO_4$ w/v for 2 h. Followed again by an additional three PBS wash cycles, pellets were dehydrated in graded acetone and embedded in Epon-Araldite.

**Supplementary Table S1. Individual vaccine status of children from Mongolia hospitalised with pneumonia colonised with vaccine serotypes where there was evidence of lineage replacement following PCV introduction.**

| | Serotype 6A | | | Serotype 6B | | | Serotype 14 | | | Serotype 19F | | | Serotype 23F | | |
|---|---|---|---|---|---|---|---|---|---|---|---|---|---|---|---|
| | **GPSC16 n/N (%)** | **GPSC853 n/N (%)** | **p-value[d]** | **GPSC23 n/N (%)** | **GPSC6 n/N (%)** | **p-value[d]** | **GPSC279 n/N (%)** | **GPSC9 n/N (%)** | **p-value[d]** | **GPSC1-2 n/N (%)** | **GPSC1-3 n/N (%)** | **p-value[d]** | **GPSC16 n/N (%)** | **n/N (%)** | **p-value[d]** |
| Pre-PCV13 period[a] | 40/76 (53) | 42/89 (47) | 0.53 | 35/52 (67) | 9/54 (17) | <0.01 | 56/94 (60) | 2/13 (15) | 0.01 | 36/53 (68) | 19/43 (44) | 0.02 | 24/33 (73) | 16/59 (27) | <0.01 |
| Undervaccinated[b] | 22/76 (29) | 26/89 (29) | 1.00 | 12/52 (23) | 19/54 (35) | 0.20 | 22/94 (24) | 5/13 (39) | 0.31 | 7/53 (13) | 8/43 (19) | 0.58 | 5/33 (15) | 20/59 (34) | 0.09 |
| Vaccinated[c] | 14/76 (18) | 21/89 (24) | 0.45 | 5/52 (10) | 26/54 (48) | <0.01 | 16/94 (17) | 6/13 (46) | 0.03 | 10/53 (19) | 16/43 (37) | 0.06 | 4/33 (12) | 23/59 (39) | 0.01 |

[a]Pre-PCV13 period includes samples obtained from children in 2015 and 2016. [b]For children admitted to hospital after vaccine introduction (after May 2016), 'Undervaccinated' refers to children <12 months of age who received 0-1 doses of PCV13 or children ≥12 months of age with no dose PCV, whereas [c]vaccinated is defined as receipt of ≥2 doses of PCV13 at <12 months of age or ≥1 dose of PCV13 administered at ≥12 months of age. . [d]Data were anaysed by proportion of children carrying each lineage by vaccine status, with each vaccine status analysed separately using Fisher's exact test.Vaccination status was not available for all children carrying each lineage.

**Supplementary Table S2. Antimicrobial resistance (AMR) genes detected by DNA microarray in vaccine serotypes where there was evidence of lineage replacement following PCV introduction[a]**

| | Serotype 6A | | | Serotype 6B | | | Serotype 14 | | | Serotype 19F[b] | | | Serotype 23F | | |
|---|---|---|---|---|---|---|---|---|---|---|---|---|---|---|---|
| **AMR gene** | **GPSC16 n/N (%)** | **GPSC853 n/N (%)** | **p-value[c]** | **GPSC23 n/N (%)** | **GPSC6 n/N (%)** | **p-value[c]** | **GPSC279 n/N (%)** | **GPSC9 n/N (%)** | **p-value[c]** | **GPSC1-2 n/N (%)** | **GPSC1-3 n/N (%)** | **p-value[c]** | **GPSC16 n/N (%)** | **GPSC101 n/N (%)** | **p-value[c]** |
| *ermB* | 45/58 (78) | 69/69 (100) | 0.0001 | 38/38 (100) | 44/44 (100) | 1.00 | 65/74 (88) | 0/12 (0) | <0.0001 | 35/35 (100) | 28/29 (97) | 1.00 | 22/22 (100) | 3/40 (8) | <0.0001 |
| *cat* | 56/58 (97) | 4/69 (6) | <0.0001 | 18/38 (34) | 4/44 (9) | <0.0001 | 19/74 (26) | 0/12 (0) | 0.06 | 1/35 (3) | 1/29 (3) | 1.00 | 21/22 (95) | 3/40 (8) | <0.0001 |
| *tetM* | 58/58 (100) | 69/69 (100) | 1.00 | 38/38 (100) | 44/44 (100) | 1.00 | 65/74 (88) | 11/12 (92) | 1.00 | 35/35 (100) | 29/29 (100) | 1.00 | 22/22 (100) | 39/40 (98) | 1.00 |
| *tetO* | 0/58 (0) | 0/69 (0) | 1.00 | 1/38 (3) | 0/44 (0) | 1.00 | 0/74 (0) | 0/12 (0) | 1.00 | 0/35 (0) | 0/29 (0) | 1.00 | 0/22 (0) | 0/40 (0) | 1.00 |
| *aphA3* | 0/58 (0) | 0/69 (0) | 1.00 | 1/38 (3) | 2/44 (5) | 1.00 | 1/74 (1) | 0/12 (0) | 1.00 | 1/35 (3) | 0/29 (0) | 1.00 | 0/22 (0) | 0/40 (0) | 1.00 |
| *sat4* | 0/58 (0) | 0/69 (0) | 1.00 | 1/38 (3) | 2/44 (5) | 1.00 | 1/74 (1) | 0/12 (0) | 1.00 | 1/35 (3) | 0/29 (0) | 1.00 | 0/22 (0) | 0/40 (0) | 1.00 |
| *mefA* | 58/58 (100) | 2/69 (3) | <0.0001 | 0/38 (0) | 0/44 (0) | 1.00 | 2/74 (3) | 0/12 (0) | 1.00 | 35/35 (100) | 27/29 (93) | 1.00 | 1/22 (5) | 19/40 (48) | 0.001 |
| Any AMR gene | 58/58 (100) | 69/69 (100) | 1.00 | 38/38 (100) | 44/44 (100) | 1.00 | 67/74 (91) | 11/12 (92) | 1.00 | 35/35 (100) | 29/29 (100) | 1.00 | 22/22 (100) | 40/40 (100) | 1.00 |
| ≥3 AMR genes[d] | 57/58 (98) | 5/69 (7) | <0.0001 | 18/38 (47) | 4/44 (9) | <0.0001 | 20/74 (27) | 0/12 (0) | 0.06 | 35/35 (100) | 27/29 (93) | 0.20 | 21/22 (95) | 2/40 (5) | <0.0001 |

[a]Only samples that contained a single pneumococcal serotype with no other species identified were included in the analysis. [b] Lineage sub-clustering within serotype 19F was based on dendrograms and samples with low intensity QC flags were excluded from analysis. [c]Data were compared using Fisher's exact test. [d]Multi-drug resistance is defined as ≥3 antimicrobial resistance genes detected. For each serotype, the lineage that was common pre-PCV is listed first, followed by the lineage that was most common post-PCV.

**Supplementary Table S3. List of primers used in this study for qRT-PCR analysis.**

| Pneumococcal gene | Amplicon size (bp) | Primer | Sequence (5'→ 3') | Reference |
|---|---|---|---|---|
| *gyrA* | 98 | Forward | CAATATGCTCGCTATCCAA | 8 |
| | | Reverse | GACGAACAACCACTTCTT | 8 |
| *wzg* | 99 | Forward | TCATTGAATCGGAGTATCC | This study |
| | | Reverse | GATTCTTAGACGTCTTAGG | This study |
| *wzy* | 83 | Forward | TAAAGTGGAGGGAATTTCG | This study |
| | | Reverse | GATTTAGAATCAGAAGAATGC | This study |
| *rmlB* | 120 | Forward | CTATCGTTCATTATGCAGC | This study |
| | | Reverse | TATCATACTTACGAGCAGC | This study |

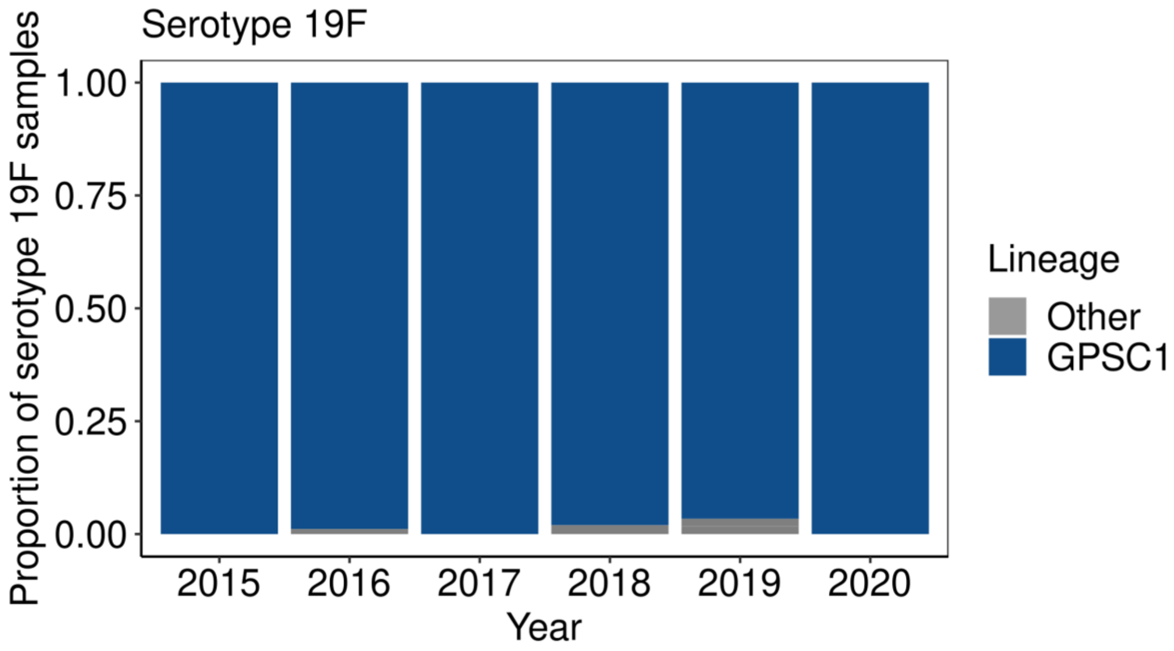

**Supplementary Figure S1. Lineage composition of vaccine serotype 19F circulating in Mongolia between 2015 and 2020.** Genetic lineage (GPSC, Global Pneumococcal Sequence Cluster) was inferred from pneumococcal-positive samples using DNA microarray.[9] Bars are coloured by lineage.

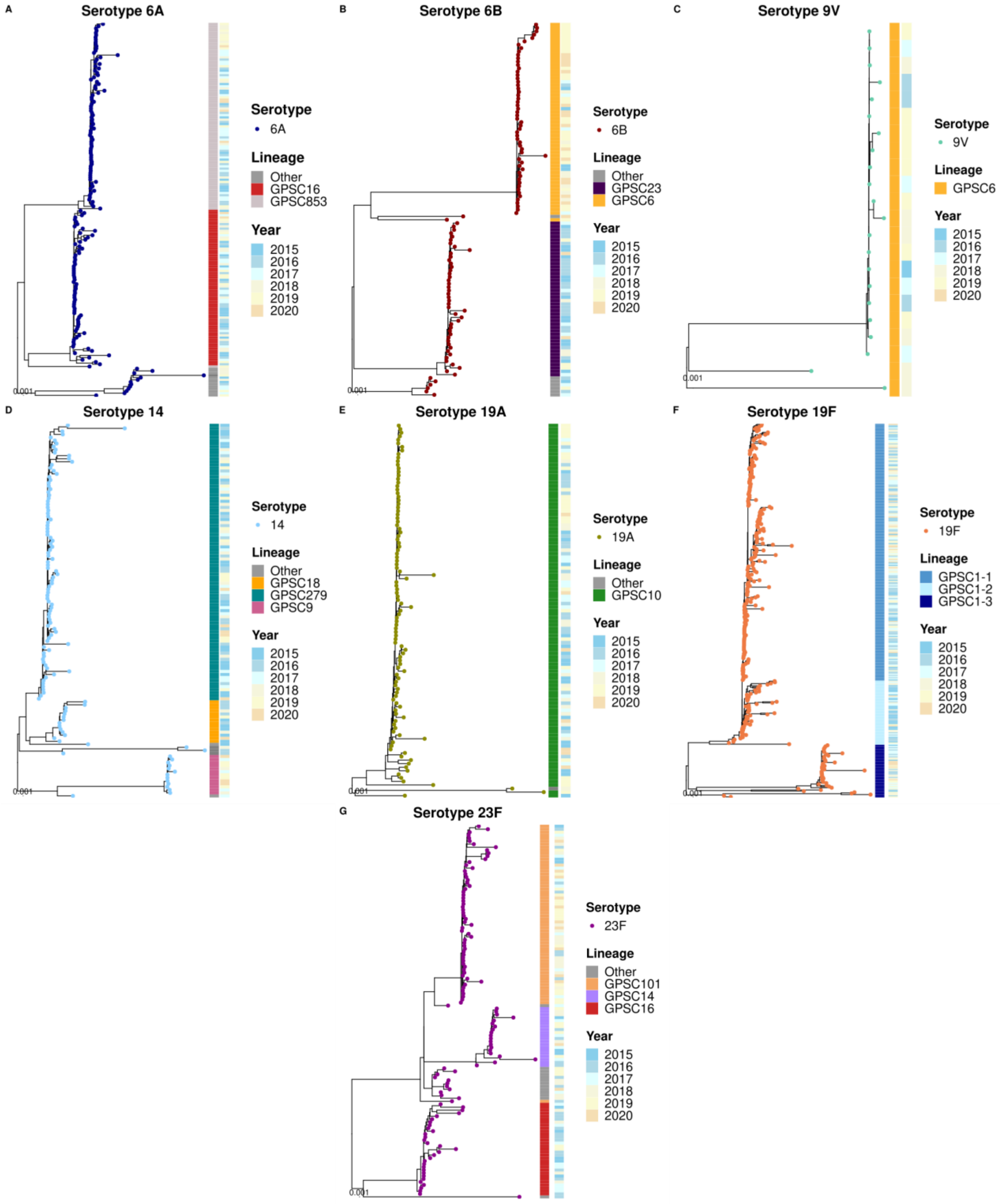

**Supplementary Figure S2. Dendrogram of pneumococcal vaccine serotypes circulating in Mongolian children hospitalised with pneumonia from 2015 to 2020.** The dendrograms are based on array-CGH analysis of the genome backbone component of microarray data,[9] scale bar represents the divergence between samples. Dendrograms are overlayed with serotype (tip points), lineage (based on Global Pneumococcal Sequence Cluster or sub-cluster; first bar) and year (second bar).

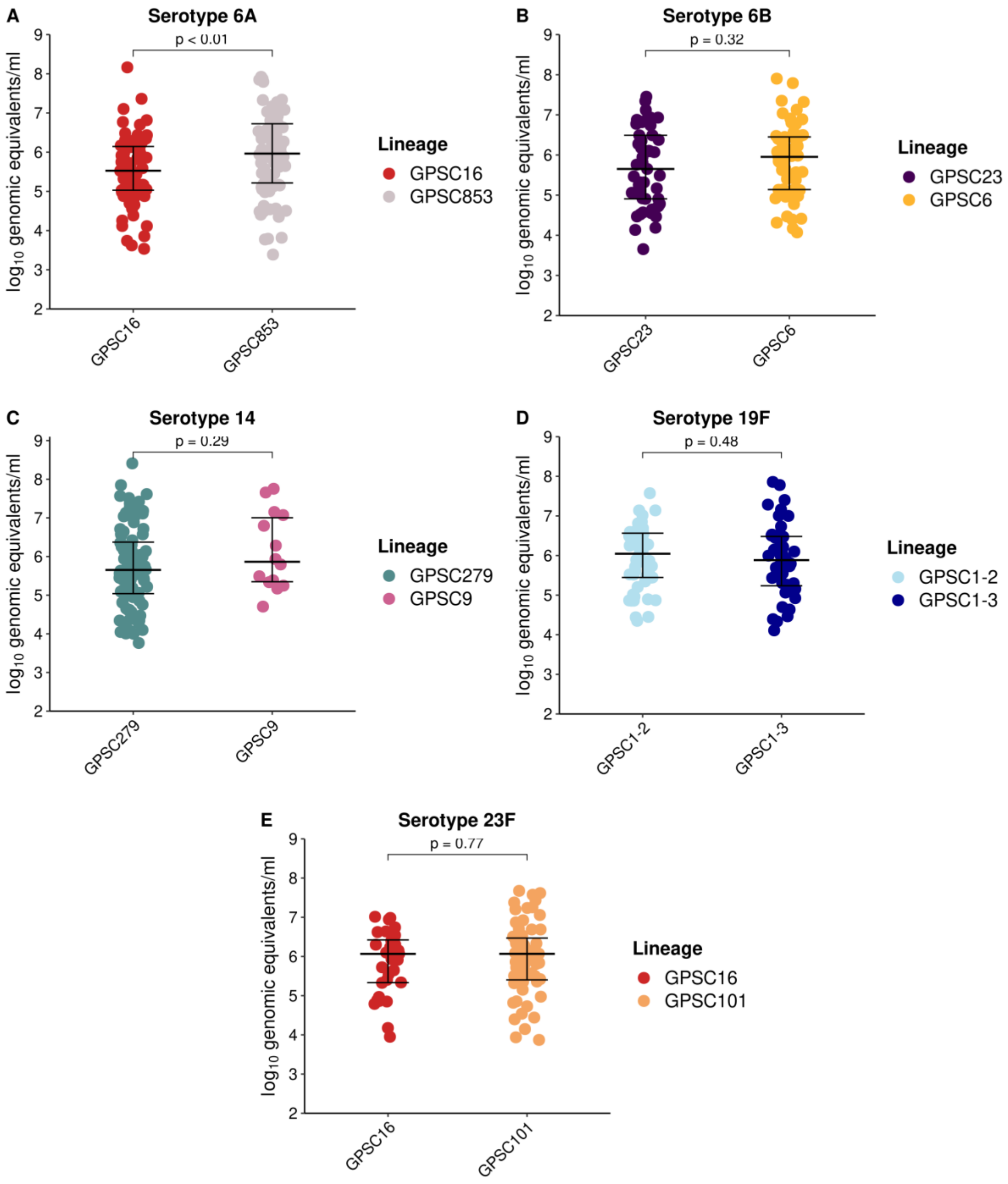

**Supplementary Figure S3. Nasopharyngeal density of the most common pre- and post-vaccine lineages of serotype 6A, 6B, 14, 19F and 23F in children with pneumonia.** Nasopharyngeal density expressed as $\log_{10}$ genome equivalents/ml. Each individual dot point represents one sample and is coloured by lineage (GPSC and/or sub-cluster). Errors bars represent the median and interquartile range. Differences in lineage density within a serotype were only found for serotype 6A ($p<0.05$) using the Mann-Whitney test (p-values shown for each comparison). GPSC, Global Pneumococcal Sequence Cluster.

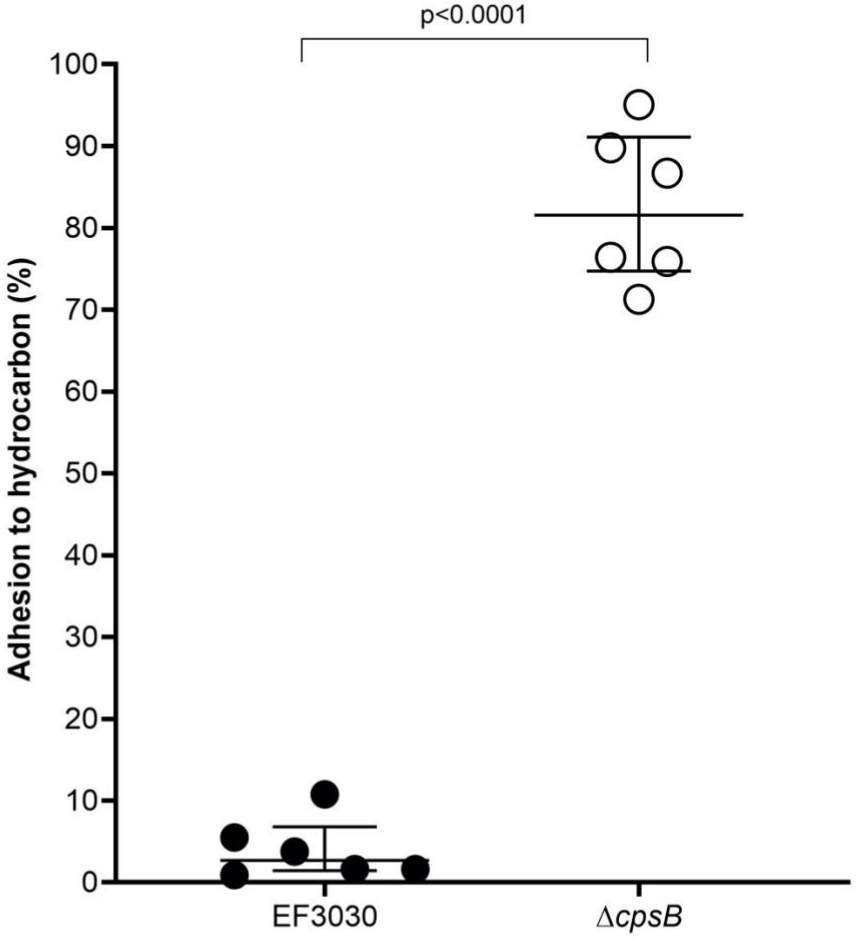

**Supplementary Figure S4. Adhesion to hydrocarbon of serotype 19F parental strain (EF3030) and capsule mutant (Δ*cpsB*) using Bacterial Adhesion to Hydrocarbon (BATH) assay.** The BATH assay measured binding of serotype 19F parent and mutant to hexadecane hydrocarbon and was expressed as a percentage of the inoculum. Each individual dot represents data from an independent experiment. Error bars represent the median and interquartile range. Differences were assessed using a paired t-test.

```
PMP1610      ATGCAAGAAAAATGGTGGCACAATGCCGTAGTCTATCAAGTCTATCCAAAGAGTTTTATG   60
PMP1666      ATGCAAGAAAAATGGTGGCACAATGCCGTAGTCTATCAAGTCTATCCAAAGAGTTTTATG   60
PMP1665      ATGCAAGAAAAATGGTGGCACAATGCCGTAGTCTATCAAGTCTATCCAAAGAGTTTTATG   60
PMP1667      ATGCAAGAAAAATGGTGGCACAATGCCGTAGTCTATCAAGTCTATCCAAAGAGTTTTATG   60
PMP1663      ATGCAAGAAAAATGGTGGCACAATGCCGTAGTCTATCAAGTCTATCCAAAGAGTTTTATG   60
PMP1664      ATGCAAGAAAAATGGTGGCACAATGCCGTAGTCTATCAAGTCTATCCAAAGAGTTTTATG   60
PMP1662      ATGCAAGAAAAATGGTGGCACAATGCCGTAGTCTATCAAGTCTATCCAAAGAGTTTTATG   60
PMP1611      ATGCAAGAAAAATGGTGGCACAATGCCGTAGTCTATCAAGTCTATCCAAAGAGTTTTATG   60
             ************************************************************

PMP1610      GATAGTAATGGAGATGGAGTTGGTGATTTGCCAGGTATTACCAGTAAGTTGGACTATCTA   120
PMP1666      GATAGTAATGGAGATGGAGTTGGTGATTTGCCAGGTATTACCAGTAAGTTGGACTATCTA   120
PMP1665      GATAGTAATGGAGATGGAGTTGGTGATTTGCCAGGTATTACCAGTAAGTTGGACTATCTA   120
PMP1667      GATAGTAATGGAGATGGAGTTGGTGATTTGCCAGGTATTACCAGTAAGTTGGACTATCTA   120
PMP1663      GATAGTAATGGAGATGGAGTTGGTGATTTGCCAGGTATTACCAGTAAGTTGGACTATCTA   120
PMP1664      GATAGTAATGGAGATGGAGTTGGTGATTTGCCAGGTATTACCAGTAAGTTGGACTATCTA   120
PMP1662      GATAGTAATGGAGATGGAGTTGGTGATTTGCCAGGTATTACCAGTAAGTTGGACTATCTA   120
PMP1611      GATAGTAATGGAGATGGAGTTGGTGATTTGCCAGGTATTACCAGTAAGTTGGACTATCTA   120
             ************************************************************

PMP1610      GCCAAGCTAGGAATCACAGCAATTTGGCTTTCTCCCGTTTATGACAGCCCTATGGATGAT   180
PMP1666      GCCAAGCTAGGAATCACAGCAATTTGGCTTTCTCCCGTTTATGACAGCCCTATGGATGAT   180
PMP1665      GCCAAGCTAGGAATCACAGCAATTTGGCTTTCTCCCGTTTATGACAGCCCTATGGATGAT   180
PMP1667      GCCAAGCTAGGAATCACAGCAATTTGGCTTTCTCCCGTTTATGACAGCCCTATGGATGAT   180
PMP1663      GCCAAGCTAGGAATCACAGCAATTTGGCTTTCTCCCGTTTATGACAGCCCTATGGATGAT   180
PMP1664      GCCAAGCTAGGAATCACAGCAATTTGGCTTTCTCCCGTTTATGACAGCCCTATGGATGAT   180
PMP1662      GCCAAGCTAGGAATCACAGCAATTTGGCTTTCTCCCGTTTATGACAGCCCTATGGATGAT   180
PMP1611      GCCAAGCTAGGAATCACAGCAATTTGGCTTTCTCCCGTTTATGACAGCCCTATGGATGAT   180
             ************************************************************

PMP1610      AATGGCTATGATATTGCTGATTATCAAGCGATTGCGGCTATTTTTGGAACCATGGAGGAC   240
PMP1666      AATGGCTATGATATTGCTGATTATCAAGCGATTGCGGCTATTTTTGGAACCATGGAGGAC   240
PMP1665      AATGGCTATGATATTGCTGATTATCAAGCGATTGCGGCTATTTTTGGAACCATGGAGGAC   240
PMP1667      AATGGCTATGATATTGCTGATTATCAAGCGATTGCGGCTATTTTTGGAACCATGGAGGAC   240
PMP1663      AATGGCTATGATATTGCTGATTATCAAGCGATTGCGGCTATTTTTGGAACCATGGAGGAC   240
PMP1664      AATGGCTATGATATTGCTGATTATCAAGCGATTGCGGCTATTTTTGGAACCATGGAGGAC   240
```

```
PMP1662      AATGGCTATGATATTGCTGATTATCAAGCGATTGCGGCTATTTTTGGAACCATGGAGGAC   240
PMP1611      AATGGCTATGATATTGCTGATTATCAAGCGATTGCGGCTATTTTTGGAACCATGGAGGAC   240
             ************************************************************

PMP1610      ATGGATCAGCTGATTGCAGAAGCTAAGAAGCGTGACATTCGTATCATCATGGACTTGGTG   300
PMP1666      ATGGATCAGCTGATTGCAGAAGCTAAGAAGCGTGACATTCGTATCATCATGGACTTGGTG   300
PMP1665      ATGGATCAGCTGATTGCAGAAGCTAAGAAGCGTGACATTCGTATCATCATGGACTTGGTG   300
PMP1667      ATGGATCAGCTGATTGCAGAAGCTAAGAAGCGTGACATTCGTATCATCATGGACTTGGTG   300
PMP1663      ATGGATCAGCTGATTGCAGAAGCTAAGAAGCGTGACATTCGTATCATCATGGACTTGGTG   300
PMP1664      ATGGATCAGCTGATTGCAGAAGCTAAGAAGCGTGACATTCGTATCATCATGGACTTGGTG   300
PMP1662      ATGGATCAGCTGATTGCAGAAGCTAAGAAGCGTGACATTCGTATCATCATGGACTTGGTG   300
PMP1611      ATGGATCAGCTGATTGCAGAAGCTAAGAAGCGTGACATTCGTATCATCATGGACTTGGTG   300
             ************************************************************

PMP1610      GTTAATCATACCTCAGATGAACATGCTTGGTTTGTCGAAGCCTGTGAAAATACTGACAGC   360
PMP1666      GTTAATCATACCTCAGATGAACATGCTTGGTTTGTCGAAGCCTGTGAAAATACTGACAGC   360
PMP1665      GTTAATCATACCTCAGATGAACATGCTTGGTTTGTCGAAGCCTGTGAAAATACTGACAGC   360
PMP1667      GTTAATCATACCTCAGATGAACATGCTTGGTTTGTCGAAGCCTGTGAAAATACTGACAGC   360
PMP1663      GTTAATCATACCTCAGATGAACATGCTTGGTTTGTCGAAGCCTGTGAAAATACTGACAGC   360
PMP1664      GTTAATCATACCTCAGATGAACATGCTTGGTTTGTCGAAGCCTGTGAAAATACTGACAGC   360
PMP1662      GTTAATCATACCTCAGATGAACATGCTTGGTTTGTCGAAGCCTGTGAAAATACTGACAGC   360
PMP1611      GTTAATCATACCTCAGATGAACATGCTTGGTTTGTCGAAGCCTGTGAAAATACTGACAGC   360
             ************************************************************

PMP1610      CCTGAGCGAGACTACTATATCTGGCGCGATGAACCCAATGACCTAGATTCTATCTTTAGT   420
PMP1666      CCTGAGCGAGACTACTATATCTGGCGCGATGAACCCAATGACCTAGATTCTATCTTTAGT   420
PMP1665      CCTGAGCGAGACTACTATATCTGGCGCGATGAACCCAATGACCTAGATTCTATCTTTAGT   420
PMP1667      CCTGAGCGAGACTACTATATCTGGCGCGATGAACCCAATGACCTAGATTCTATCTTTAGT   420
PMP1663      CCTGAGCGAGACTACTATATCTGGCGCGATGAACCCAATGACCTAGATTCTATCTTTAGT   420
PMP1664      CCTGAGCGAGACTACTATATCTGGCGCGATGAACCCAATGACCTAGATTCTATCTTTAGT   420
PMP1662      CCTGAGCGAGACTACTATATCTGGCGCGATGAACCCAATGACCTAGATTCTATCTTTAGT   420
PMP1611      CCTGAGCGAGACTACTATATCTGGCGCGATGAACCCAATGACCTAGATTCTATCTTTAGT   420
             ************************************************************

PMP1610      GGGTCTGCTTGGGAATACGATGAAAAGTCAGGTCAATACTATCTCCACTTTTTCAGCAAA   480
PMP1666      GGGTCTGCTTGGGAATACGATGAAAAGTCAGGTCAATACTATCTCCACTTTTTCAGCAAA   480
```

```
PMP1665      GGGTCTGCTTGGGAATACGATGAAAAGTCAGGTCAATACTATCTCCACTTTTTCAGCAAA   480
PMP1667      GGGTCTGCTTGGGAATACGATGAAAAGTCAGGTCAATACTATCTCCACTTTTTCAGCAAA   480
PMP1663      GGGTCTGCTTGGGAATACGATGAAAAGTCAGGTCAATACTATCTCCACTTTTTCAGCAAA   480
PMP1664      GGGTCTGCTTGGGAATACGATGAAAAGTCAGGTCAATACTATCTCCACTTTTTCAGCAAA   480
PMP1662      GGGTCTGCTTGGGAATACGATGAAAAGTCAGGTCAATACTATCTCCACTTTTTCAGCAAA   480
PMP1611      GGGTCTGCTTGGGAATACGATGAAAAGTCAGGTCAATACTATCTCCACTTTTTCAGCAAA   480
             ************************************************************

PMP1610      AAACAGCCGGATCTCAACTGGGAAAATGAAAAACTTCGCCAGAAAATTTATGAGATGATG   540
PMP1666      AAACAGCCGGATCTCAACTGGGAAAATGAAAAACTTCGCCAGAAAATTTATGAGATGATG   540
PMP1665      AAACAGCCGGATCTCAACTGGGAAAATGAAAAACTTCGCCAGAAAATTTATGAGATGATG   540
PMP1667      AAACAGCCGGATCTCAACTGGGAAAATGAAAAACTTCGCCAGAAAATTTATGAGATGATG   540
PMP1663      AAACAGCCGGATCTCAACTGGGAAAATGAAAAACTTCGCCAGAAAATTTATGAGATGATG   540
PMP1664      AAACAGCCGGATCTCAACTGGGAAAATGAAAAACTTCGCCAGAAAATTTATGAGATGATG   540
PMP1662      AAACAGCCGGATCTCAACTGGGAAAATGAAAAACTTCGCCAGAAAATTTATGAGATGATG   540
PMP1611      AAACAGCCGGATCTCAACTGGGAAAATGAAAAACTTCGCCAGAAAATTTATGAGATGATG   540
             ************************************************************

PMP1610      AACTTCTGGATTGATAAAGGTATTGGTGGTTTCCGTATGGATGTTATTGACATGATTGGC   600
PMP1666      AACTTCTGGATTGATAAAGGTATTGGTGGTTTCCGTATGGATGTTATTGACATGATTGGC   600
PMP1665      AACTTCTGGATTGATAAAGGTATTGGTGGTTTCCGTATGGATGTTATTGACATGATTGGC   600
PMP1667      AACTTCTGGATTGATAAAGGTATTGGTGGTTTCCGTATGGATGTTATTGACATGATTGGC   600
PMP1663      AACTTCTGGATTGATAAAGGTATTGGTGGTTTCCGTATGGATGTTATTGACATGATTGGC   600
PMP1664      AACTTCTGGATTGATAAAGGTATTGGTGGTTTCCGTATGGATGTTATTGACATGATTGGC   600
PMP1662      AACTTCTGGATTGATAAAGGTATTGGTGGTTTCCGTATGGATGTTATTGACATGATTGGC   600
PMP1611      AACTTCTGGATTGATAAAGGTATTGGTGGTTTCCGTATGGATGTTATTGACATGATTGGC   600
             ************************************************************

PMP1610      AAAATTCCTGACGAGAAGGTAGTCAATAATGGTCCTATGCTCCATCCCTATCTCAAGGAA   660
PMP1666      AAAATTCCTGACGAGAAGGTAGTCAATAATGGTCCTATGCTCCATCCCTATCTCAAGGAA   660
PMP1665      AAAATTCCTGACGAGAAGGTAGTCAATAATGGTCCTATGCTCCATCCCTATCTCAAGGAA   660
PMP1667      AAAATTCCTGACGAGAAGGTAGTCAATAATGGTCCTATGCTCCATCCCTATCTCAAGGAA   660
PMP1663      AAAATTCCTGACGAGAAGGTAGTCAATAATGGTCCTATGCTCCATCCCTATCTCAAGGAA   660
PMP1664      AAAATTCCTGACGAGAAGGTAGTCAATAATGGTCCTATGCTCCATCCCTATCTCAAGGAA   660
PMP1662      AAAATTCCTGACGAGAAGGTAGTCAATAATGGTCCTATGCTCCATCCCTATCTCAAGGAA   660
PMP1611      AAAATTCCTGACGAGAAGGTAGTCAATAATGGTCCTATGCTCCATCCCTATCTCAAGGAA   660
```

```
            ************************************************************

PMP1610     ATGAATCAGGCGACCTTTGGAGATAAGGATCTCTTGACAGTAGGGGAGACTTGGGGAGCA   720
PMP1666     ATGAATCAGGCGACCTTTGGAGATAAGGATCTCTTGACAGTAGGGGAGACTTGGGGAGCA   720
PMP1665     ATGAATCAGGCGACCTTTGGAGATAAGGATCTCTTGACAGTAGGGGAGACTTGGGGAGCA   720
PMP1667     ATGAATCAGGCGACCTTTGGAGATAAGGATCTCTTGACAGTAGGGGAGACTTGGGGAGCA   720
PMP1663     ATGAATCAGGCGACCTTTGGAGATAAGGATCTCTTGACAGTAGGGGAGACTTGGGGAGCA   720
PMP1664     ATGAATCAGGCGACCTTTGGAGATAAGGATCTCTTGACAGTAGGGGAGACTTGGGGAGCA   720
PMP1662     ATGAATCAGGCGACCTTTGGAGATAAGGATCTCTTGACAGTAGGGGAGACTTGGGGAGCA   720
PMP1611     ATGAATCAGGCGACCTTTGGAGATAAGGATCTCTTGACAGTAGGGGAGACTTGGGGAGCA   720
            ************************************************************

PMP1610     ACGCCAGAGATTGCCAAGCTCTACTCTGATCCAAAGGGGCAAGAATTGTCTATGGTCTTC   780
PMP1666     ACGCCAGAGATTGCCAAGCTCTACTCTGATCCAAAGGGGCAAGAATTGTCTATGGTCTTC   780
PMP1665     ACGCCAGAGATTGCCAAGCTCTACTCTGATCCAAAGGGGCAAGAATTGTCTATGGTCTTC   780
PMP1667     ACGCCAGAGATTGCCAAGCTCTACTCTGATCCAAAGGGGCAAGAATTGTCTATGGTCTTC   780
PMP1663     ACGCCAGAGATTGCCAAGCTCTACTCTGATCCAAAGGGGCAAGAATTGTCTATGGTCTTC   780
PMP1664     ACGCCAGAGATTGCCAAGCTCTACTCTGATCCAAAGGGGCAAGAATTGTCTATGGTCTTC   780
PMP1662     ACGCCAGAGATTGCCAAGCTCTACTCTGATCCAAAGGGGCAAGAATTGTCTATGGTCTTC   780
PMP1611     ACGCCAGAGATTGCCAAGCTCTACTCTGATCCAAAGGGGCAAGAATTGTCTATGGTCTTC   780
            ************************************************************

PMP1610     CAGTTTGAACATATCGGTCTTCAGTATCAGGAAGGTCAGCCTAAATGGCACTATCAAAAA   840
PMP1666     CAGTTTGAACATATCGGTCTTCAGTATCAGGAAGGTCAGCCTAAATGGCACTATCAAAAA   840
PMP1665     CAGTTTGAACATATCGGTCTTCAGTATCAGGAAGGTCAGCCTAAATGGCACTATCAAAAA   840
PMP1667     CAGTTTGAACATATCGGTCTTCAGTATCAGGAAGGTCAGCCTAAATGGCACTATCAAAAA   840
PMP1663     CAGTTTGAACATATCGGTCTTCAGTATCAGGAAGGTCAGCCTAAATGGCACTATCAAAAA   840
PMP1664     CAGTTTGAACATATCGGTCTTCAGTATCAGGAAGGTCAGCCTAAATGGCACTATCAAAAA   840
PMP1662     CAGTTTGAACATATCGGTCTTCAGTATCAGGAAGGTCAGCCTAAATGGCACTATCAAAAA   840
PMP1611     CAGTTTGAACATATCGGTCTTCAGTATCAGGAAGGTCAGCCTAAATGGCACTATCAAAAA   840
            ************************************************************

PMP1610     GAGCTGAATATCGCTAAGTTAAAAGAAATCTTCAACAAATGGCAGACAGAGTTAGGAGTT   900
PMP1666     GAGCTGAATATCGCTAAGTTAAAAGAAATCTTCAACAAATGGCAGACAGAGTTAGGAGTT   900
PMP1665     GAGCTGAATATCGCTAAGTTAAAAGAAATCTTCAACAAATGGCAGACAGAGTTAGGAGTT   900
PMP1667     GAGCTGAATATCGCTAAGTTAAAAGAAATCTTCAACAAATGGCAGACAGAGTTAGGAGTT   900
```

PMP1663 GAGCTGAATATCGCTAAGTTAAAAGAAATCTTCAACAAATGGCAGACAGAGTTAGGAGTT 900
PMP1664 GAGCTGAATATCGCTAAGTTAAAAGAAATCTTCAACAAATGGCAGACAGAGTTAGGAGTT 900
PMP1662 GAGCTGAATATCGCTAAGTTAAAAGAAATCTTCAACAAATGGCAGACAGAGTTAGGAGTT 900
PMP1611 GAGCTGAATATCGCTAAGTTAAAAGAAATCTTCAACAAATGGCAGACAGAGTTAGGAGTT 900
************************************************************

PMP1610 GAGGACGGCTGGAATTCCCTCTTCTGGAACAACCATGACCTCCCTCGTATTGTCTCAATC 960
PMP1666 GAGGACGGCTGGAATTCCCTCTTCTGGAACAACCATGACCTCCCTCGTATTGTCTCAATC 960
PMP1665 GAGGACGGCTGGAATTCCCTCTTCTGGAACAACCATGACCTCCCTCGTATTGTCTCAATC 960
PMP1667 GAGGACGGCTGGAATTCCCTCTTCTGGAACAACCATGACCTCCCTCGTATTGTCTCAATC 960
PMP1663 GAGGACGGCTGGAATTCCCTCTTCTGGAACAACCATGACCTCCCTCGTATTGTCTCAATC 960
PMP1664 GAGGACGGCTGGAATTCCCTCTTCTGGAACAACCATGACCTCCCTCGTATTGTCTCAATC 960
PMP1662 GAGGACGGCTGGAATTCCCTCTTCTGGAACAACCATGACCTCCCTCGTATTGTCTCAATC 960
PMP1611 GAGGACGGCTGGAATTCCCTCTTCTGGAACAACCATGACCTCCCTCGTATTGTCTCAATC 960
************************************************************

PMP1610 TGGGGAAATGACCAAGAATACCGCGAAAAATCTGCCAAAGCCTTTGCAATCTTGCTTCAT 1020
PMP1666 TGGGGAAATGACCAAGAATACCGCGAAAAATCTGCCAAAGCCTTTGCAATCTTGCTTCAT 1020
PMP1665 TGGGGAAATGACCAAGAATACCGCGAAAAATCTGCCAAAGCCTTTGCAATCTTGCTTCAT 1020
PMP1667 TGGGGAAATGACCAAGAATACCGCGAAAAATCTGCCAAAGCCTTTGCAATCTTGCTTCAT 1020
PMP1663 TGGGGAAATGACCAAGAATACCGCGAAAAATCTGCCAAAGCCTTTGCAATCTTGCTTCAT 1020
PMP1664 TGGGGAAATGACCAAGAATACCGCGAAAAATCTGCCAAAGCCTTTGCAATCTTGCTTCAT 1020
PMP1662 TGGGGAAATGACCAAGAATACCGCGAAAAATCTGCCAAAGCCTTTGCAATCTTGCTTCAT 1020
PMP1611 TGGGGAAATGACCAAGAATACCGCGAAAAATCTGCCAAAGCCTTTGCAATCTTGCTTCAT 1020
************************************************************

PMP1610 CTTATGAGAGGAACTCCTTATATCTACCAAGGTGAGGAGATTGGGATGACCAACTATCCG 1080
PMP1666 CTTATGAGAGGAACTCCTTATATCTACCAAGGTGAGGAGATTGGGATGACCAACTATCCG 1080
PMP1665 CTTATGAGAGGAACTCCTTATATCTACCAAGGTGAGGAGATTGGGATGACCAACTATCCG 1080
PMP1667 CTTATGAGAGGAACTCCTTATATCTACCAAGGTGAGGAGATTGGGATGACCAACTATCCG 1080
PMP1663 CTTATGAGAGGAACTCCTTATATCTACCAAGGTGAGGAGATTGGGATGACCAACTATCCG 1080
PMP1664 CTTATGAGAGGAACTCCTTATATCTACCAAGGTGAGGAGATTGGGATGACCAACTATCCG 1080
PMP1662 CTTATGAGAGGAACTCCTTATATCTACCAAGGTGAGGAGATTGGGATGACCAACTATCCG 1080
PMP1611 CTTATGAGAGGAACTCCTTATATCTACCAAGGTGAGGAGATTGGGATGACCAACTATCCG 1080
************************************************************

```
PMP1610      TTTGAAACACTGGATCAAGTAGAAGATATTGAATCTCTCAACTATGCGCGTGAGGCTCTT   1140
PMP1666      TTTGAAACACTGGATCAAGTAGAAGATATTGAATCTCTCAACTATGCGCGTGAGGCTCTT   1140
PMP1665      TTTGAAACACTGGATCAAGTAGAAGATATTGAATCTCTCAACTATGCGCGTGAGGCTCTT   1140
PMP1667      TTTGAAACACTGGATCAAGTAGAAGATATTGAATCTCTCAACTATGCGCGTGAGGCTCTT   1140
PMP1663      TTTGAAACACTGGATCAAGTAGAAGATATTGAATCTCTCAACTATGCGCGTGAGGCTCTT   1140
PMP1664      TTTGAAACACTGGATCAAGTAGAAGATATTGAATCTCTCAACTATGCGCGTGAGGCTCTT   1140
PMP1662      TTTGAAACACTGGATCAAGTAGAAGATATTGAATCTCTCAACTATGCGCGTGAGGCTCTT   1140
PMP1611      TTTGAAACACTGGATCAAGTAGAAGATATTGAATCTCTCAACTATGCGCGTGAGGCTCTT   1140
             ************************************************************

PMP1610      GAAAAAGGTGTTCCGATTGAAGAAATCATGGACAGTATCCGTGTTATTGGACGTGACAAT   1200
PMP1666      GAAAAAGGTGTTCCGATTGAAGAAATCATGGACAGTATCCGTGTTATTGGACGTGACAAT   1200
PMP1665      GAAAAAGGTGTTCCGATTGAAGAAATCATGGACAGTATCCGTGTTATTGGACGTGACAAT   1200
PMP1667      GAAAAAGGTGTTCCGATTGAAGAAATCATGGACAGTATCCGTGTTATTGGACGTGACAAT   1200
PMP1663      GAAAAAGGTGTTCCGATTGAAGAAATCATGGACAGTATCCGTGTTATTGGACGTGACAAT   1200
PMP1664      GAAAAAGGTGTTCCGATTGAAGAAATCATGGACAGTATCCGTGTTATTGGACGTGACAAT   1200
PMP1662      GAAAAAGGTGTTCCGATTGAAGAAATCATGGACAGTATCCGTGTTATTGGACGTGACAAT   1200
PMP1611      GAAAAAGGTGTTCCGATTGAAGAAATCATGGACAGTATCCGTGTTATTGGACGTGACAAT   1200
             ************************************************************

PMP1610      GCCCGTACCCCTATGCAATGGGACGAGAGCAAAAACGCTGGTTTCTCAACAGGTCAACCT   1260
PMP1666      GCCCGTACCCCTATGCAATGGGACGAGAGCAAAAACGCTGGTTTCTCAACAGGTCAACCT   1260
PMP1665      GCCCGTACCCCTATGCAATGGGACGAGAGCAAAAACGCTGGTTTCTCAACAGGTCAACCT   1260
PMP1667      GCCCGTACCCCTATGCAATGGGACGAGAGCAAAAACGCTGGTTTCTCAACAGGTCAACCT   1260
PMP1663      GCCCGTACCCCTATGCAATGGGACGAGAGCAAAAACGCTGGTTTCTCAACAGGTCAACCT   1260
PMP1664      GCCCGTACCCCTATGCAATGGGACGAGAGCAAAAACGCTGGTTTCTCAACAGGTCAACCT   1260
PMP1662      GCCCGTACCCCTATGCAATGGGACGAGAGCAAAAACGCTGGTTTCTCAACAGGTCAACCT   1260
PMP1611      GCCCGTACCCCTATGCAATGGGACGAGAGCAAAAACGCTGGTTTCTCAACAGGTCAACCT   1260
             ************************************************************

PMP1610      TGGTTGGCGGTTAATCCAAATTACGAGATGATCAATGTCCAAGAAGCGCTGGCAAATCCA   1320
PMP1666      TGGTTGGCGGTTAATCCAAATTACGAGATGATCAATGTCCAAGAAGCGCTGGCAAATCCA   1320
PMP1665      TGGTTGGCGGTTAATCCAAATTACGAGATGATCAATGTCCAAGAAGCGCTGGCAAATCCA   1320
PMP1667      TGGTTGGCGGTTAATCCAAATTACGAGATGATCAATGTCCAAGAAGCGCTGGCAAATCCA   1320
PMP1663      TGGTTGGCGGTTAATCCAAATTACGAGATGATCAATGTCCAAGAAGCGCTGGCAAATCCA   1320
PMP1664      TGGTTGGCGGTTAATCCAAATTACGAGATGATCAATGTCCAAGAAGCGCTGGCAAATCCA   1320
```

```
PMP1662      TGGTTGGCGGTTAATCCAAATTACGAGATGATCAATGTCCAAGAAGCGCTGGCAAATCCA    1320
PMP1611      TGGTTGGCGGTTAATCCAAATTACGAGATGATCAATGTCCAAGAAGCGCTGGCAAATCCA    1320
             ************************************************************

PMP1610      GATTCTATTTTCTATACCTATCAGAAACTGGTCCAAATTCGCAAGGAGAATAGCTGGCTA    1380
PMP1666      GATTCTATTTTCTATACCTATCAGAAACTGGTCCAAATTCGCAAGGAGAATAGCTGGCTA    1380
PMP1665      GATTCTATTTTCTATACCTATCAGAAACTGGTCCAAATTCGCAAGGAGAATAGCTGGCTA    1380
PMP1667      GATTCTATTTTCTATACCTATCAGAAACTGGTCCAAATTCGCAAGGAGAATAGCTGGCTA    1380
PMP1663      GATTCTATTTTCTATACCTATCAGAAACTGGTCCAAATTCGCAAGGAGAATAGCTGGCTA    1380
PMP1664      GATTCTATTTTCTATACCTATCAGAAACTGGTCCAAATTCGCAAGGAGAATAGCTGGCTA    1380
PMP1662      GATTCTATTTTCTATACCTATCAGAAACTGGTCCAAATTCGCAAGGAGAATAGCTGGCTA    1380
PMP1611      GATTCTATTTTCTATACCTATCAGAAACTGGTCCAAATTCGCAAGGAGAATAGCTGGCTA    1380
             ************************************************************

PMP1610      GTTCGAGCTGACTTTGAATTGCTTGATACGGCTGATAAGGTCTTTGCTTATATACGTAAG    1440
PMP1666      GTTCGAGCTGACTTTGAATTGCTTGATACGGCTGATAAGGTCTTTGCTTATATACGTAAG    1440
PMP1665      GTTCGAGCTGACTTTGAATTGCTTGATACGGCTGATAAGGTCTTTGCTTATATACGTAAG    1440
PMP1667      GTTCGAGCTGACTTTGAATTGCTTGATACGGCTGATAAGGTCTTTGCTTATATACGTAAG    1440
PMP1663      GTTCGAGCTGACTTTGAATTGCTTGATACGGCTGATAAGGTCTTTGCTTATATACGTAAG    1440
PMP1664      GTTCGAGCTGACTTTGAATTGCTTGATACGGCTGATAAGGTCTTTGCTTATATACGTAAG    1440
PMP1662      GTTCGAGCTGACTTTGAATTGCTTGATACGGCTGATAAGGTCTTTGCTTATATACGTAAG    1440
PMP1611      GTTCGAGCTGACTTTGAATTGCTTGATACGGCTGATAAGGTCTTTGCTTATATACGTAAG    1440
             ************************************************************

PMP1610      GATGGCGACCGTCGCTTCCTAGTCGTGGCTAATTTATCCAATGACAAACAAAACTTTTCA    1500
PMP1666      GATGGCGACCGTCGCTTCCTAGTCGTGGCTAATTTATCCAATGACAAACAAAACTTTTCA    1500
PMP1665      GATGGCGACCGTCGCTTCCTAGTCGTGGCTAATTTATCCAATGACAAACAAAACTTTTCA    1500
PMP1667      GATGGCGACCGTCGCTTCCTAGTCGTGGCTAATTTATCCAATGACAAACAAAACTTTTCA    1500
PMP1663      GATGGCGACCGTCGCTTCCTAGTCGTGGCTAATTTATCCAATGACAAACAAAACTTTTCA    1500
PMP1664      GATGGCGACCGTCGCTTCCTAGTCGTGGCTAATTTATCCAATGACAAACAAAACTTTTCA    1500
PMP1662      GATGGCGACCGTCGCTTCCTAGTCGTGGCTAATTTATCCAATGACAAACAAAACTTTTCA    1500
PMP1611      GATGGCGACCGTCGCTTCCTAGTCGTGGCTAATTTATCCAATGACAAACAAAACTTTTCA    1500
             ************************************************************

PMP1610      GTAGATGGAAAAGTTAGATCTGTCTTGATTGAAAACACTGCGGCTAAAGAAGTACTTGAA    1560
PMP1666      GTAGATGGAAAAGTTAGATCTGTCTTGATTGAAAACACTGCGGCTAAAGAAGTACTTGAA    1560
```

```
PMP1665      GTAGATGGAAAAGTTAGATCTGTCTTGATTGAAAACACTGCGGCTAAAGAAGTACTTGAA    1560
PMP1667      GTAGATGGAAAAGTTAGATCTGTCTTGATTGAAAACACTGCGGCTAAAGAAGTACTTGAA    1560
PMP1663      GTAGATGGAAAAGTTAGATCTGTCTTGATTGAAAACACTGCGGCTAAAGAAGTACTTGAA    1560
PMP1664      GTAGATGGAAAAGTTAGATCTGTCTTGATTGAAAACACTGCGGCTAAAGAAGTACTTGAA    1560
PMP1662      GTAGATGGAAAAGTTAGATCTGTCTTGATTGAAAACACTGCGGCTAAAGAAGTACTTGAA    1560
PMP1611      GTAGATGGAAAAGTTAGATCTGTCTTGATTGAAAACACTGCGGCTAAAGAAGTACTTGAA    1560
             ************************************************************

PMP1610      AAACAGGTCTTGGCTCCATGGGATGCTTTCTGTGTGGAAATGACTGATTAGAATGAGCAA    1620
PMP1666      AAACAGGTCTTGGCTCCATGGGATGCTTTCTGTGTGGAAATGACTGATTAGAATGAGCAA    1620
PMP1665      AAACAGGTCTTGGCTCCATGGGATGCTTTCTGTGTGGAAATGACTGATTAGAATGAGCAA    1620
PMP1667      AAACAGGTCTTGGCTCCATGGGATGCTTTCTGTGTGGAAATGACTGATTAGAATGAGCAA    1620
PMP1663      AAACAGGTCTTGGCTCCATGGGATGCTTTCTGTGTGGAAATGACTGATTAGAATGAGCAA    1620
PMP1664      AAACAGGTCTTGGCTCCATGGGATGCTTTCTGTGTGGAAATGACTGATTAGAATGAGCAA    1620
PMP1662      AAACAGGTCTTGGCTCCATGGGATGCTTTCTGTGTGGAAATGACTGATTAGAATGAGCAA    1620
PMP1611      AAACAGGTCTTGGCTCCATGGGATGCTTTCTGTGTGGAAATGACTGATTAGAATGAGCAA    1620
             ************************************************************

PMP1610      GAGCCTGGGACAAAATAGTTCTCAATCACAAAAAAGCTAGAGATTTCCAATTGTGAAACT    1680
PMP1666      GAGCCTGGGACAAAATAGTTCTCAATCACAAAAAAGCTAGAGATTTCCAATTGTGAAACT    1680
PMP1665      GAGCCTGGGACAAAATAGTTCTCAATCACAAAAAAGCTAGAGATTTCCAATTGTGAAACT    1680
PMP1667      GAGCCTGGGACAAAATAGTTCTCAATCACAAAAAAGCTAGAGATTTCCAATTGTGAAACT    1680
PMP1663      GAGCCTGGGACAAAATAGTTCTCAATCACAAAAAAGCTAGAGATTTCCAATTGTGAAACT    1680
PMP1664      GAGCCTGGGACAAAATAGTTCTCAATCACAAAAAAGCTAGAGATTTCCAATTGTGAAACT    1680
PMP1662      GAGCCTGGGACAAAATAGTTCTCAATCACAAAAAAGCTAGAGATTTCCAATTGTGAAACT    1680
PMP1611      GAGCCTGGGACAAAATAGTTCTCAATCACAAAAAAGCTAGAGATTTCCAATTGTGAAACT    1680
             ************************************************************

PMP1610      GTACTGCCCCCCAAAAGTTAGACAATTAATTTATCCGAAGGATTTAGTTCTGTATTGCAC    1740
PMP1666      GTACTGCCCCCCAAAAGTTAGACAATTAATTTATCCGAAGGATTTAGTTCTGTATTGCAC    1740
PMP1665      GTACTGCCCCCCAAAAGTTAGACAATTAATTTATCCGAAGGATTTAGTTCTGTATTGCAC    1740
PMP1667      GTACTGCCCCCCAAAAGTTAGACAATTAATTTATCCGAAGGATTTAGTTCTGTATTGCAC    1740
PMP1663      GTACTGCCCCCCAAAAGTTAGACAATTAATTTATCCGAAGGATTTAGTTCTGTATTGCAC    1740
PMP1664      GTACTGCCCCCCAAAAGTTAGACAATTAATTTATCCGAAGGATTTAGTTCTGTATTGCAC    1740
PMP1662      GTACTGCCCCCCAAAAGTTAGACAATTAATTTATCCGAAGGATTTAGTTCTGTATTGCAC    1740
PMP1611      GTACTGCCCCCCAAAAGTTAGACAATTAATTTATCCGAAGGATTTAGTTCTGTATTGCAC    1740
```

```
              ************************************************************

PMP1610       AGGACTAAGTCCTTTTAGTTTTACCTTAATTCGTTTGTTGTTGTAGTAATCAATGTAGTC    1800
PMP1666       AGGACTAAGTCCTTTTAGTTTTACCTTAATTCGTTTGTTGTTGTAGTAATCAATGTAGTC    1800
PMP1665       AGGACTAAGTCCTTTTAGTTTTACCTTAATTCGTTTGTTGTTGTAGTAATCAATGTAGTC    1800
PMP1667       AGGACTAAGTCCTTTTAGTTTTACCTTAATTCGTTTGTTGTTGTAGTAATCAATGTAGTC    1800
PMP1663       AGGACTAAGTCCTTTTAGTTTTACCTTAATTCGTTTGTTGTTGTAGTAATCAATGTAGTC    1800
PMP1664       AGGACTAAGTCCTTTTAGTTTTACCTTAATTCGTTTGTTGTTGTAGTAATCAATGTAGTC    1800
PMP1662       AGGACTAAGTCCTTTTAGTTTTACCTTAATTCGTTTGTTGTTGTAGTAATCAATGTAGTC    1800
PMP1611       AGGACTAAGTCCTTTTAGTTTTACCTTAATTCGTTTGTTGTTGTAGTAATCAATGTAGTC    1800
              ************************************************************

PMP1610       CACAATAGCTTGTTCAAGGTTTTCTAAAGATCTAAAGTTCTTCTCATAACCATAAAACAT    1860
PMP1666       CACAATAGCTTGTTCAAGGTTTTCTAAAGATCTAAAGTTCTTCTCATAACCATAAAACAT    1860
PMP1665       CACAATAGCTTGTTCAAGGTTTTCTAAAGATCTAAAGTTCTTCTCATAACCATAAAACAT    1860
PMP1667       CACAATAGCTTGTTCAAGGTTTTCTAAAGATCTAAAGTTCTTCTCATAACCATAAAACAT    1860
PMP1663       CACAATAGCTTGTTCAAGGTTTTCTAAAGATCTAAAGTTCTTCTCATAACCATAAAACAT    1860
PMP1664       CACAATAGCTTGTTCAAGGTTTTCTAAAGATCTAAAGTTCTTCTCATAACCATAAAACAT    1860
PMP1662       CACAATAGCTTGTTCAAGGTTTTCTAAAGATCTAAAGTTCTTCTCATAACCATAAAACAT    1860
PMP1611       CACAATAGCTTGTTCAAGGTTTTCTAAAGATCTAAAGTTCTTCTCATAACCATAAAACAT    1860
              ************************************************************

PMP1610       TTCGGATTTCAAGATGCCAAAGAAGGACTCCATCATGCCATTGTCTGGGCTGTTTCCTTT    1920
PMP1666       TTCGGATTTCAAGATGCCAAAGAAGGACTCCATCATGCCATTGTCTGGGCTGTTTCCTTT    1920
PMP1665       TTCGGATTTCAAGATGCCAAAGAAGGACTCCATCATGCCATTGTCTGGGCTGTTTCCTTT    1920
PMP1667       TTCGGATTTCAAGATGCCAAAGAAGGACTCCATCATGCCATTGTCTGGGCTGTTTCCTTT    1920
PMP1663       TTCGGATTTCAAGATGCCAAAGAAGGACTCCATCATGCCATTGTCTGGGCTGTTTCCTTT    1920
PMP1664       TTCGGATTTCAAGATGCCAAAGAAGGACTCCATCATGCCATTGTCTGGGCTGTTTCCTTT    1920
PMP1662       TTCGGATTTCAAGATGCCAAAGAAGGACTCCATCATGCCATTGTCTGGGCTGTTTCCTTT    1920
PMP1611       TTCGGATTTCAAGATGCCAAAGAAGGACTCCATCATGCCATTGTCTGGGCTGTTTCCTTT    1920
              ************************************************************

PMP1610       GCGTGACATAGATGGCTGAATTCCCTTACTCTCTAAAAACCTGTGATAAAAATCGTGTTG    1980
PMP1666       GCGTGACATAGATGGCTGAATTCCCTTACTCTCTAAAAACCTGTGATAAAAATCGTGTTG    1980
PMP1665       GCGTGACATAGATGGCTGAATTCCCTTACTCTCTAAAAACCTGTGATAAAAATCGTGTTG    1980
PMP1667       GCGTGACATAGATGGCTGAATTCCCTTACTCTCTAAAAACCTGTGATAAAAATCGTGTTG    1980
```

```
PMP1663      GCGTGACATAGATGGCTGAATTCCCTTACTCTCTAAAAACCTGTGATAAAAATCGTGTTG   1980
PMP1664      GCGTGACATAGATGGCTGAATTCCCTTACTCTCTAAAAACCTGTGATAAAAATCGTGTTG   1980
PMP1662      GCGTGACATAGATGGCTGAATTCCCTTACTCTCTAAAAACCTGTGATAAAAATCGTGTTG   1980
PMP1611      GCGTGACATAGATGGCTGAATTCCCTTACTCTCTAAAAACCTGTGATAAAAATCGTGTTG   1980
             ************************************************************

PMP1610      GTATTGCCAGCCTTGATCGCTATGGAGAATGGTATTCTCATAATGTTTCTCTGTAAAGGC   2040
PMP1666      GTATTGCCAGCCTTGATCGCTATGGAGAATGGTATTCTCATAATGTTTCTCTGTAAAGGC   2040
PMP1665      GTATTGCCAGCCTTGATCGCTATGGAGAATGGTATTCTCATAATGTTTCTCTGTAAAGGC   2040
PMP1667      GTATTGCCAGCCTTGATCGCTATGGAGAATGGTATTCTCATAATGTTTCTCTGTAAAGGC   2040
PMP1663      GTATTGCCAGCCTTGATCGCTATGGAGAATGGTATTCTCATAATGTTTCTCTGTAAAGGC   2040
PMP1664      GTATTGCCAGCCTTGATCGCTATGGAGAATGGTATTCTCATAATGTTTCTCTGTAAAGGC   2040
PMP1662      GTATTGCCAGCCTTGATCGCTATGGAGAATGGTATTCTCATAATGTTTCTCTGTAAAGGC   2040
PMP1611      GTATTGCCAGCCTTGATCGCTATGGAGAATGGTATTCTCATAATGTTTCTCTGTAAAGGC   2040
             ************************************************************

PMP1610      CTGGTTTAACATAGCTTTCACTTGTTCTAAGTTTGGTGAAGTTGAAAGATTATAGGCGAA   2100
PMP1666      CTGGTTTAACATAGCTTTCACTTGTTCTAAGTTTGGTGAAGTTGAAAGATTATAGGCGAA   2100
PMP1665      CTGGTTTAACATAGCTTTCACTTGTTCTAAGTTTGGTGAAGTTGAAAGATTATAGGCGAA   2100
PMP1667      CTGGTTTAACATAGCTTTCACTTGTTCTAAGTTTGGTGAAGTTGAAAGATTATAGGCGAA   2100
PMP1663      CTGGTTTAACATAGCTTTCACTTGTTCTAAGTTTGGTGAAGTTGAAAGATTATAGGCGAA   2100
PMP1664      CTGGTTTAACATAGCTTTCACTTGTTCTAAGTTTGGTGAAGTTGAAAGATTATAGGCGAA   2100
PMP1662      CTGGTTTAACATAGCTTTCACTTGTTCTAAGTTTGGTGAAGTTGAAAGATTATAGGCGAA   2100
PMP1611      CTGGTTTAACATAGCTTTCACTTGTTCTAAGTTTGGTGAAGTTGAAAGATTATAGGCGAA   2100
             ************************************************************

PMP1610      ATTTCGCTGTTAAAGCCATCTAAAACAGGCGATAAATAGAGTTTCTGTGTGCTATTTGGA   2160
PMP1666      ATTTCGCTGTTAAAGCCATCTAAAACAGGCGATAAATAGAGTTTCTGTGTGCTATTTGGA   2160
PMP1665      ATTTCGCTGTTAAAGCCATCTAAAACAGGCGATAAATAGAGTTTCTGTGTGCTATTTGGA   2160
PMP1667      ATTTCGCTGTTAAAGCCATCTAAAACAGGCGATAAATAGAGTTTCTGTGTGCTATTTGGA   2160
PMP1663      ATTTCGCTGTTAAAGCCATCTAAAACAGGCGATAAATAGAGTTTCTGTGTGCTATTTGGA   2160
PMP1664      ATTTCGCTGTTAAAGCCATCTAAAACAGGCGATAAATAGAGTTTCTGTGTGCTATTTGGA   2160
PMP1662      ATTTCGCTGTTAAAGCCATCTAAAACAGGCGATAAATAGAGTTTCTGTGTGCTATTTGGA   2160
PMP1611      ATTTCGCTGTTAAAGCCATCTAAAACAGGCGATAAATAGAGTTTCTGTGTGCTATTTGGA   2160
             ************************************************************
```

```
PMP1610      ATGGCAAACTCTGTCACATCCGTATAGCACTTTTCCATTGGTTTGGTTGCTTCAAACTGA   2220
PMP1666      ATGGCAAACTCTGTCACATCCGTATAGCACTTTTCCATTGGTTTGGTTGCTTCAAACTGA   2220
PMP1665      ATGGCAAACTCTGTCACATCCGTATAGCACTTTTCCATTGGTTTGGTTGCTTCAAACTGA   2220
PMP1667      ATGGCAAACTCTGTCACATCCGTATAGCACTTTTCCATTGGTTTGGTTGCTTCAAACTGA   2220
PMP1663      ATGGCAAACTCTGTCACATCCGTATAGCACTTTTCCATTGGTTTGGTTGCTTCAAACTGA   2220
PMP1664      ATGGCAAACTCTGTCACATCCGTATAGCACTTTTCCATTGGTTTGGTTGCTTCAAACTGA   2220
PMP1662      ATGGCAAACTCTGTCACATCCGTATAGCACTTTTCCATTGGTTTGGTTGCTTCAAACTGA   2220
PMP1611      ATGGCAAACTCTGTCACATCCGTATAGCACTTTTCCATTGGTTTGGTTGCTTCAAACTGA   2220
             ************************************************************

PMP1610      CGTTGAATGAGATTCTCTGCTTTCTTGCCAATCTCTCCTTGGTAGGAGGAATACTTTCGT   2280
PMP1666      CGTTGAATGAGATTCTCTGCTTTCTTGCCAATCTCTCCTTGGTAGGAGGAATACTTTCGT   2280
PMP1665      CGTTGAATGAGATTCTCTGCTTTCTTGCCAATCTCTCCTTGGTAGGAGGAATACTTTCGT   2280
PMP1667      CGTTGAATGAGATTCTCTGCTTTCTTGCCAATCTCTCCTTGGTAGGAGGAATACTTTCGT   2280
PMP1663      CGTTGAATGAGATTCTCTGCTTTCTTGCCAATCTCTCCTTGGTAGGAGGAATACTTTCGT   2280
PMP1664      CGTTGAATGAGATTCTCTGCTTTCTTGCCAATCTCTCCTTGGTAGGAGGAATACTTTCGT   2280
PMP1662      CGTTGAATGAGATTCTCTGCTTTCTTGCCAATCTCTCCTTGGTAGGAGGAATACTTTCGT   2280
PMP1611      CGTTGAATGAGATTCTCTGCTTTCTTGCCAATCTCTCCTTGGTAGGAGGAATACTTTCGT   2280
             ************************************************************

PMP1610      TTCCGACGAATTCGAGCTGTTAAACCAAGAACTTTCATCAGACGTTGAACTTTCTTATGA   2340
PMP1666      TTCCGACGAATTCGAGCTGTTAAACCAAGAACTTTCATCAGACGTTGAACTTTCTTATGA   2340
PMP1665      TTCCGACGAATTCGAGCTGTTAAACCAAGAACTTTCATCAGACGTTGAACTTTCTTATGA   2340
PMP1667      TTCCGACGAATTCGAGCTGTTAAACCAAGAACTTTCATCAGACGTTGAACTTTCTTATGA   2340
PMP1663      TTCCGACGAATTCGAGCTGTTAAACCAAGAACTTTCATCAGACGTTGAACTTTCTTATGA   2340
PMP1664      TTCCGACGAATTCGAGCTGTTAAACCAAGAACTTTCATCAGACGTTGAACTTTCTTATGA   2340
PMP1662      TTCCGACGAATTCGAGCTGTTAAACCAAGAACTTTCATCAGACGTTGAACTTTCTTATGA   2340
PMP1611      TTCCGACGAATTCGAGCTGTTAAACCAAGAACTTTCATCAGACGTTGAACTTTCTTATGA   2340
             ************************************************************

PMP1610      TTCACCACAAAACCACGATTTCTTAGTTCTAAATGGATTCGACGATAGCCATAGTTGCCT   2400
PMP1666      TTCACCACAAAACCACGATTTCTTAGTTCTAAATGGATTCGACGATAGCCATAGTTGCCT   2400
PMP1665      TTCACCACAAAACCACGATTTCTTAGTTCTAAATGGATTCGACGATAGCCATAGTTGCCT   2400
PMP1667      TTCACCACAAAACCACGATTTCTTAGTTCTAAATGGATTCGACGATAGCCATAGTTGCCT   2400
PMP1663      TTCACCACAAAACCACGATTTCTTAGTTCTAAATGGATTCGACGATAGCCATAGTTGCCT   2400
PMP1664      TTCACCACAAAACCACGATTTCTTAGTTCTAAATGGATTCGACGATAGCCATAGTTGCCT   2400
```

```
PMP1662      TTCACCACAAAACCACGATTTCTTAGTTCTAAATGGATTCGACGATAGCCATAGTTGCCT   2400
PMP1611      TTCACCACAAAACCACGATTTCTTAGTTCTAAATGGATTCGACGATAGCCATAGTTGCCT   2400
             ************************************************************

PMP1610      TTATGTTCATTATAAATGGCCTGAATTTCGGTTTTAATCTCTTTATCTTTATCAACCCCA   2460
PMP1666      TTATGTTCATTATAAATGGCCTGAATTTCGGTTTTAATCTCTTTATCTTTATCAACCCCA   2460
PMP1665      TTATGTTCATTATAAATGGCCTGAATTTCGGTTTTAATCTCTTTATCTTTATCAACCCCA   2460
PMP1667      TTATGTTCATTATAAATGGCCTGAATTTCGGTTTTAATCTCTTTATCTTTATCAACCCCA   2460
PMP1663      TTATGTTCATTATAAATGGCCTGAATTTCGGTTTTAATCTCTTTATCTTTATCAACCCCA   2460
PMP1664      TTATGTTCATTATAAATGGCCTGAATTTCGGTTTTAATCTCTTTATCTTTATCAACCCCA   2460
PMP1662      TTATGTTCATTATAAATGGCCTGAATTTCGGTTTTAATCTCTTTATCTTTATCAACCCCA   2460
PMP1611      TTATGTTCATTATAAATGGCCTGAATTTCGGTTTTAATCTCTTTATCTTTATCAACCCCA   2460
             ************************************************************

PMP1610      TCCAGTTGTTTCAACTGATAATAGTAAGTCGAGCGAGCTAAACGTGCTGTTTCAAGAAGA   2520
PMP1666      TCCAGTTGTTTCAACTGATAATAGTAAGTCGAGCGAGCTAAACGTGCTGTTTCAAGAAGA   2520
PMP1665      TCCAGTTGTTTCAACTGATAATAGTAAGTCGAGCGAGCTAAACGTGCTGTTTCAAGAAGA   2520
PMP1667      TCCAGTTGTTTCAACTGATAATAGTAAGTCGAGCGAGCTAAACGTGCTGTTTCAAGAAGA   2520
PMP1663      TCCAGTTGTTTCAACTGATAATAGTAAGTCGAGCGAGCTAAACGTGCTGTTTCAAGAAGA   2520
PMP1664      TCCAGTTGTTTCAACTGATAATAGTAAGTCGAGCGAGCTAAACGTGCTGTTTCAAGAAGA   2520
PMP1662      TCCAGTTGTTTCAACTGATAATAGTAAGTCGAGCGAGCTAAACGTGCTGTTTCAAGAAGA   2520
PMP1611      TCCAGTTGTTTCAACTGATAATAGTAAGTCGAGCGAGCTAAACGTGCTGTTTCAAGAAGA   2520
             ************************************************************

PMP1610      AAATCTAGTCGAAATCCTCCTGAAACCATTTCTCTAACTGTTTCTGCCTTTCTCGCTCTA   2580
PMP1666      AAATCTAGTCGAAATCCTCCTGAAACCATTTCTCTAACTGTTTCTGCCTTTCTCGCTCTA   2580
PMP1665      AAATCTAGTCGAAATCCTCCTGAAACCATTTCTCTAACTGTTTCTGCCTTTCTCGCTCTA   2580
PMP1667      AAATCTAGTCGAAATCCTCCTGAAACCATTTCTCTAACTGTTTCTGCCTTTCTCGCTCTA   2580
PMP1663      AAATCTAGTCGAAATCCTCCTGAAACCATTTCTCTAACTGTTTCTGCCTTTCTCGCTCTA   2580
PMP1664      AAATCTAGTCGAAATCCTCCTGAAACCATTTCTCTAACTGTTTCTGCCTTTCTCGCTCTA   2580
PMP1662      AAATCTAGTCGAAATCCTCCTGAAACCATTTCTCTAACTGTTTCTGCCTTTCTCGCTCTA   2580
PMP1611      AAATCTAGTCGAAATCCTCCTGAAACCATTTCTCTAACTGTTTCTGCCTTTCTCGCTCTA   2580
             ************************************************************

PMP1610      GGGCTTCGTCCCTTTCCTCTAACTCTTTTAACTTTTTTAAGTAAGCCACCTCAGTCCGTA   2640
PMP1666      GGGCTTCGTCCCTTTCCTCTAACTCTTTTAACTTTTTTAAGTAAGCCACCTCAGTCCGTA   2640
```

```
PMP1665      GGGCTTCGTCCCTTTCCTCTAACTCTTTTAACTTTTTTAAGTAAGCCACCTCAGTCCGTA    2640
PMP1667      GGGCTTCGTCCCTTTCCTCTAACTCTTTTAACTTTTTTAAGTAAGCCACCTCAGTCCGTA    2640
PMP1663      GGGCTTCGTCCCTTTCCTCTAACTCTTTTAACTTTTTTAAGTAAGCCACCTCAGTCCGTA    2640
PMP1664      GGGCTTCGTCCCTTTCCTCTAACTCTTTTAACTTTTTTAAGTAAGCCACCTCAGTCCGTA    2640
PMP1662      GGGCTTCGTCCCTTTCCTCTAACTCTTTTAACTTTTTTAAGTAAGCCACCTCAGTCCGTA    2640
PMP1611      GGGCTTCGTCCCTTTCCTCTAACTCTTTTAACTTTTTTAAGTAAGCCACCTCAGTCCGTA    2640
             ************************************************************

PMP1610      AGCGTTCATTCTCCTCTTGAAGTCGTTCTAATTCTGTCATTTCTTCCCAAGTTTTCTTTC    2700
PMP1666      AGCGTTCATTCTCCTCTTGAAGTCGTTCTAATTCTGTCATTTCTTCCCAAGTTTTCTTTC    2700
PMP1665      AGCGTTCATTCTCCTCTTGAAGTCGTTCTAATTCTGTCATTTCTTCCCAAGTTTTCTTTC    2700
PMP1667      AGCGTTCATTCTCCTCTTGAAGTCGTTCTAATTCTGTCATTTCTTCCCAAGTTTTCTTTC    2700
PMP1663      AGCGTTCATTCTCCTCTTGAAGTCGTTCTAATTCTGTCATTTCTTCCCAAGTTTTCTTTC    2700
PMP1664      AGCGTTCATTCTCCTCTTGAAGTCGTTCTAATTCTGTCATTTCTTCCCAAGTTTTCTTTC    2700
PMP1662      AGCGTTCATTCTCCTCTTGAAGTCGTTCTAATTCTGTCATTTCTTCCCAAGTTTTCTTTC    2700
PMP1611      AGCGTTCATTCTCCTCTTGAAGTCGTTCTAATTCTGTCATTTCTTCCCAAGTTTTCTTTC    2700
             ************************************************************

PMP1610      GTTTACGTTCCATTTTAGCTGGTCTCCCTCTTGTTTTCTCAACAATAGTATACCCGTTTT    2760
PMP1666      GTTTACGTTCCATTTTAGCTGGTCTCCCTCTTGTTTTCTCAACAATAGTATACCCGTTTT    2760
PMP1665      GTTTACGTTCCATTTTAGCTGGTCTCCCTCTTGTTTTCTCAACAATAGTATACCCGTTTT    2760
PMP1667      GTTTACGTTCCATTTTAGCTGGTCTCCCTCTTGTTTTCTCAACAATAGTATACCCGTTTT    2760
PMP1663      GTTTACGTTCCATTTTAGCTGGTCTCCCTCTTGTTTTCTCAACAATAGTATACCCGTTTT    2760
PMP1664      GTTTACGTTCCATTTTAGCTGGTCTCCCTCTTGTTTTCTCAACAATAGTATACCCGTTTT    2760
PMP1662      GTTTACGTTCCATTTTAGCTGGTCTCCCTCTTGTTTTCTCAACAATAGTATACCCGTTTT    2760
PMP1611      GTTTACGTTCCATTTTAGCTGGTCTCCCTCTTGTTTTCTCAACAATAGTATACCCGTTTT    2760
             ************************************************************

PMP1610      TCTTATATTGCGCCAGCCAATTTGGAAGCATACCACGATTTGGGAGAGCATATTTGAGAG    2820
PMP1666      TCTTATATTGCGCCAGCCAATTTGGAAGCATACCACGATTTGGGAGAGCATATTTGAGAG    2820
PMP1665      TCTTATATTGCGCCAGCCAATTTGGAAGCATACCACGATTTGGGAGAGCATATTTGAGAG    2820
PMP1667      TCTTATATTGCGCCAGCCAATTTGGAAGCATACCACGATTTGGGAGAGCATATTTGAGAG    2820
PMP1663      TCTTATATTGCGCCAGCCAATTTGGAAGCATACCACGATTTGGGAGAGCATATTTGAGAG    2820
PMP1664      TCTTATATTGCGCCAGCCAATTTGGAAGCATACCACGATTTGGGAGAGCATATTTGAGAG    2820
PMP1662      TCTTATATTGCGCCAGCCAATTTGGAAGCATACCACGATTTGGGAGAGCATATTTGAGAG    2820
PMP1611      TCTTATATTGCGCCAGCCAATTTGGAAGCATACCACGATTTGGGAGAGCATATTTGAGAG    2820
```

```
             ************************************************************

PMP1610      AAACTCTATCTTGAGACCAGTTTGCATGTAAAACTTTATCAATCATTTCTTGTTTTAGTT    2880
PMP1666      AAACTCTATCTTGAGACCAGTTTGCATGTAAAACTTTATCAATCATTTCTTGTTTTAGTT    2880
PMP1665      AAACTCTATCTTGAGACCAGTTTGCATGTAAAACTTTATCAATCATTTCTTGTTTTAGTT    2880
PMP1667      AAACTCTATCTTGAGACCAGTTTGCATGTAAAACTTTATCAATCATTTCTTGTTTTAGTT    2880
PMP1663      AAACTCTATCTTGAGACCAGTTTGCATGTAAAACTTTATCAATCATTTCTTGTTTTAGTT    2880
PMP1664      AAACTCTATCTTGAGACCAGTTTGCATGTAAAACTTTATCAATCATTTCTTGTTTTAGTT    2880
PMP1662      AAACTCTATCTTGAGACCAGTTTGCATGTAAAACTTTATCAATCATTTCTTGTTTTAGTT    2880
PMP1611      AAACTCTATCTTGAGACCAGTTTGCATGTAAAACTTTATCAATCATTTCTTGTTTTAGTT    2880
             ************************************************************

PMP1610      CTGGAGAATAGTATTGATTTTTCCCTTTTTTGACGAACTCTATTCCGTAACGATCAATCA    2940
PMP1666      CTGGAGAATAGTATTGATTTTTCCCTTTTTTGACGAACTCTATTCCGTAACGATCAATCA    2940
PMP1665      CTGGAGAATAGTATTGATTTTTCCCTTTTTTGACGAACTCTATTCCGTAACGATCAATCA    2940
PMP1667      CTGGAGAATAGTATTGATTTTTCCCTTTTTTGACGAACTCTATTCCGTAACGATCAATCA    2940
PMP1663      CTGGAGAATAGTATTGATTTTTCCCTTTTTTGACGAACTCTATTCCGTAACGATCAATCA    2940
PMP1664      CTGGAGAATAGTATTGATTTTTCCCTTTTTTGACGAACTCTATTCCGTAACGATCAATCA    2940
PMP1662      CTGGAGAATAGTATTGATTTTTCCCTTTTTTGACGAACTCTATTCCGTAACGATCAATCA    2940
PMP1611      CTGGAGAATAGTATTGATTTTTCCCTTTTTTGACGAACTCTATTCCGTAACGATCAATCA    2940
             ************************************************************

PMP1610      ATTTAATCATATACCTAAGATTAGAATTATTTATCCCAAATTTATTTGAAAGCTTCTCTA    3000
PMP1666      ATTTAATCATATACCTAAGATTAGAATTATTTATCCCAAATTTATTTGAAAGCTTCTCTA    3000
PMP1665      ATTTAATCATATACCTAAGATTAGAATTATTTATCCCAAATTTATTTGAAAGCTTCTCTA    3000
PMP1667      ATTTAATCATATACCTAAGATTAGAATTATTTATCCCAAATTTATTTGAAAGCTTCTCTA    3000
PMP1663      ATTTAATCATATACCTAAGATTAGAATTATTTATCCCAAATTTATTTGAAAGCTTCTCTA    3000
PMP1664      ATTTAATCATATACCTAAGATTAGAATTATTTATCCCAAATTTATTTGAAAGCTTCTCTA    3000
PMP1662      ATTTAATCATATACCTAAGATTAGAATTATTTATCCCAAATTTATTTGAAAGCTTCTCTA    3000
PMP1611      ATTTAATCATATACCTAAGATTAGAATTATTTATCCCAAATTTATTTGAAAGCTTCTCTA    3000
             ************************************************************

PMP1610      AGCTATATCCTTGTTTTCTAAGTTCATAGATCTGAACTTTATCATCATAAGTTAATTTCA    3060
PMP1666      AGCTATATCCTTGTTTTCTAAGTTCATAGATCTGAACTTTATCATCATAAGTTAATTTCA    3060
PMP1665      AGCTATATCCTTGTTTTCTAAGTTCATAGATCTGAACTTTATCATCATAAGTTAATTTCA    3060
PMP1667      AGCTATATCCTTGTTTTCTAAGTTCATAGATCTGAACTTTATCATCATAAGTTAATTTCA    3060
```

```
PMP1663      AGCTATATCCTTGTTTTCTAAGTTCATAGATCTGAACTTTATCATCATAAGTTAATTTCA   3060
PMP1664      AGCTATATCCTTGTTTTCTAAGTTCATAGATCTGAACTTTATCATCATAAGTTAATTTCA   3060
PMP1662      AGCTATATCCTTGTTTTCTAAGTTCATAGATCTGAACTTTATCATCATAAGTTAATTTCA   3060
PMP1611      AGCTATATCCTTGTTTTCTAAGTTCATAGATCTGAACTTTATCATCATAAGTTAATTTCA   3060
             ************************************************************

PMP1610      TAATAAAAACACCCCAAAAGTTAGATTTTTTCTGTCTAACTTTTGGGGTGCAGTTCAAAC   3120
PMP1666      TAATAAAAACACCCCAAAAGTTAGATTTTTTCTGTCTAACTTTTGGGGTGCAGTTCAAAC   3120
PMP1665      TAATAAAAACACCCCAAAAGTTAGATTTTTTCTGTCTAACTTTTGGGGTGCAGTTCAAAC   3120
PMP1667      TAATAAAAACACCCCAAAAGTTAGATTTTTTCTGTCTAACTTTTGGGGTGCAGTTCAAAC   3120
PMP1663      TAATAAAAACACCCCAAAAGTTAGATTTTTTCTGTCTAACTTTTGGGGTGCAGTTCAAAC   3120
PMP1664      TAATAAAAACACCCCAAAAGTTAGATTTTTTCTGTCTAACTTTTGGGGTGCAGTTCAAAC   3120
PMP1662      TAATAAAAACACCCCAAAAGTTAGATTTTTTCTGTCTAACTTTTGGGGTGCAGTTCAAAC   3120
PMP1611      TAATAAAAACACCCCAAAAGTTAGATTTTTTCTGTCTAACTTTTGGGGTGCAGTTCAAAC   3120
             ************************************************************

PMP1610      TCTAGCTTTTTAATTTTGAGTCGTTCTCTTATTGTATTTTAGGAGGTTATTGGCCATAAG   3180
PMP1666      TCTAGCTTTTTAATTTTGAGTCGTTCTCTTATTGTATTTTAGGAGGTTATTGGCCATAAG   3180
PMP1665      TCTAGCTTTTTAATTTTGAGTCGTTCTCTTATTGTATTTTAGGAGGTTATTGGCCATAAG   3180
PMP1667      TCTAGCTTTTTAATTTTGAGTCGTTCTCTTATTGTATTTTAGGAGGTTATTGGCCATAAG   3180
PMP1663      TCTAGCTTTTTAATTTTGAGTCGTTCTCTTATTGTATTTTAGGAGGTTATTGGCCATAAG   3180
PMP1664      TCTAGCTTTTTAATTTTGAGTCGTTCTCTTATTGTATTTTAGGAGGTTATTGGCCATAAG   3180
PMP1662      TCTAGCTTTTTAATTTTGAGTCGTTCTCTTATTGTATTTTAGGAGGTTATTGGCCATAAG   3180
PMP1611      TCTAGCTTTTTAATTTTGAGTCGTTCTCTTATTGTATTTTAGGAGGTTATTGGCCATAAG   3180
             ************************************************************

PMP1610      TACCAATCCCATGTCAATTCTCACTTGACGCTTGCCCCTCAGATTACATCTCTTGTAACC   3240
PMP1666      TACCAATCCCATGTCAATTCTCACTTGACGCTTGCCCCTCAGATTACATCTCTTGTAACC   3240
PMP1665      TACCAATCCCATGTCAATTCTCACTTGACGCTTGCCCCTCAGATTACATCTCTTGTAACC   3240
PMP1667      TACCAATCCCATGTCAATTCTCACTTGACGCTTGCCCCTCAGATTACATCTCTTGTAACC   3240
PMP1663      TACCAATCCCATGTCAATTCTCACTTGACGCTTGCCCCTCAGATTACATCTCTTGTAACC   3240
PMP1664      TACCAATCCCATGTCAATTCTCACTTGACGCTTGCCCCTCAGATTACATCTCTTGTAACC   3240
PMP1662      TACCAATCCCATGTCAATTCTCACTTGACGCTTGCCCCTCAGATTACATCTCTTGTAACC   3240
PMP1611      TACCAATCCCATGTCAATTCTCACTTGACGCTTGCCCCTCAGATTACATCTCTTGTAACC   3240
             ************************************************************
```

```
PMP1610     CAAACAAGCCTTTATCTGCCCAAAGACAGGTTCCACATCAATCTTGCGTTGAGCGAAAAT   3300
PMP1666     CAAACAAGCCTTTATCTGCCCAAAGACAGGTTCCACATCAATCTTGCGTTGAGCGAAAAT   3300
PMP1665     CAAACAAGCCTTTATCTGCCCAAAGACAGGTTCCACATCAATCTTGCGTTGAGCGAAAAT   3300
PMP1667     CAAACAAGCCTTTATCTGCCCAAAGACAGGTTCCACATCAATCTTGCGTTGAGCGAAAAT   3300
PMP1663     CAAACAAGCCTTTATCTGCCCAAAGACAGGTTCCACATCAATCTTGCGTTGAGCGAAAAT   3300
PMP1664     CAAACAAGCCTTTATCTGCCCAAAGACAGGTTCCACATCAATCTTGCGTTGAGCGAAAAT   3300
PMP1662     CAAACAAGCCTTTATCTGCCCAAAGACAGGTTCCACATCAATCTTGCGTTGAGCGAAAAT   3300
PMP1611     CAAACAAGCCTTTATCTGCCCAAAGACAGGTTCCACATCAATCTTGCGTTGAGCGAAAAT   3300
            ************************************************************

PMP1610     CTGTCTACCTTGGGGAGATAAAAGCGCTTGACATTCTTTCGCCTTCAAGTTTTGATAGCC   3360
PMP1666     CTGTCTACCTTGGGGAGATAAAAGCGCTTGACATTCTTTCGCCTTCAAGTTTTGATAGCC   3360
PMP1665     CTGTCTACCTTGGGGAGATAAAAGCGCTTGACATTCTTTCGCCTTCAAGTTTTGATAGCC   3360
PMP1667     CTGTCTACCTTGGGGAGATAAAAGCGCTTGACATTCTTTCGCCTTCAAGTTTTGATAGCC   3360
PMP1663     CTGTCTACCTTGGGGAGATAAAAGCGCTTGACATTCTTTCGCCTTCAAGTTTTGATAGCC   3360
PMP1664     CTGTCTACCTTGGGGAGATAAAAGCGCTTGACATTCTTTCGCCTTCAAGTTTTGATAGCC   3360
PMP1662     CTGTCTACCTTGGGGAGATAAAAGCGCTTGACATTCTTTCGCCTTCAAGTTTTGATAGCC   3360
PMP1611     CTGTCTACCTTGGGGAGATAAAAGCGCTTGACATTCTTTCGCCTTCAAGTTTTGATAGCC   3360
            ************************************************************

PMP1610     TTCGTTCATATACAGTCCCTTTTGAGGGGCTGATTCAGGTTCGTCGGCGTAGTAAACCTT   3420
PMP1666     TTCGTTCATATACAGTCCCTTTTGAGGGGCTGATTCAGGTTCGTCGGCGTAGTAAACCTT   3420
PMP1665     TTCGTTCATATACAGTCCCTTTTGAGGGGCTGATTCAGGTTCGTCGGCGTAGTAAACCTT   3420
PMP1667     TTCGTTCATATACAGTCCCTTTTGAGGGGCTGATTCAGGTTCGTCGGCGTAGTAAACCTT   3420
PMP1663     TTCGTTCATATACAGTCCCTTTTGAGGGGCTGATTCAGGTTCGTCGGCGTAGTAAACCTT   3420
PMP1664     TTCGTTCATATACAGTCCCTTTTGAGGGGCTGATTCAGGTTCGTCGGCGTAGTAAACCTT   3420
PMP1662     TTCGTTCATATACAGTCCCTTTTGAGGGGCTGATTCAGGTTCGTCGGCGTAGTAAACCTT   3420
PMP1611     TTCGTTCATATACAGTCCCTTTTGAGGGGCTGATTCAGGTTCGTCGGCGTAGTAAACCTT   3420
            ************************************************************

PMP1610     GATTTCTTGTTGAAAGTCTGTCTGTGTTTTCTGATGTTTGGTATGGTGAAAACGATAATA   3480
PMP1666     GATTTCTTGTTGAAAGTCTGTCTGTGTTTTCTGATGTTTGGTATGGTGAAAACGATAATA   3480
PMP1665     GATTTCTTGTTGAAAGTCTGTCTGTGTTTTCTGATGTTTGGTATGGTGAAAACGATAATA   3480
PMP1667     GATTTCTTGTTGAAAGTCTGTCTGTGTTTTCTGATGTTTGGTATGGTGAAAACGATAATA   3480
PMP1663     GATTTCTTGTTGAAAGTCTGTCTGTGTTTTCTGATGTTTGGTATGGTGAAAACGATAATA   3480
PMP1664     GATTTCTTGTTGAAAGTCTGTCTGTGTTTTCTGATGTTTGGTATGGTGAAAACGATAATA   3480
```

```
PMP1662      GATTTCTTGTTGAAAGTCTGTCTGTGTTTTCTGATGTTTGGTATGGTGAAAACGATAATA   3480
PMP1611      GATTTCTTGTTGAAAGTCTGTCTGTGTTTTCTGATGTTTGGTATGGTGAAAACGATAATA   3480
             ************************************************************

PMP1610      CCAGCCATCAGGATGTGTATAGCTATCCTCCTTGTCATTATAGTGCCAATTCGCTAAGTT   3540
PMP1666      CCAGCCATCAGGATGTGTATAGCTATCCTCCTTGTCATTATAGTGCCAATTCGCTAAGTT   3540
PMP1665      CCAGCCATCAGGATGTGTATAGCTATCCTCCTTGTCATTATAGTGCCAATTCGCTAAGTT   3540
PMP1667      CCAGCCATCAGGATGTGTATAGCTATCCTCCTTGTCATTATAGTGCCAATTCGCTAAGTT   3540
PMP1663      CCAGCCATCAGGATGTGTATAGCTATCCTCCTTGTCATTATAGTGCCAATTCGCTAAGTT   3540
PMP1664      CCAGCCATCAGGATGTGTATAGCTATCCTCCTTGTCATTATAGTGCCAATTCGCTAAGTT   3540
PMP1662      CCAGCCATCAGGATGTGTATAGCTATCCTCCTTGTCATTATAGTGCCAATTCGCTAAGTT   3540
PMP1611      CCAGCCATCAGGATGTGTATAGCTATCCTCCTTGTCATTATAGTGCCAATTCGCTAAGTT   3540
             ************************************************************

PMP1610      TCTAGCTGACTGTTTATAGCCTCTCTTCTGTTCCTTATCAAACATGGCATATTTAATCAG   3600
PMP1666      TCTAGCTGACTGTTTATAGCCTCTCTTCTGTTCCTTATCAAACATGGCATATTTAATCAG   3600
PMP1665      TCTAGCTGACTGTTTATAGCCTCTCTTCTGTTCCTTATCAAACATGGCATATTTAATCAG   3600
PMP1667      TCTAGCTGACTGTTTATAGCCTCTCTTCTGTTCCTTATCAAACATGGCATATTTAATCAG   3600
PMP1663      TCTAGCTGACTGTTTATAGCCTCTCTTCTGTTCCTTATCAAACATGGCATATTTAATCAG   3600
PMP1664      TCTAGCTGACTGTTTATAGCCTCTCTTCTGTTCCTTATCAAACATGGCATATTTAATCAG   3600
PMP1662      TCTAGCTGACTGTTTATAGCCTCTCTTCTGTTCCTTATCAAACATGGCATATTTAATCAG   3600
PMP1611      TCTAGCTGACTGTTTATAGCCTCTCTTCTGTTCCTTATCAAACATGGCATATTTAATCAG   3600
             ************************************************************

PMP1610      ATGGTTTACCTCCTTTTCATCTAAACGAAGGAGGTTCTCTTCACTTCCATATCCAGCATC   3660
PMP1666      ATGGTTTACCTCCTTTTCATCTAAACGAAGGAGGTTCTCTTCACTTCCATATCCAGCATC   3660
PMP1665      ATGGTTTACCTCCTTTTCATCTAAACGAAGGAGGTTCTCTTCACTTCCATATCCAGCATC   3660
PMP1667      ATGGTTTACCTCCTTTTCATCTAAACGAAGGAGGTTCTCTTCACTTCCATATCCAGCATC   3660
PMP1663      ATGGTTTACCTCCTTTTCATCTAAACGAAGGAGGTTCTCTTCACTTCCATATCCAGCATC   3660
PMP1664      ATGGTTTACCTCCTTTTCATCTAAACGAAGGAGGTTCTCTTCACTTCCATATCCAGCATC   3660
PMP1662      ATGGTTTACCTCCTTTTCATCTAAACGAAGGAGGTTCTCTTCACTTCCATATCCAGCATC   3660
PMP1611      ATGGTTTACCTCCTTTTCATCTAAACGAAGGAGGTTCTCTTCACTTCCATATCCAGCATC   3660
             ************************************************************

PMP1610      TGCGACAACTGTCTTCAAGTCATGCGGATAGGTTTCAAGGAATGGCAGAAGAGTCTTGGT   3720
PMP1666      TGCGACAACTGTCTTCAAGTCATGCGGATAGGTTTCAAGGAATGGCAGAAGAGTCTTGGT   3720
```

```
PMP1665      TGCGACAACTGTCTTCAAGTCATGCGGATAGGTTTCAAGGAATGGCAGAAGAGTCTTGGT    3720
PMP1667      TGCGACAACTGTCTTCAAGTCATGCGGATAGGTTTCAAGGAATGGCAGAAGAGTCTTGGT    3720
PMP1663      TGCGACAACTGTCTTCAAGTCATGCGGATAGGTTTCAAGGAATGGCAGAAGAGTCTTGGT    3720
PMP1664      TGCGACAACTGTCTTCAAGTCATGCGGATAGGTTTCAAGGAATGGCAGAAGAGTCTTGGT    3720
PMP1662      TGCGACAACTGTCTTCAAGTCATGCGGATAGGTTTCAAGGAATGGCAGAAGAGTCTTGGT    3720
PMP1611      TGCGACAACTGTCTTCAAGTCATGCGGATAGGTTTCAAGGAATGGCAGAAGAGTCTTGGT    3720
             ************************************************************

PMP1610      ATCTGTCGGATTTGAAAAGACATCATAGTGAAGAACAAATTGGTTTTCCGTAGCGATTTG    3780
PMP1666      ATCTGTCGGATTTGAAAAGACATCATAGTGAAGAACAAATTGGTTTTCCGTAGCGATTTG    3780
PMP1665      ATCTGTCGGATTTGAAAAGACATCATAGTGAAGAACAAATTGGTTTTCCGTAGCGATTTG    3780
PMP1667      ATCTGTCGGATTTGAAAAGACATCATAGTGAAGAACAAATTGGTTTTCCGTAGCGATTTG    3780
PMP1663      ATCTGTCGGATTTGAAAAGACATCATAGTGAAGAACAAATTGGTTTTCCGTAGCGATTTG    3780
PMP1664      ATCTGTCGGATTTGAAAAGACATCATAGTGAAGAACAAATTGGTTTTCCGTAGCGATTTG    3780
PMP1662      ATCTGTCGGATTTGAAAAGACATCATAGTGAAGAACAAATTGGTTTTCCGTAGCGATTTG    3780
PMP1611      ATCTGTCGGATTTGAAAAGACATCATAGTGAAGAACAAATTGGTTTTCCGTAGCGATTTG    3780
             ************************************************************

PMP1610      AAGATTATAAGCAGCCTTGAGTTGACCATTTTTCATATGATCTTCCTTCATCCGCATAAA    3840
PMP1666      AAGATTATAAGCAGCCTTGAGTTGACCATTTTTCATATGATCTTCCTTCATCCGCATAAA    3840
PMP1665      AAGATTATAAGCAGCCTTGAGTTGACCATTTTTCATATGATCTTCCTTCATCCGCATAAA    3840
PMP1667      AAGATTATAAGCAGCCTTGAGTTGACCATTTTTCATATGATCTTCCTTCATCCGCATAAA    3840
PMP1663      AAGATTATAAGCAGCCTTGAGTTGACCATTTTTCATATGATCTTCCTTCATCCGCATAAA    3840
PMP1664      AAGATTATAAGCAGCCTTGAGTTGACCATTTTTCATATGATCTTCCTTCATCCGCATAAA    3840
PMP1662      AAGATTATAAGCAGCCTTGAGTTGACCATTTTTCATATGATCTTCCTTCATCCGCATAAA    3840
PMP1611      AAGATTATAAGCAGCCTTGAGTTGACCATTTTTCATATGATCTTCCTTCATCCGCATAAA    3840
             ************************************************************

PMP1610      AGTGGCATCTGGATCGGTTTTGGAAAAACTGTTAGGCCCTTGAAATGTCTCCTGATAGTT    3900
PMP1666      AGTGGCATCTGGATCGGTTTTGGAAAAACTGTTAGGCCCTTGAAATGTCTCCTGATAGTT    3900
PMP1665      AGTGGCATCTGGATCGGTTTTGGAAAAACTGTTAGGCCCTTGAAATGTCTCCTGATAGTT    3900
PMP1667      AGTGGCATCTGGATCGGTTTTGGAAAAACTGTTAGGCCCTTGAAATGTCTCCTGATAGTT    3900
PMP1663      AGTGGCATCTGGATCGGTTTTGGAAAAACTGTTAGGCCCTTGAAATGTCTCCTGATAGTT    3900
PMP1664      AGTGGCATCTGGATCGGTTTTGGAAAAACTGTTAGGCCCTTGAAATGTCTCCTGATAGTT    3900
PMP1662      AGTGGCATCTGGATCGGTTTTGGAAAAACTGTTAGGCCCTTGAAATGTCTCCTGATAGTT    3900
PMP1611      AGTGGCATCTGGATCGGTTTTGGAAAAACTGTTAGGCCCTTGAAATGTCTCCTGATAGTT    3900
```

```
            ************************************************************

PMP1610     CTCATATTTTTCAGCACGTACTGAAAAATCCTCCTTTAAACTCTTCATAATCCTTTTTAT    3960
PMP1666     CTCATATTTTTCAGCACGTACTGAAAAATCCTCCTTTAAACTCTTCATAATCCTTTTTAT    3960
PMP1665     CTCATATTTTTCAGCACGTACTGAAAAATCCTCCTTTAAACTCTTCATAATCCTTTTTAT    3960
PMP1667     CTCATATTTTTCAGCACGTACTGAAAAATCCTCCTTTAAACTCTTCATAATCCTTTTTAT    3960
PMP1663     CTCATATTTTTCAGCACGTACTGAAAAATCCTCCTTTAAACTCTTCATAATCCTTTTTAT    3960
PMP1664     CTCATATTTTTCAGCACGTACTGAAAAATCCTCCTTTAAACTCTTCATAATCCTTTTTAT    3960
PMP1662     CTCATATTTTTCAGCACGTACTGAAAAATCCTCCTTTAAACTCTTCATAATCCTTTTTAT    3960
PMP1611     CTCATATTTTTCAGCACGTACTGAAAAATCCTCCTTTAAACTCTTCATAATCCTTTTTAT    3960
            ************************************************************

PMP1610     TTTTATGAAGATATTGTTTGAAAGATGTGAGTTTCCACGGATGGGTTTGTGGAGGGATAT    4020
PMP1666     TTTTATGAAGATATTGTTTGAAAGATGTGAGTTTCCACGGATGGGTTTGTGGAGGGATAT    4020
PMP1665     TTTTATGAAGATATTGTTTGAAAGATGTGAGTTTCCACGGATGGGTTTGTGGAGGGATAT    4020
PMP1667     TTTTATGAAGATATTGTTTGAAAGATGTGAGTTTCCACGGATGGGTTTGTGGAGGGATAT    4020
PMP1663     TTTTATGAAGATATTGTTTGAAAGATGTGAGTTTCCACGGATGGGTTTGTGGAGGGATAT    4020
PMP1664     TTTTATGAAGATATTGTTTGAAAGATGTGAGTTTCCACGGATGGGTTTGTGGAGGGATAT    4020
PMP1662     TTTTATGAAGATATTGTTTGAAAGATGTGAGTTTCCACGGATGGGTTTGTGGAGGGATAT    4020
PMP1611     TTTTATGAAGATATTGTTTGAAAGATGTGAGTTTCCACGGATGGGTTTGTGGAGGGATAT    4020
            ************************************************************

PMP1610     ACTTGCGTCTTTCTTTTTTTTGTTCTGGTTCTTGTTCAAAGTTTTTCGAATAGAGTTCAT    4080
PMP1666     ACTTGCGTCTTTCTTTTTTTTGTTCTGGTTCTTGTTCAAAGTTTTTCGAATAGAGTTCAT    4080
PMP1665     ACTTGCGTCTTTCTTTTTTTTGTTCTGGTTCTTGTTCAAAGTTTTTCGAATAGAGTTCAT    4080
PMP1667     ACTTGCGTCTTTCTTTTTTTTGTTCTGGTTCTTGTTCAAAGTTTTTCGAATAGAGTTCAT    4080
PMP1663     ACTTGCGTCTTTCTTTTTTTTGTTCTGGTTCTTGTTCAAAGTTTTTCGAATAGAGTTCAT    4080
PMP1664     ACTTGCGTCTTTCTTTTTTTTGTTCTGGTTCTTGTTCAAAGTTTTTCGAATAGAGTTCAT    4080
PMP1662     ACTTGCGTCTTTCTTTTTTTTGTTCTGGTTCTTGTTCAAAGTTTTTCGAATAGAGTTCAT    4080
PMP1611     ACTTGCGTCTTTCTTTTTTTTGTTCTGGTTCTTGTTCAAAGTTTTTCGAATAGAGTTCAT    4080
            ************************************************************

PMP1610     GATTTAGTAGCTCCTTTGTGTGATAGATTTTGTCAGCGATATTGAGGTAGATGTCACCAT    4140
PMP1666     GATTTAGTAGCTCCTTTGTGTGATAGATTTTGTCAGCGATATTGAGGTAGATGTCACCAT    4140
PMP1665     GATTTAGTAGCTCCTTTGTGTGATAGATTTTGTCAGCGATATTGAGGTAGATGTCACCAT    4140
PMP1667     GATTTAGTAGCTCCTTTGTGTGATAGATTTTGTCAGCGATATTGAGGTAGATGTCACCAT    4140
```

```
PMP1663      GATTTAGTAGCTCCTTTGTGTGATAGATTTTGTCAGCGATATTGAGGTAGATGTCACCAT   4140
PMP1664      GATTTAGTAGCTCCTTTGTGTGATAGATTTTGTCAGCGATATTGAGGTAGATGTCACCAT   4140
PMP1662      GATTTAGTAGCTCCTTTGTGTGATAGATTTTGTCAGCGATATTGAGGTAGATGTCACCAT   4140
PMP1611      GATTTAGTAGCTCCTTTGTGTGATAGATTTTGTCAGCGATATTGAGGTAGATGTCACCAT   4140
             ************************************************************

PMP1610      CAAATGCTTTTATAACTAATGCTTTTGTCTTTCTGATGAAATAGACTTCTTTTCCTTGCT   4200
PMP1666      CAAATGCTTTTATAACTAATGCTTTTGTCTTTCTGATGAAATAGACTTCTTTTCCTTGCT   4200
PMP1665      CAAATGCTTTTATAACTAATGCTTTTGTCTTTCTGATGAAATAGACTTCTTTTCCTTGCT   4200
PMP1667      CAAATGCTTTTATAACTAATGCTTTTGTCTTTCTGATGAAATAGACTTCTTTTCCTTGCT   4200
PMP1663      CAAATGCTTTTATAACTAATGCTTTTGTCTTTCTGATGAAATAGACTTCTTTTCCTTGCT   4200
PMP1664      CAAATGCTTTTATAACTAATGCTTTTGTCTTTCTGATGAAATAGACTTCTTTTCCTTGCT   4200
PMP1662      CAAATGCTTTTATAACTAATGCTTTTGTCTTTCTGATGAAATAGACTTCTTTTCCTTGCT   4200
PMP1611      CAAATGCTTTTATAACTAATGCTTTTGTCTTTCTGATGAAATAGACTTCTTTTCCTTGCT   4200
             ************************************************************

PMP1610      CGGTAGGGATATAGCAACGATTTTGGAATCGGATATGGTGTCCACTATCGACGACTCTCT   4260
PMP1666      CGGTAGGGATATAGCAACGATTTTGGAATCGGATATGGTGTCCACTATCGACGACTCTCT   4260
PMP1665      CGGTAGGGATATAGCAACGATTTTGGAATCGGATATGGTGTCCACTATCGACGACTCTCT   4260
PMP1667      CGGTAGGGATATAGCAACGATTTTGGAATCGGATATGGTGTCCACTATCGACGACTCTCT   4260
PMP1663      CGGTAGGGATATAGCAACGATTTTGGAATCGGATATGGTGTCCACTATCGACGACTCTCT   4260
PMP1664      CGGTAGGGATATAGCAACGATTTTGGAATCGGATATGGTGTCCACTATCGACGACTCTCT   4260
PMP1662      CGGTAGGGATATAGCAACGATTTTGGAATCGGATATGGTGTCCACTATCGACGACTCTCT   4260
PMP1611      CGGTAGGGATATAGCAACGATTTTGGAATCGGATATGGTGTCCACTATCGACGACTCTCT   4260
             ************************************************************

PMP1610      CCCGCCAGTCTAGCTAGAATGAGATTTCGTTCAGAGGGCTTAGGAGCCTCCTCAAAGACA   4320
PMP1666      CCCGCCAGTCTAGCTAGAATGAGATTTCGTTCAGAGGGCTTAGGAGCCTCCTCAAAGACA   4320
PMP1665      CCCGCCAGTCTAGCTAGAATGAGATTTCGTTCAGAGGGCTTAGGAGCCTCCTCAAAGACA   4320
PMP1667      CCCGCCAGTCTAGCTAGAATGAGATTTCGTTCAGAGGGCTTAGGAGCCTCCTCAAAGACA   4320
PMP1663      CCCGCCAGTCTAGCTAGAATGAGATTTCGTTCAGAGGGCTTAGGAGCCTCCTCAAAGACA   4320
PMP1664      CCCGCCAGTCTAGCTAGAATGAGATTTCGTTCAGAGGGCTTAGGAGCCTCCTCAAAGACA   4320
PMP1662      CCCGCCAGTCTAGCTAGAATGAGATTTCGTTCAGAGGGCTTAGGAGCCTCCTCAAAGACA   4320
PMP1611      CCCGCCAGTCTAGCTAGAATGAGATTTCGTTCAGAGGGCTTAGGAGCCTCCTCAAAGACA   4320
             ************************************************************
```

```
PMP1610      GAGAGTTTTGTCTTGTTTCCAAACTGTTCATTAAAGGTTTGGATATAGGAAGGTAGAAAG   4380
PMP1666      GAGAGTTTTGTCTTGTTTCCAAACTGTTCATTAAAGGTTTGGATATAGGAAGGTAGAAAG   4380
PMP1665      GAGAGTTTTGTCTTGTTTCCAAACTGTTCATTAAAGGTTTGGATATAGGAAGGTAGAAAG   4380
PMP1667      GAGAGTTTTGTCTTGTTTCCAAACTGTTCATTAAAGGTTTGGATATAGGAAGGTAGAAAG   4380
PMP1663      GAGAGTTTTGTCTTGTTTCCAAACTGTTCATTAAAGGTTTGGATATAGGAAGGTAGAAAG   4380
PMP1664      GAGAGTTTTGTCTTGTTTCCAAACTGTTCATTAAAGGTTTGGATATAGGAAGGTAGAAAG   4380
PMP1662      GAGAGTTTTGTCTTGTTTCCAAACTGTTCATTAAAGGTTTGGATATAGGAAGGTAGAAAG   4380
PMP1611      GAGAGTTTTGTCTTGTTTCCAAACTGTTCATTAAAGGTTTGGATATAGGAAGGTAGAAAG   4380
             ************************************************************

PMP1610      GTATTGGCTTCTTCCAAGGTATGAATATTGTTTCGTTCCAGTTCGATAGGCAGGCGAGAT   4440
PMP1666      GTATTGGCTTCTTCCAAGGTATGAATATTGTTTCGTTCCAGTTCGATAGGCAGGCGAGAT   4440
PMP1665      GTATTGGCTTCTTCCAAGGTATGAATATTGTTTCGTTCCAGTTCGATAGGCAGGCGAGAT   4440
PMP1667      GTATTGGCTTCTTCCAAGGTATGAATATTGTTTCGTTCCAGTTCGATAGGCAGGCGAGAT   4440
PMP1663      GTATTGGCTTCTTCCAAGGTATGAATATTGTTTCGTTCCAGTTCGATAGGCAGGCGAGAT   4440
PMP1664      GTATTGGCTTCTTCCAAGGTATGAATATTGTTTCGTTCCAGTTCGATAGGCAGGCGAGAT   4440
PMP1662      GTATTGGCTTCTTCCAAGGTATGAATATTGTTTCGTTCCAGTTCGATAGGCAGGCGAGAT   4440
PMP1611      GTATTGGCTTCTTCCAAGGTATGAATATTGTTTCGTTCCAGTTCGATAGGCAGGCGAGAT   4440
             ************************************************************

PMP1610      TGTAGTGTCTGATTGAGTCTTTCGACTCTCCCTTTAGCTTGAGGGATAGAGGTTGTCTCA   4500
PMP1666      TGTAGTGTCTGATTGAGTCTTTCGACTCTCCCTTTAGCTTGAGGGATAGAGGTTGTCTCA   4500
PMP1665      TGTAGTGTCTGATTGAGTCTTTCGACTCTCCCTTTAGCTTGAGGGATAGAGGTTGTCTCA   4500
PMP1667      TGTAGTGTCTGATTGAGTCTTTCGACTCTCCCTTTAGCTTGAGGGATAGAGGTTGTCTCA   4500
PMP1663      TGTAGTGTCTGATTGAGTCTTTCGACTCTCCCTTTAGCTTGAGGGATAGAGGTTGTCTCA   4500
PMP1664      TGTAGTGTCTGATTGAGTCTTTCGACTCTCCCTTTAGCTTGAGGGATAGAGGTTGTCTCA   4500
PMP1662      TGTAGTGTCTGATTGAGTCTTTCGACTCTCCCTTTAGCTTGAGGGATAGAGGTTGTCTCA   4500
PMP1611      TGTAGTGTCTGATTGAGTCTTTCGACTCTCCCTTTAGCTTGAGGGATAGAGGTTGTCTCA   4500
             ************************************************************

PMP1610      AGGAGAATTCCCAGTTGGTGACAGGCGTATCCAAATTGTGTATGGGTGTCGTCCTCCATT   4560
PMP1666      AGGAGAATTCCCAGTTGGTGACAGGCGTATCCAAATTGTGTATGGGTGTCGTCCTCCATT   4560
PMP1665      AGGAGAATTCCCAGTTGGTGACAGGCGTATCCAAATTGTGTATGGGTGTCGTCCTCCATT   4560
PMP1667      AGGAGAATTCCCAGTTGGTGACAGGCGTATCCAAATTGTGTATGGGTGTCGTCCTCCATT   4560
PMP1663      AGGAGAATTCCCAGTTGGTGACAGGCGTATCCAAATTGTGTATGGGTGTCGTCCTCCATT   4560
PMP1664      AGGAGAATTCCCAGTTGGTGACAGGCGTATCCAAATTGTGTATGGGTGTCGTCCTCCATT   4560
```

```
PMP1662      AGGAGAATTCCCAGTTGGTGACAGGCGTATCCAAATTGTGTATGGGTGTCGTCCTCCATT   4560
PMP1611      AGGAGAATTCCCAGTTGGTGACAGGCGTATCCAAATTGTGTATGGGTGTCGTCCTCCATT   4560
             ************************************************************

PMP1610      TTCTTAGAGTTGGAGGCTTGATAGGTAAAAACAGTTCTCTTATCTGTTTTAATTTGAAGA   4620
PMP1666      TTCTTAGAGTTGGAGGCTTGATAGGTAAAAACAGTTCTCTTATCTGTTTTAATTTGAAGA   4620
PMP1665      TTCTTAGAGTTGGAGGCTTGATAGGTAAAAACAGTTCTCTTATCTGTTTTAATTTGAAGA   4620
PMP1667      TTCTTAGAGTTGGAGGCTTGATAGGTAAAAACAGTTCTCTTATCTGTTTTAATTTGAAGA   4620
PMP1663      TTCTTAGAGTTGGAGGCTTGATAGGTAAAAACAGTTCTCTTATCTGTTTTAATTTGAAGA   4620
PMP1664      TTCTTAGAGTTGGAGGCTTGATAGGTAAAAACAGTTCTCTTATCTGTTTTAATTTGAAGA   4620
PMP1662      TTCTTAGAGTTGGAGGCTTGATAGGTAAAAACAGTTCTCTTATCTGTTTTAATTTGAAGA   4620
PMP1611      TTCTTAGAGTTGGAGGCTTGATAGGTAAAAACAGTTCTCTTATCTGTTTTAATTTGAAGA   4620
             ************************************************************

PMP1610      GGAATGCCGTGGTTGGCTAAGATTTGTTCGAGGACATGATAGTAAGCATTCAAGGTCTCT   4680
PMP1666      GGAATGCCGTGGTTGGCTAAGATTTGTTCGAGGACATGATAGTAAGCATTCAAGGTCTCT   4680
PMP1665      GGAATGCCGTGGTTGGCTAAGATTTGTTCGAGGACATGATAGTAAGCATTCAAGGTCTCT   4680
PMP1667      GGAATGCCGTGGTTGGCTAAGATTTGTTCGAGGACATGATAGTAAGCATTCAAGGTCTCT   4680
PMP1663      GGAATGCCGTGGTTGGCTAAGATTTGTTCGAGGACATGATAGTAAGCATTCAAGGTCTCT   4680
PMP1664      GGAATGCCGTGGTTGGCTAAGATTTGTTCGAGGACATGATAGTAAGCATTCAAGGTCTCT   4680
PMP1662      GGAATGCCGTGGTTGGCTAAGATTTGTTCGAGGACATGATAGTAAGCATTCAAGGTCTCT   4680
PMP1611      GGAATGCCGTGGTTGGCTAAGATTTGTTCGAGGACATGATAGTAAGCATTCAAGGTCTCT   4680
             ************************************************************

PMP1610      TGTTTGTCAAAATAAGCGCCTAGGATATTGCCAGAAGCATCATCAATGGCTAGGTGTAAG   4740
PMP1666      TGTTTGTCAAAATAAGCGCCTAGGATATTGCCAGAAGCATCATCAATGGCTAGGTGTAAG   4740
PMP1665      TGTTTGTCAAAATAAGCGCCTAGGATATTGCCAGAAGCATCATCAATGGCTAGGTGTAAG   4740
PMP1667      TGTTTGTCAAAATAAGCGCCTAGGATATTGCCAGAAGCATCATCAATGGCTAGGTGTAAG   4740
PMP1663      TGTTTGTCAAAATAAGCGCCTAGGATATTGCCAGAAGCATCATCAATGGCTAGGTGTAAG   4740
PMP1664      TGTTTGTCAAAATAAGCGCCTAGGATATTGCCAGAAGCATCATCAATGGCTAGGTGTAAG   4740
PMP1662      TGTTTGTCAAAATAAGCGCCTAGGATATTGCCAGAAGCATCATCAATGGCTAGGTGTAAG   4740
PMP1611      TGTTTGTCAAAATAAGCGCCTAGGATATTGCCAGAAGCATCATCAATGGCTAGGTGTAAG   4740
             ************************************************************

PMP1610      TTAGAGGTTTCTACTCCAAACCAGGCATGAGGGCTGGCATCCATTTGGATGAGTTCTCCA   4800
PMP1666      TTAGAGGTTTCTACTCCAAACCAGGCATGAGGGCTGGCATCCATTTGGATGAGTTCTCCA   4800
```

```
PMP1665      TTAGAGGTTTCTACTCCAAACCAGGCATGAGGGCTGGCATCCATTTGGATGAGTTCTCCA    4800
PMP1667      TTAGAGGTTTCTACTCCAAACCAGGCATGAGGGCTGGCATCCATTTGGATGAGTTCTCCA    4800
PMP1663      TTAGAGGTTTCTACTCCAAACCAGGCATGAGGGCTGGCATCCATTTGGATGAGTTCTCCA    4800
PMP1664      TTAGAGGTTTCTACTCCAAACCAGGCATGAGGGCTGGCATCCATTTGGATGAGTTCTCCA    4800
PMP1662      TTAGAGGTTTCTACTCCAAACCAGGCATGAGGGCTGGCATCCATTTGGATGAGTTCTCCA    4800
PMP1611      TTAGAGGTTTCTACTCCAAACCAGGCATGAGGGCTGGCATCCATTTGGATGAGTTCTCCA    4800
             ************************************************************

PMP1610      GCAAATTTTTTTCTGGGTCTACTAGGATGTACCTTTTTAGGGTCTTCCAGGAAGTTTTCA    4860
PMP1666      GCAAATTTTTTTCTGGGTCTACTAGGATGTACCTTTTTAGGGTCTTCCAGGAAGTTTTCA    4860
PMP1665      GCAAATTTTTTTCTGGGTCTACTAGGATGTACCTTTTTAGGGTCTTCCAGGAAGTTTTCA    4860
PMP1667      GCAAATTTTTTTCTGGGTCTACTAGGATGTACCTTTTTAGGGTCTTCCAGGAAGTTTTCA    4860
PMP1663      GCAAATTTTTTTCTGGGTCTACTAGGATGTACCTTTTTAGGGTCTTCCAGGAAGTTTTCA    4860
PMP1664      GCAAATTTTTTTCTGGGTCTACTAGGATGTACCTTTTTAGGGTCTTCCAGGAAGTTTTCA    4860
PMP1662      GCAAATTTTTTTCTGGGTCTACTAGGATGTACCTTTTTAGGGTCTTCCAGGAAGTTTTCA    4860
PMP1611      GCAAATTTTTTTCTGGGTCTACTAGGATGTACCTTTTTAGGGTCTTCCAGGAAGTTTTCA    4860
             ************************************************************

PMP1610      GCTGTCGGTAAGATTGGATTGTCTAGGGGTTGCTTGGGGTTCAGTTTAGCTTGTTTTCTT    4920
PMP1666      GCTGTCGGTAAGATTGGATTGTCTAGGGGTTGCTTGGGGTTCAGTTTAGCTTGTTTTCTT    4920
PMP1665      GCTGTCGGTAAGATTGGATTGTCTAGGGGTTGCTTGGGGTTCAGTTTAGCTTGTTTTCTT    4920
PMP1667      GCTGTCGGTAAGATTGGATTGTCTAGGGGTTGCTTGGGGTTCAGTTTAGCTTGTTTTCTT    4920
PMP1663      GCTGTCGGTAAGATTGGATTGTCTAGGGGTTGCTTGGGGTTCAGTTTAGCTTGTTTTCTT    4920
PMP1664      GCTGTCGGTAAGATTGGATTGTCTAGGGGTTGCTTGGGGTTCAGTTTAGCTTGTTTTCTT    4920
PMP1662      GCTGTCGGTAAGATTGGATTGTCTAGGGGTTGCTTGGGGTTCAGTTTAGCTTGTTTTCTT    4920
PMP1611      GCTGTCGGTAAGATTGGATTGTCTAGGGGTTGCTTGGGGTTCAGTTTAGCTTGTTTTCTT    4920
             ************************************************************

PMP1610      ACTCTCTTCTTTGTCTTTCTGTGAGACTTAGGCGACAGGATATTTTTCTTATAGAGTATT    4980
PMP1666      ACTCTCTTCTTTGTCTTTCTGTGAGACTTAGGCGACAGGATATTTTTCTTATAGAGTATT    4980
PMP1665      ACTCTCTTCTTTGTCTTTCTGTGAGACTTAGGCGACAGGATATTTTTCTTATAGAGTATT    4980
PMP1667      ACTCTCTTCTTTGTCTTTCTGTGAGACTTAGGCGACAGGATATTTTTCTTATAGAGTATT    4980
PMP1663      ACTCTCTTCTTTGTCTTTCTGTGAGACTTAGGCGACAGGATATTTTTCTTATAGAGTATT    4980
PMP1664      ACTCTCTTCTTTGTCTTTCTGTGAGACTTAGGCGACAGGATATTTTTCTTATAGAGTATT    4980
PMP1662      ACTCTCTTCTTTGTCTTTCTGTGAGACTTAGGCGACAGGATATTTTTCTTATAGAGTATT    4980
PMP1611      ACTCTCTTCTTTGTCTTTCTGTGAGACTTAGGCGACAGGATATTTTTCTTATAGAGTATT    4980
```

```
            ************************************************************

PMP1610     TTTCTAACAGTTGTATCAGAGAGCTGAATTCCTTCTTCTTCAGCTAGCAATTCACAGAAA   5040
PMP1666     TTTCTAACAGTTGTATCAGAGAGCTGAATTCCTTCTTCTTCAGCTAGCAATTCACAGAAA   5040
PMP1665     TTTCTAACAGTTGTATCAGAGAGCTGAATTCCTTCTTCTTCAGCTAGCAATTCACAGAAA   5040
PMP1667     TTTCTAACAGTTGTATCAGAGAGCTGAATTCCTTCTTCTTCAGCTAGCAATTCACAGAAA   5040
PMP1663     TTTCTAACAGTTGTATCAGAGAGCTGAATTCCTTCTTCTTCAGCTAGCAATTCACAGAAA   5040
PMP1664     TTTCTAACAGTTGTATCAGAGAGCTGAATTCCTTCTTCTTCAGCTAGCAATTCACAGAAA   5040
PMP1662     TTTCTAACAGTTGTATCAGAGAGCTGAATTCCTTCTTCTTCAGCTAGCAATTCACAGAAA   5040
PMP1611     TTTCTAACAGTTGTATCAGAGAGCTGAATTCCTTCTTCTTCAGCTAGCAATTCACAGAAA   5040
            ************************************************************

PMP1610     TGACGGACATTTGGTTTATATGTTTGATAGGAGAGGTATTTCTTTAGGATACGTTCTTTG   5100
PMP1666     TGACGGACATTTGGTTTATATGTTTGATAGGAGAGGTATTTCTTTAGGATACGTTCTTTG   5100
PMP1665     TGACGGACATTTGGTTTATATGTTTGATAGGAGAGGTATTTCTTTAGGATACGTTCTTTG   5100
PMP1667     TGACGGACATTTGGTTTATATGTTTGATAGGAGAGGTATTTCTTTAGGATACGTTCTTTG   5100
PMP1663     TGACGGACATTTGGTTTATATGTTTGATAGGAGAGGTATTTCTTTAGGATACGTTCTTTG   5100
PMP1664     TGACGGACATTTGGTTTATATGTTTGATAGGAGAGGTATTTCTTTAGGATACGTTCTTTG   5100
PMP1662     TGACGGACATTTGGTTTATATGTTTGATAGGAGAGGTATTTCTTTAGGATACGTTCTTTG   5100
PMP1611     TGACGGACATTTGGTTTATATGTTTGATAGGAGAGGTATTTCTTTAGGATACGTTCTTTG   5100
            ************************************************************

PMP1610     ATTTCATCAGGGATTGCATGTTTTGGTTTTCGATTTCTGTTTCCGTGTCTGAAGGCTTCT   5160
PMP1666     ATTTCATCAGGGATTGCATGTTTTGGTTTTCGATTTCTGTTTCCGTGTCTGAAGGCTTCT   5160
PMP1665     ATTTCATCAGGGATTGCATGTTTTGGTTTTCGATTTCTGTTTCCGTGTCTGAAGGCTTCT   5160
PMP1667     ATTTCATCAGGGATTGCATGTTTTGGTTTTCGATTTCTGTTTCCGTGTCTGAAGGCTTCT   5160
PMP1663     ATTTCATCAGGGATTGCATGTTTTGGTTTTCGATTTCTGTTTCCGTGTCTGAAGGCTTCT   5160
PMP1664     ATTTCATCAGGGATTGCATGTTTTGGTTTTCGATTTCTGTTTCCGTGTCTGAAGGCTTCT   5160
PMP1662     ATTTCATCAGGGATTGCATGTTTTGGTTTTCGATTTCTGTTTCCGTGTCTGAAGGCTTCT   5160
PMP1611     ATTTCATCAGGGATTGCATGTTTTGGTTTTCGATTTCTGTTTCCGTGTCTGAAGGCTTCT   5160
            ************************************************************

PMP1610     TTTTCTTTCTGTTGATAGGCTAGTAGCAGACGATTGATTTGTCTTTCAGAAAGATTGAGT   5220
PMP1666     TTTTCTTTCTGTTGATAGGCTAGTAGCAGACGATTGATTTGTCTTTCAGAAAGATTGAGT   5220
PMP1665     TTTTCTTTCTGTTGATAGGCTAGTAGCAGACGATTGATTTGTCTTTCAGAAAGATTGAGT   5220
PMP1667     TTTTCTTTCTGTTGATAGGCTAGTAGCAGACGATTGATTTGTCTTTCAGAAAGATTGAGT   5220
```

```
PMP1663      TTTTCTTTCTGTTGATAGGCTAGTAGCAGACGATTGATTTGTCTTTCAGAAAGATTGAGT   5220
PMP1664      TTTTCTTTCTGTTGATAGGCTAGTAGCAGACGATTGATTTGTCTTTCAGAAAGATTGAGT   5220
PMP1662      TTTTCTTTCTGTTGATAGGCTAGTAGCAGACGATTGATTTGTCTTTCAGAAAGATTGAGT   5220
PMP1611      TTTTCTTTCTGTTGATAGGCTAGTAGCAGACGATTGATTTGTCTTTCAGAAAGATTGAGT   5220
             ************************************************************

PMP1610      TCGACACAGGCCCGTTTCTTTGTTTTCTTTCCTTGGGCTATAGCTTTTATCACAAGATAT   5280
PMP1666      TCGACACAGGCCCGTTTCTTTGTTTTCTTTCCTTGGGCTATAGCTTTTATCACAAGATAT   5280
PMP1665      TCGACACAGGCCCGTTTCTTTGTTTTCTTTCCTTGGGCTATAGCTTTTATCACAAGATAT   5280
PMP1667      TCGACACAGGCCCGTTTCTTTGTTTTCTTTCCTTGGGCTATAGCTTTTATCACAAGATAT   5280
PMP1663      TCGACACAGGCCCGTTTCTTTGTTTTCTTTCCTTGGGCTATAGCTTTTATCACAAGATAT   5280
PMP1664      TCGACACAGGCCCGTTTCTTTGTTTTCTTTCCTTGGGCTATAGCTTTTATCACAAGATAT   5280
PMP1662      TCGACACAGGCCCGTTTCTTTGTTTTCTTTCCTTGGGCTATAGCTTTTATCACAAGATAT   5280
PMP1611      TCGACACAGGCCCGTTTCTTTGTTTTCTTTCCTTGGGCTATAGCTTTTATCACAAGATAT   5280
             ************************************************************

PMP1610      TTTTTCGTTTCATTCATATTCAGTTGGATCCTTTTCATATGACTATTCTACCAAATGGGA   5340
PMP1666      TTTTTCGTTTCATTCATATTCAGTTGGATCCTTTTCATATGACTATTCTACCAAATGGGA   5340
PMP1665      TTTTTCGTTTCATTCATATTCAGTTGGATCCTTTTCATATGACTATTCTACCAAATGGGA   5340
PMP1667      TTTTTCGTTTCATTCATATTCAGTTGGATCCTTTTCATATGACTATTCTACCAAATGGGA   5340
PMP1663      TTTTTCGTTTCATTCATATTCAGTTGGATCCTTTTCATATGACTATTCTACCAAATGGGA   5340
PMP1664      TTTTTCGTTTCATTCATATTCAGTTGGATCCTTTTCATATGACTATTCTACCAAATGGGA   5340
PMP1662      TTTTTCGTTTCATTCATATTCAGTTGGATCCTTTTCATATGACTATTCTACCAAATGGGA   5340
PMP1611      TTTTTCGTTTCATTCATATTCAGTTGGATCCTTTTCATATGACTATTCTACCAAATGGGA   5340
             ************************************************************

PMP1610      CATTTTTACGTTCGGTTTACTAAAGACATTATCACATTCGAATTACACAAGATGCAGATA   5400
PMP1666      CATTTTTACGTTCGGTTTACTAAAGACATTATCACATTCGAATTACACAAGATGCAGATA   5400
PMP1665      CATTTTTACGTTCGGTTTACTAAAGACATTATCACATTCGAATTACACAAGATGCAGATA   5400
PMP1667      CATTTTTACGTTCGGTTTACTAAAGACATTATCACATTCGAATTACACAAGATGCAGATA   5400
PMP1663      CATTTTTACGTTCGGTTTACTAAAGACATTATCACATTCGAATTACACAAGATGCAGATA   5400
PMP1664      CATTTTTACGTTCGGTTTACTAAAGACATTATCACATTCGAATTACACAAGATGCAGATA   5400
PMP1662      CATTTTTACGTTCGGTTTACTAAAGACATTATCACATTCGAATTACACAAGATGCAGATA   5400
PMP1611      CATTTTTACGTTCGGTTTACTAAAGACATTATCACATTCGAATTACACAAGATGCAGATA   5400
             ************************************************************
```

```
PMP1610      GTGAAAGAAAGGTGTAGACATTACCGTAAAAAAGTGATACAATTATAAGATGTTCAATGT   5460
PMP1666      GTGAAAGAAAGGTGTAGACATTACCGTAAAAAAGTGATACAATTATAAGATGTTCAATGT   5460
PMP1665      GTGAAAGAAAGGTGTAGACATTACCGTAAAAAAGTGATACAATTATAAGATGTTCAATGT   5460
PMP1667      GTGAAAGAAAGGTGTAGACATTACCGTAAAAAAGTGATACAATTATAAGATGTTCAATGT   5460
PMP1663      GTGAAAGAAAGGTGTAGACATTACCGTAAAAAAGTGATACAATTATAAGATGTTCAATGT   5460
PMP1664      GTGAAAGAAAGGTGTAGACATTACCGTAAAAAAGTGATACAATTATAAGATGTTCAATGT   5460
PMP1662      GTGAAAGAAAGGTGTAGACATTACCGTAAAAAAGTGATACAATTATAAGATGTTCAATGT   5460
PMP1611      GTGAAAGAAAGGTGTAGACATTACCGTAAAAAAGTGATACAATTATAAGATGTTCAATGT   5460
             ************************************************************

PMP1610      ATAGGTGTTAATCATGAGTAGACGTTTTAAGAAATCAGGTTCACAGAAAGTGAAGCGAAG   5520
PMP1666      ATAGGTGTTAATCATGAGTAGACGTTTTAAGAAATCAGGTTCACAGAAAGTGAAGCGAAG   5520
PMP1665      ATAGGTGTTAATCATGAGTAGACGTTTTAAGAAATCAGGTTCACAGAAAGTGAAGCGAAG   5520
PMP1667      ATAGGTGTTAATCATGAGTAGACGTTTTAAGAAATCAGGTTCACAGAAAGTGAAGCGAAG   5520
PMP1663      ATAGGTGTTAATCATGAGTAGACGTTTTAAGAAATCAGGTTCACAGAAAGTGAAGCGAAG   5520
PMP1664      ATAGGTGTTAATCATGAGTAGACGTTTTAAGAAATCAGGTTCACAGAAAGTGAAGCGAAG   5520
PMP1662      ATAGGTGTTAATCATGAGTAGACGTTTTAAGAAATCAGGTTCACAGAAAGTGAAGCGAAG   5520
PMP1611      ATAGGTGTTAATCATGAGTAGACGTTTTAAGAAATCAGGTTCACAGAAAGTGAAGCGAAG   5520
             ************************************************************

PMP1610      TGTTAATATAGTTTTGTTGACTATTTATTTATTGTTAGTTTGTTTTTTATTGTTCTTAAT   5580
PMP1666      TGTTAATATAGTTTTGTTGACTATTTATTTATTGTTAGTTTGTTTTTTATTGTTCTTAAT   5580
PMP1665      TGTTAATATAGTTTTGTTGACTATTTATTTATTGTTAGTTTGTTTTTTATTGTTCTTAAT   5580
PMP1667      TGTTAATATAGTTTTGTTGACTATTTATTTATTGTTAGTTTGTTTTTTATTGTTCTTAAT   5580
PMP1663      TGTTAATATAGTTTTGTTGACTATTTATTTATTGTTAGTTTGTTTTTTATTGTTCTTAAT   5580
PMP1664      TGTTAATATAGTTTTGTTGACTATTTATTTATTGTTAGTTTGTTTTTTATTGTTCTTAAT   5580
PMP1662      TGTTAATATAGTTTTGTTGACTATTTATTTATTGTTAGTTTGTTTTTTATTGTTCTTAAT   5580
PMP1611      TGTTAATATAGTTTTGTTGACTATTTATTTATTGTTAGTTTGTTTTTTATTGTTCTTAAT   5580
             ************************************************************

PMP1610      CTTTAAGTACAATATCCTTGCTTTTAGATATCTTAATCTAGTGGCAACTGCCTTTGTTCT   5640
PMP1666      CTTTAAGTACAATATCCTTGCTTTTAGATATCTTAATCTAGTGGCAACTGCCTTTGTTCT   5640
PMP1665      CTTTAAGTACAATATCCTTGCTTTTAGATATCTTAATCTAGTGGCAACTGCCTTTGTTCT   5640
PMP1667      CTTTAAGTACAATATCCTTGCTTTTAGATATCTTAATCTAGTGGCAACTGCCTTTGTTCT   5640
PMP1663      CTTTAAGTACAATATCCTTGCTTTTAGATATCTTAATCTAGTGGCAACTGCCTTTGTTCT   5640
PMP1664      CTTTAAGTACAATATCCTTGCTTTTAGATATCTTAATCTAGTGGCAACTGCCTTTGTTCT   5640
```

```
PMP1662      CTTTAAGTACAATATCCTTGCTTTTAGATATCTTAATCTAGTGGCAACTGCCTTTGTTCT   5640
PMP1611      CTTTAAGTACAATATCCTTGCTTTTAGATATCTTAATCTAGTGGCAACTGCCTTTGTTCT   5640
             ************************************************************

PMP1610      ACTAGTTGCCTTAATAGGGCTACTCTCGATTATCTATAAAAAAGCTGAAAAGTTTACCAT   5700
PMP1666      ACTAGTTGCCTTAATAGGGCTACTCTCGATTATCTATAAAAAAGCTGAAAAGTTTACCAT   5700
PMP1665      ACTAGTTGCCTTAATAGGGCTACTCTCGATTATCTATAAAAAAGCTGAAAAGTTTACCAT   5700
PMP1667      ACTAGTTGCCTTAATAGGGCTACTCTCGATTATCTATAAAAAAGCTGAAAAGTTTACCAT   5700
PMP1663      ACTAGTTGCCTTAATAGGGCTACTCTCGATTATCTATAAAAAAGCTGAAAAGTTTACCAT   5700
PMP1664      ACTAGTTGCCTTAATAGGGCTACTCTCGATTATCTATAAAAAAGCTGAAAAGTTTACCAT   5700
PMP1662      ACTAGTTGCCTTAATAGGGCTACTCTCGATTATCTATAAAAAAGCTGAAAAGTTTACCAT   5700
PMP1611      ACTAGTTGCCTTAATAGGGCTACTCTCGATTATCTATAAAAAAGCTGAAAAGTTTACCAT   5700
             ************************************************************

PMP1610      TTTTCTGTTGGTGCTCTCTATTCTTGTCAGCTCAGTGTCGCTCTTTGCAGTACAGCAGTT   5760
PMP1666      TTTTCTGTTGGTGCTCTCTATTCTTGTCAGCTCAGTGTCGCTCTTTGCAGTACAGCAGTT   5760
PMP1665      TTTTCTGTTGGTGCTCTCTATTCTTGTCAGCTCAGTGTCGCTCTTTGCAGTACAGCAGTT   5760
PMP1667      TTTTCTGTTGGTGCTCTCTATTCTTGTCAGCTCAGTGTCGCTCTTTGCAGTACAGCAGTT   5760
PMP1663      TTTTCTGTTGGTGCTCTCTATTCTTGTCAGCTCAGTGTCGCTCTTTGCAGTACAGCAGTT   5760
PMP1664      TTTTCTGTTGGTGCTCTCTATTCTTGTCAGCTCAGTGTCGCTCTTTGCAGTACAGCAGTT   5760
PMP1662      TTTTCTGTTGGTGCTCTCTATTCTTGTCAGCTCAGTGTCGCTCTTTGCAGTACAGCAGTT   5760
PMP1611      TTTTCTGTTGGTGCTCTCTATTCTTGTCAGCTCAGTGTCGCTCTTTGCAGTACAGCAGTT   5760
             ************************************************************

PMP1610      TGTTGGACTGACCAATCGTTTAAATGCGACTTCTAATTACTCAGAATATTCGCTCAGTGT   5820
PMP1666      TGTTGGACTGACCAATCGTTTAAATGCGACTTCTAATTACTCAGAATATTCGCTCAGTGT   5820
PMP1665      TGTTGGACTGACCAATCGTTTAAATGCGACTTCTAATTACTCAGAATATTCGCTCAGTGT   5820
PMP1667      TGTTGGACTGACCAATCGTTTAAATGCGACTTCTAATTACTCAGAATATTCGCTCAGTGT   5820
PMP1663      TGTTGGACTGACCAATCGTTTAAATGCGACTTCTAATTACTCAGAATATTCGCTCAGTGT   5820
PMP1664      TGTTGGACTGACCAATCGTTTAAATGCGACTTCTAATTACTCAGAATATTCGCTCAGTGT   5820
PMP1662      TGTTGGACTGACCAATCGTTTAAATGCGACTTCTAATTACTCAGAATATTCGCTCAGTGT   5820
PMP1611      TGTTGGACTGACCAATCGTTTAAATGCGACTTCTAATTACTCAGAATATTCGCTCAGTGT   5820
             ************************************************************

PMP1610      CGCTGTTTTAGCAGATAGTGAGATTGAGAATGTTACGCAACTGACGAGTGTGACAGCACC   5880
PMP1666      CGCTGTTTTAGCAGATAGTGAGATTGAGAATGTTACGCAACTGACGAGTGTGACAGCACC   5880
```

```
PMP1665     CGCTGTTTTAGCAGATAGTGAGATTGAGAATGTTACGCAACTGACGAGTGTGACAGCACC    5880
PMP1667     CGCTGTTTTAGCAGATAGTGAGATTGAGAATGTTACGCAACTGACGAGTGTGACAGCACC    5880
PMP1663     CGCTGTTTTAGCAGATAGTGAGATTGAGAATGTTACGCAACTGACGAGTGTGACAGCACC    5880
PMP1664     CGCTGTTTTAGCAGATAGTGAGATTGAGAATGTTACGCAACTGACGAGTGTGACAGCACC    5880
PMP1662     CGCTGTTTTAGCAGATAGTGAGATTGAGAATGTTACGCAACTGACGAGTGTGACAGCACC    5880
PMP1611     CGCTGTTTTAGCAGATAGTGAGATTGAGAATGTTACGCAACTGACGAGTGTGACAGCACC    5880
            ************************************************************

PMP1610     GACTGGGACTGATAATGAAAACATTCAAAAACTACTAGCTGATATCAAATCAAGTCAGAA    5940
PMP1666     GACTGGGACTGATAATGAAAACATTCAAAAACTACTAGCTGATATCAAATCAAGTCAGAA    5940
PMP1665     GACTGGGACTGATAATGAAAACATTCAAAAACTACTAGCTGATATCAAATCAAGTCAGAA    5940
PMP1667     GACTGGGACTGATAATGAAAACATTCAAAAACTACTAGCTGATATCAAATCAAGTCAGAA    5940
PMP1663     GACTGGGACTGATAATGAAAACATTCAAAAACTACTAGCTGATATCAAATCAAGTCAGAA    5940
PMP1664     GACTGGGACTGATAATGAAAACATTCAAAAACTACTAGCTGATATCAAATCAAGTCAGAA    5940
PMP1662     GACTGGGACTGATAATGAAAACATTCAAAAACTACTAGCTGATATCAAATCAAGTCAGAA    5940
PMP1611     GACTGGGACTGATAATGAAAACATTCAAAAACTACTAGCTGATATCAAATCAAGTCAGAA    5940
            ************************************************************

PMP1610     TATCGATTTGACGGTTAATCAAAGTTCGTCTTACTTGTCAGCTTACAGGAGTTTGATTGC    6000
PMP1666     TATCGATTTGACGGTTAATCAAAGTTCGTCTTACTTGTCAGCTTACAGGAGTTTGATTGC    6000
PMP1665     TATCGATTTGACGGTTAATCAAAGTTCGTCTTACTTGTCAGCTTACAGGAGTTTGATTGC    6000
PMP1667     TATCGATTTGACGGTTAATCAAAGTTCGTCTTACTTGTCAGCTTACAGGAGTTTGATTGC    6000
PMP1663     TATCGATTTGACGGTTAATCAAAGTTCGTCTTACTTGTCAGCTTACAGGAGTTTGATTGC    6000
PMP1664     TATCGATTTGACGGTTAATCAAAGTTCGTCTTACTTGTCAGCTTACAGGAGTTTGATTGC    6000
PMP1662     TATCGATTTGACGGTTAATCAAAGTTCGTCTTACTTGTCAGCTTACAGGAGTTTGATTGC    6000
PMP1611     TATCGATTTGACGGTTAATCAAAGTTCGTCTTACTTGTCAGCTTACAGGAGTTTGATTGC    6000
            ************************************************************

PMP1610     AGGAGAGACTAAGGCCATTGTCTTAAATAGTGTCTTTGAAAATATCATTGAATCGGAGTA    6060
PMP1666     AGGAGAGACTAAGGCCATTGTCTTAAATAGTGTCTTTGAAAATATCATTGAATCGGAGTA    6060
PMP1665     AGGAGAGACTAAGGCCATTGTCTTAAATAGTGTCTTTGAAAATATCATTGAATCGGAGTA    6060
PMP1667     AGGAGAGACTAAGGCCATTGTCTTAAATAGTGTCTTTGAAAATATCATTGAATCGGAGTA    6060
PMP1663     AGGAGAGACTAAGGCCATTGTCTTAAATAGTGTCTTTGAAAATATCATTGAATCGGAGTA    6060
PMP1664     AGGAGAGACTAAGGCCATTGTCTTAAATAGTGTCTTTGAAAATATCATTGAATCGGAGTA    6060
PMP1662     AGGAGAGACTAAGGCCATTGTCTTAAATAGTGTCTTTGAAAATATCATTGAATCGGAGTA    6060
PMP1611     AGGAGAGACTAAGGCCATTGTCTTAAATAGTGTCTTTGAAAATATCATTGAATCGGAGTA    6060
```

```
              ************************************************************

PMP1610       TCCAGATCACGCATCGAAGATAAAAAAGATTTATACCAAGGGATTCATTAAAAAAGTAGA   6120
PMP1666       TCCAGATCACGCATCGAAGATAAAAAAGATTTATACCAAGGGATTCATTAAAAAAGTAGA   6120
PMP1665       TCCAGATCACGCATCGAAGATAAAAAAGATTTATACCAAGGGATTCATTAAAAAAGTAGA   6120
PMP1667       TCCAGATCACGCATCGAAGATAAAAAAGATTTATACCAAGGGATTCATTAAAAAAGTAGA   6120
PMP1663       TCCAGATCACGCATCGAAGATAAAAAAGATTTATACCAAGGGATTCATTAAAAAAGTAGA   6120
PMP1664       TCCAGATCACGCATCGAAGATAAAAAAGATTTATACCAAGGGATTCATTAAAAAAGTAGA   6120
PMP1662       TCCAGATCACGCATCGAAGATAAAAAAGATTTATACCAAGGGATTCATTAAAAAAGTAGA   6120
PMP1611       TCCAGATCACGCATCGAAGATAAAAAAGATTTATACCAAGGGATTCATTAAAAAAGTAGA   6120
              ************************************************************

PMP1610       AGCTCCTAAGACGTCTAAGAATCAGTCTTTCAATATCTATGTTAGTGGAATTGATACTTA   6180
PMP1666       AGCTCCTAAGACGTCTAAGAATCAGTCTTTCAATATCTATGTTAGTGGAATTGATACTTA   6180
PMP1665       AGCTCCTAAGACGTCTAAGAATCAGTCTTTCAATATCTATGTTAGTGGAATTGATACTTA   6180
PMP1667       AGCTCCTAAGACGTCTAAGAATCAGTCTTTCAATATCTATGTTAGTGGAATTGATACTTA   6180
PMP1663       AGCTCCTAAGACGTCTAAGAATCAGTCTTTCAATATCTATGTTAGTGGAATTGATACTTA   6180
PMP1664       AGCTCCTAAGACGTCTAAGAATCAGTCTTTCAATATCTATGTTAGTGGAATTGATACTTA   6180
PMP1662       AGCTCCTAAGACGTCTAAGAATCAGTCTTTCAATATCTATGTTAGTGGAATTGATACTTA   6180
PMP1611       AGCTCCTAAGACGTCTAAGAATCAGTCTTTCAATATCTATGTTAGTGGAATTGATACTTA   6180
              ************************************************************

PMP1610       TGGTCCAATTAGTTCGGTGTCGCGTTCAGATGTCAATATCTTGATGACTGTCAATCGAGA   6240
PMP1666       TGGTCCAATTAGTTCGGTGTCGCGTTCAGATGTCAATATCTTGATGACTGTCAATCGAGA   6240
PMP1665       TGGTCCAATTAGTTCGGTGTCGCGTTCAGATGTCAATATCTTGATGACTGTCAATCGAGA   6240
PMP1667       TGGTCCAATTAGTTCGGTGTCGCGTTCAGATGTCAATATCTTGATGACTGTCAATCGAGA   6240
PMP1663       TGGTCCAATTAGTTCGGTGTCGCGTTCAGATGTCAATATCTTGATGACTGTCAATCGAGA   6240
PMP1664       TGGTCCAATTAGTTCGGTGTCGCGTTCAGATGTCAATATCTTGATGACTGTCAATCGAGA   6240
PMP1662       TGGTCCAATTAGTTCGGTGTCGCGTTCAGATGTCAATATCTTGATGACTGTCAATCGAGA   6240
PMP1611       TGGTCCAATTAGTTCGGTGTCGCGTTCAGATGTCAATATCTTGATGACTGTCAATCGAGA   6240
              ************************************************************

PMP1610       TACCAAGAAAATTCTCTTGACCACAACGCCACGTGATGCTTATGTACCAATCGCAGATGG   6300
PMP1666       TACCAAGAAAATTCTCTTGACCACAACGCCACGTGATGCTTATGTACCAATCGCAGATGG   6300
PMP1665       TACCAAGAAAATTCTCTTGACCACAACGCCACGTGATGCTTATGTACCAATCGCAGATGG   6300
PMP1667       TACCAAGAAAATTCTCTTGACCACAACGCCACGTGATGCTTATGTACCAATCGCAGATGG   6300
```

```
PMP1663       TACCAAGAAAATTCTCTTGACCACAACGCCACGTGATGCTTATGTACCAATCGCAGATGG   6300
PMP1664       TACCAAGAAAATTCTCTTGACCACAACGCCACGTGATGCTTATGTACCAATCGCAGATGG   6300
PMP1662       TACCAAGAAAATTCTCTTGACCACAACGCCACGTGATGCTTATGTACCAATCGCAGATGG   6300
PMP1611       TACCAAGAAAATTCTCTTGACCACAACGCCACGTGATGCTTATGTACCAATCGCAGATGG   6300
              ************************************************************

PMP1610       TGGAAATAATCAAAAAGATAAATTGACTCATGCGGGCATTTATGGAGTTGATTCGTCCAT   6360
PMP1666       TGGAAATAATCAAAAAGATAAATTGACTCATGCGGGCATTTATGGAGTTGATTCGTCCAT   6360
PMP1665       TGGAAATAATCAAAAAGATAAATTGACTCATGCGGGCATTTATGGAGTTGATTCGTCCAT   6360
PMP1667       TGGAAATAATCAAAAAGATAAATTGACTCATGCGGGCATTTATGGAGTTGATTCGTCCAT   6360
PMP1663       TGGAAATAATCAAAAAGATAAATTGACTCATGCGGGCATTTATGGAGTTGATTCGTCCAT   6360
PMP1664       TGGAAATAATCAAAAAGATAAATTGACTCATGCGGGCATTTATGGAGTTGATTCGTCCAT   6360
PMP1662       TGGAAATAATCAAAAAGATAAATTGACTCATGCGGGCATTTATGGAGTTGATTCGTCCAT   6360
PMP1611       TGGAAATAATCAAAAAGATAAATTGACTCATGCGGGCATTTATGGAGTTGATTCGTCCAT   6360
              ************************************************************

PMP1610       TCACACCTTAGAAAATCTCTATGGAGTGGATATCAATTACTATGTGCGATTGAACTTCAC   6420
PMP1666       TCACACCTTAGAAAATCTCTATGGAGTGGATATCAATTACTATGTGCGATTGAACTTCAC   6420
PMP1665       TCACACCTTAGAAAATCTCTATGGAGTGGATATCAATTACTATGTGCGATTGAACTTCAC   6420
PMP1667       TCACACCTTAGAAAATCTCTATGGAGTGGATATCAATTACTATGTGCGATTGAACTTCAC   6420
PMP1663       TCACACCTTAGAAAATCTCTATGGAGTGGATATCAATTACTATGTGCGATTGAACTTCAC   6420
PMP1664       TCACACCTTAGAAAATCTCTATGGAGTGGATATCAATTACTATGTGCGATTGAACTTCAC   6420
PMP1662       TCACACCTTAGAAAATCTCTATGGAGTGGATATCAATTACTATGTGCGATTGAACTTCAC   6420
PMP1611       TCACACCTTAGAAAATCTCTATGGAGTGGATATCAATTACTATGTGCGATTGAACTTCAC   6420
              ************************************************************

PMP1610       TTCGTTTTTGAAATTGATTGATTTGTTGGGTGGGGTAGATGTTTATAATGATCAGGAATT   6480
PMP1666       TTCGTTTTTGAAATTGATTGATTTGTTGGGTGGGGTAGATGTTTATAATGATCAGGAATT   6480
PMP1665       TTCGTTTTTGAAATTGATTGATTTGTTGGGTGGGGTAGATGTTTATAATGATCAGGAATT   6480
PMP1667       TTCGTTTTTGAAATTGATTGATTTGTTGGGTGGGGTAGATGTTTATAATGATCAGGAATT   6480
PMP1663       TTCGTTTTTGAAATTGATTGATTTGTTGGGTGGGGTAGATGTTTATAATGATCAGGAATT   6480
PMP1664       TTCGTTTTTGAAATTGATTGATTTGTTGGGTGGGGTAGATGTTTATAATGATCAGGAATT   6480
PMP1662       TTCGTTTTTGAAATTGATTGATTTGTTGGGTGGGGTAGATGTTTATAATGATCAGGAATT   6480
PMP1611       TTCGTTTTTGAAATTGATTGATTTGTTGGGTGGGGTAGATGTTTATAATGATCAGGAATT   6480
              ************************************************************
```

```
PMP1610      CACAGCTCTTGCTAATAAAAAACACTATTCTATTGGTAATGTCCATTTAGATTCAGAAGA    6540
PMP1666      CACAGCTCTTGCTAATAAAAAACACTATTCTATTGGTAATGTCCATTTAGATTCAGAAGA    6540
PMP1665      CACAGCTCTTGCTAATAAAAAACACTATTCTATTGGTAATGTCCATTTAGATTCAGAAGA    6540
PMP1667      CACAGCTCTTGCTAATAAAAAACACTATTCTATTGGTAATGTCCATTTAGATTCAGAAGA    6540
PMP1663      CACAGCTCTTGCTAATAAAAAACACTATTCTATTGGTAATGTCCATTTAGATTCAGAAGA    6540
PMP1664      CACAGCTCTTGCTAATAAAAAACACTATTCTATTGGTAATGTCCATTTAGATTCAGAAGA    6540
PMP1662      CACAGCTCTTGCTAATAAAAAACACTATTCTATTGGTAATGTCCATTTAGATTCAGAAGA    6540
PMP1611      CACAGCTCTTGCTAATAAAAAACACTATTCTATTGGTAATGTCCATTTAGATTCAGAAGA    6540
             ************************************************************

PMP1610      GGCACTCGCTTTTGTTCGTGAGCGCTATTCCCTAGCGGATGGTGATCGTGACCGTGGGCG    6600
PMP1666      GGCACTCGCTTTTGTTCGTGAGCGCTATTCCCTAGCGGATGGTGATCGTGACCGTGGGCG    6600
PMP1665      GGCACTCGCTTTTGTTCGTGAGCGCTATTCCCTAGCGGATGGTGATCGTGACCGTGGGCG    6600
PMP1667      GGCACTCGCTTTTGTTCGTGAGCGCTATTCCCTAGCGGATGGTGATCGTGACCGTGGGCG    6600
PMP1663      GGCACTCGCTTTTGTTCGTGAGCGCTATTCCCTAGCGGATGGTGATCGTGACCGTGGGCG    6600
PMP1664      GGCACTCGCTTTTGTTCGTGAGCGCTATTCCCTAGCGGATGGTGATCGTGACCGTGGGCG    6600
PMP1662      GGCACTCGCTTTTGTTCGTGAGCGCTATTCCCTAGCGGATGGTGATCGTGACCGTGGGCG    6600
PMP1611      GGCACTCGCTTTTGTTCGTGAGCGCTATTCCCTAGCGGATGGTGATCGTGACCGTGGGCG    6600
             ************************************************************

PMP1610      CAATCAACAAAAGGTGATTGTGGCTATCCTTCAAAAATTAACTTCGACCGAAGTACTGAA    6660
PMP1666      CAATCAACAAAAGGTGATTGTGGCTATCCTTCAAAAATTAACTTCGACCGAAGTACTGAA    6660
PMP1665      CAATCAACAAAAGGTGATTGTGGCTATCCTTCAAAAATTAACTTCGACCGAAGTACTGAA    6660
PMP1667      CAATCAACAAAAGGTGATTGTGGCTATCCTTCAAAAATTAACTTCGACCGAAGTACTGAA    6660
PMP1663      CAATCAACAAAAGGTGATTGTGGCTATCCTTCAAAAATTAACTTCGACCGAAGTACTGAA    6660
PMP1664      CAATCAACAAAAGGTGATTGTGGCTATCCTTCAAAAATTAACTTCGACCGAAGTACTGAA    6660
PMP1662      CAATCAACAAAAGGTGATTGTGGCTATCCTTCAAAAATTAACTTCGACCGAAGTACTGAA    6660
PMP1611      CAATCAACAAAAGGTGATTGTGGCTATCCTTCAAAAATTAACTTCGACCGAAGTACTGAA    6660
             ************************************************************

PMP1610      AAATTATAGTACGATCATTGATAGCTTGCAAGATTCTATCCAAACAAACATGCCACTTGA    6720
PMP1666      AAATTATAGTACGATCATTGATAGCTTGCAAGATTCTATCCAAACAAACATGCCACTTGA    6720
PMP1665      AAATTATAGTACGATCATTGATAGCTTGCAAGATTCTATCCAAACAAACATGCCACTTGA    6720
PMP1667      AAATTATAGTACGATCATTGATAGCTTGCAAGATTCTATCCAAACAAACATGCCACTTGA    6720
PMP1663      AAATTATAGTACGATCATTGATAGCTTGCAAGATTCTATCCAAACAAACATGCCACTTGA    6720
PMP1664      AAATTATAGTACGATCATTGATAGCTTGCAAGATTCTATCCAAACAAACATGCCACTTGA    6720
```

```
PMP1662      AAATTATAGTACGATCATTGATAGCTTGCAAGATTCTATCCAAACAAACATGCCACTTGA    6720
PMP1611      AAATTATAGTACGATCATTGATAGCTTGCAAGATTCTATCCAAACAAACATGCCACTTGA    6720
             ************************************************************

PMP1610      GACCATGATAAACTTGGTCAATGCTCAGTTAGAAAGTGGTGGAACGTACAAAGTAAATTC    6780
PMP1666      GACCATGATAAACTTGGTCAATGCTCAGTTAGAAAGTGGTGGAACGTACAAAGTAAATTC    6780
PMP1665      GACCATGATAAACTTGGTCAATGCTCAGTTAGAAAGTGGTGGAACGTACAAAGTAAATTC    6780
PMP1667      GACCATGATAAACTTGGTCAATGCTCAGTTAGAAAGTGGTGGAACGTACAAAGTAAATTC    6780
PMP1663      GACCATGATAAACTTGGTCAATGCTCAGTTAGAAAGTGGTGGAACGTACAAAGTAAATTC    6780
PMP1664      GACCATGATAAACTTGGTCAATGCTCAGTTAGAAAGTGGTGGAACGTACAAAGTAAATTC    6780
PMP1662      GACCATGATAAACTTGGTCAATGCTCAGTTAGAAAGTGGTGGAACGTACAAAGTAAATTC    6780
PMP1611      GACCATGATAAACTTGGTCAATGCTCAGTTAGAAAGTGGTGGAACGTACAAAGTAAATTC    6780
             ************************************************************

PMP1610      GCAAGACTTGAAGGGTAGGGGACGGACGGATCTTCCTTCCTATGCGATGCCAGATAGTAA    6840
PMP1666      GCAAGACTTGAAGGGTAGGGGACGGACGGATCTTCCTTCCTATGCGATGCCAGATAGTAA    6840
PMP1665      GCAAGACTTGAAGGGTAGGGGACGGACGGATCTTCCTTCCTATGCGATGCCAGATAGTAA    6840
PMP1667      GCAAGACTTGAAGGGTAGGGGACGGACGGATCTTCCTTCCTATGCGATGCCAGATAGTAA    6840
PMP1663      GCAAGACTTGAAGGGTAGGGGACGGACGGATCTTCCTTCCTATGCGATGCCAGATAGTAA    6840
PMP1664      GCAAGACTTGAAGGGTAGGGGACGGACGGATCTTCCTTCCTATGCGATGCCAGATAGTAA    6840
PMP1662      GCAAGACTTGAAGGGTAGGGGACGGACGGATCTTCCTTCCTATGCGATGCCAGATAGTAA    6840
PMP1611      GCAAGACTTGAAGGGTAGGGGACGGACGGATCTTCCTTCCTATGCGATGCCAGATAGTAA    6840
             ************************************************************

PMP1610      CCTCTATGTGATGGAAATTAACGACAGTAGCCTTGCATCTGTCAAAACGGCTATTCAGGA    6900
PMP1666      CCTCTATGTGATGGAAATTAACGACAGTAGCCTTGCATCTGTCAAAACGGCTATTCAGGA    6900
PMP1665      CCTCTATGTGATGGAAATTAACGACAGTAGCCTTGCATCTGTCAAAACGGCTATTCAGGA    6900
PMP1667      CCTCTATGTGATGGAAATTAACGACAGTAGCCTTGCATCTGTCAAAACGGCTATTCAGGA    6900
PMP1663      CCTCTATGTGATGGAAATTAACGACAGTAGCCTTGCATCTGTCAAAACGGCTATTCAGGA    6900
PMP1664      CCTCTATGTGATGGAAATTAACGACAGTAGCCTTGCATCTGTCAAAACGGCTATTCAGGA    6900
PMP1662      CCTCTATGTGATGGAAATTAACGACAGTAGCCTTGCATCTGTCAAAACGGCTATTCAGGA    6900
PMP1611      CCTCTATGTGATGGAAATTAACGACAGTAGCCTTGCATCTGTCAAAACGGCTATTCAGGA    6900
             ************************************************************

PMP1610      CGTGTTGGAGGGCAGATGAAATGATTGATATTCATTCGCACATTGTCTTTGATGTAGATG    6960
PMP1666      CGTGTTGGAGGGCAGATGAAATGATTGATATTCATTCGCACATTGTCTTTGATGTAGATG    6960
```

```
PMP1665      CGTGTTGGAGGGCAGATGAAATGATTGATATTCATTCGCACATTGTCTTTGATGTAGATG   6960
PMP1667      CGTGTTGGAGGGCAGATGAAATGATTGATATTCATTCGCACATTGTCTTTGATGTAGATG   6960
PMP1663      CGTGTTGGAGGGCAGATGAAATGATTGATATTCATTCGCACATTGTCTTTGATGTAGATG   6960
PMP1664      CGTGTTGGAGGGCAGATGAAATGATTGATATTCATTCGCACATTGTCTTTGATGTAGATG   6960
PMP1662      CGTGTTGGAGGGCAGATGAAATGATTGATATTCATTCGCACATTGTCTTTGATGTAGATG   6960
PMP1611      CGTGTTGGAGGGCAGATGAAATGATTGATATTCATTCGCACATTGTCTTTGATGTAGATG   6960
             ************************************************************

PMP1610      ATGGTCCCAAGTCAAGAGAGGAAAGTAAGGCTCTCTTGACAGAAGCCTACAGGCAGGGGG   7020
PMP1666      ATGGTCCCAAGTCAAGAGAGGAAAGTAAGGCTCTCTTGACAGAAGCCTACAGGCAGGGGG   7020
PMP1665      ATGGTCCCAAGTCAAGAGAGGAAAGTAAGGCTCTCTTGACAGAAGCCTACAGGCAGGGGG   7020
PMP1667      ATGGTCCCAAGTCAAGAGAGGAAAGTAAGGCTCTCTTGACAGAAGCCTACAGGCAGGGGG   7020
PMP1663      ATGGTCCCAAGTCAAGAGAGGAAAGTAAGGCTCTCTTGACAGAAGCCTACAGGCAGGGGG   7020
PMP1664      ATGGTCCCAAGTCAAGAGAGGAAAGTAAGGCTCTCTTGACAGAAGCCTACAGGCAGGGGG   7020
PMP1662      ATGGTCCCAAGTCAAGAGAGGAAAGTAAGGCTCTCTTGACAGAAGCCTACAGGCAGGGGG   7020
PMP1611      ATGGTCCCAAGTCAAGAGAGGAAAGTAAGGCTCTCTTGACAGAAGCCTACAGGCAGGGGG   7020
             ************************************************************

PMP1610      TGCGAACCATTGTCTCTACCTCTCACCGTCGCAAGGGCATGTTTGAAACTCCAGAAGAGA   7080
PMP1666      TGCGAACCATTGTCTCTACCTCTCACCGTCGCAAGGGCATGTTTGAAACTCCAGAAGAGA   7080
PMP1665      TGCGAACCATTGTCTCTACCTCTCACCGTCGCAAGGGCATGTTTGAAACTCCAGAAGAGA   7080
PMP1667      TGCGAACCATTGTCTCTACCTCTCACCGTCGCAAGGGCATGTTTGAAACTCCAGAAGAGA   7080
PMP1663      TGCGAACCATTGTCTCTACCTCTCACCGTCGCAAGGGCATGTTTGAAACTCCAGAAGAGA   7080
PMP1664      TGCGAACCATTGTCTCTACCTCTCACCGTCGCAAGGGCATGTTTGAAACTCCAGAAGAGA   7080
PMP1662      TGCGAACCATTGTCTCTACCTCTCACCGTCGCAAGGGCATGTTTGAAACTCCAGAAGAGA   7080
PMP1611      TGCGAACCATTGTCTCTACCTCTCACCGTCGCAAGGGCATGTTTGAAACTCCAGAAGAGA   7080
             ************************************************************

PMP1610      AGATAGCAGAAAACTTTCTTCAGGTTCGGGAAATAGCTAAGGAAGTCGCGAGTGACTTGG   7140
PMP1666      AGATAGCAGAAAACTTTCTTCAGGTTCGGGAAATAGCTAAGGAAGTCGCGAGTGACTTGG   7140
PMP1665      AGATAGCAGAAAACTTTCTTCAGGTTCGGGAAATAGCTAAGGAAGTCGCGAGTGACTTGG   7140
PMP1667      AGATAGCAGAAAACTTTCTTCAGGTTCGGGAAATAGCTAAGGAAGTCGCGAGTGACTTGG   7140
PMP1663      AGATAGCAGAAAACTTTCTTCAGGTTCGGGAAATAGCTAAGGAAGTCGCGAGTGACTTGG   7140
PMP1664      AGATAGCAGAAAACTTTCTTCAGGTTCGGGAAATAGCTAAGGAAGTCGCGAGTGACTTGG   7140
PMP1662      AGATAGCAGAAAACTTTCTTCAGGTTCGGGAAATAGCTAAGGAAGTCGCGAGTGACTTGG   7140
PMP1611      AGATAGCAGAAAACTTTCTTCAGGTTCGGGAAATAGCTAAGGAAGTCGCGAGTGACTTGG   7140
```

```
                ************************************************************

PMP1610         TCATTGCTTATGGGGCTGAAATTTACTACACGCCAGATGTTTTGGATAAGCTGGAAAACA    7200
PMP1666         TCATTGCTTATGGGGCTGAAATTTACTACACGCCAGATGTTTTGGATAAGCTGGAAAACA    7200
PMP1665         TCATTGCTTATGGGGCTGAAATTTACTACACGCCAGATGTTTTGGATAAGCTGGAAAACA    7200
PMP1667         TCATTGCTTATGGGGCTGAAATTTACTACACGCCAGATGTTTTGGATAAGCTGGAAAACA    7200
PMP1663         TCATTGCTTATGGGGCTGAAATTTACTACACGCCAGATGTTTTGGATAAGCTGGAAAACA    7200
PMP1664         TCATTGCTTATGGGGCTGAAATTTACTACACGCCAGATGTTTTGGATAAGCTGGAAAACA    7200
PMP1662         TCATTGCTTATGGGGCTGAAATTTACTACACGCCAGATGTTTTGGATAAGCTGGAAAACA    7200
PMP1611         TCATTGCTTATGGGGCTGAAATTTACTACACGCCAGATGTTTTGGATAAGCTGGAAAACA    7200
                ************************************************************

PMP1610         ATCGGATTCCGACCCTCAATAATAGTCGTTATGCCTTGATAGAGTTTAGTATGAACACTC    7260
PMP1666         ATCGGATTCCGACCCTCAATAATAGTCGTTATGCCTTGATAGAGTTTAGTATGAACACTC    7260
PMP1665         ATCGGATTCCGACCCTCAATAATAGTCGTTATGCCTTGATAGAGTTTAGTATGAACACTC    7260
PMP1667         ATCGGATTCCGACCCTCAATAATAGTCGTTATGCCTTGATAGAGTTTAGTATGAACACTC    7260
PMP1663         ATCGGATTCCGACCCTCAATAATAGTCGTTATGCCTTGATAGAGTTTAGTATGAACACTC    7260
PMP1664         ATCGGATTCCGACCCTCAATAATAGTCGTTATGCCTTGATAGAGTTTAGTATGAACACTC    7260
PMP1662         ATCGGATTCCGACCCTCAATAATAGTCGTTATGCCTTGATAGAGTTTAGTATGAACACTC    7260
PMP1611         ATCGGATTCCGACCCTCAATAATAGTCGTTATGCCTTGATAGAGTTTAGTATGAACACTC    7260
                ************************************************************

PMP1610         CTTATCGCGATATTCATAGTGCCTTGAATAAAATATTGATGTTGGGAATTACTCCCGTCA    7320
PMP1666         CTTATCGCGATATTCATAGTGCCTTGAATAAAATATTGATGTTGGGAATTACTCCCGTCA    7320
PMP1665         CTTATCGCGATATTCATAGTGCCTTGAATAAAATATTGATGTTGGGAATTACTCCCGTCA    7320
PMP1667         CTTATCGCGATATTCATAGTGCCTTGAATAAAATATTGATGTTGGGAATTACTCCCGTCA    7320
PMP1663         CTTATCGCGATATTCATAGTGCCTTGAATAAAATATTGATGTTGGGAATTACTCCCGTCA    7320
PMP1664         CTTATCGCGATATTCATAGTGCCTTGAATAAAATATTGATGTTGGGAATTACTCCCGTCA    7320
PMP1662         CTTATCGCGATATTCATAGTGCCTTGAATAAAATATTGATGTTGGGAATTACTCCCGTCA    7320
PMP1611         CTTATCGCGATATTCATAGTGCCTTGAATAAAATATTGATGTTGGGAATTACTCCCGTCA    7320
                ************************************************************

PMP1610         TTGCCCACATAGAGCGCTATGATGTTCTTGAAAATAATGAAAAACGCGTTCGAGAGCTGA    7380
PMP1666         TTGCCCACATAGAGCGCTATGATGTTCTTGAAAATAATGAAAAACGCGTTCGAGAGCTGA    7380
PMP1665         TTGCCCACATAGAGCGCTATGATGTTCTTGAAAATAATGAAAAACGCGTTCGAGAGCTGA    7380
PMP1667         TTGCCCACATAGAGCGCTATGATGTTCTTGAAAATAATGAAAAACGCGTTCGAGAGCTGA    7380
```

```
PMP1663       TTGCCCACATAGAGCGCTATGATGTTCTTGAAAATAATGAAAAACGCGTTCGAGAGCTGA    7380
PMP1664       TTGCCCACATAGAGCGCTATGATGTTCTTGAAAATAATGAAAAACGCGTTCGAGAGCTGA    7380
PMP1662       TTGCCCACATAGAGCGCTATGATGTTCTTGAAAATAATGAAAAACGCGTTCGAGAGCTGA    7380
PMP1611       TTGCCCACATAGAGCGCTATGATGTTCTTGAAAATAATGAAAAACGCGTTCGAGAGCTGA    7380
              ************************************************************

PMP1610       TCGATATGGGCTGTTACACGCAAATAAATAGTTCACATGTCCTCAAATCCAAACTTTTTG    7440
PMP1666       TCGATATGGGCTGTTACACGCAAATAAATAGTTCACATGTCCTCAAATCCAAACTTTTTG    7440
PMP1665       TCGATATGGGCTGTTACACGCAAATAAATAGTTCACATGTCCTCAAATCCAAACTTTTTG    7440
PMP1667       TCGATATGGGCTGTTACACGCAAATAAATAGTTCACATGTCCTCAAATCCAAACTTTTTG    7440
PMP1663       TCGATATGGGCTGTTACACGCAAATAAATAGTTCACATGTCCTCAAATCCAAACTTTTTG    7440
PMP1664       TCGATATGGGCTGTTACACGCAAATAAATAGTTCACATGTCCTCAAATCCAAACTTTTTG    7440
PMP1662       TCGATATGGGCTGTTACACGCAAATAAATAGTTCACATGTCCTCAAATCCAAACTTTTTG    7440
PMP1611       TCGATATGGGCTGTTACACGCAAATAAATAGTTCACATGTCCTCAAATCCAAACTTTTTG    7440
              ************************************************************

PMP1610       GAGAACCTTATAAATTCATGAAAAAAAGAGCGCAGTATTTCTTGGAGCGTGATTTGGTTC    7500
PMP1666       GAGAACCTTATAAATTCATGAAAAAAAGAGCGCAGTATTTCTTGGAGCGTGATTTGGTTC    7500
PMP1665       GAGAACCTTATAAATTCATGAAAAAAAGAGCGCAGTATTTCTTGGAGCGTGATTTGGTTC    7500
PMP1667       GAGAACCTTATAAATTCATGAAAAAAAGAGCGCAGTATTTCTTGGAGCGTGATTTGGTTC    7500
PMP1663       GAGAACCTTATAAATTCATGAAAAAAAGAGCGCAGTATTTCTTGGAGCGTGATTTGGTTC    7500
PMP1664       GAGAACCTTATAAATTCATGAAAAAAAGAGCGCAGTATTTCTTGGAGCGTGATTTGGTTC    7500
PMP1662       GAGAACCTTATAAATTCATGAAAAAAAGAGCGCAGTATTTCTTGGAGCGTGATTTGGTTC    7500
PMP1611       GAGAACCTTATAAATTCATGAAAAAAAGAGCGCAGTATTTCTTGGAGCGTGATTTGGTTC    7500
              ************************************************************

PMP1610       ATATCATTGCAAGTGATATGCATAATGTGGACGGCAGACCCCCCCATATGGCAGAAGCAT    7560
PMP1666       ATATCATTGCAAGTGATATGCATAATGTGGACGGCAGACCCCCCCATATGGCAGAAGCAT    7560
PMP1665       ATATCATTGCAAGTGATATGCATAATGTGGACGGCAGACCCCCCCATATGGCAGAAGCAT    7560
PMP1667       ATATCATTGCAAGTGATATGCATAATGTGGACGGCAGACCCCCCCATATGGCAGAAGCAT    7560
PMP1663       ATATCATTGCAAGTGATATGCATAATGTGGACGGCAGACCCCCCCATATGGCAGAAGCAT    7560
PMP1664       ATATCATTGCAAGTGATATGCATAATGTGGACGGCAGACCCCCCCATATGGCAGAAGCAT    7560
PMP1662       ATATCATTGCAAGTGATATGCATAATGTGGACGGCAGACCCCCCCATATGGCAGAAGCAT    7560
PMP1611       ATATCATTGCAAGTGATATGCATAATGTGGACGGCAGACCCCCCCATATGGCAGAAGCAT    7560
              ************************************************************
```

```
PMP1610     ATGACCTTGTTTCCCAAAAATACGGAGAAGCGAAGGCTCAGGAACTTTTTATAGACAATC   7620
PMP1666     ATGACCTTGTTTCCCAAAAATACGGAGAAGCGAAGGCTCAGGAACTTTTTATAGACAATC   7620
PMP1665     ATGACCTTGTTTCCCAAAAATACGGAGAAGCGAAGGCTCAGGAACTTTTTATAGACAATC   7620
PMP1667     ATGACCTTGTTTCCCAAAAATACGGAGAAGCGAAGGCTCAGGAACTTTTTATAGACAATC   7620
PMP1663     ATGACCTTGTTTCCCAAAAATACGGAGAAGCGAAGGCTCAGGAACTTTTTATAGACAATC   7620
PMP1664     ATGACCTTGTTTCCCAAAAATACGGAGAAGCGAAGGCTCAGGAACTTTTTATAGACAATC   7620
PMP1662     ATGACCTTGTTTCCCAAAAATACGGAGAAGCGAAGGCTCAGGAACTTTTTATAGACAATC   7620
PMP1611     ATGACCTTGTTTCCCAAAAATACGGAGAAGCGAAGGCTCAGGAACTTTTTATAGACAATC   7620
            ************************************************************

PMP1610     CTCGAAAAAATTGTAATGGATCAACTAATTTAGGAGAAATGATGAAAGAACAAAACACGAT   7680
PMP1666     CTCGAAAAAATTGTAATGGATCAACTAATTTAGGAGAAATGATGAAAGAACAAAACACGAT   7680
PMP1665     CTCGAAAAAATTGTAATGGATCAACTAATTTAGGAGAAATGATGAAAGAACAAAACACGAT   7680
PMP1667     CTCGAAAAAATTGTAATGGATCAACTAATTTAGGAGAAATGATGAAAGAACAAAACACGAT   7680
PMP1663     CTCGAAAAAATTGTAATGGATCAACTAATTTAGGAGAAATGATGAAAGAACAAAACACGAT   7680
PMP1664     CTCGAAAAAATTGTAATGGATCAACTAATTTAGGAGAAATGATGAAAGAACAAAACACGAT   7680
PMP1662     CTCGAAAAAATTGTAATGGATCAACTAATTTAGGAGAAATGATGAAAGAACAAAACACGAT   7680
PMP1611     CTCGAAAAAATTGTAATGGATCAACTAATTTAGGAGAAATGATGAAAGAACAAAACACGAT   7680
            ************************************************************

PMP1610     AGAAATCGATGTATTTCAAGTACTTAAAACCTTGTGGAAACACAAGTTAATAATTTTATT   7740
PMP1666     AGAAATCGATGTATTTCAAGTACTTAAAACCTTGTGGAAACACAAGTTAATAATTTTATT   7740
PMP1665     AGAAATCGATGTATTTCAAGTACTTAAAACCTTGTGGAAACACAAGTTAATAATTTTATT   7740
PMP1667     AGAAATCGATGTATTTCAAGTACTTAAAACCTTGTGGAAACACAAGTTAATAATTTTATT   7740
PMP1663     AGAAATCGATGTATTTCAAGTACTTAAAACCTTGTGGAAACACAAGTTAATAATTTTATT   7740
PMP1664     AGAAATCGATGTATTTCAAGTACTTAAAACCTTGTGGAAACACAAGTTAATAATTTTATT   7740
PMP1662     AGAAATCGATGTATTTCAAGTACTTAAAACCTTGTGGAAACACAAGTTAATAATTTTATT   7740
PMP1611     AGAAATCGATGTATTTCAAGTACTTAAAACCTTGTGGAAACACAAGTTAATAATTTTATT   7740
            ************************************************************

PMP1610     AGTGGCGCTTGTGACAGGGGCGGGAGCTTTTGCATATAGCACTTTTATTGTTAAACCAGA   7800
PMP1666     AGTGGCGCTTGTGACAGGGGCGGGAGCTTTTGCATATAGCACTTTTATTGTTAAACCAGA   7800
PMP1665     AGTGGCGCTTGTGACAGGGGCGGGAGCTTTTGCATATAGCACTTTTATTGTTAAACCAGA   7800
PMP1667     AGTGGCGCTTGTGACAGGGGCGGGAGCTTTTGCATATAGCACTTTTATTGTTAAACCAGA   7800
PMP1663     AGTGGCGCTTGTGACAGGGGCGGGAGCTTTTGCATATAGCACTTTTATTGTTAAACCAGA   7800
PMP1664     AGTGGCGCTTGTGACAGGGGCGGGAGCTTTTGCATATAGCACTTTTATTGTTAAACCAGA   7800
```

```
PMP1662       AGTGGCGCTTGTGACAGGGGCGGGAGCTTTTGCATATAGCACTTTTATTGTTAAACCAGA   7800
PMP1611       AGTGGCGCTTGTGACAGGGGCGGGAGCTTTTGCATATAGCACTTTTATTGTTAAACCAGA   7800
              ************************************************************

PMP1610       ATATACGAGCACCACGCGTATTTACGTCGTCAACCGTAATCAAGAAGGTAAGTCGGGACT   7860
PMP1666       ATATACGAGCACCACGCGTATTTACGTCGTCAACCGTAATCAAGAAGGTAAGTCGGGACT   7860
PMP1665       ATATACGAGCACCACGCGTATTTACGTCGTCAACCGTAATCAAGAAGGTAAGTCGGGACT   7860
PMP1667       ATATACGAGCACCACGCGTATTTACGTCGTCAACCGTAATCAAGAAGGTAAGTCGGGACT   7860
PMP1663       ATATACGAGCACCACGCGTATTTACGTCGTCAACCGTAATCAAGAAGGTAAGTCGGGACT   7860
PMP1664       ATATACGAGCACCACGCGTATTTACGTCGTCAACCGTAATCAAGAAGGTAAGTCGGGACT   7860
PMP1662       ATATACGAGCACCACGCGTATTTACGTCGTCAACCGTAATCAAGAAGGTAAGTCGGGACT   7860
PMP1611       ATATACGAGCACCACGCGTATTTACGTCGTCAACCGTAATCAAGAAGGTAAGTCGGGACT   7860
              ************************************************************

PMP1610       GACGAATCAGGACTTGCAGGCAGGAACTTATCTGGTAAAAGACTACCGCGAAATTATCCT   7920
PMP1666       GACGAATCAGGACTTGCAGGCAGGAACTTATCTGGTAAAAGACTACCGCGAAATTATCCT   7920
PMP1665       GACGAATCAGGACTTGCAGGCAGGAACTTATCTGGTAAAAGACTACCGCGAAATTATCCT   7920
PMP1667       GACGAATCAGGACTTGCAGGCAGGAACTTATCTGGTAAAAGACTACCGCGAAATTATCCT   7920
PMP1663       GACGAATCAGGACTTGCAGGCAGGAACTTATCTGGTAAAAGACTACCGCGAAATTATCCT   7920
PMP1664       GACGAATCAGGACTTGCAGGCAGGAACTTATCTGGTAAAAGACTACCGCGAAATTATCCT   7920
PMP1662       GACGAATCAGGACTTGCAGGCAGGAACTTATCTGGTAAAAGACTACCGCGAAATTATCCT   7920
PMP1611       GACGAATCAGGACTTGCAGGCAGGAACTTATCTGGTAAAAGACTACCGCGAAATTATCCT   7920
              ************************************************************

PMP1610       TTCGCAAGATGTATTGGAAAAGGTAGCGACAAATTTGAAATTGGATATGCCAGCAAAAGC   7980
PMP1666       TTCGCAAGATGTATTGGAAAAGGTAGCGACAAATTTGAAATTGGATATGCCAGCAAAAGC   7980
PMP1665       TTCGCAAGATGTATTGGAAAAGGTAGCGACAAATTTGAAATTGGATATGCCAGCAAAAGC   7980
PMP1667       TTCGCAAGATGTATTGGAAAAGGTAGCGACAAATTTGAAATTGGATATGCCAGCAAAAGC   7980
PMP1663       TTCGCAAGATGTATTGGAAAAGGTAGCGACAAATTTGAAATTGGATATGCCAGCAAAAGC   7980
PMP1664       TTCGCAAGATGTATTGGAAAAGGTAGCGACAAATTTGAAATTGGATATGCCAGCAAAAGC   7980
PMP1662       TTCGCAAGATGTATTGGAAAAGGTAGCGACAAATTTGAAATTGGATATGCCAGCAAAAGC   7980
PMP1611       TTCGCAAGATGTATTGGAAAAGGTAGCGACAAATTTGAAATTGGATATGCCAGCAAAAGC   7980
              ************************************************************

PMP1610       GTTAACTAGCAAAGTTCAAGTGACTGTACCAACCGACACTCGTATCGTCTCAATCTCTGT   8040
PMP1666       GTTAACTAGCAAAGTTCAAGTGACTGTACCAACCGACACTCGTATCGTCTCAATCTCTGT   8040
```

```
PMP1665      GTTAACTAGCAAAGTTCAAGTGACTGTACCAACCGACACTCGTATCGTCTCAATCTCTGT   8040
PMP1667      GTTAACTAGCAAAGTTCAAGTGACTGTACCAACCGACACTCGTATCGTCTCAATCTCTGT   8040
PMP1663      GTTAACTAGCAAAGTTCAAGTGACTGTACCAACCGACACTCGTATCGTCTCAATCTCTGT   8040
PMP1664      GTTAACTAGCAAAGTTCAAGTGACTGTACCAACCGACACTCGTATCGTCTCAATCTCTGT   8040
PMP1662      GTTAACTAGCAAAGTTCAAGTGACTGTACCAACCGACACTCGTATCGTCTCAATCTCTGT   8040
PMP1611      GTTAACTAGCAAAGTTCAAGTGACTGTACCAACCGACACTCGTATCGTCTCAATCTCTGT   8040
             ************************************************************

PMP1610      CAAGGATAAAGAACCAGAGGAAGCCAGTCGCATTGCTAATTCTCTACGAGAAGTTGCTGC   8100
PMP1666      CAAGGATAAAGAACCAGAGGAAGCCAGTCGCATTGCTAATTCTCTACGAGAAGTTGCTGC   8100
PMP1665      CAAGGATAAAGAACCAGAGGAAGCCAGTCGCATTGCTAATTCTCTACGAGAAGTTGCTGC   8100
PMP1667      CAAGGATAAAGAACCAGAGGAAGCCAGTCGCATTGCTAATTCTCTACGAGAAGTTGCTGC   8100
PMP1663      CAAGGATAAAGAACCAGAGGAAGCCAGTCGCATTGCTAATTCTCTACGAGAAGTTGCTGC   8100
PMP1664      CAAGGATAAAGAACCAGAGGAAGCCAGTCGCATTGCTAATTCTCTACGAGAAGTTGCTGC   8100
PMP1662      CAAGGATAAAGAACCAGAGGAAGCCAGTCGCATTGCTAATTCTCTACGAGAAGTTGCTGC   8100
PMP1611      CAAGGATAAAGAACCAGAGGAAGCCAGTCGCATTGCTAATTCTCTACGAGAAGTTGCTGC   8100
             ************************************************************

PMP1610      AGGAAAGATCGTCGCTGTAACGCGAGTATCTGATGTAACGACGCTTGAAGAAGCGCGGCC   8160
PMP1666      AGGAAAGATCGTCGCTGTAACGCGAGTATCTGATGTAACGACGCTTGAAGAAGCGCGGCC   8160
PMP1665      AGGAAAGATCGTCGCTGTAACGCGAGTATCTGATGTAACGACGCTTGAAGAAGCGCGGCC   8160
PMP1667      AGGAAAGATCGTCGCTGTAACGCGAGTATCTGATGTAACGACGCTTGAAGAAGCGCGGCC   8160
PMP1663      AGGAAAGATCGTCGCTGTAACGCGAGTATCTGATGTAACGACGCTTGAAGAAGCGCGGCC   8160
PMP1664      AGGAAAGATCGTCGCTGTAACGCGAGTATCTGATGTAACGACGCTTGAAGAAGCGCGGCC   8160
PMP1662      AGGAAAGATCGTCGCTGTAACGCGAGTATCTGATGTAACGACGCTTGAAGAAGCGCGGCC   8160
PMP1611      AGGAAAGATCGTCGCTGTAACGCGAGTATCTGATGTAACGACGCTTGAAGAAGCGCGGCC   8160
             ************************************************************

PMP1610      AGCTACGACTCCCTCTTCTCCCAATGTTCGACGTAACACCTTAGTTGGTTTTCTTGGAGG   8220
PMP1666      AGCTACGACTCCCTCTTCTCCCAATGTTCGACGTAACACCTTAGTTGGTTTTCTTGGAGG   8220
PMP1665      AGCTACGACTCCCTCTTCTCCCAATGTTCGACGTAACACCTTAGTTGGTTTTCTTGGAGG   8220
PMP1667      AGCTACGACTCCCTCTTCTCCCAATGTTCGACGTAACACCTTAGTTGGTTTTCTTGGAGG   8220
PMP1663      AGCTACGACTCCCTCTTCTCCCAATGTTCGACGTAACACCTTAGTTGGTTTTCTTGGAGG   8220
PMP1664      AGCTACGACTCCCTCTTCTCCCAATGTTCGACGTAACACCTTAGTTGGTTTTCTTGGAGG   8220
PMP1662      AGCTACGACTCCCTCTTCTCCCAATGTTCGACGTAACACCTTAGTTGGTTTTCTTGGAGG   8220
PMP1611      AGCTACGACTCCCTCTTCTCCCAATGTTCGACGTAACACCTTAGTTGGTTTTCTTGGAGG   8220
```

```
            ************************************************************

PMP1610     AGCCGTCGTAACAGTAATTACTGTTCTTTTGATTGAGTTGCTCGATACCCGTGTGAAACG   8280
PMP1666     AGCCGTCGTAACAGTAATTACTGTTCTTTTGATTGAGTTGCTCGATACCCGTGTGAAACG   8280
PMP1665     AGCCGTCGTAACAGTAATTACTGTTCTTTTGATTGAGTTGCTCGATACCCGTGTGAAACG   8280
PMP1667     AGCCGTCGTAACAGTAATTACTGTTCTTTTGATTGAGTTGCTCGATACCCGTGTGAAACG   8280
PMP1663     AGCCGTCGTAACAGTAATTACTGTTCTTTTGATTGAGTTGCTCGATACCCGTGTGAAACG   8280
PMP1664     AGCCGTCGTAACAGTAATTACTGTTCTTTTGATTGAGTTGCTCGATACCCGTGTGAAACG   8280
PMP1662     AGCCGTCGTAACAGTAATTACTGTTCTTTTGATTGAGTTGCTCGATACCCGTGTGAAACG   8280
PMP1611     AGCCGTCGTAACAGTAATTACTGTTCTTTTGATTGAGTTGCTCGATACCCGTGTGAAACG   8280
            ************************************************************

PMP1610     TCCTGAAGAAGTTGAAGAGGTGCTGCAAGTGCCACTTCTAGGAGTCGTTCCAGATTTGGA   8340
PMP1666     TCCTGAAGAAGTTGAAGAGGTGCTGCAAGTGCCACTTCTAGGAGTCGTTCCAGATTTGGA   8340
PMP1665     TCCTGAAGAAGTTGAAGAGGTGCTGCAAGTGCCACTTCTAGGAGTCGTTCCAGATTTGGA   8340
PMP1667     TCCTGAAGAAGTTGAAGAGGTGCTGCAAGTGCCACTTCTAGGAGTCGTTCCAGATTTGGA   8340
PMP1663     TCCTGAAGAAGTTGAAGAGGTGCTGCAAGTGCCACTTCTAGGAGTCGTTCCAGATTTGGA   8340
PMP1664     TCCTGAAGAAGTTGAAGAGGTGCTGCAAGTGCCACTTCTAGGAGTCGTTCCAGATTTGGA   8340
PMP1662     TCCTGAAGAAGTTGAAGAGGTGCTGCAAGTGCCACTTCTAGGAGTCGTTCCAGATTTGGA   8340
PMP1611     TCCTGAAGAAGTTGAAGAGGTGCTGCAAGTGCCACTTCTAGGAGTCGTTCCAGATTTGGA   8340
            ************************************************************

PMP1610     TAAAATGAAATAGGAGGAAGTTATGCCAACGTTAGAAATCTCACAGGTGAAATTAGAACT   8400
PMP1666     TAAAATGAAATAGGAGGAAGTTATGCCAACGTTAGAAATCTCACAGGTGAAATTAGAACT   8400
PMP1665     TAAAATGAAATAGGAGGAAGTTATGCCAACGTTAGAAATCTCACAGGTGAAATTAGAACT   8400
PMP1667     TAAAATGAAATAGGAGGAAGTTATGCCAACGTTAGAAATCTCACAGGTGAAATTAGAACT   8400
PMP1663     TAAAATGAAATAGGAGGAAGTTATGCCAACGTTAGAAATCTCACAGGTGAAATTAGAACT   8400
PMP1664     TAAAATGAAATAGGAGGAAGTTATGCCAACGTTAGAAATCTCACAGGTGAAATTAGAACT   8400
PMP1662     TAAAATGAAATAGGAGGAAGTTATGCCAACGTTAGAAATCTCACAGGTGAAATTAGAACT   8400
PMP1611     TAAAATGAAATAGGAGGAAGTTATGCCAACGTTAGAAATCTCACAGGTGAAATTAGAACT   8400
            ************************************************************

PMP1610     TGCGAAAAAGGCAGAGGAATATTATAATGCTTTGTGCACGAACCTACAGTTAAGTGGAGA   8460
PMP1666     TGCGAAAAAGGCAGAGGAATATTATAATGCTTTGTGCACGAACCTACAGTTAAGTGGAGA   8460
PMP1665     TGCGAAAAAGGCAGAGGAATATTATAATGCTTTGTGCACGAACCTACAGTTAAGTGGAGA   8460
PMP1667     TGCGAAAAAGGCAGAGGAATATTATAATGCTTTGTGCACGAACCTACAGTTAAGTGGAGA   8460
```

```
PMP1663       TGCGAAAAAGGCAGAGGAATATTATAATGCTTTGTGCACGAACCTACAGTTAAGTGGAGA    8460
PMP1664       TGCGAAAAAGGCAGAGGAATATTATAATGCTTTGTGCACGAACCTACAGTTAAGTGGAGA    8460
PMP1662       TGCGAAAAAGGCAGAGGAATATTATAATGCTTTGTGCACGAACCTACAGTTAAGTGGAGA    8460
PMP1611       TGCGAAAAAGGCAGAGGAATATTATAATGCTTTGTGCACGAACCTACAGTTAAGTGGAGA    8460
              ************************************************************

PMP1610       TGATTTGAAAGTATTTTCTATCACTTCTGTGAAACCAGGAGAAGGAAAAACAACGACTTC    8520
PMP1666       TGATTTGAAAGTATTTTCTATCACTTCTGTGAAACCAGGAGAAGGAAAAACAACGACTTC    8520
PMP1665       TGATTTGAAAGTATTTTCTATCACTTCTGTGAAACCAGGAGAAGGAAAAACAACGACTTC    8520
PMP1667       TGATTTGAAAGTATTTTCTATCACTTCTGTGAAACCAGGAGAAGGAAAAACAACGACTTC    8520
PMP1663       TGATTTGAAAGTATTTTCTATCACTTCTGTGAAACCAGGAGAAGGAAAAACAACGACTTC    8520
PMP1664       TGATTTGAAAGTATTTTCTATCACTTCTGTGAAACCAGGAGAAGGAAAAACAACGACTTC    8520
PMP1662       TGATTTGAAAGTATTTTCTATCACTTCTGTGAAACCAGGAGAAGGAAAAACAACGACTTC    8520
PMP1611       TGATTTGAAAGTATTTTCTATCACTTCTGTGAAACCAGGAGAAGGAAAAACAACGACTTC    8520
              ************************************************************

PMP1610       CACCAATATCGCTCGGGCTTTTGCGCGTGCAGGTTACAAAACGTTGCTGATTGATGCAGA    8580
PMP1666       CACCAATATCGCTCGGGCTTTTGCGCGTGCAGGTTACAAAACGTTGCTGATTGATGCAGA    8580
PMP1665       CACCAATATCGCTCGGGCTTTTGCGCGTGCAGGTTACAAAACGTTGCTGATTGATGCAGA    8580
PMP1667       CACCAATATCGCTCGGGCTTTTGCGCGTGCAGGTTACAAAACGTTGCTGATTGATGCAGA    8580
PMP1663       CACCAATATCGCTCGGGCTTTTGCGCGTGCAGGTTACAAAACGTTGCTGATTGATGCAGA    8580
PMP1664       CACCAATATCGCTCGGGCTTTTGCGCGTGCAGGTTACAAAACGTTGCTGATTGATGCAGA    8580
PMP1662       CACCAATATCGCTCGGGCTTTTGCGCGTGCAGGTTACAAAACGTTGCTGATTGATGCAGA    8580
PMP1611       CACCAATATCGCTCGGGCTTTTGCGCGTGCAGGTTACAAAACGTTGCTGATTGATGCAGA    8580
              ************************************************************

PMP1610       CATGCGTAACTCCGTGATGTCAGGTGTCTTTAAATCAAGGGAAAGGATTACCGGGCTAAC    8640
PMP1666       CATGCGTAACTCCGTGATGTCAGGTGTCTTTAAATCAAGGGAAAGGATTACCGGGCTAAC    8640
PMP1665       CATGCGTAACTCCGTGATGTCAGGTGTCTTTAAATCAAGGGAAAGGATTACCGGGCTAAC    8640
PMP1667       CATGCGTAACTCCGTGATGTCAGGTGTCTTTAAATCAAGGGAAAGGATTACCGGGCTAAC    8640
PMP1663       CATGCGTAACTCCGTGATGTCAGGTGTCTTTAAATCAAGGGAAAGGATTACCGGGCTAAC    8640
PMP1664       CATGCGTAACTCCGTGATGTCAGGTGTCTTTAAATCAAGGGAAAGGATTACCGGGCTAAC    8640
PMP1662       CATGCGTAACTCCGTGATGTCAGGTGTCTTTAAATCAAGGGAAAGGATTACCGGGCTAAC    8640
PMP1611       CATGCGTAACTCCGTGATGTCAGGTGTCTTTAAATCAAGGGAAAGGATTACCGGGCTAAC    8640
              ************************************************************
```

```
PMP1610      AGAATTTCTATCAGGGACTACAGACCTGTCACAGGGACTTTGTGATACCAATATTGAGAA    8700
PMP1666      AGAATTTCTATCAGGGACTACAGACCTGTCACAGGGACTTTGTGATACCAATATTGAGAA    8700
PMP1665      AGAATTTCTATCAGGGACTACAGACCTGTCACAGGGACTTTGTGATACCAATATTGAGAA    8700
PMP1667      AGAATTTCTATCAGGGACTACAGACCTGTCACAGGGACTTTGTGATACCAATATTGAGAA    8700
PMP1663      AGAATTTCTATCAGGGACTACAGACCTGTCACAGGGACTTTGTGATACCAATATTGAGAA    8700
PMP1664      AGAATTTCTATCAGGGACTACAGACCTGTCACAGGGACTTTGTGATACCAATATTGAGAA    8700
PMP1662      AGAATTTCTATCAGGGACTACAGACCTGTCACAGGGACTTTGTGATACCAATATTGAGAA    8700
PMP1611      AGAATTTCTATCAGGGACTACAGACCTGTCACAGGGACTTTGTGATACCAATATTGAGAA    8700
             ************************************************************

PMP1610      TCTCTTTGTGATTCAGGCTGGCTCTGTGTCACCAAATCCGACAGCCCTTCTTCAAAGTAG    8760
PMP1666      TCTCTTTGTGATTCAGGCTGGCTCTGTGTCACCAAATCCGACAGCCCTTCTTCAAAGTAG    8760
PMP1665      TCTCTTTGTGATTCAGGCTGGCTCTGTGTCACCAAATCCGACAGCCCTTCTTCAAAGTAG    8760
PMP1667      TCTCTTTGTGATTCAGGCTGGCTCTGTGTCACCAAATCCGACAGCCCTTCTTCAAAGTAG    8760
PMP1663      TCTCTTTGTGATTCAGGCTGGCTCTGTGTCACCAAATCCGACAGCCCTTCTTCAAAGTAG    8760
PMP1664      TCTCTTTGTGATTCAGGCTGGCTCTGTGTCACCAAATCCGACAGCCCTTCTTCAAAGTAG    8760
PMP1662      TCTCTTTGTGATTCAGGCTGGCTCTGTGTCACCAAATCCGACAGCCCTTCTTCAAAGTAG    8760
PMP1611      TCTCTTTGTGATTCAGGCTGGCTCTGTGTCACCAAATCCGACAGCCCTTCTTCAAAGTAG    8760
             ************************************************************

PMP1610      GAATTTCAGTACAATGCTTGAAACCGTGCGTAAATATTTTGACTATATCGTCGTAGACAC    8820
PMP1666      GAATTTCAGTACAATGCTTGAAACCGTGCGTAAATATTTTGACTATATCGTCGTAGACAC    8820
PMP1665      GAATTTCAGTACAATGCTTGAAACCGTGCGTAAATATTTTGACTATATCGTCGTAGACAC    8820
PMP1667      GAATTTCAGTACAATGCTTGAAACCGTGCGTAAATATTTTGACTATATCGTCGTAGACAC    8820
PMP1663      GAATTTCAGTACAATGCTTGAAACCGTGCGTAAATATTTTGACTATATCGTCGTAGACAC    8820
PMP1664      GAATTTCAGTACAATGCTTGAAACCGTGCGTAAATATTTTGACTATATCGTCGTAGACAC    8820
PMP1662      GAATTTCAGTACAATGCTTGAAACCGTGCGTAAATATTTTGACTATATCGTCGTAGACAC    8820
PMP1611      GAATTTCAGTACAATGCTTGAAACCGTGCGTAAATATTTTGACTATATCGTCGTAGACAC    8820
             ************************************************************

PMP1610      TGCTCCTGTCGGTGTCGTGATTGATGCGGCTATCATTACGCAGAAATGTGATGCTTCTAT    8880
PMP1666      TGCTCCTGTCGGTGTCGTGATTGATGCGGCTATCATTACGCAGAAATGTGATGCTTCTAT    8880
PMP1665      TGCTCCTGTCGGTGTCGTGATTGATGCGGCTATCATTACGCAGAAATGTGATGCTTCTAT    8880
PMP1667      TGCTCCTGTCGGTGTCGTGATTGATGCGGCTATCATTACGCAGAAATGTGATGCTTCTAT    8880
PMP1663      TGCTCCTGTCGGTGTCGTGATTGATGCGGCTATCATTACGCAGAAATGTGATGCTTCTAT    8880
PMP1664      TGCTCCTGTCGGTGTCGTGATTGATGCGGCTATCATTACGCAGAAATGTGATGCTTCTAT    8880
```

```
PMP1662      TGCTCCTGTCGGTGTCGTGATTGATGCGGCTATCATTACGCAGAAATGTGATGCTTCTAT   8880
PMP1611      TGCTCCTGTCGGTGTCGTGATTGATGCGGCTATCATTACGCAGAAATGTGATGCTTCTAT   8880
             ************************************************************

PMP1610      TTTAGTGACGAAGGCAGGCGAAACAAAGCGACGGGATATTCAAAAAGCGAAAGAACAGAT   8940
PMP1666      TTTAGTGACGAAGGCAGGCGAAACAAAGCGACGGGATATTCAAAAAGCGAAAGAACAGAT   8940
PMP1665      TTTAGTGACGAAGGCAGGCGAAACAAAGCGACGGGATATTCAAAAAGCGAAAGAACAGAT   8940
PMP1667      TTTAGTGACGAAGGCAGGCGAAACAAAGCGACGGGATATTCAAAAAGCGAAAGAACAGAT   8940
PMP1663      TTTAGTGACGAAGGCAGGCGAAACAAAGCGACGGGATATTCAAAAAGCGAAAGAACAGAT   8940
PMP1664      TTTAGTGACGAAGGCAGGCGAAACAAAGCGACGGGATATTCAAAAAGCGAAAGAACAGAT   8940
PMP1662      TTTAGTGACGAAGGCAGGCGAAACAAAGCGACGGGATATTCAAAAAGCGAAAGAACAGAT   8940
PMP1611      TTTAGTGACGAAGGCAGGCGAAACAAAGCGACGGGATATTCAAAAAGCGAAAGAACAGAT   8940
             ************************************************************

PMP1610      AGAACAAACTGGGAAGCCGTGTTTAGGAGTTGTGTTGAATAAATTCGATACTTCAGTAGA   9000
PMP1666      AGAACAAACTGGGAAGCCGTGTTTAGGAGTTGTGTTGAATAAATTCGATACTTCAGTAGA   9000
PMP1665      AGAACAAACTGGGAAGCCGTGTTTAGGAGTTGTGTTGAATAAATTCGATACTTCAGTAGA   9000
PMP1667      AGAACAAACTGGGAAGCCGTGTTTAGGAGTTGTGTTGAATAAATTCGATACTTCAGTAGA   9000
PMP1663      AGAACAAACTGGGAAGCCGTGTTTAGGAGTTGTGTTGAATAAATTCGATACTTCAGTAGA   9000
PMP1664      AGAACAAACTGGGAAGCCGTGTTTAGGAGTTGTGTTGAATAAATTCGATACTTCAGTAGA   9000
PMP1662      AGAACAAACTGGGAAGCCGTGTTTAGGAGTTGTGTTGAATAAATTCGATACTTCAGTAGA   9000
PMP1611      AGAACAAACTGGGAAGCCGTGTTTAGGAGTTGTGTTGAATAAATTCGATACTTCAGTAGA   9000
             ************************************************************

PMP1610      CGAATACGGTTCTTATGGAAGTTATGGGAAAAAGAAAAAATAGGTTGGGGGATAGAGATG   9060
PMP1666      CGAATACGGTTCTTATGGAAGTTATGGGAAAAAGAAAAAATAGGTTGGGGGATAGAGATG   9060
PMP1665      CGAATACGGTTCTTATGGAAGTTATGGGAAAAAGAAAAAATAGGTTGGGGGATAGAGATG   9060
PMP1667      CGAATACGGTTCTTATGGAAGTTATGGGAAAAAGAAAAAATAGGTTGGGGGATAGAGATG   9060
PMP1663      CGAATACGGTTCTTATGGAAGTTATGGGAAAAAGAAAAAATAGGTTGGGGGATAGAGATG   9060
PMP1664      CGAATACGGTTCTTATGGAAGTTATGGGAAAAAGAAAAAATAGGTTGGGGGATAGAGATG   9060
PMP1662      CGAATACGGTTCTTATGGAAGTTATGGGAAAAAGAAAAAATAGGTTGGGGGATAGAGATG   9060
PMP1611      CGAATACGGTTCTTATGGAAGTTATGGGAAAAAGAAAAAATAGGTTGGGGGATAGAGATG   9060
             ************************************************************

PMP1610      AATGGAAAACTAGTAAAGCCTTCATTGGCCATAGTCCAGAGTTTTCTTGTTATTTTATTG   9120
PMP1666      AATGGAAAACTAGTAAAGCCTTCATTGGCCATAGTCCAGAGTTTTCTTGTTATTTTATTG   9120
```

```
PMP1665      AATGGAAAACTAGTAAAGCCTTCATTGGCCATAGTCCAGAGTTTTCTTGTTATTTTATTG   9120
PMP1667      AATGGAAAACTAGTAAAGCCTTCATTGGCCATAGTCCAGAGTTTTCTTGTTATTTTATTG   9120
PMP1663      AATGGAAAACTAGTAAAGCCTTCATTGGCCATAGTCCAGAGTTTTCTTGTTATTTTATTG   9120
PMP1664      AATGGAAAACTAGTAAAGCCTTCATTGGCCATAGTCCAGAGTTTTCTTGTTATTTTATTG   9120
PMP1662      AATGGAAAACTAGTAAAGCCTTCATTGGCCATAGTCCAGAGTTTTCTTGTTATTTTATTG   9120
PMP1611      AATGGAAAACTAGTAAAGCCTTCATTGGCCATAGTCCAGAGTTTTCTTGTTATTTTATTG   9120
             ************************************************************

PMP1610      GCTTATCTACTTAGCACTGTGAGAGAAACAGAGATTGTTTCAACAACAGCTATTGTACTT   9180
PMP1666      GCTTATCTACTTAGCACTGTGAGAGAAACAGAGATTGTTTCAACAACAGCTATTGTACTT   9180
PMP1665      GCTTATCTACTTAGCACTGTGAGAGAAACAGAGATTGTTTCAACAACAGCTATTGTACTT   9180
PMP1667      GCTTATCTACTTAGCACTGTGAGAGAAACAGAGATTGTTTCAACAACAGCTATTGTACTT   9180
PMP1663      GCTTATCTACTTAGCACTGTGAGAGAAACAGAGATTGTTTCAACAACAGCTATTGTACTT   9180
PMP1664      GCTTATCTACTTAGCACTGTGAGAGAAACAGAGATTGTTTCAACAACAGCTATTGTACTT   9180
PMP1662      GCTTATCTACTTAGCACTGTGAGAGAAACAGAGATTGTTTCAACAACAGCTATTGTACTT   9180
PMP1611      GCTTATCTACTTAGCACTGTGAGAGAAACAGAGATTGTTTCAACAACAGCTATTGTACTT   9180
             ************************************************************

PMP1610      TATATCCTCCACTATTTTGCCTTTTATATCAGTGATTATGGACAGGATTTCTTTAAAAGG   9240
PMP1666      TATATCCTCCACTATTTTGCCTTTTATATCAGTGATTATGGACAGGATTTCTTTAAAAGG   9240
PMP1665      TATATCCTCCACTATTTTGCCTTTTATATCAGTGATTATGGACAGGATTTCTTTAAAAGG   9240
PMP1667      TATATCCTCCACTATTTTGCCTTTTATATCAGTGATTATGGACAGGATTTCTTTAAAAGG   9240
PMP1663      TATATCCTCCACTATTTTGCCTTTTATATCAGTGATTATGGACAGGATTTCTTTAAAAGG   9240
PMP1664      TATATCCTCCACTATTTTGCCTTTTATATCAGTGATTATGGACAGGATTTCTTTAAAAGG   9240
PMP1662      TATATCCTCCACTATTTTGCCTTTTATATCAGTGATTATGGACAGGATTTCTTTAAAAGG   9240
PMP1611      TATATCCTCCACTATTTTGCCTTTTATATCAGTGATTATGGACAGGATTTCTTTAAAAGG   9240
             ************************************************************

PMP1610      GGATATTTGATTGAACTTGTCCAGACATTGAAATATATCCTATTCTTTGCGCTAGCGATT   9300
PMP1666      GGATATTTGATTGAACTTGTCCAGACATTGAAATATATCCTATTCTTTGCGCTAGCGATT   9300
PMP1665      GGATATTTGATTGAACTTGTCCAGACATTGAAATATATCCTATTCTTTGCGCTAGCGATT   9300
PMP1667      GGATATTTGATTGAACTTGTCCAGACATTGAAATATATCCTATTCTTTGCGCTAGCGATT   9300
PMP1663      GGATATTTGATTGAACTTGTCCAGACATTGAAATATATCCTATTCTTTGCGCTAGCGATT   9300
PMP1664      GGATATTTGATTGAACTTGTCCAGACATTGAAATATATCCTATTCTTTGCGCTAGCGATT   9300
PMP1662      GGATATTTGATTGAACTTGTCCAGACATTGAAATATATCCTATTCTTTGCGCTAGCGATT   9300
PMP1611      GGATATTTGATTGAACTTGTCCAGACATTGAAATATATCCTATTCTTTGCGCTAGCGATT   9300
```

```
               ************************************************************

PMP1610        AGTATTTCTAATTTTTTCTTAGAAGATCGATTTAGTATTTCCAGAAGAGGCATGATTTAC    9360
PMP1666        AGTATTTCTAATTTTTTCTTAGAAGATCGATTTAGTATTTCCAGAAGAGGCATGATTTAC    9360
PMP1665        AGTATTTCTAATTTTTTCTTAGAAGATCGATTTAGTATTTCCAGAAGAGGCATGATTTAC    9360
PMP1667        AGTATTTCTAATTTTTTCTTAGAAGATCGATTTAGTATTTCCAGAAGAGGCATGATTTAC    9360
PMP1663        AGTATTTCTAATTTTTTCTTAGAAGATCGATTTAGTATTTCCAGAAGAGGCATGATTTAC    9360
PMP1664        AGTATTTCTAATTTTTTCTTAGAAGATCGATTTAGTATTTCCAGAAGAGGCATGATTTAC    9360
PMP1662        AGTATTTCTAATTTTTTCTTAGAAGATCGATTTAGTATTTCCAGAAGAGGCATGATTTAC    9360
PMP1611        AGTATTTCTAATTTTTTCTTAGAAGATCGATTTAGTATTTCCAGAAGAGGCATGATTTAC    9360
               ************************************************************

PMP1610        TTCCTCCTATTACATGTTTTCTTAGTCTATGTGCTAAACCGATTTATCAAGTGGTATTGG    9420
PMP1666        TTCCTCCTATTACATGTTTTCTTAGTCTATGTGCTAAACCGATTTATCAAGTGGTATTGG    9420
PMP1665        TTCCTCCTATTACATGTTTTCTTAGTCTATGTGCTAAACCGATTTATCAAGTGGTATTGG    9420
PMP1667        TTCCTCCTATTACATGTTTTCTTAGTCTATGTGCTAAACCGATTTATCAAGTGGTATTGG    9420
PMP1663        TTCCTCCTATTACATGTTTTCTTAGTCTATGTGCTAAACCGATTTATCAAGTGGTATTGG    9420
PMP1664        TTCCTCCTATTACATGTTTTCTTAGTCTATGTGCTAAACCGATTTATCAAGTGGTATTGG    9420
PMP1662        TTCCTCCTATTACATGTTTTCTTAGTCTATGTGCTAAACCGATTTATCAAGTGGTATTGG    9420
PMP1611        TTCCTCCTATTACATGTTTTCTTAGTCTATGTGCTAAACCGATTTATCAAGTGGTATTGG    9420
               ************************************************************

PMP1610        AAGCGGGCTTATCCCAACTTTAAAGGAAGTAAGAAGATTCTCCTACTTACAGCAACTTCT    9480
PMP1666        AAGCGGGCTTATCCCAACTTTAAAGGAAGTAAGAAGATTCTCCTACTTACAGCAACTTCT    9480
PMP1665        AAGCGGGCTTATCCCAACTTTAAAGGAAGTAAGAAGATTCTCCTACTTACAGCAACTTCT    9480
PMP1667        AAGCGGGCTTATCCCAACTTTAAAGGAAGTAAGAAGATTCTCCTACTTACAGCAACTTCT    9480
PMP1663        AAGCGGGCTTATCCCAACTTTAAAGGAAGTAAGAAGATTCTCCTACTTACAGCAACTTCT    9480
PMP1664        AAGCGGGCTTATCCCAACTTTAAAGGAAGTAAGAAGATTCTCCTACTTACAGCAACTTCT    9480
PMP1662        AAGCGGGCTTATCCCAACTTTAAAGGAAGTAAGAAGATTCTCCTACTTACAGCAACTTCT    9480
PMP1611        AAGCGGGCTTATCCCAACTTTAAAGGAAGTAAGAAGATTCTCCTACTTACAGCAACTTCT    9480
               ************************************************************

PMP1610        CGTGTCGAAAAGGTATTGGATAGACTAATAGAATCAGATGATGTTGTTGGGGAGTTGGTA    9540
PMP1666        CGTGTCGAAAAGGTATTGGATAGACTAATAGAATCAGATGATGTTGTTGGGGAGTTGGTA    9540
PMP1665        CGTGTCGAAAAGGTATTGGATAGACTAATAGAATCAGATGATGTTGTTGGGGAGTTGGTA    9540
PMP1667        CGTGTCGAAAAGGTATTGGATAGACTAATAGAATCAGATGATGTTGTTGGGGAGTTGGTA    9540
```

```
PMP1663      CGTGTCGAAAAGGTATTGGATAGACTAATAGAATCAGATGATGTTGTTGGGGAGTTGGTA   9540
PMP1664      CGTGTCGAAAAGGTATTGGATAGACTAATAGAATCAGATGATGTTGTTGGGGAGTTGGTA   9540
PMP1662      CGTGTCGAAAAGGTATTGGATAGACTAATAGAATCAGATGATGTTGTTGGGGAGTTGGTA   9540
PMP1611      CGTGTCGAAAAGGTATTGGATAGACTAATAGAATCAGATGATGTTGTTGGGGAGTTGGTA   9540
             ************************************************************

PMP1610      GCCGTCAGTGTTTTAGATAAACCAGATTTTCAGCATGATTATTTAAAGGTTGTAGCAGAG   9600
PMP1666      GCCGTCAGTGTTTTAGATAAACCAGATTTTCAGCATGATTATTTAAAGGTTGTAGCAGAG   9600
PMP1665      GCCGTCAGTGTTTTAGATAAACCAGATTTTCAGCATGATTATTTAAAGGTTGTAGCAGAG   9600
PMP1667      GCCGTCAGTGTTTTAGATAAACCAGATTTTCAGCATGATTATTTAAAGGTTGTAGCAGAG   9600
PMP1663      GCCGTCAGTGTTTTAGATAAACCAGATTTTCAGCATGATTATTTAAAGGTTGTAGCAGAG   9600
PMP1664      GCCGTCAGTGTTTTAGATAAACCAGATTTTCAGCATGATTATTTAAAGGTTGTAGCAGAG   9600
PMP1662      GCCGTCAGTGTTTTAGATAAACCAGATTTTCAGCATGATTATTTAAAGGTTGTAGCAGAG   9600
PMP1611      GCCGTCAGTGTTTTAGATAAACCAGATTTTCAGCATGATTATTTAAAGGTTGTAGCAGAG   9600
             ************************************************************

PMP1610      GGGGAGATCGTAAACTTTGCGACTCATGAGGTGGTCGATGAAGTCTTTATCAATCTTCCA   9660
PMP1666      GGGGAGATCGTAAACTTTGCGACTCATGAGGTGGTCGATGAAGTCTTTATCAATCTTCCA   9660
PMP1665      GGGGAGATCGTAAACTTTGCGACTCATGAGGTGGTCGATGAAGTCTTTATCAATCTTCCA   9660
PMP1667      GGGGAGATCGTAAACTTTGCGACTCATGAGGTGGTCGATGAAGTCTTTATCAATCTTCCA   9660
PMP1663      GGGGAGATCGTAAACTTTGCGACTCATGAGGTGGTCGATGAAGTCTTTATCAATCTTCCA   9660
PMP1664      GGGGAGATCGTAAACTTTGCGACTCATGAGGTGGTCGATGAAGTCTTTATCAATCTTCCA   9660
PMP1662      GGGGAGATCGTAAACTTTGCGACTCATGAGGTGGTCGATGAAGTCTTTATCAATCTTCCA   9660
PMP1611      GGGGAGATCGTAAACTTTGCGACTCATGAGGTGGTCGATGAAGTCTTTATCAATCTTCCA   9660
             ************************************************************

PMP1610      AGTGAAAAAATACAATATTGGAGAGCTTGTCTCTCAGTTTGAAACGATGGGAGTTGATGTA   9720
PMP1666      AGTGAAAAAATACAATATTGGAGAGCTTGTCTCTCAGTTTGAAACGATGGGAGTTGATGTA   9720
PMP1665      AGTGAAAAAATACAATATTGGAGAGCTTGTCTCTCAGTTTGAAACGATGGGAGTTGATGTA   9720
PMP1667      AGTGAAAAAATACAATATTGGAGAGCTTGTCTCTCAGTTTGAAACGATGGGAGTTGATGTA   9720
PMP1663      AGTGAAAAAATACAATATTGGAGAGCTTGTCTCTCAGTTTGAAACGATGGGAGTTGATGTA   9720
PMP1664      AGTGAAAAAATACAATATTGGAGAGCTTGTCTCTCAGTTTGAAACGATGGGAGTTGATGTA   9720
PMP1662      AGTGAAAAAATACAATATTGGAGAGCTTGTCTCTCAGTTTGAAACGATGGGAGTTGATGTA   9720
PMP1611      AGTGAAAAAATACAATATTGGAGAGCTTGTCTCTCAGTTTGAAACGATGGGAGTTGATGTA   9720
             ************************************************************
```

```
PMP1610      ACAGTCAATCTAAATGCTTTCGATTGTATCTTGGCACATAACAAGCAAATTTGTGAGATG    9780
PMP1666      ACAGTCAATCTAAATGCTTTCGATTGTATCTTGGCACATAACAAGCAAATTTGTGAGATG    9780
PMP1665      ACAGTCAATCTAAATGCTTTCGATTGTATCTTGGCACATAACAAGCAAATTTGTGAGATG    9780
PMP1667      ACAGTCAATCTAAATGCTTTCGATTGTATCTTGGCACATAACAAGCAAATTTGTGAGATG    9780
PMP1663      ACAGTCAATCTAAATGCTTTCGATTGTATCTTGGCACATAACAAGCAAATTTGTGAGATG    9780
PMP1664      ACAGTCAATCTAAATGCTTTCGATTGTATCTTGGCACATAACAAGCAAATTTGTGAGATG    9780
PMP1662      ACAGTCAATCTAAATGCTTTCGATTGTATCTTGGCACATAACAAGCAAATTTGTGAGATG    9780
PMP1611      ACAGTCAATCTAAATGCTTTCGATTGTATCTTGGCACATAACAAGCAAATTTGTGAGATG    9780
             ************************************************************

PMP1610      GCAGGACTAAACGTTGTGACTTTTTCTACAACATTTTATAAGACTAGCCATGTGATTGCT    9840
PMP1666      GCAGGACTAAACGTTGTGACTTTTTCTACAACATTTTATAAGACTAGCCATGTGATTGCT    9840
PMP1665      GCAGGACTAAACGTTGTGACTTTTTCTACAACATTTTATAAGACTAGCCATGTGATTGCT    9840
PMP1667      GCAGGACTAAACGTTGTGACTTTTTCTACAACATTTTATAAGACTAGCCATGTGATTGCT    9840
PMP1663      GCAGGACTAAACGTTGTGACTTTTTCTACAACATTTTATAAGACTAGCCATGTGATTGCT    9840
PMP1664      GCAGGACTAAACGTTGTGACTTTTTCTACAACATTTTATAAGACTAGCCATGTGATTGCT    9840
PMP1662      GCAGGACTAAACGTTGTGACTTTTTCTACAACATTTTATAAGACTAGCCATGTGATTGCT    9840
PMP1611      GCAGGACTAAACGTTGTGACTTTTTCTACAACATTTTATAAGACTAGCCATGTGATTGCT    9840
             ************************************************************

PMP1610      AAGCGGGTTATTGATATTATCGGTTCCCTGGTAGGTTTGATACTATGTGGTCTAGTCAGT    9900
PMP1666      AAGCGGGTTATTGATATTATCGGTTCCCTGGTAGGTTTGATACTATGTGGTCTAGTCAGT    9900
PMP1665      AAGCGGGTTATTGATATTATCGGTTCCCTGGTAGGTTTGATACTATGTGGTCTAGTCAGT    9900
PMP1667      AAGCGGGTTATTGATATTATCGGTTCCCTGGTAGGTTTGATACTATGTGGTCTAGTCAGT    9900
PMP1663      AAGCGGGTTATTGATATTATCGGTTCCCTGGTAGGTTTGATACTATGTGGTCTAGTCAGT    9900
PMP1664      AAGCGGGTTATTGATATTATCGGTTCCCTGGTAGGTTTGATACTATGTGGTCTAGTCAGT    9900
PMP1662      AAGCGGGTTATTGATATTATCGGTTCCCTGGTAGGTTTGATACTATGTGGTCTAGTCAGT    9900
PMP1611      AAGCGGGTTATTGATATTATCGGTTCCCTGGTAGGTTTGATACTATGTGGTCTAGTCAGT    9900
             ************************************************************

PMP1610      ATTGTACTGGTTCCTTTGATTCGAAAGGATGGGGGCTCTGCTATTTTTGCTCAGACGCGT    9960
PMP1666      ATTGTACTGGTTCCTTTGATTCGAAAGGATGGGGGCTCTGCTATTTTTGCTCAGACGCGT    9960
PMP1665      ATTGTACTGGTTCCTTTGATTCGAAAGGATGGGGGCTCTGCTATTTTTGCTCAGACGCGT    9960
PMP1667      ATTGTACTGGTTCCTTTGATTCGAAAGGATGGGGGCTCTGCTATTTTTGCTCAGACGCGT    9960
PMP1663      ATTGTACTGGTTCCTTTGATTCGAAAGGATGGGGGCTCTGCTATTTTTGCTCAGACGCGT    9960
PMP1664      ATTGTACTGGTTCCTTTGATTCGAAAGGATGGGGGCTCTGCTATTTTTGCTCAGACGCGT    9960
```

```
PMP1662      ATTGTACTGGTTCCTTTGATTCGAAAGGATGGGGGCTCTGCTATTTTTGCTCAGACGCGT    9960
PMP1611      ATTGTACTGGTTCCTTTGATTCGAAAGGATGGGGGCTCTGCTATTTTTGCTCAGACGCGT    9960
             ************************************************************

PMP1610      ATTGGGAAAAATGGTCGCCATTTCACTTTTTACAAGTTTCGCTCTATGTGTGTGGATGCT    10020
PMP1666      ATTGGGAAAAATGGTCGCCATTTCACTTTTTACAAGTTTCGCTCTATGTGTGTGGATGCT    10020
PMP1665      ATTGGGAAAAATGGTCGCCATTTCACTTTTTACAAGTTTCGCTCTATGTGTGTGGATGCT    10020
PMP1667      ATTGGGAAAAATGGTCGCCATTTCACTTTTTACAAGTTTCGCTCTATGTGTGTGGATGCT    10020
PMP1663      ATTGGGAAAAATGGTCGCCATTTCACTTTTTACAAGTTTCGCTCTATGTGTGTGGATGCT    10020
PMP1664      ATTGGGAAAAATGGTCGCCATTTCACTTTTTACAAGTTTCGCTCTATGTGTGTGGATGCT    10020
PMP1662      ATTGGGAAAAATGGTCGCCATTTCACTTTTTACAAGTTTCGCTCTATGTGTGTGGATGCT    10020
PMP1611      ATTGGGAAAAATGGTCGCCATTTCACTTTTTACAAGTTTCGCTCTATGTGTGTGGATGCT    10020
             ************************************************************

PMP1610      GAGGAGAAAAAAAGAGAACTCATGGAACAAAATACCATGCAGGGTGGAATGTTTAAGGTG    10080
PMP1666      GAGGAGAAAAAAAGAGAACTCATGGAACAAAATACCATGCAGGGTGGAATGTTTAAGGTG    10080
PMP1665      GAGGAGAAAAAAAGAGAACTCATGGAACAAAATACCATGCAGGGTGGAATGTTTAAGGTG    10080
PMP1667      GAGGAGAAAAAAAGAGAACTCATGGAACAAAATACCATGCAGGGTGGAATGTTTAAGGTG    10080
PMP1663      GAGGAGAAAAAAAGAGAACTCATGGAACAAAATACCATGCAGGGTGGAATGTTTAAGGTG    10080
PMP1664      GAGGAGAAAAAAAGAGAACTCATGGAACAAAATACCATGCAGGGTGGAATGTTTAAGGTG    10080
PMP1662      GAGGAGAAAAAAAGAGAACTCATGGAACAAAATACCATGCAGGGTGGAATGTTTAAGGTG    10080
PMP1611      GAGGAGAAAAAAAGAGAACTCATGGAACAAAATACCATGCAGGGTGGAATGTTTAAGGTG    10080
             ************************************************************

PMP1610      GATGAGGATCCACGTATCACGAAAATTGGTCATTTTATACGGAAGACGAGCTTGGACGAG    10140
PMP1666      GATGAGGATCCACGTATCACGAAAATTGGTCATTTTATACGGAAGACGAGCTTGGACGAG    10140
PMP1665      GATGAGGATCCACGTATCACGAAAATTGGTCATTTTATACGGAAGACGAGCTTGGACGAG    10140
PMP1667      GATGAGGATCCACGTATCACGAAAATTGGTCATTTTATACGGAAGACGAGCTTGGACGAG    10140
PMP1663      GATGAGGATCCACGTATCACGAAAATTGGTCATTTTATACGGAAGACGAGCTTGGACGAG    10140
PMP1664      GATGAGGATCCACGTATCACGAAAATTGGTCATTTTATACGGAAGACGAGCTTGGACGAG    10140
PMP1662      GATGAGGATCCACGTATCACGAAAATTGGTCATTTTATACGGAAGACGAGCTTGGACGAG    10140
PMP1611      GATGAGGATCCACGTATCACGAAAATTGGTCATTTTATACGGAAGACGAGCTTGGACGAG    10140
             ************************************************************

PMP1610      CTACCACAGTTTTACAATGTTCTAAAGGGAGATATGAGTTTGGTAGGGACACGACCACCA    10200
PMP1666      CTACCACAGTTTTACAATGTTCTAAAGGGAGATATGAGTTTGGTAGGGACACGACCACCA    10200
```

```
PMP1665      CTACCACAGTTTTACAATGTTCTAAAGGGAGATATGAGTTTGGTAGGGACACGACCACCA   10200
PMP1667      CTACCACAGTTTTACAATGTTCTAAAGGGAGATATGAGTTTGGTAGGGACACGACCACCA   10200
PMP1663      CTACCACAGTTTTACAATGTTCTAAAGGGAGATATGAGTTTGGTAGGGACACGACCACCA   10200
PMP1664      CTACCACAGTTTTACAATGTTCTAAAGGGAGATATGAGTTTGGTAGGGACACGACCACCA   10200
PMP1662      CTACCACAGTTTTACAATGTTCTAAAGGGAGATATGAGTTTGGTAGGGACACGACCACCA   10200
PMP1611      CTACCACAGTTTTACAATGTTCTAAAGGGAGATATGAGTTTGGTAGGGACACGACCACCA   10200
             ************************************************************

PMP1610      ACAGTGGACGAGTATGAGCACTATACCCCAGAACAAAAACGTCGCCTAAGTTTTAAACCT   10260
PMP1666      ACAGTGGACGAGTATGAGCACTATACCCCAGAACAAAAACGTCGCCTAAGTTTTAAACCT   10260
PMP1665      ACAGTGGACGAGTATGAGCACTATACCCCAGAACAAAAACGTCGCCTAAGTTTTAAACCT   10260
PMP1667      ACAGTGGACGAGTATGAGCACTATACCCCAGAACAAAAACGTCGCCTAAGTTTTAAACCT   10260
PMP1663      ACAGTGGACGAGTATGAGCACTATACCCCAGAACAAAAACGTCGCCTAAGTTTTAAACCT   10260
PMP1664      ACAGTGGACGAGTATGAGCACTATACCCCAGAACAAAAACGTCGCCTAAGTTTTAAACCT   10260
PMP1662      ACAGTGGACGAGTATGAGCACTATACCCCAGAACAAAAACGTCGCCTAAGTTTTAAACCT   10260
PMP1611      ACAGTGGACGAGTATGAGCACTATACCCCAGAACAAAAACGTCGCCTAAGTTTTAAACCT   10260
             ************************************************************

PMP1610      GGCATAACAGGTCTATGGCAGGTCAGCGGACGAAGCGAGATCAAGAATTTCGATGAGGTT   10320
PMP1666      GGCATAACAGGTCTATGGCAGGTCAGCGGACGAAGCGAGATCAAGAATTTCGATGAGGTT   10320
PMP1665      GGCATAACAGGTCTATGGCAGGTCAGCGGACGAAGCGAGATCAAGAATTTCGATGAGGTT   10320
PMP1667      GGCATAACAGGTCTATGGCAGGTCAGCGGACGAAGCGAGATCAAGAATTTCGATGAGGTT   10320
PMP1663      GGCATAACAGGTCTATGGCAGGTCAGCGGACGAAGCGAGATCAAGAATTTCGATGAGGTT   10320
PMP1664      GGCATAACAGGTCTATGGCAGGTCAGCGGACGAAGCGAGATCAAGAATTTCGATGAGGTT   10320
PMP1662      GGCATAACAGGTCTATGGCAGGTCAGCGGACGAAGCGAGATCAAGAATTTCGATGAGGTT   10320
PMP1611      GGCATAACAGGTCTATGGCAGGTCAGCGGACGAAGCGAGATCAAGAATTTCGATGAGGTT   10320
             ************************************************************

PMP1610      GTCAAATTAGATGTGGCCTATATAGACGGTTGGACAATTTGGAAAGATATTGAAATTTTA   10380
PMP1666      GTCAAATTAGATGTGGCCTATATAGACGGTTGGACAATTTGGAAAGATATTGAAATTTTA   10380
PMP1665      GTCAAATTAGATGTGGCCTATATAGACGGTTGGACAATTTGGAAAGATATTGAAATTTTA   10380
PMP1667      GTCAAATTAGATGTGGCCTATATAGACGGTTGGACAATTTGGAAAGATATTGAAATTTTA   10380
PMP1663      GTCAAATTAGATGTGGCCTATATAGACGGTTGGACAATTTGGAAAGATATTGAAATTTTA   10380
PMP1664      GTCAAATTAGATGTGGCCTATATAGACGGTTGGACAATTTGGAAAGATATTGAAATTTTA   10380
PMP1662      GTCAAATTAGATGTGGCCTATATAGACGGTTGGACAATTTGGAAAGATATTGAAATTTTA   10380
PMP1611      GTCAAATTAGATGTGGCCTATATAGACGGTTGGACAATTTGGAAAGATATTGAAATTTTA   10380
```

```
            ************************************************************

PMP1610     TTGAAGACAGTTAAAGTTGTATTGATGAAGGATGGAGCGAAGTAGAGAGTTTCTCTCTGA   10440
PMP1666     TTGAAGACAGTTAAAGTTGTATTGATGAAGGATGGAGCGAAGTAGAGAGTTTCTCTCTGA   10440
PMP1665     TTGAAGACAGTTAAAGTTGTATTGATGAAGGATGGAGCGAAGTAGAGAGTTTCTCTCTGA   10440
PMP1667     TTGAAGACAGTTAAAGTTGTATTGATGAAGGATGGAGCGAAGTAGAGAGTTTCTCTCTGA   10440
PMP1663     TTGAAGACAGTTAAAGTTGTATTGATGAAGGATGGAGCGAAGTAGAGAGTTTCTCTCTGA   10440
PMP1664     TTGAAGACAGTTAAAGTTGTATTGATGAAGGATGGAGCGAAGTAGAGAGTTTCTCTCTGA   10440
PMP1662     TTGAAGACAGTTAAAGTTGTATTGATGAAGGATGGAGCGAAGTAGAGAGTTTCTCTCTGA   10440
PMP1611     TTGAAGACAGTTAAAGTTGTATTGATGAAGGATGGAGCGAAGTAGAGAGTTTCTCTCTGA   10440
            ************************************************************

PMP1610     CCACAATCGTGCAATCCTAATAATAGCGAGATAATAAAATTGTCTTGCTTTTAACTCGGA   10500
PMP1666     CCACAATCGTGCAATCCTAATAATAGCGAGATAATAAAATTGTCTTGCTTTTAACTCGGA   10500
PMP1665     CCACAATCGTGCAATCCTAATAATAGCGAGATAATAAAATTGTCTTGCTTTTAACTCGGA   10500
PMP1667     CCACAATCGTGCAATCCTAATAATAGCGAGATAATAAAATTGTCTTGCTTTTAACTCGGA   10500
PMP1663     CCACAATCGTGCAATCCTAATAATAGCGAGATAATAAAATTGTCTTGCTTTTAACTCGGA   10500
PMP1664     CCACAATCGTGCAATCCTAATAATAGCGAGATAATAAAATTGTCTTGCTTTTAACTCGGA   10500
PMP1662     CCACAATCGTGCAATCCTAATAATAGCGAGATAATAAAATTGTCTTGCTTTTAACTCGGA   10500
PMP1611     CCACAATCGTGCAATCCTAATAATAGCGAGATAATAAAATTGTCTTGCTTTTAACTCGGA   10500
            ************************************************************

PMP1610     CTCTAAATTTAGGAGTTCTAGTGATAATATTAACAATAAAGGTGAGGTGAAAAATGGATA   10560
PMP1666     CTCTAAATTTAGGAGTTCTAGTGATAATATTAACAATAAAGGTGAGGTGAAAAATGGATA   10560
PMP1665     CTCTAAATTTAGGAGTTCTAGTGATAATATTAACAATAAAGGTGAGGTGAAAAATGGATA   10560
PMP1667     CTCTAAATTTAGGAGTTCTAGTGATAATATTAACAATAAAGGTGAGGTGAAAAATGGATA   10560
PMP1663     CTCTAAATTTAGGAGTTCTAGTGATAATATTAACAATAAAGGTGAGGTGAAAAATGGATA   10560
PMP1664     CTCTAAATTTAGGAGTTCTAGTGATAATATTAACAATAAAGGTGAGGTGAAAAATGGATA   10560
PMP1662     CTCTAAATTTAGGAGTTCTAGTGATAATATTAACAATAAAGGTGAGGTGAAAAATGGATA   10560
PMP1611     CTCTAAATTTAGGAGTTCTAGTGATAATATTAACAATAAAGGTGAGGTGAAAAATGGATA   10560
            ************************************************************

PMP1610     TAGTATATGCGACAGATAATAATTTTGTAGATGTATTGAGTGCATCTATCAAGTCGCTTT   10620
PMP1666     TAGTATATGCGACAGATAATAATTTTGTAGATGTATTGAGTGCATCTATCAAGTCGCTTT   10620
PMP1665     TAGTATATGCGACAGATAATAATTTTGTAGATGTATTGAGTGCATCTATCAAGTCGCTTT   10620
PMP1667     TAGTATATGCGACAGATAATAATTTTGTAGATGTATTGAGTGCATCTATCAAGTCGCTTT   10620
```

```
PMP1663      TAGTATATGCGACAGATAATAATTTTGTAGATGTATTGAGTGCATCTATCAAGTCGCTTT   10620
PMP1664      TAGTATATGCGACAGATAATAATTTTGTAGATGTATTGAGTGCATCTATCAAGTCGCTTT   10620
PMP1662      TAGTATATGCGACAGATAATAATTTTGTAGATGTATTGAGTGCATCTATCAAGTCGCTTT   10620
PMP1611      TAGTATATGCGACAGATAATAATTTTGTAGATGTATTGAGTGCATCTATCAAGTCGCTTT   10620
             ************************************************************

PMP1610      ACACTACTAATTCAGATTTGGATTTAAATTTATGGATTATTGCTGATAAAGTTTCGGATA   10680
PMP1666      ACACTACTAATTCAGATTTGGATTTAAATTTATGGATTATTGCTGATAAAGTTTCGGATA   10680
PMP1665      ACACTACTAATTCAGATTTGGATTTAAATTTATGGATTATTGCTGATAAAGTTTCGGATA   10680
PMP1667      ACACTACTAATTCAGATTTGGATTTAAATTTATGGATTATTGCTGATAAAGTTTCGGATA   10680
PMP1663      ACACTACTAATTCAGATTTGGATTTAAATTTATGGATTATTGCTGATAAAGTTTCGGATA   10680
PMP1664      ACACTACTAATTCAGATTTGGATTTAAATTTATGGATTATTGCTGATAAAGTTTCGGATA   10680
PMP1662      ACACTACTAATTCAGATTTGGATTTAAATTTATGGATTATTGCTGATAAAGTTTCGGATA   10680
PMP1611      ACACTACTAATTCAGATTTGGATTTAAATTTATGGATTATTGCTGATAAAGTTTCGGATA   10680
             ************************************************************

PMP1610      GAAATAAAGAAAAGATAAATAGATTATCAAAACAATTTGCGCAGAGAGAAATTAATTGGA   10740
PMP1666      GAAATAAAGAAAAGATAAATAGATTATCAAAACAATTTGCGCAGAGAGAAATTAATTGGA   10740
PMP1665      GAAATAAAGAAAAGATAAATAGATTATCAAAACAATTTGCGCAGAGAGAAATTAATTGGA   10740
PMP1667      GAAATAAAGAAAAGATAAATAGATTATCAAAACAATTTGCGCAGAGAGAAATTAATTGGA   10740
PMP1663      GAAATAAAGAAAAGATAAATAGATTATCAAAACAATTTGCGCAGAGAGAAATTAATTGGA   10740
PMP1664      GAAATAAAGAAAAGATAAATAGATTATCAAAACAATTTGCGCAGAGAGAAATTAATTGGA   10740
PMP1662      GAAATAAAGAAAAGATAAATAGATTATCAAAACAATTTGCGCAGAGAGAAATTAATTGGA   10740
PMP1611      GAAATAAAGAAAAGATAAATAGATTATCAAAACAATTTGCGCAGAGAGAAATTAATTGGA   10740
             ************************************************************

PMP1610      TAGAGAACGTTGAGATTCCATTTAAATTACATTTAGATAGGGGATCAATTAGTTCATTTA   10800
PMP1666      TAGAGAACGTTGAGATTCCATTTAAATTACATTTAGATAGGGGATCAATTAGTTCATTTA   10800
PMP1665      TAGAGAACGTTGAGATTCCATTTAAATTACATTTAGATAGGGGATCAATTAGTTCATTTA   10800
PMP1667      TAGAGAACGTTGAGATTCCATTTAAATTACATTTAGATAGGGGATCAATTAGTTCATTTA   10800
PMP1663      TAGAGAACGTTGAGATTCCATTTAAATTACATTTAGATAGGGGATCAATTAGTTCATTTA   10800
PMP1664      TAGAGAACGTTGAGATTCCATTTAAATTACATTTAGATAGGGGATCAATTAGTTCATTTA   10800
PMP1662      TAGAGAACGTTGAGATTCCATTTAAATTACATTTAGATAGGGGATCAATTAGTTCATTTA   10800
PMP1611      TAGAGAACGTTGAGATTCCATTTAAATTACATTTAGATAGGGGATCAATTAGTTCATTTA   10800
             ************************************************************
```

```
PMP1610      GCAGATTATTTCTGGGAAGTGTTCTTCCATCTTCAATGAGTAAAGTTCTCTATCTTGATA   10860
PMP1666      GCAGATTATTTCTGGGAAGTGTTCTTCCATCTTCAATGAGTAAAGTTCTCTATCTTGATA   10860
PMP1665      GCAGATTATTTCTGGGAAGTGTTCTTCCATCTTCAATGAGTAAAGTTCTCTATCTTGATA   10860
PMP1667      GCAGATTATTTCTGGGAAGTGTTCTTCCATCTTCAATGAGTAAAGTTCTCTATCTTGATA   10860
PMP1663      GCAGATTATTTCTGGGAAGTGTTCTTCCATCTTCAATGAGTAAAGTTCTCTATCTTGATA   10860
PMP1664      GCAGATTATTTCTGGGAAGTGTTCTTCCATCTTCAATGAGTAAAGTTCTCTATCTTGATA   10860
PMP1662      GCAGATTATTTCTGGGAAGTGTTCTTCCATCTTCAATGAGTAAAGTTCTCTATCTTGATA   10860
PMP1611      GCAGATTATTTCTGGGAAGTGTTCTTCCATCTTCAATGAGTAAAGTTCTCTATCTTGATA   10860
             ************************************************************

PMP1610      GTGATATTATTGTAATGGATTCTTTACGAAGTATTCTTGATATTGATTTTAAAGATAAAA   10920
PMP1666      GTGATATTATTGTAATGGATTCTTTACGAAGTATTCTTGATATTGATTTTAAAGATAAAA   10920
PMP1665      GTGATATTATTGTAATGGATTCTTTACGAAGTATTCTTGATATTGATTTTAAAGATAAAA   10920
PMP1667      GTGATATTATTGTAATGGATTCTTTACGAAGTATTCTTGATATTGATTTTAAAGATAAAA   10920
PMP1663      GTGATATTATTGTAATGGATTCTTTACGAAGTATTCTTGATATTGATTTTAAAGATAAAA   10920
PMP1664      GTGATATTATTGTAATGGATTCTTTACGAAGTATTCTTGATATTGATTTTAAAGATAAAA   10920
PMP1662      GTGATATTATTGTAATGGATTCTTTACGAAGTATTCTTGATATTGATTTTAAAGATAAAA   10920
PMP1611      GTGATATTATTGTAATGGATTCTTTACGAAGTATTCTTGATATTGATTTTAAAGATAAAA   10920
             ************************************************************

PMP1610      TTCTCTATGGGGTAAATGATACTTTTAATAAAGAATACAAGCAGGTGTTGGGGATACCAA   10980
PMP1666      TTCTCTATGGGGTAAATGATACTTTTAATAAAGAATACAAGCAGGTGTTGGGGATACCAA   10980
PMP1665      TTCTCTATGGGGTAAATGATACTTTTAATAAAGAATACAAGCAGGTGTTGGGGATACCAA   10980
PMP1667      TTCTCTATGGGGTAAATGATACTTTTAATAAAGAATACAAGCAGGTGTTGGGGATACCAA   10980
PMP1663      TTCTCTATGGGGTAAATGATACTTTTAATAAAGAATACAAGCAGGTGTTGGGGATACCAA   10980
PMP1664      TTCTCTATGGGGTAAATGATACTTTTAATAAAGAATACAAGCAGGTGTTGGGGATACCAA   10980
PMP1662      TTCTCTATGGGGTAAATGATACTTTTAATAAAGAATACAAGCAGGTGTTGGGGATACCAA   10980
PMP1611      TTCTCTATGGGGTAAATGATACTTTTAATAAAGAATACAAGCAGGTGTTGGGGATACCAA   10980
             ************************************************************

PMP1610      TCGACAAACCAATGTTTAATGCTGGAGTTATGCTGATTAATTTAGAGTTATGGAGAAATA   11040
PMP1666      TCGACAAACCAATGTTTAATGCTGGAGTTATGCTGATTAATTTAGAGTTATGGAGAAATA   11040
PMP1665      TCGACAAACCAATGTTTAATGCTGGAGTTATGCTGATTAATTTAGAGTTATGGAGAAATA   11040
PMP1667      TCGACAAACCAATGTTTAATGCTGGAGTTATGCTGATTAATTTAGAGTTATGGAGAAATA   11040
PMP1663      TCGACAAACCAATGTTTAATGCTGGAGTTATGCTGATTAATTTAGAGTTATGGAGAAATA   11040
PMP1664      TCGACAAACCAATGTTTAATGCTGGAGTTATGCTGATTAATTTAGAGTTATGGAGAAATA   11040
```

```
PMP1662       TCGACAAACCAATGTTTAATGCTGGAGTTATGCTGATTAATTTAGAGTTATGGAGAAATA    11040
PMP1611       TCGACAAACCAATGTTTAATGCTGGAGTTATGCTGATTAATTTAGAGTTATGGAGAAATA    11040
              ************************************************************

PMP1610       ATAACGTCGAAGAAAAATTTTTGCAAGTAATTCAAAAGTTTAATGGTACTATATTACAAG    11100
PMP1666       ATAACGTCGAAGAAAAATTTTTGCAAGTAATTCAAAAGTTTAATGGTACTATATTACAAG    11100
PMP1665       ATAACGTCGAAGAAAAATTTTTGCAAGTAATTCAAAAGTTTAATGGTACTATATTACAAG    11100
PMP1667       ATAACGTCGAAGAAAAATTTTTGCAAGTAATTCAAAAGTTTAATGGTACTATATTACAAG    11100
PMP1663       ATAACGTCGAAGAAAAATTTTTGCAAGTAATTCAAAAGTTTAATGGTACTATATTACAAG    11100
PMP1664       ATAACGTCGAAGAAAAATTTTTGCAAGTAATTCAAAAGTTTAATGGTACTATATTACAAG    11100
PMP1662       ATAACGTCGAAGAAAAATTTTTGCAAGTAATTCAAAAGTTTAATGGTACTATATTACAAG    11100
PMP1611       ATAACGTCGAAGAAAAATTTTTGCAAGTAATTCAAAAGTTTAATGGTACTATATTACAAG    11100
              ************************************************************

PMP1610       GTGATTTAGGAGTTTTAAATGCAGTTTTATATAACTCATTTGGTGTACTTCCTCCAGAAT    11160
PMP1666       GTGATTTAGGAGTTTTAAATGCAGTTTTATATAACTCATTTGGTGTACTTCCTCCAGAAT    11160
PMP1665       GTGATTTAGGAGTTTTAAATGCAGTTTTATATAACTCATTTGGTGTACTTCCTCCAGAAT    11160
PMP1667       GTGATTTAGGAGTTTTAAATGCAGTTTTATATAACTCATTTGGTGTACTTCCTCCAGAAT    11160
PMP1663       GTGATTTAGGAGTTTTAAATGCAGTTTTATATAACTCATTTGGTGTACTTCCTCCAGAAT    11160
PMP1664       GTGATTTAGGAGTTTTAAATGCAGTTTTATATAACTCATTTGGTGTACTTCCTCCAGAAT    11160
PMP1662       GTGATTTAGGAGTTTTAAATGCAGTTTTATATAACTCATTTGGTGTACTTCCTCCAGAAT    11160
PMP1611       GTGATTTAGGAGTTTTAAATGCAGTTTTATATAACTCATTTGGTGTACTTCCTCCAGAAT    11160
              ************************************************************

PMP1610       ATAATTATATGACCATATTTGAAGATTTGACTTATGAAGAAATGATAGTTTTTAAAAAAC    11220
PMP1666       ATAATTATATGACCATATTTGAAGATTTGACTTATGAAGAAATGATAGTTTTTAAAAAAC    11220
PMP1665       ATAATTATATGACCATATTTGAAGATTTGACTTATGAAGAAATGATAGTTTTTAAAAAAC    11220
PMP1667       ATAATTATATGACCATATTTGAAGATTTGACTTATGAAGAAATGATAGTTTTTAAAAAAC    11220
PMP1663       ATAATTATATGACCATATTTGAAGATTTGACTTATGAAGAAATGATAGTTTTTAAAAAAC    11220
PMP1664       ATAATTATATGACCATATTTGAAGATTTGACTTATGAAGAAATGATAGTTTTTAAAAAAC    11220
PMP1662       ATAATTATATGACCATATTTGAAGATTTGACTTATGAAGAAATGATAGTTTTTAAAAAAC    11220
PMP1611       ATAATTATATGACCATATTTGAAGATTTGACTTATGAAGAAATGATAGTTTTTAAAAAAC    11220
              ************************************************************

PMP1610       CAATTAACTATTATTCAAAAGAAGAAATTAAAAATGCTAGAGAACGTATAGTCTTGCGCC    11280
PMP1666       CAATTAACTATTATTCAAAAGAAGAAATTAAAAATGCTAGAGAACGTATAGTCTTGCGCC    11280
```

```
PMP1665      CAATTAACTATTATTCAAAAGAAGAAATTAAAAATGCTAGAGAACGTATAGTCTTGCGCC   11280
PMP1667      CAATTAACTATTATTCAAAAGAAGAAATTAAAAATGCTAGAGAACGTATAGTCTTGCGCC   11280
PMP1663      CAATTAACTATTATTCAAAAGAAGAAATTAAAAATGCTAGAGAACGTATAGTCTTGCGCC   11280
PMP1664      CAATTAACTATTATTCAAAAGAAGAAATTAAAAATGCTAGAGAACGTATAGTCTTGCGCC   11280
PMP1662      CAATTAACTATTATTCAAAAGAAGAAATTAAAAATGCTAGAGAACGTATAGTCTTGCGCC   11280
PMP1611      CAATTAACTATTATTCAAAAGAAGAAATTAAAAATGCTAGAGAACGTATAGTCTTGCGCC   11280
             ************************************************************

PMP1610      ATTTCACAACGTGTTTTCTATCACTCAGACCTTGGCAAGAAAATAGTGAGGTGGCGCATG   11340
PMP1666      ATTTCACAACGTGTTTTCTATCACTCAGACCTTGGCAAGAAAATAGTGAGGTGGCGCATG   11340
PMP1665      ATTTCACAACGTGTTTTCTATCACTCAGACCTTGGCAAGAAAATAGTGAGGTGGCGCATG   11340
PMP1667      ATTTCACAACGTGTTTTCTATCACTCAGACCTTGGCAAGAAAATAGTGAGGTGGCGCATG   11340
PMP1663      ATTTCACAACGTGTTTTCTATCACTCAGACCTTGGCAAGAAAATAGTGAGGTGGCGCATG   11340
PMP1664      ATTTCACAACGTGTTTTCTATCACTCAGACCTTGGCAAGAAAATAGTGAGGTGGCGCATG   11340
PMP1662      ATTTCACAACGTGTTTTCTATCACTCAGACCTTGGCAAGAAAATAGTGAGGTGGCGCATG   11340
PMP1611      ATTTCACAACGTGTTTTCTATCACTCAGACCTTGGCAAGAAAATAGTGAGGTGGCGCATG   11340
             ************************************************************

PMP1610      TGGAAATATTTAAAAAATACTATAGAGGAACATACAAACAAGTATCTCCATCTAAGTTAT   11400
PMP1666      TGGAAATATTTAAAAAATACTATAGAGGAACATACAAACAAGTATCTCCATCTAAGTTAT   11400
PMP1665      TGGAAATATTTAAAAAATACTATAGAGGAACATACAAACAAGTATCTCCATCTAAGTTAT   11400
PMP1667      TGGAAATATTTAAAAAATACTATAGAGGAACATACAAACAAGTATCTCCATCTAAGTTAT   11400
PMP1663      TGGAAATATTTAAAAAATACTATAGAGGAACATACAAACAAGTATCTCCATCTAAGTTAT   11400
PMP1664      TGGAAATATTTAAAAAATACTATAGAGGAACATACAAACAAGTATCTCCATCTAAGTTAT   11400
PMP1662      TGGAAATATTTAAAAAATACTATAGAGGAACATACAAACAAGTATCTCCATCTAAGTTAT   11400
PMP1611      TGGAAATATTTAAAAAATACTATAGAGGAACATACAAACAAGTATCTCCATCTAAGTTAT   11400
             ************************************************************

PMP1610      CAAGAATTTATAAAATTTTACCGAAAAAAATGTCGCTCTATTTACTAGGTTTTATTCAAT   11460
PMP1666      CAAGAATTTATAAAATTTTACCGAAAAAAATGTCGCTCTATTTACTAGGTTTTATTCAAT   11460
PMP1665      CAAGAATTTATAAAATTTTACCGAAAAAAATGTCGCTCTATTTACTAGGTTTTATTCAAT   11460
PMP1667      CAAGAATTTATAAAATTTTACCGAAAAAAATGTCGCTCTATTTACTAGGTTTTATTCAAT   11460
PMP1663      CAAGAATTTATAAAATTTTACCGAAAAAAATGTCGCTCTATTTACTAGGTTTTATTCAAT   11460
PMP1664      CAAGAATTTATAAAATTTTACCGAAAAAAATGTCGCTCTATTTACTAGGTTTTATTCAAT   11460
PMP1662      CAAGAATTTATAAAATTTTACCGAAAAAAATGTCGCTCTATTTACTAGGTTTTATTCAAT   11460
PMP1611      CAAGAATTTATAAAATTTTACCGAAAAAAATGTCGCTCTATTTACTAGGTTTTATTCAAT   11460
```

```
              ************************************************************

PMP1610       CAAAAGTGCGTCCAAAACTGTATAGAATTTTGAAATAAGATGGCTAGTAGTAATGGCAAA   11520
PMP1666       CAAAAGTGCGTCCAAAACTGTATAGAATTTTGAAATAAGATGGCTAGTAGTAATGGCAAA   11520
PMP1665       CAAAAGTGCGTCCAAAACTGTATAGAATTTTGAAATAAGATGGCTAGTAGTAATGGCAAA   11520
PMP1667       CAAAAGTGCGTCCAAAACTGTATAGAATTTTGAAATAAGATGGCTAGTAGTAATGGCAAA   11520
PMP1663       CAAAAGTGCGTCCAAAACTGTATAGAATTTTGAAATAAGATGGCTAGTAGTAATGGCAAA   11520
PMP1664       CAAAAGTGCGTCCAAAACTGTATAGAATTTTGAAATAAGATGGCTAGTAGTAATGGCAAA   11520
PMP1662       CAAAAGTGCGTCCAAAACTGTATAGAATTTTGAAATAAGATGGCTAGTAGTAATGGCAAA   11520
PMP1611       CAAAAGTGCGTCCAAAACTGTATAGAATTTTGAAATAAGATGGCTAGTAGTAATGGCAAA   11520
              ************************************************************

PMP1610       AAAGAAAGAATTTATTGAAAGTTCAAGAAAAATTGAAATTATTAAAAGAGTTTATAAAAA   11580
PMP1666       AAAGAAAGAATTTATTGAAAGTTCAAGAAAAATTGAAATTATTAAAAGAGTTTATAAAAA   11580
PMP1665       AAAGAAAGAATTTATTGAAAGTTCAAGAAAAATTGAAATTATTAAAAGAGTTTATAAAAA   11580
PMP1667       AAAGAAAGAATTTATTGAAAGTTCAAGAAAAATTGAAATTATTAAAAGAGTTTATAAAAA   11580
PMP1663       AAAGAAAGAATTTATTGAAAGTTCAAGAAAAATTGAAATTATTAAAAGAGTTTATAAAAA   11580
PMP1664       AAAGAAAGAATTTATTGAAAGTTCAAGAAAAATTGAAATTATTAAAAGAGTTTATAAAAA   11580
PMP1662       AAAGAAAGAATTTATTGAAAGTTCAAGAAAAATTGAAATTATTAAAAGAGTTTATAAAAA   11580
PMP1611       AAAGAAAGAATTTATTGAAAGTTCAAGAAAAATTGAAATTATTAAAAGAGTTTATAAAAA   11580
              ************************************************************

PMP1610       TATGTTCTAAAAATAAAATGAAGTATTTTGCTTCTGGAGGTAGTCTATTGGAAGCTGCAA   11640
PMP1666       TATGTTCTAAAAATAAAATGAAGTATTTTGCTTCTGGAGGTAGTCTATTGGAAGCTGCAA   11640
PMP1665       TATGTTCTAAAAATAAAATGAAGTATTTTGCTTCTGGAGGTAGTCTATTGGAAGCTGCAA   11640
PMP1667       TATGTTCTAAAAATAAAATGAAGTATTTTGCTTCTGGAGGTAGTCTATTGGAAGCTGCAA   11640
PMP1663       TATGTTCTAAAAATAAAATGAAGTATTTTGCTTCTGGAGGTAGTCTATTGGAAGCTGCAA   11640
PMP1664       TATGTTCTAAAAATAAAATGAAGTATTTTGCTTCTGGAGGTAGTCTATTGGAAGCTGCAA   11640
PMP1662       TATGTTCTAAAAATAAAATGAAGTATTTTGCTTCTGGAGGTAGTCTATTGGAAGCTGCAA   11640
PMP1611       TATGTTCTAAAAATAAAATGAAGTATTTTGCTTCTGGAGGTAGTCTATTGGAAGCTGCAA   11640
              ************************************************************

PMP1610       GATACAAAGGTTTTATTCCTTGGAATGATGATATGGCTCTAGGTTTACCAAGAAAGCATT   11700
PMP1666       GATACAAAGGTTTTATTCCTTGGAATGATGATATGGCTCTAGGTTTACCAAGAAAGCATT   11700
PMP1665       GATACAAAGGTTTTATTCCTTGGAATGATGATATGGCTCTAGGTTTACCAAGAAAGCATT   11700
PMP1667       GATACAAAGGTTTTATTCCTTGGAATGATGATATGGCTCTAGGTTTACCAAGAAAGCATT   11700
```

```
PMP1663     GATACAAAGGTTTTATTCCTTGGAATGATGATATGGCTCTAGGTTTACCAAGAAAGCATT   11700
PMP1664     GATACAAAGGTTTTATTCCTTGGAATGATGATATGGCTCTAGGTTTACCAAGAAAGCATT   11700
PMP1662     GATACAAAGGTTTTATTCCTTGGAATGATGATATGGCTCTAGGTTTACCAAGAAAGCATT   11700
PMP1611     GATACAAAGGTTTTATTCCTTGGAATGATGATATGGCTCTAGGTTTACCAAGAAAGCATT   11700
            ************************************************************

PMP1610     TTGAAAAGTTTATAAATGAAATAGATTTTGAAAAATATAATAAAAATTATATTTTAGAAA   11760
PMP1666     TTGAAAAGTTTATAAATGAAATAGATTTTGAAAAATATAATAAAAATTATATTTTAGAAA   11760
PMP1665     TTGAAAAGTTTATAAATGAAATAGATTTTGAAAAATATAATAAAAATTATATTTTAGAAA   11760
PMP1667     TTGAAAAGTTTATAAATGAAATAGATTTTGAAAAATATAATAAAAATTATATTTTAGAAA   11760
PMP1663     TTGAAAAGTTTATAAATGAAATAGATTTTGAAAAATATAATAAAAATTATATTTTAGAAA   11760
PMP1664     TTGAAAAGTTTATAAATGAAATAGATTTTGAAAAATATAATAAAAATTATATTTTAGAAA   11760
PMP1662     TTGAAAAGTTTATAAATGAAATAGATTTTGAAAAATATAATAAAAATTATATTTTAGAAA   11760
PMP1611     TTGAAAAGTTTATAAATGAAATAGATTTTGAAAAATATAATAAAAATTATATTTTAGAAA   11760
            ************************************************************

PMP1610     GTTCGGAAATGAATTTAGGTATTTTTCAATATAAATTAAAATCTGCTATTTTAATTTTAG   11820
PMP1666     GTTCGGAAATGAATTTAGGTATTTTTCAATATAAATTAAAATCTGCTATTTTAATTTTAG   11820
PMP1665     GTTCGGAAATGAATTTAGGTATTTTTCAATATAAATTAAAATCTGCTATTTTAATTTTAG   11820
PMP1667     GTTCGGAAATGAATTTAGGTATTTTTCAATATAAATTAAAATCTGCTATTTTAATTTTAG   11820
PMP1663     GTTCGGAAATGAATTTAGGTATTTTTCAATATAAATTAAAATCTGCTATTTTAATTTTAG   11820
PMP1664     GTTCGGAAATGAATTTAGGTATTTTTCAATATAAATTAAAATCTGCTATTTTAATTTTAG   11820
PMP1662     GTTCGGAAATGAATTTAGGTATTTTTCAATATAAATTAAAATCTGCTATTTTAATTTTAG   11820
PMP1611     GTTCGGAAATGAATTTAGGTATTTTTCAATATAAATTAAAATCTGCTATTTTAATTTTAG   11820
            ************************************************************

PMP1610     GGAAATCGTATGACGTATGTTTGAATTTATTTCTGTTAGACAGGATGTCAGAATTGGTTA   11880
PMP1666     GGAAATCGTATGACGTATGTTTGAATTTATTTCTGTTAGACAGGATGTCAGAATTGGTTA   11880
PMP1665     GGAAATCGTATGACGTATGTTTGAATTTATTTCTGTTAGACAGGATGTCAGAATTGGTTA   11880
PMP1667     GGAAATCGTATGACGTATGTTTGAATTTATTTCTGTTAGACAGGATGTCAGAATTGGTTA   11880
PMP1663     GGAAATCGTATGACGTATGTTTGAATTTATTTCTGTTAGACAGGATGTCAGAATTGGTTA   11880
PMP1664     GGAAATCGTATGACGTATGTTTGAATTTATTTCTGTTAGACAGGATGTCAGAATTGGTTA   11880
PMP1662     GGAAATCGTATGACGTATGTTTGAATTTATTTCTGTTAGACAGGATGTCAGAATTGGTTA   11880
PMP1611     GGAAATCGTATGACGTATGTTTGAATTTATTTCTGTTAGACAGGATGTCAGAATTGGTTA   11880
            ************************************************************
```

```
PMP1610      TTGATGGTGAGTCAGGTTTTAAAGTCCCACTATATAATTTAGAAGTGGCTGTAGATAGAA   11940
PMP1666      TTGATGGTGAGTCAGGTTTTAAAGTCCCACTATATAATTTAGAAGTGGCTGTAGATAGAA   11940
PMP1665      TTGATGGTGAGTCAGGTTTTAAAGTCCCACTATATAATTTAGAAGTGGCTGTAGATAGAA   11940
PMP1667      TTGATGGTGAGTCAGGTTTTAAAGTCCCACTATATAATTTAGAAGTGGCTGTAGATAGAA   11940
PMP1663      TTGATGGTGAGTCAGGTTTTAAAGTCCCACTATATAATTTAGAAGTGGCTGTAGATAGAA   11940
PMP1664      TTGATGGTGAGTCAGGTTTTAAAGTCCCACTATATAATTTAGAAGTGGCTGTAGATAGAA   11940
PMP1662      TTGATGGTGAGTCAGGTTTTAAAGTCCCACTATATAATTTAGAAGTGGCTGTAGATAGAA   11940
PMP1611      TTGATGGTGAGTCAGGTTTTAAAGTCCCACTATATAATTTAGAAGTGGCTGTAGATAGAA   11940
             ************************************************************

PMP1610      GTAGAAGTATTATTGAGAATAGAGAACTAGCCAATGAGTTAGGTAGTGTTGCTTTCCAAA   12000
PMP1666      GTAGAAGTATTATTGAGAATAGAGAACTAGCCAATGAGTTAGGTAGTGTTGCTTTCCAAA   12000
PMP1665      GTAGAAGTATTATTGAGAATAGAGAACTAGCCAATGAGTTAGGTAGTGTTGCTTTCCAAA   12000
PMP1667      GTAGAAGTATTATTGAGAATAGAGAACTAGCCAATGAGTTAGGTAGTGTTGCTTTCCAAA   12000
PMP1663      GTAGAAGTATTATTGAGAATAGAGAACTAGCCAATGAGTTAGGTAGTGTTGCTTTCCAAA   12000
PMP1664      GTAGAAGTATTATTGAGAATAGAGAACTAGCCAATGAGTTAGGTAGTGTTGCTTTCCAAA   12000
PMP1662      GTAGAAGTATTATTGAGAATAGAGAACTAGCCAATGAGTTAGGTAGTGTTGCTTTCCAAA   12000
PMP1611      GTAGAAGTATTATTGAGAATAGAGAACTAGCCAATGAGTTAGGTAGTGTTGCTTTCCAAA   12000
             ************************************************************

PMP1610      GAGTTCAATCTATATTTGAAATAAAAGAAAAAGTGTCAGAGCTAGAGAATATATTCATGA   12060
PMP1666      GAGTTCAATCTATATTTGAAATAAAAGAAAAAGTGTCAGAGCTAGAGAATATATTCATGA   12060
PMP1665      GAGTTCAATCTATATTTGAAATAAAAGAAAAAGTGTCAGAGCTAGAGAATATATTCATGA   12060
PMP1667      GAGTTCAATCTATATTTGAAATAAAAGAAAAAGTGTCAGAGCTAGAGAATATATTCATGA   12060
PMP1663      GAGTTCAATCTATATTTGAAATAAAAGAAAAAGTGTCAGAGCTAGAGAATATATTCATGA   12060
PMP1664      GAGTTCAATCTATATTTGAAATAAAAGAAAAAGTGTCAGAGCTAGAGAATATATTCATGA   12060
PMP1662      GAGTTCAATCTATATTTGAAATAAAAGAAAAAGTGTCAGAGCTAGAGAATATATTCATGA   12060
PMP1611      GAGTTCAATCTATATTTGAAATAAAAGAAAAAGTGTCAGAGCTAGAGAATATATTCATGA   12060
             ************************************************************

PMP1610      GTTTAGGAGAAGATGATAATGTCAATATATAAACTTTATAAAGATATTGAAAGAAAAACG   12120
PMP1666      GTTTAGGAGAAGATGATAATGTCAATATATAAACTTTATAAAGATATTGAAAGAAAAACG   12120
PMP1665      GTTTAGGAGAAGATGATAATGTCAATATATAAACTTTATAAAGATATTGAAAGAAAAACG   12120
PMP1667      GTTTAGGAGAAGATGATAATGTCAATATATAAACTTTATAAAGATATTGAAAGAAAAACG   12120
PMP1663      GTTTAGGAGAAGATGATAATGTCAATATATAAACTTTATAAAGATATTGAAAGAAAAACG   12120
PMP1664      GTTTAGGAGAAGATGATAATGTCAATATATAAACTTTATAAAGATATTGAAAGAAAAACG   12120
```

```
PMP1662      GTTTAGGAGAAGATGATAATGTCAATATATAAACTTTATAAAGATATTGAAAGAAAAACG    12120
PMP1611      GTTTAGGAGAAGATGATAATGTCAATATATAAACTTTATAAAGATATTGAAAGAAAAACG    12120
             ************************************************************

PMP1610      ATGTCGCCTGCTAAAAAAGCAATGGCTAAAAATGATTATTTTGCGTTTTATGTTGGAAGA    12180
PMP1666      ATGTCGCCTGCTAAAAAAGCAATGGCTAAAAATGATTATTTTGCGTTTTATGTTGGAAGA    12180
PMP1665      ATGTCGCCTGCTAAAAAAGCAATGGCTAAAAATGATTATTTTGCGTTTTATGTTGGAAGA    12180
PMP1667      ATGTCGCCTGCTAAAAAAGCAATGGCTAAAAATGATTATTTTGCGTTTTATGTTGGAAGA    12180
PMP1663      ATGTCGCCTGCTAAAAAAGCAATGGCTAAAAATGATTATTTTGCGTTTTATGTTGGAAGA    12180
PMP1664      ATGTCGCCTGCTAAAAAAGCAATGGCTAAAAATGATTATTTTGCGTTTTATGTTGGAAGA    12180
PMP1662      ATGTCGCCTGCTAAAAAAGCAATGGCTAAAAATGATTATTTTGCGTTTTATGTTGGAAGA    12180
PMP1611      ATGTCGCCTGCTAAAAAAGCAATGGCTAAAAATGATTATTTTGCGTTTTATGTTGGAAGA    12180
             ************************************************************

PMP1610      CCGTTATCCTATCTTTTAACAGTTCCTTTTTTAAAAACGAATATTACTCCCAATCAAGTA    12240
PMP1666      CCGTTATCCTATCTTTTAACAGTTCCTTTTTTAAAAACGAATATTACTCCCAATCAAGTA    12240
PMP1665      CCGTTATCCTATCTTTTAACAGTTCCTTTTTTAAAAACGAATATTACTCCCAATCAAGTA    12240
PMP1667      CCGTTATCCTATCTTTTAACAGTTCCTTTTTTAAAAACGAATATTACTCCCAATCAAGTA    12240
PMP1663      CCGTTATCCTATCTTTTAACAGTTCCTTTTTTAAAAACGAATATTACTCCCAATCAAGTA    12240
PMP1664      CCGTTATCCTATCTTTTAACAGTTCCTTTTTTAAAAACGAATATTACTCCCAATCAAGTA    12240
PMP1662      CCGTTATCCTATCTTTTAACAGTTCCTTTTTTAAAAACGAATATTACTCCCAATCAAGTA    12240
PMP1611      CCGTTATCCTATCTTTTAACAGTTCCTTTTTTAAAAACGAATATTACTCCCAATCAAGTA    12240
             ************************************************************

PMP1610      TCTTATTTATCTATAGCCCCTTTGATTCTTGGATTTCTGACAATGACATTTACAACTAAT    12300
PMP1666      TCTTATTTATCTATAGCCCCTTTGATTCTTGGATTTCTGACAATGACATTTACAACTAAT    12300
PMP1665      TCTTATTTATCTATAGCCCCTTTGATTCTTGGATTTCTGACAATGACATTTACAACTAAT    12300
PMP1667      TCTTATTTATCTATAGCCCCTTTGATTCTTGGATTTCTGACAATGACATTTACAACTAAT    12300
PMP1663      TCTTATTTATCTATAGCCCCTTTGATTCTTGGATTTCTGACAATGACATTTACAACTAAT    12300
PMP1664      TCTTATTTATCTATAGCCCCTTTGATTCTTGGATTTCTGACAATGACATTTACAACTAAT    12300
PMP1662      TCTTATTTATCTATAGCCCCTTTGATTCTTGGATTTCTGACAATGACATTTACAACTAAT    12300
PMP1611      TCTTATTTATCTATAGCCCCTTTGATTCTTGGATTTCTGACAATGACATTTACAACTAAT    12300
             ************************************************************

PMP1610      TTCATTCTATTATTGCTGGCATGGTTTCTATTTTTTTTATGGAACTTACTAGATGGAGTA    12360
PMP1666      TTCATTCTATTATTGCTGGCATGGTTTCTATTTTTTTTATGGAACTTACTAGATGGAGTA    12360
```

```
PMP1665      TTCATTCTATTATTGCTGGCATGGTTTCTATTTTTTTTATGGAACTTACTAGATGGAGTA   12360
PMP1667      TTCATTCTATTATTGCTGGCATGGTTTCTATTTTTTTTATGGAACTTACTAGATGGAGTA   12360
PMP1663      TTCATTCTATTATTGCTGGCATGGTTTCTATTTTTTTTATGGAACTTACTAGATGGAGTA   12360
PMP1664      TTCATTCTATTATTGCTGGCATGGTTTCTATTTTTTTTATGGAACTTACTAGATGGAGTA   12360
PMP1662      TTCATTCTATTATTGCTGGCATGGTTTCTATTTTTTTTATGGAACTTACTAGATGGAGTA   12360
PMP1611      TTCATTCTATTATTGCTGGCATGGTTTCTATTTTTTTTATGGAACTTACTAGATGGAGTA   12360
             ************************************************************

PMP1610      GATGGGAACTTAGCTAGATATCGGGAGCAATACTCGAAGGATGGAAGTGTAGTAGATGCA   12420
PMP1666      GATGGGAACTTAGCTAGATATCGGGAGCAATACTCGAAGGATGGAAGTGTAGTAGATGCA   12420
PMP1665      GATGGGAACTTAGCTAGATATCGGGAGCAATACTCGAAGGATGGAAGTGTAGTAGATGCA   12420
PMP1667      GATGGGAACTTAGCTAGATATCGGGAGCAATACTCGAAGGATGGAAGTGTAGTAGATGCA   12420
PMP1663      GATGGGAACTTAGCTAGATATCGGGAGCAATACTCGAAGGATGGAAGTGTAGTAGATGCA   12420
PMP1664      GATGGGAACTTAGCTAGATATCGGGAGCAATACTCGAAGGATGGAAGTGTAGTAGATGCA   12420
PMP1662      GATGGGAACTTAGCTAGATATCGGGAGCAATACTCGAAGGATGGAAGTGTAGTAGATGCA   12420
PMP1611      GATGGGAACTTAGCTAGATATCGGGAGCAATACTCGAAGGATGGAAGTGTAGTAGATGCA   12420
             ************************************************************

PMP1610      ATGGCTGGCTATGTAGCCATGGTGTTGACGTATTTCGGTGCAGGAATAGTAGCTGCTCAT   12480
PMP1666      ATGGCAGGCTATGTGGCTATGGTGCTGACGTATTTTGGTGCAGGAATAGTAGCAACTCAT   12480
PMP1665      ATGGCAGGCTATGTGGCTATGGTGCTGACGTATTTTGGTGCAGGAATAGTAGCAACTCAT   12480
PMP1667      ATGGCAGGCTATGTGGCTATGGTGCTGACGTATTTTGGTGCAGGAATAGTAGCAACTCAT   12480
PMP1663      ATGGCAGGCTATGTGGCTATGGTGCTGACGTATTTTGGTGCAGGAATAGTAGCAACTCAT   12480
PMP1664      ATGGCAGGCTATGTGGCTATGGTGCTGACGTATTTTGGTGCAGGAATAGTAGCAACTCAT   12480
PMP1662      ATGGCAGGCTATGTGGCTATGGTGCTGACGTATTTTGGTGCAGGAATAGTAGCAACTCAT   12480
PMP1611      ATGGCAGGCTATGTGGCTATGGTGCTGACGTATTTTGGTGCAGGAATAGTAGCAACTCAT   12480
             ***** ******** ** ****** ********** *****************  *****

PMP1610      TTAAACGACTCAGATATCTATATAATTTTGGGTGCATTATCTGGGATTTCATTGATTTTT   12540
PMP1666      CTAAATGGCTCAGATATGTATGTGATTTTGGGTGCTTTATCTGGAATTTCTTTGATTTTT   12540
PMP1665      CTAAATGGCTCAGATATGTATGTGATTTTGGGTGCTTTATCTGGAATTTCTTTGATTTTT   12540
PMP1667      CTAAATGGCTCAGATATGTATGTGATTTTGGGTGCTTTATCTGGAATTTCTTTGATTTTT   12540
PMP1663      CTAAATGGCTCAGATATGTATGTGATTTTGGGTGCTTTATCTGGAATTTCTTTGATTTTT   12540
PMP1664      CTAAATGGCTCAGATATGTATGTGATTTTGGGTGCTTTATCTGGAATTTCTTTGATTTTT   12540
PMP1662      CTAAATGGCTCAGATATGTATGTGATTTTGGGTGCTTTATCTGGAATTTCTTTGATTTTT   12540
PMP1611      CTAAATGGCTCAGATATGTATGTGATTTTGGGTGCTTTATCTGGAATTTCTTTGATTTTT   12540
```

```
             **** * ********* *** * *********** ******** ***** *********

PMP1610      CCAAGGTTAGTGATGCATAAGTATATCAATACAGTAGCTCAAGATGAGTCTGTGAGTAGC    12600
PMP1666      CCAAGATTGGTGATGCATAAGTATATCAACACAGTAGCACGAAATGAGTCTGTCAATAAC    12600
PMP1665      CCAAGATTGGTGATGCATAAGTATATCAACACAGTAGCACGAAATGAGTCTGTCAATAAC    12600
PMP1667      CCAAGATTGGTGATGCATAAGTATATCAACACAGTAGCACGAAATGAGTCTGTCAATAAC    12600
PMP1663      CCAAGATTGGTGATGCATAAGTATATCAACACAGTAGCACGAAATGAGTCTGTCAATAAC    12600
PMP1664      CCAAGATTGGTGATGCATAAGTATATCAACACAGTAGCACGAAATGAGTCTGTCAATAAC    12600
PMP1662      CCAAGATTGGTGATGCATAAGTATATCAACACAGTAGCACGAAATGAGTCTGTCAATAAC    12600
PMP1611      CCAAGATTGGTGATGCATAAGTATATCAACACAGTAGCACGAAATGAGTCTGTCAATAAC    12600
             ***** ** ******************** ********* * * ********* * ** *

PMP1610      ATTAAAGATAAATCTGATTTTAATACTATAAAAATACTGGCTCTAAACATGACATCAATT    12660
PMP1666      ATTAAAGATAAATCAAATTTTAGTACTATAAAACTACTGGCTTTAAATATGACATCAATT    12660
PMP1665      ATTAAAGATAAATCAAATTTTAGTACTATAAAACTACTGGCTTTAAATATGACATCAATT    12660
PMP1667      ATTAAAGATAAATCAAATTTTAGTACTATAAAACTACTGGCTTTAAATATGACATCAATT    12660
PMP1663      ATTAAAGATAAATCAAATTTTAGTACTATAAAACTACTGGCTTTAAATATGACATCAATT    12660
PMP1664      ATTAAAGATAAATCAAATTTTAGTACTATAAAACTACTGGCTTTAAATATGACATCAATT    12660
PMP1662      ATTAAAGATAAATCAAATTTTAGTACTATAAAACTACTGGCTTTAAATATGACATCAATT    12660
PMP1611      ATTAAAGATAAATCAAATTTTAGTACTATAAAACTACTGGCTTTAAATATGACATCAATT    12660
             **************  ****** ********** ******** **** ************

PMP1610      ACAGGAATTCCGCAGGTTTTACTGCTATTAACTATTTTAACAAATCAGTGGGTACTTTTT    12720
PMP1666      ACAGGTATTCCTCAGGTTTTACTACTAGTAACGATTTTAACAAATCAGTGGGAATTTTTT    12720
PMP1665      ACAGGTATTCCTCAGGTTTTACTACTAGTAACGATTTTAACAAATCAGTGGGAATTTTTT    12720
PMP1667      ACAGGTATTCCTCAGGTTTTACTACTAGTAACGATTTTAACAAATCAGTGGGAATTTTTT    12720
PMP1663      ACAGGTATTCCTCAGGTTTTACTACTAGTAACGATTTTAACAAATCAGTGGGAATTTTTT    12720
PMP1664      ACAGGTATTCCTCAGGTTTTACTACTAGTAACGATTTTAACAAATCAGTGGGAATTTTTT    12720
PMP1662      ACAGGTATTCCTCAGGTTTTACTACTAGTAACGATTTTAACAAATCAGTGGGAATTTTTT    12720
PMP1611      ACAGGTATTCCTCAGGTTTTACTACTAGTAACGATTTTAACAAATCAGTGGGAATTTTTT    12720
             ***** ***** *********** *** **** ******************* * *****

PMP1610      ACTTTAGTATATTTCACGATTAATTTTTTATTAATGATATTTTCTTTGTATTCATTATTC    12780
PMP1666      ACTTTAGTATATTTCACGATTAATTTTTTATTAATGATATTTTCGTTATATTCATTATTT    12780
PMP1665      ACTTTAGTATATTTCACGATTAATTTTTTATTAATGATATTTTCGTTATATTCATTATTT    12780
PMP1667      ACTTTAGTATATTTCACGATTAATTTTTTATTAATGATATTTTCGTTATATTCATTATTT    12780
```

```
PMP1663      ACTTTAGTATATTTCACGATTAATTTTTTATTAATGATATTTTCGTTATATTCATTATTT    12780
PMP1664      ACTTTAGTATATTTCACGATTAATTTTTTATTAATGATATTTTCGTTATATTCATTATTT    12780
PMP1662      ACTTTAGTATATTTCACGATTAATTTTTTATTAATGATATTTTCGTTATATTCATTATTT    12780
PMP1611      ACTTTAGTATATTTCACGATTAATTTTTTATTAATGATATTTTCGTTATATTCATTATTT    12780
             ******************************************** ** ***********

PMP1610      AAAAAGGAGAATGTTTAGAAATGGGAAAGTCAGTTGCAATTTTAATGACCACCTATAATG    12840
PMP1666      AAAAAGGAGAATGTTTAGAGATGGGAAAATCAGTTGCAATTTTAATGACCACTTATAATG    12840
PMP1665      AAAAAGGAGAATGTTTAGAGATGGGAAAATCAGTTGCAATTTTAATGACCACTTATAATG    12840
PMP1667      AAAAAGGAGAATGTTTAGAGATGGGAAAATCAGTTGCAATTTTAATGACCACTTATAATG    12840
PMP1663      AAAAAGGAGAATGTTTAGAGATGGGAAAATCAGTTGCAATTTTAATGACCACTTATAATG    12840
PMP1664      AAAAAGGAGAATGTTTAGAGATGGGAAAATCAGTTGCAATTTTAATGACCACTTATAATG    12840
PMP1662      AAAAAGGAGAATGTTTAGAGATGGGAAAATCAGTTGCAATTTTAATGACCACTTATAATG    12840
PMP1611      AAAAAGGAGAATGTTTAGAGATGGGAAAATCAGTTGCAATTTTAATGACCACTTATAATG    12840
             ******************* ******** *********************** *******

PMP1610      GTGAGCGATATTTGTCACAACAGATTGATAGTATTAGGTCTCAAACATTCACTAATTGGA    12900
PMP1666      GTGAGAGATATTTGTCACAACAGATTGATAGTATTAGGTCTCAAACATTTACCAATTGGA    12900
PMP1665      GTGAGAGATATTTGTCACAACAGATTGATAGTATTAGGTCTCAAACATTTACCAATTGGA    12900
PMP1667      GTGAGAGATATTTGTCACAACAGATTGATAGTATTAGGTCTCAAACATTTACCAATTGGA    12900
PMP1663      GTGAGAGATATTTGTCACAACAGATTGATAGTATTAGGTCTCAAACATTTACCAATTGGA    12900
PMP1664      GTGAGAGATATTTGTCACAACAGATTGATAGTATTAGGTCTCAAACATTTACCAATTGGA    12900
PMP1662      GTGAGAGATATTTGTCACAACAGATTGATAGTATTAGGTCTCAAACATTTACCAATTGGA    12900
PMP1611      GTGAGAGATATTTGTCACAACAGATTGATAGTATTAGGTCTCAAACATTTACCAATTGGA    12900
             ***** ******************************************* ** *******

PMP1610      CGCTTTTTATTAGGGATGATGGATCAAAAGATAAAACAATAGAAGTAATACAGAGGTATT    12960
PMP1666      CGCTTTTTATTAGAGATGATGGATCAAAAGATAAGACAGTAGAAGTAATACAGAGGTATT    12960
PMP1665      CGCTTTTTATTAGAGATGATGGATCAAAAGATAAGACAGTAGAAGTAATACAGAGGTATT    12960
PMP1667      CGCTTTTTATTAGAGATGATGGATCAAAAGATAAGACAGTAGAAGTAATACAGAGGTATT    12960
PMP1663      CGCTTTTTATTAGAGATGATGGATCAAAAGATAAGACAGTAGAAGTAATACAGAGGTATT    12960
PMP1664      CGCTTTTTATTAGAGATGATGGATCAAAAGATAAGACAGTAGAAGTAATACAGAGGTATT    12960
PMP1662      CGCTTTTTATTAGAGATGATGGATCAAAAGATAAGACAGTAGAAGTAATACAGAGGTATT    12960
PMP1611      CGCTTTTTATTAGAGATGATGGATCAAAAGATAAGACAGTAGAAGTAATACAGAGGTATT    12960
             ************* ******************** *** *********************
```

```
PMP1610      CTAAGATAGATGATAGAATTAGATTAGTTGAAAATCCCTCAAAGTTTCATGGAGCTTATT   13020
PMP1666      CTAAGATAGATGATAGAATTAGATTAGTTGAAAATCCCTCAAAGTTTCATGGAGCTTATT   13020
PMP1665      CTAAGATAGATGATAGAATTAGATTAGTTGAAAATCCCTCAAAGTTTCATGGAGCTTATT   13020
PMP1667      CTAAGATAGATGATAGAATTAGATTAGTTGAAAATCCCTCAAAGTTTCATGGAGCTTATT   13020
PMP1663      CTAAGATAGATGATAGAATTAGATTAGTTGAAAATCCCTCAAAGTTTCATGGAGCTTATT   13020
PMP1664      CTAAGATAGATGATAGAATTAGATTAGTTGAAAATCCCTCAAAGTTTCATGGAGCTTATT   13020
PMP1662      CTAAGATAGATGATAGAATTAGATTAGTTGAAAATCCCTCAAAGTTTCATGGAGCTTATT   13020
PMP1611      CTAAGATAGATGATAGAATTAGATTAGTTGAAAATCCCTCAAAGTTTCATGGAGCTTATT   13020
             ************************************************************

PMP1610      ACAATTTTTTTAATCTAATTGAATACGTTAAAAACAATTATCAATTTGATTATTACTTTT   13080
PMP1666      ATAATTTTTTTAATCTAATTGAATACGTTAAAAACAATTATCAATTTGATTATTACTTTT   13080
PMP1665      ATAATTTTTTTAATCTAATTGAATACGTTAAAAACAATTATCAATTTGATTATTACTTTT   13080
PMP1667      ATAATTTTTTTAATCTAATTGAATACGTTAAAAACAATTATCAATTTGATTATTACTTTT   13080
PMP1663      ATAATTTTTTTAATCTAATTGAATACGTTAAAAACAATTATCAATTTGATTATTACTTTT   13080
PMP1664      ATAATTTTTTTAATCTAATTGAATACGTTAAAAACAATTATCAATTTGATTATTACTTTT   13080
PMP1662      ATAATTTTTTTAATCTAATTGAATACGTTAAAAACAATTATCAATTTGATTATTACTTTT   13080
PMP1611      ATAATTTTTTTAATCTAATTGAATACGTTAAAAACAATTATCAATTTGATTATTACTTTT   13080
             * **********************************************************

PMP1610      TTTGTGATCAAGATGATATTTGGAAAGAGCACAAGTTAGAAATACAGCTGTTAAGATTTT   13140
PMP1666      TTTGTGATCAAGATGATATTTGGAAAGAGCACAAGCTAGAAATACAGCTGTTAAGATTTT   13140
PMP1665      TTTGTGATCAAGATGATATTTGGAAAGAGCACAAGCTAGAAATACAGCTGTTAAGATTTT   13140
PMP1667      TTTGTGATCAAGATGATATTTGGAAAGAGCACAAGCTAGAAATACAGCTGTTAAGATTTT   13140
PMP1663      TTTGTGATCAAGATGATATTTGGAAAGAGCACAAGCTAGAAATACAGCTGTTAAGATTTT   13140
PMP1664      TTTGTGATCAAGATGATATTTGGAAAGAGCACAAGCTAGAAATACAGCTGTTAAGATTTT   13140
PMP1662      TTTGTGATCAAGATGATATTTGGAAAGAGCACAAGCTAGAAATACAGCTGTTAAGATTTT   13140
PMP1611      TTTGTGATCAAGATGATATTTGGAAAGAGCACAAGCTAGAAATACAGCTGTTAAGATTTT   13140
             *********************************** ************************

PMP1610      CTAAAGATGACATGCCAGAGATGGTTTACTCTGATCTGTCAACGATTGATGCCAGTAATA   13200
PMP1666      CTAAGGATGAAATGCCAGAGATGGTTTACTCTGATATGTCAACGATTGATGCCAATAATA   13200
PMP1665      CTAAGGATGAAATGCCAGAGATGGTTTACTCTGATATGTCAACGATTGATGCCAATAATA   13200
PMP1667      CTAAGGATGAAATGCCAGAGATGGTTTACTCTGATATGTCAACGATTGATGCCAATAATA   13200
PMP1663      CTAAGGATGAAATGCCAGAGATGGTTTACTCTGATATGTCAACGATTGATGCCAATAATA   13200
PMP1664      CTAAGGATGAAATGCCAGAGATGGTTTACTCTGATATGTCAACGATTGATGCCAATAATA   13200
```

```
PMP1662       CTAAGGATGAAATGCCAGAGATGGTTTACTCTGATATGTCAACGATTGATGCCAATAATA    13200
PMP1611       CTAAGGATGAAATGCCAGAGATGGTTTACTCTGATATGTCAACGATTGATGCCAATAATA    13200
              **** ***** ************************ ****************** *****

PMP1610       ATTTGATAGATATTAGTATAAATAAAATAATGGGGATTGAATTACCGAACATAAATAATT    13260
PMP1666       AGTTGATAGATATTAGTATAAATAACATAATGGGGATTGAATTACCGAACATAAATAATT    13260
PMP1665       AGTTGATAGATATTAGTATAAATAACATAATGGGGATTGAATTACCGAACATAAATAATT    13260
PMP1667       AGTTGATAGATATTAGTATAAATAACATAATGGGGATTGAATTACCGAACATAAATAATT    13260
PMP1663       AGTTGATAGATATTAGTATAAATAACATAATGGGGATTGAATTACCGAACATAAATAATT    13260
PMP1664       AGTTGATAGATATTAGTATAAATAACATAATGGGGATTGAATTACCGAACATAAATAATT    13260
PMP1662       AGTTGATAGATATTAGTATAAATAACATAATGGGGATTGAATTACCGAACATAAATAATT    13260
PMP1611       AGTTGATAGATATTAGTATAAATAACATAATGGGGATTGAATTACCGAACATAAATAATT    13260
              * *********************** **********************************

PMP1610       TGTATTTTATTCATGCCTATATCTGGGGGTGTACTGCAGGTTTTAATCATGCATTGCTAG    13320
PMP1666       TGTATTTTATTCATGCCTATATCTGGGGGTGTACGGCAGGCTTTAATCATGCATTGTTAG    13320
PMP1665       TGTATTTTATTCATGCCTATATCTGGGGGTGTACGGCAGGCTTTAATCATGCATTGTTAG    13320
PMP1667       TGTATTTTATTCATGCCTATATCTGGGGGTGTACGGCAGGCTTTAATCATGCATTGTTAG    13320
PMP1663       TGTATTTTATTCATGCCTATATCTGGGGGTGTACGGCAGGCTTTAATCATGCATTGTTAG    13320
PMP1664       TGTATTTTATTCATGCCTATATCTGGGGGTGTACGGCAGGCTTTAATCATGCATTGTTAG    13320
PMP1662       TGTATTTTATTCATGCCTATATCTGGGGGTGTACGGCAGGCTTTAATCATGCATTGTTAG    13320
PMP1611       TGTATTTTATTCATGCCTATATCTGGGGGTGTACGGCAGGCTTTAATCATGCATTGTTAG    13320
              ********************************** ***** *************** ***

PMP1610       AGATGGTTCCTTCAGTTGATATTGATAAAGATTATTTATATATAGAAAAACTGTCTCATG    13380
PMP1666       AGATGGTTCCTTCAGTTGATATTGATAAAGATTATTTATATATAGAAAAACTGTCTCATG    13380
PMP1665       AGATGGTTCCTTCAGTTGATATTGATAAAGATTATTTATATATAGAAAAACTGTCTCATG    13380
PMP1667       AGATGGTTCCTTCAGTTGATATTGATAAAGATTATTTATATATAGAAAAACTGTCTCATG    13380
PMP1663       AGATGGTTCCTTCAGTTGATATTGATAAAGATTATTTATATATAGAAAAACTGTCTCATG    13380
PMP1664       AGATGGTTCCTTCAGTTGATATTGATAAAGATTATTTATATATAGAAAAACTGTCTCATG    13380
PMP1662       AGATGGTTCCTTCAGTTGATATTGATAAAGATTATTTATATATAGAAAAACTGTCTCATG    13380
PMP1611       AGATGGTTCCTTCAGTTGATATTGATAAAGATTATTTATATATAGAAAAACTGTCTCATG    13380
              ************************************************************

PMP1610       ATAATTATTTTGCAAAGTTTGCACTAGAGTATGGGAAGGTGTTGTTCTGCCCTGAGCAAC    13440
PMP1666       ATAATTATTTTGCAAAGTTTGCACTAGAGTATGGGAAGGTGTTGTTCTGCCCTGAGCAAC    13440
```

```
PMP1665       ATAATTATTTTGCAAAGTTTGCACTAGAGTATGGGAAGGTGTTGTTCTGCCCTGAGCAAC    13440
PMP1667       ATAATTATTTTGCAAAGTTTGCACTAGAGTATGGGAAGGTGTTGTTCTGCCCTGAGCAAC    13440
PMP1663       ATAATTATTTTGCAAAGTTTGCACTAGAGTATGGGAAGGTGTTGTTCTGCCCTGAGCAAC    13440
PMP1664       ATAATTATTTTGCAAAGTTTGCACTAGAGTATGGGAAGGTGTTGTTCTGCCCTGAGCAAC    13440
PMP1662       ATAATTATTTTGCAAAGTTTGCACTAGAGTATGGGAAGGTGTTGTTCTGCCCTGAGCAAC    13440
PMP1611       ATAATTATTTTGCAAAGTTTGCACTAGAGTATGGGAAGGTGTTGTTCTGCCCTGAGCAAC    13440
              ************************************************************

PMP1610       TGGTCTTGTATCGAAGACATGGACATAATGTAACAACTAGTCATCATTTTAAATTATCTC    13500
PMP1666       TGGTCTTGTATCGAAGACACGGACATAATGTAACAACTAGTCATCATTTTAAATTATCTC    13500
PMP1665       TGGTCTTGTATCGAAGACACGGACATAATGTAACAACTAGTCATCATTTTAAATTATCTC    13500
PMP1667       TGGTCTTGTATCGAAGACACGGACATAATGTAACAACTAGTCATCATTTTAAATTATCTC    13500
PMP1663       TGGTCTTGTATCGAAGACACGGACATAATGTAACAACTAGTCATCATTTTAAATTATCTC    13500
PMP1664       TGGTCTTGTATCGAAGACACGGACATAATGTAACAACTAGTCATCATTTTAAATTATCTC    13500
PMP1662       TGGTCTTGTATCGAAGACACGGACATAATGTAACAACTAGTCATCATTTTAAATTATCTC    13500
PMP1611       TGGTCTTGTATCGAAGACACGGACATAATGTAACAACTAGTCATCATTTTAAATTATCTC    13500
              ******************* ****************************************

PMP1610       CGCTAAATGTTTTCAGAAAGGCTATATTGGGTTTCAATGAATTGGCACTTACACATGCTG    13560
PMP1666       CGCTAAATATTCTCAGAAAGGCTATTTTGGGTTTCAATGAATTGGCACTTACACATGCTG    13560
PMP1665       CGCTAAATATTCTCAGAAAGGCTATTTTGGGTTTCAATGAATTGGCACTTACACATGCTG    13560
PMP1667       CGCTAAATATTCTCAGAAAGGCTATTTTGGGTTTCAATGAATTGGCACTTACACATGCTG    13560
PMP1663       CGCTAAATATTCTCAGAAAGGCTATTTTGGGTTTCAATGAATTGGCACTTACACATGCTG    13560
PMP1664       CGCTAAATATTCTCAGAAAGGCTATTTTGGGTTTCAATGAATTGGCACTTACACATGCTG    13560
PMP1662       CGCTAAATATTCTCAGAAAGGCTATTTTGGGTTTCAATGAATTGGCACTTACACATGCTG    13560
PMP1611       CGCTAAATATTCTCAGAAAGGCTATTTTGGGTTTCAATGAATTGGCACTTACACATGCTG    13560
              ******** ** ************ ***********************************

PMP1610       GGGTATATAATCAAACTCTTTATATGCTAAAAAAAGCTTCTGGAAAAAATCCTTTAAGTG    13620
PMP1666       GGGTATATAATCAAACTCTTTATATGCTAAAAAAAGCTTCTGAAAAAAGTCCTTTAAGTG    13620
PMP1665       GGGTATATAATCAAACTCTTTATATGCTAAAAAAAGCTTCTGAAAAAAGTCCTTTAAGTG    13620
PMP1667       GGGTATATAATCAAACTCTTTATATGCTAAAAAAAGCTTCTGAAAAAAGTCCTTTAAGTG    13620
PMP1663       GGGTATATAATCAAACTCTTTATATGCTAAAAAAAGCTTCTGAAAAAAGTCCTTTAAGTG    13620
PMP1664       GGGTATATAATCAAACTCTTTATATGCTAAAAAAAGCTTCTGAAAAAAGTCCTTTAAGTG    13620
PMP1662       GGGTATATAATCAAACTCTTTATATGCTAAAAAAAGCTTCTGAAAAAAGTCCTTTAAGTG    13620
PMP1611       GGGTATATAATCAAACTCTTTATATGCTAAAAAAAGCTTCTGAAAAAAGTCCTTTAAGTG    13620
```

```
              ****************************************** ***** ***********

PMP1610       ATAGACTACTTGAAATTCAGGAAGTAATCAAAATTGGAGGATTAAAAGGTGTGAGATATT   13680
PMP1666       ATAGACTGCTTGAAATTCAGGAAGTAATCAAAATTGGAGGATTAAAAGGTGTGAGATATT   13680
PMP1665       ATAGACTGCTTGAAATTCAGGAAGTAATCAAAATTGGAGGATTAAAAGGTGTGAGATATT   13680
PMP1667       ATAGACTGCTTGAAATTCAGGAAGTAATCAAAATTGGAGGATTAAAAGGTGTGAGATATT   13680
PMP1663       ATAGACTGCTTGAAATTCAGGAAGTAATCAAAATTGGAGGATTAAAAGGTGTGAGATATT   13680
PMP1664       ATAGACTGCTTGAAATTCAGGAAGTAATCAAAATTGGAGGATTAAAAGGTGTGAGATATT   13680
PMP1662       ATAGACTGCTTGAAATTCAGGAAGTAATCAAAATTGGAGGATTAAAAGGTGTGAGATATT   13680
PMP1611       ATAGACTGCTTGAAATTCAGGAAGTAATCAAAATTGGAGGATTAAAAGGTGTGAGATATT   13680
              ******* ****************************************************

PMP1610       TCTATCAGAATCGAATTTCTCGAAAGCAACTCGTAAGAACCATCGGCTTATATACCATCA   13740
PMP1666       TCTGTCAGAATCGAATTTCTCGAAAGCAACTCGTAAGAACCATCGGCTTATATACCATCA   13740
PMP1665       TCTGTCAGAATCGAATTTCTCGAAAGCAACTCGTAAGAACCATCGGCTTATATACCATCA   13740
PMP1667       TCTGTCAGAATCGAATTTCTCGAAAGCAACTCGTAAGAACCATCGGCTTATATACCATCA   13740
PMP1663       TCTGTCAGAATCGAATTTCTCGAAAGCAACTCGTAAGAACCATCGGCTTATATACCATCA   13740
PMP1664       TCTGTCAGAATCGAATTTCTCGAAAGCAACTCGTAAGAACCATCGGCTTATATACCATCA   13740
PMP1662       TCTGTCAGAATCGAATTTCTCGAAAGCAACTCGTAAGAACCATCGGCTTATATACCATCA   13740
PMP1611       TCTGTCAGAATCGAATTTCTCGAAAGCAACTCGTAAGAACCATCGGCTTATATACCATCA   13740
              *** ********************************************************

PMP1610       TGCTTTTTGGCACCTATAAAAAATATATTATGAAAGAACTCTCATAATGCTTTTAAATTT   13800
PMP1666       TGCTTTTTGGCACCTATAAAAAATATATTATGAAAGAACTCTCATAATGCTTTTAAATTT   13800
PMP1665       TGCTTTTTGGCACCTATAAAAAATATATTATGAAAGAACTCTCATAATGCTTTTAAATTT   13800
PMP1667       TGCTTTTTGGCACCTATAAAAAATATATTATGAAAGAACTCTCATAATGCTTTTAAATTT   13800
PMP1663       TGCTTTTTGGCACCTATAAAAAATATATTATGAAAGAACTCTCATAATGCTTTTAAATTT   13800
PMP1664       TGCTTTTTGGCACCTATAAAAAATATATTATGAAAGAACTCTCATAATGCTTTTAAATTT   13800
PMP1662       TGCTTTTTGGCACCTATAAAAAATATATTATGAAAGAACTCTCATAATGCTTTTAAATTT   13800
PMP1611       TGCTTTTTGGCACCTATAAAAAATATATTATGAAAGAACTCTCATAATGCTTTTAAATTT   13800
              ************************************************************

PMP1610       CTTATTCATATTTATTTTTCTATTAATTATCATTACATTTATATTATTTGAGGGAGATTT   13860
PMP1666       CTTATTCATATTTATTTTTCTATTAATTATCATTACATTTATATTATTTGAGGGAGATTT   13860
PMP1665       CTTATTCATATTTATTTTTCTATTAATTATCATTACATTTATATTATTTGAGGGAGATTT   13860
PMP1667       CTTATTCATATTTATTTTTCTATTAATTATCATTACATTTATATTATTTGAGGGAGATTT   13860
```

```
PMP1663      CTTATTCATATTTATTTTTCTATTAATTATCATTACATTTATATTATTTGAGGGAGATTT   13860
PMP1664      CTTATTCATATTTATTTTTCTATTAATTATCATTACATTTATATTATTTGAGGGAGATTT   13860
PMP1662      CTTATTCATATTTATTTTTCTATTAATTATCATTACATTTATATTATTTGAGGGAGATTT   13860
PMP1611      CTTATTCATATTTATTTTTCTATTAATTATCATTACATTTATATTATTTGAGGGAGATTT   13860
             ************************************************************

PMP1610      GTTTCAACCCGCAGTAATTTTAACACTTGCTTATTTTATTTCGATTGCAAGTGCTCTAGT   13920
PMP1666      GTTTCAACCCGCAGTAATTTTAACACTTGCTTATTTTATTTCGATTGCAAGTGCTCTAGT   13920
PMP1665      GTTTCAACCCGCAGTAATTTTAACACTTGCTTATTTTATTTCGATTGCAAGTGCTCTAGT   13920
PMP1667      GTTTCAACCCGCAGTAATTTTAACACTTGCTTATTTTATTTCGATTGCAAGTGCTCTAGT   13920
PMP1663      GTTTCAACCCGCAGTAATTTTAACACTTGCTTATTTTATTTCGATTGCAAGTGCTCTAGT   13920
PMP1664      GTTTCAACCCGCAGTAATTTTAACACTTGCTTATTTTATTTCGATTGCAAGTGCTCTAGT   13920
PMP1662      GTTTCAACCCGCAGTAATTTTAACACTTGCTTATTTTATTTCGATTGCAAGTGCTCTAGT   13920
PMP1611      GTTTCAACCCGCAGTAATTTTAACACTTGCTTATTTTATTTCGATTGCAAGTGCTCTAGT   13920
             ************************************************************

PMP1610      TAATAGAAATGTTTGGGGAACAGAACTCCATTTCAAAACCTTTGGTTTGATATTGCTAGG   13980
PMP1666      TAATAGAAATGTTTGGGGAACAGAACTCCATTTCAAAACCTTTGGTTTGATATTGCTAGG   13980
PMP1665      TAATAGAAATGTTTGGGGAACAGAACTCCATTTCAAAACCTTTGGTTTGATATTGCTAGG   13980
PMP1667      TAATAGAAATGTTTGGGGAACAGAACTCCATTTCAAAACCTTTGGTTTGATATTGCTAGG   13980
PMP1663      TAATAGAAATGTTTGGGGAACAGAACTCCATTTCAAAACCTTTGGTTTGATATTGCTAGG   13980
PMP1664      TAATAGAAATGTTTGGGGAACAGAACTCCATTTCAAAACCTTTGGTTTGATATTGCTAGG   13980
PMP1662      TAATAGAAATGTTTGGGGAACAGAACTCCATTTCAAAACCTTTGGTTTGATATTGCTAGG   13980
PMP1611      TAATAGAAATGTTTGGGGAACAGAACTCCATTTCAAAACCTTTGGTTTGATATTGCTAGG   13980
             ************************************************************

PMP1610      GGTTGCTACATTTATTATAGTTTCCTTGTTGACAAAATTGTCGTACAAACCTAAAGTGGA   14040
PMP1666      GGTTGCTACATTTATTATAGTTTCCTTGTTGACAAAATTGTCGTACAAACCTAAAGTGGA   14040
PMP1665      GGTTGCTACATTTATTATAGTTTCCTTGTTGACAAAATTGTCGTACAAACCTAAAGTGGA   14040
PMP1667      GGTTGCTACATTTATTATAGTTTCCTTGTTGACAAAATTGTCGTACAAACCTAAAGTGGA   14040
PMP1663      GGTTGCTACATTTATTATAGTTTCCTTGTTGACAAAATTGTCGTACAAACCTAAAGTGGA   14040
PMP1664      GGTTGCTACATTTATTATAGTTTCCTTGTTGACAAAATTGTCGTACAAACCTAAAGTGGA   14040
PMP1662      GGTTGCTACATTTATTATAGTTTCCTTGTTGACAAAATTGTCGTACAAACCTAAAGTGGA   14040
PMP1611      GGTTGCTACATTTATTATAGTTTCCTTGTTGACAAAATTGTCGTACAAACCTAAAGTGGA   14040
             ************************************************************
```

```
PMP1610      GGGAATTTCGTATAAAGAATTAAAAGAAATAAATCCTTCAAAGATAATATATGGCATTCT   14100
PMP1666      GGGAATTTCGTATAAAGAATTAAAAGAAATAAATCCTTCAAAGATAATATATGGCATTCT   14100
PMP1665      GGGAATTTCGTATAAAGAATTAAAAGAAATAAATCCTTCAAAGATAATATATGGCATTCT   14100
PMP1667      GGGAATTTCGTATAAAGAATTAAAAGAAATAAATCCTTCAAAGATAATATATGGCATTCT   14100
PMP1663      GGGAATTTCGTATAAAGAATTAAAAGAAATAAATCCTTCAAAGATAATATATGGCATTCT   14100
PMP1664      GGGAATTTCGTATAAAGAATTAAAAGAAATAAATCCTTCAAAGATAATATATGGCATTCT   14100
PMP1662      GGGAATTTCGTATAAAGAATTAAAAGAAATAAATCCTTCAAAGATAATATATGGCATTCT   14100
PMP1611      GGGAATTTCGTATAAAGAATTAAAAGAAATAAATCCTTCAAAGATAATATATGGCATTCT   14100
             ************************************************************

PMP1610      TCTGATTCTAAATCTTGTTATGCTATTTCTTTATATCCATGAAATTCAGAAAGTGGTACT   14160
PMP1666      TCTGATTCTAAATCTTGTTATGCTATTTCTTTATATCCATGAAATTCAGAAAGTGGTACT   14160
PMP1665      TCTGATTCTAAATCTTGTTATGCTATTTCTTTATATCCATGAAATTCAGAAAGTGGTACT   14160
PMP1667      TCTGATTCTAAATCTTGTTATGCTATTTCTTTATATCCATGAAATTCAGAAAGTGGTACT   14160
PMP1663      TCTGATTCTAAATCTTGTTATGCTATTTCTTTATATCCATGAAATTCAGAAAGTGGTACT   14160
PMP1664      TCTGATTCTAAATCTTGTTATGCTATTTCTTTATATCCATGAAATTCAGAAAGTGGTACT   14160
PMP1662      TCTGATTCTAAATCTTGTTATGCTATTTCTTTATATCCATGAAATTCAGAAAGTGGTACT   14160
PMP1611      TCTGATTCTAAATCTTGTTATGCTATTTCTTTATATCCATGAAATTCAGAAAGTGGTACT   14160
             ************************************************************

PMP1610      GTTTTCAGGTAGAGGTTTTTCTAATATTACAGATTTGATAAGTAACTATAGGTACCTATC   14220
PMP1666      GTTTTCAGGTAGAGGTTTTTCTAATATTACAGATTTGATAAGTAACTATAGGTACCTATC   14220
PMP1665      GTTTTCAGGTAGAGGTTTTTCTAATATTACAGATTTGATAAGTAACTATAGGTACCTATC   14220
PMP1667      GTTTTCAGGTAGAGGTTTTTCTAATATTACAGATTTGATAAGTAACTATAGGTACCTATC   14220
PMP1663      GTTTTCAGGTAGAGGTTTTTCTAATATTACAGATTTGATAAGTAACTATAGGTACCTATC   14220
PMP1664      GTTTTCAGGTAGAGGTTTTTCTAATATTACAGATTTGATAAGTAACTATAGGTACCTATC   14220
PMP1662      GTTTTCAGGTAGAGGTTTTTCTAATATTACAGATTTGATAAGTAACTATAGGTACCTATC   14220
PMP1611      GTTTTCAGGTAGAGGTTTTTCTAATATTACAGATTTGATAAGTAACTATAGGTACCTATC   14220
             ************************************************************

PMP1610      TTATTATTCAAATGAAGTAGAAGATCGTGTAAGTGGAATGATTAATCAACTAGCTAAAAT   14280
PMP1666      TTATTATTCAAATGAAGTAGAAGATCGTGTAAGTGGAATGATTAATCAACTAGCTAAAAT   14280
PMP1665      TTATTATTCAAATGAAGTAGAAGATCGTGTAAGTGGAATGATTAATCAACTAGCTAAAAT   14280
PMP1667      TTATTATTCAAATGAAGTAGAAGATCGTGTAAGTGGAATGATTAATCAACTAGCTAAAAT   14280
PMP1663      TTATTATTCAAATGAAGTAGAAGATCGTGTAAGTGGAATGATTAATCAACTAGCTAAAAT   14280
PMP1664      TTATTATTCAAATGAAGTAGAAGATCGTGTAAGTGGAATGATTAATCAACTAGCTAAAAT   14280
```

```
PMP1662      TTATTATTCAAATGAAGTAGAAGATCGTGTAAGTGGAATGATTAATCAACTAGCTAAAAT   14280
PMP1611      TTATTATTCAAATGAAGTAGAAGATCGTGTAAGTGGAATGATTAATCAACTAGCTAAAAT   14280
             ************************************************************

PMP1610      TATTCCAGCGACTACATTTGTTTCTTTATATATATTTATAAATAATTATTTTATAACGAA   14340
PMP1666      TATTCCAGCGACTACATTTGTTTCTTTATATATATTTATAAATAATTATTTTATAACGAA   14340
PMP1665      TATTCCAGCGACTACATTTGTTTCTTTATATATATTTATAAATAATTATTTTATAACGAA   14340
PMP1667      TATTCCAGCGACTACATTTGTTTCTTTATATATATTTATAAATAATTATTTTATAACGAA   14340
PMP1663      TATTCCAGCGACTACATTTGTTTCTTTATATATATTTATAAATAATTATTTTATAACGAA   14340
PMP1664      TATTCCAGCGACTACATTTGTTTCTTTATATATATTTATAAATAATTATTTTATAACGAA   14340
PMP1662      TATTCCAGCGACTACATTTGTTTCTTTATATATATTTATAAATAATTATTTTATAACGAA   14340
PMP1611      TATTCCAGCGACTACATTTGTTTCTTTATATATATTTATAAATAATTATTTTATAACGAA   14340
             ************************************************************

PMP1610      GCAAATAAAGAAAAATTTCATTTATTTGATTCCAATAGCTATATTCTTTGTCTATGCAAT   14400
PMP1666      GCAAATAAAGAAAAATTTCATTTATTTGATTCCAATAGCTATATTCTTTGTCTATGCAAT   14400
PMP1665      GCAAATAAAGAAAAATTTCATTTATTTGATTCCAATAGCTATATTCTTTGTCTATGCAAT   14400
PMP1667      GCAAATAAAGAAAAATTTCATTTATTTGATTCCAATAGCTATATTCTTTGTCTATGCAAT   14400
PMP1663      GCAAATAAAGAAAAATTTCATTTATTTGATTCCAATAGCTATATTCTTTGTCTATGCAAT   14400
PMP1664      GCAAATAAAGAAAAATTTCATTTATTTGATTCCAATAGCTATATTCTTTGTCTATGCAAT   14400
PMP1662      GCAAATAAAGAAAAATTTCATTTATTTGATTCCAATAGCTATATTCTTTGTCTATGCAAT   14400
PMP1611      GCAAATAAAGAAAAATTTCATTTATTTGATTCCAATAGCTATATTCTTTGTCTATGCAAT   14400
             ************************************************************

PMP1610      CATTAGTGGTGGTAGACTGCCCCTTATAAGGTTAGTTATTGGAACTCTGTTGATATTGTA   14460
PMP1666      CATTAGTGGTGGTAGACTGCCCCTTATAAGGTTAGTTATTGGAACTCTGTTGATATTGTA   14460
PMP1665      CATTAGTGGTGGTAGACTGCCCCTTATAAGGTTAGTTATTGGAACTCTGTTGATATTGTA   14460
PMP1667      CATTAGTGGTGGTAGACTGCCCCTTATAAGGTTAGTTATTGGAACTCTGTTGATATTGTA   14460
PMP1663      CATTAGTGGTGGTAGACTGCCCCTTATAAGGTTAGTTATTGGAACTCTGTTGATATTGTA   14460
PMP1664      CATTAGTGGTGGTAGACTGCCCCTTATAAGGTTAGTTATTGGAACTCTGTTGATATTGTA   14460
PMP1662      CATTAGTGGTGGTAGACTGCCCCTTATAAGGTTAGTTATTGGAACTCTGTTGATATTGTA   14460
PMP1611      CATTAGTGGTGGTAGACTGCCCCTTATAAGGTTAGTTATTGGAACTCTGTTGATATTGTA   14460
             ************************************************************

PMP1610      TATATACTCTGTGTACGGGAGTCATAAATCTCAACTTACCAGAAGTTTTAAAATGATTAC   14520
PMP1666      TATATACTCTGTGTACGGGAGTCATAAATCTCAACTTACCAGAAGTTTTAAAATGATTAC   14520
```

```
PMP1665      TATATACTCTGTGTACGGGAGTCATAAATCTCAACTTACCAGAAGTTTTAAAATGATTAC    14520
PMP1667      TATATACTCTGTGTACGGGAGTCATAAATCTCAACTTACCAGAAGTTTTAAAATGATTAC    14520
PMP1663      TATATACTCTGTGTACGGGAGTCATAAATCTCAACTTACCAGAAGTTTTAAAATGATTAC    14520
PMP1664      TATATACTCTGTGTACGGGAGTCATAAATCTCAACTTACCAGAAGTTTTAAAATGATTAC    14520
PMP1662      TATATACTCTGTGTACGGGAGTCATAAATCTCAACTTACCAGAAGTTTTAAAATGATTAC    14520
PMP1611      TATATACTCTGTGTACGGGAGTCATAAATCTCAACTTACCAGAAGTTTTAAAATGATTAC    14520
             ************************************************************

PMP1610      TCGCTCTCTTTTTGCATTTCTTATGTTGATAGTTTTATTCTTTCTTTTAAAATTTGTATT    14580
PMP1666      TCGCTCTCTTTTTGCATTTCTTATGTTGATAGTTTTATTCTTTCTTTTAAAATTTGTATT    14580
PMP1665      TCGCTCTCTTTTTGCATTTCTTATGTTGATAGTTTTATTCTTTCTTTTAAAATTTGTATT    14580
PMP1667      TCGCTCTCTTTTTGCATTTCTTATGTTGATAGTTTTATTCTTTCTTTTAAAATTTGTATT    14580
PMP1663      TCGCTCTCTTTTTGCATTTCTTATGTTGATAGTTTTATTCTTTCTTTTAAAATTTGTATT    14580
PMP1664      TCGCTCTCTTTTTGCATTTCTTATGTTGATAGTTTTATTCTTTCTTTTAAAATTTGTATT    14580
PMP1662      TCGCTCTCTTTTTGCATTTCTTATGTTGATAGTTTTATTCTTTCTTTTAAAATTTGTATT    14580
PMP1611      TCGCTCTCTTTTTGCATTTCTTATGTTGATAGTTTTATTCTTTCTTTTAAAATTTGTATT    14580
             ************************************************************

PMP1610      AGGGCGTTCTTCTCAGGAAGATTTTATCAGTTACATCACTCGTTATATGGGAGGCTCAAT    14640
PMP1666      AGGGCGTTCTTCTCAGGAAGATTTTATCAGTTACATCACTCGTTATATGGGAGGCTCAAT    14640
PMP1665      AGGGCGTTCTTCTCAGGAAGATTTTATCAGTTACATCACTCGTTATATGGGAGGCTCAAT    14640
PMP1667      AGGGCGTTCTTCTCAGGAAGATTTTATCAGTTACATCACTCGTTATATGGGAGGCTCAAT    14640
PMP1663      AGGGCGTTCTTCTCAGGAAGATTTTATCAGTTACATCACTCGTTATATGGGAGGCTCAAT    14640
PMP1664      AGGGCGTTCTTCTCAGGAAGATTTTATCAGTTACATCACTCGTTATATGGGAGGCTCAAT    14640
PMP1662      AGGGCGTTCTTCTCAGGAAGATTTTATCAGTTACATCACTCGTTATATGGGAGGCTCAAT    14640
PMP1611      AGGGCGTTCTTCTCAGGAAGATTTTATCAGTTACATCACTCGTTATATGGGAGGCTCAAT    14640
             ************************************************************

PMP1610      TCAACTATTTGATTTATTTGTTATAGATCCGATACGACGTAACAAAGAACTAGGTGCAGA    14700
PMP1666      TCAACTATTTGATTTATTTGTTATAGATCCGATACGACGTAACAAAGAACTAGGTGCAGA    14700
PMP1665      TCAACTATTTGATTTATTTGTTATAGATCCGATACGACGTAACAAAGAACTAGGTGCAGA    14700
PMP1667      TCAACTATTTGATTTATTTGTTATAGATCCGATACGACGTAACAAAGAACTAGGTGCAGA    14700
PMP1663      TCAACTATTTGATTTATTTGTTATAGATCCGATACGACGTAACAAAGAACTAGGTGCAGA    14700
PMP1664      TCAACTATTTGATTTATTTGTTATAGATCCGATACGACGTAACAAAGAACTAGGTGCAGA    14700
PMP1662      TCAACTATTTGATTTATTTGTTATAGATCCGATACGACGTAACAAAGAACTAGGTGCAGA    14700
PMP1611      TCAACTATTTGATTTATTTGTTATAGATCCGATACGACGTAACAAAGAACTAGGTGCAGA    14700
```

```
            ************************************************************

PMP1610     AACTTTTTCGGGAATTTATGAGATGCTTGCAAAATTAGGATTTGATAATAATATTATAAA   14760
PMP1666     AACTTTTTCGGGAATTTATGAGATGCTTGCAAAATTAGGATTTGATAATAATATTATAAA   14760
PMP1665     AACTTTTTCGGGAATTTATGAGATGCTTGCAAAATTAGGATTTGATAATAATATTATAAA   14760
PMP1667     AACTTTTTCGGGAATTTATGAGATGCTTGCAAAATTAGGATTTGATAATAATATTATAAA   14760
PMP1663     AACTTTTTCGGGAATTTATGAGATGCTTGCAAAATTAGGATTTGATAATAATATTATAAA   14760
PMP1664     AACTTTTTCGGGAATTTATGAGATGCTTGCAAAATTAGGATTTGATAATAATATTATAAA   14760
PMP1662     AACTTTTTCGGGAATTTATGAGATGCTTGCAAAATTAGGATTTGATAATAATATTATAAA   14760
PMP1611     AACTTTTTCGGGAATTTATGAGATGCTTGCAAAATTAGGATTTGATAATAATATTATAAA   14760
            ************************************************************

PMP1610     AGGCTTAGAATGGAGAATATCGCCTAATTATTATTCTTTAGGGAATGTGTATACTGCAAT   14820
PMP1666     AGGCTTAGAATGGAGAATATCGCCTAATTATTATTCTTTAGGGAATGTGTATACTGCAAT   14820
PMP1665     AGGCTTAGAATGGAGAATATCGCCTAATTATTATTCTTTAGGGAATGTGTATACTGCAAT   14820
PMP1667     AGGCTTAGAATGGAGAATATCGCCTAATTATTATTCTTTAGGGAATGTGTATACTGCAAT   14820
PMP1663     AGGCTTAGAATGGAGAATATCGCCTAATTATTATTCTTTAGGGAATGTGTATACTGCAAT   14820
PMP1664     AGGCTTAGAATGGAGAATATCGCCTAATTATTATTCTTTAGGGAATGTGTATACTGCAAT   14820
PMP1662     AGGCTTAGAATGGAGAATATCGCCTAATTATTATTCTTTAGGGAATGTGTATACTGCAAT   14820
PMP1611     AGGCTTAGAATGGAGAATATCGCCTAATTATTATTCTTTAGGGAATGTGTATACTGCAAT   14820
            ************************************************************

PMP1610     TAGACGCTATTATTCAGACTTTGGTGTAATTGGTATTGTAATTTGTCAGAGTTTTACAGC   14880
PMP1666     TAGACGCTATTATTCAGACTTTGGTGTAATTGGTATTGTAATTTGTCAGAGTTTTACAGC   14880
PMP1665     TAGACGCTATTATTCAGACTTTGGTGTAATTGGTATTGTAATTTGTCAGAGTTTTACAGC   14880
PMP1667     TAGACGCTATTATTCAGACTTTGGTGTAATTGGTATTGTAATTTGTCAGAGTTTTACAGC   14880
PMP1663     TAGACGCTATTATTCAGACTTTGGTGTAATTGGTATTGTAATTTGTCAGAGTTTTACAGC   14880
PMP1664     TAGACGCTATTATTCAGACTTTGGTGTAATTGGTATTGTAATTTGTCAGAGTTTTACAGC   14880
PMP1662     TAGACGCTATTATTCAGACTTTGGTGTAATTGGTATTGTAATTTGTCAGAGTTTTACAGC   14880
PMP1611     TAGACGCTATTATTCAGACTTTGGTGTAATTGGTATTGTAATTTGTCAGAGTTTTACAGC   14880
            ************************************************************

PMP1610     ATGGTTGTATACTTTAGGTTATGAAAAAATTAGACATCATTCTTTAGTTACAAATGGTCA   14940
PMP1666     ATGGTTGTATACTTTAGGTTATGAAAAAATTAGACATCATTCTTTAGTTACAAATGGTCA   14940
PMP1665     ATGGTTGTATACTTTAGGTTATGAAAAAATTAGACATCATTCTTTAGTTACAAATGGTCA   14940
PMP1667     ATGGTTGTATACTTTAGGTTATGAAAAAATTAGACATCATTCTTTAGTTACAAATGGTCA   14940
```

```
PMP1663     ATGGTTGTATACTTTAGGTTATGAAAAAATTAGACATCATTCTTTAGTTACAAATGGTCA   14940
PMP1664     ATGGTTGTATACTTTAGGTTATGAAAAAATTAGACATCATTCTTTAGTTACAAATGGTCA   14940
PMP1662     ATGGTTGTATACTTTAGGTTATGAAAAAATTAGACATCATTCTTTAGTTACAAATGGTCA   14940
PMP1611     ATGGTTGTATACTTTAGGTTATGAAAAAATTAGACATCATTCTTTAGTTACAAATGGTCA   14940
            ************************************************************

PMP1610     AAGATTTAGGTTGATTCTATTAGCAGCTTCATTTTATCCATTATTTTTAAATAGTATCGA   15000
PMP1666     AAGATTTAGGTTGATTCTATTAGCAGCTTCATTTTATCCATTATTTTTAAATAGTATCGA   15000
PMP1665     AAGATTTAGGTTGATTCTATTAGCAGCTTCATTTTATCCATTATTTTTAAATAGTATCGA   15000
PMP1667     AAGATTTAGGTTGATTCTATTAGCAGCTTCATTTTATCCATTATTTTTAAATAGTATCGA   15000
PMP1663     AAGATTTAGGTTGATTCTATTAGCAGCTTCATTTTATCCATTATTTTTAAATAGTATCGA   15000
PMP1664     AAGATTTAGGTTGATTCTATTAGCAGCTTCATTTTATCCATTATTTTTAAATAGTATCGA   15000
PMP1662     AAGATTTAGGTTGATTCTATTAGCAGCTTCATTTTATCCATTATTTTTAAATAGTATCGA   15000
PMP1611     AAGATTTAGGTTGATTCTATTAGCAGCTTCATTTTATCCATTATTTTTAAATAGTATCGA   15000
            ************************************************************

PMP1610     GGATGTGTTTTATATTTCAATGGTTACCATTGGATATGGAATCCAAATTGTTATCTTTTA   15060
PMP1666     GGATGTGTTTTATATTTCAATGGTTACCATTGGATATGGAATCCAAATTGTTATCTTTTA   15060
PMP1665     GGATGTGTTTTATATTTCAATGGTTACCATTGGATATGGAATCCAAATTGTTATCTTTTA   15060
PMP1667     GGATGTGTTTTATATTTCAATGGTTACCATTGGATATGGAATCCAAATTGTTATCTTTTA   15060
PMP1663     GGATGTGTTTTATATTTCAATGGTTACCATTGGATATGGAATCCAAATTGTTATCTTTTA   15060
PMP1664     GGATGTGTTTTATATTTCAATGGTTACCATTGGATATGGAATCCAAATTGTTATCTTTTA   15060
PMP1662     GGATGTGTTTTATATTTCAATGGTTACCATTGGATATGGAATCCAAATTGTTATCTTTTA   15060
PMP1611     GGATGTGTTTTATATTTCAATGGTTACCATTGGATATGGAATCCAAATTGTTATCTTTTA   15060
            ************************************************************

PMP1610     TCTGGTTTTTTGGGTTCTTCTGAAAGTTCAGGTTGACTTTAACAAAGGTAAATTAAGGAT   15120
PMP1666     TCTGGTTTTTTGGGTTCTTCTGAAAGTTCAGGTTGACTTTAACAAAGGTAAATTAAGGAT   15120
PMP1665     TCTGGTTTTTTGGGTTCTTCTGAAAGTTCAGGTTGACTTTAACAAAGGTAAATTAAGGAT   15120
PMP1667     TCTGGTTTTTTGGGTTCTTCTGAAAGTTCAGGTTGACTTTAACAAAGGTAAATTAAGGAT   15120
PMP1663     TCTGGTTTTTTGGGTTCTTCTGAAAGTTCAGGTTGACTTTAACAAAGGTAAATTAAGGAT   15120
PMP1664     TCTGGTTTTTTGGGTTCTTCTGAAAGTTCAGGTTGACTTTAACAAAGGTAAATTAAGGAT   15120
PMP1662     TCTGGTTTTTTGGGTTCTTCTGAAAGTTCAGGTTGACTTTAACAAAGGTAAATTAAGGAT   15120
PMP1611     TCTGGTTTTTTGGGTTCTTCTGAAAGTTCAGGTTGACTTTAACAAAGGTAAATTAAGGAT   15120
            ************************************************************
```

```
PMP1610        AAATAGATGAATTTAGCGCTAATGTATTGAGTTATATAAAGGGATAGATTTGGTAGTATT    15180
PMP1666        AAATAGATGAATTTAGCGCTAATGTATTGAGTTATATAAAGGGATAGATTTGGTAGTATT    15180
PMP1665        AAATAGATGAATTTAGCGCTAATGTATTGAGTTATATAAAGGGATAGATTTGGTAGTATT    15180
PMP1667        AAATAGATGAATTTAGCGCTAATGTATTGAGTTATATAAAGGGATAGATTTGGTAGTATT    15180
PMP1663        AAATAGATGAATTTAGCGCTAATGTATTGAGTTATATAAAGGGATAGATTTGGTAGTATT    15180
PMP1664        AAATAGATGAATTTAGCGCTAATGTATTGAGTTATATAAAGGGATAGATTTGGTAGTATT    15180
PMP1662        AAATAGATGAATTTAGCGCTAATGTATTGAGTTATATAAAGGGATAGATTTGGTAGTATT    15180
PMP1611        AAATAGATGAATTTAGCGCTAATGTATTGAGTTATATAAAGGGATAGATTTGGTAGTATT    15180
               ************************************************************

PMP1610        TGTAGGAATTTTAATTGGAGGAAGAGAGCCTTGAATGAGAATTACAAATCTTCTGAAACA    15240
PMP1666        TGTAGGAATTTTAATTGGAGGAAGAGAGCCTTGAATGAGAATTACAAATCTTCTGAAACA    15240
PMP1665        TGTAGGAATTTTAATTGGAGGAAGAGAGCCTTGAATGAGAATTACAAATCTTCTGAAACA    15240
PMP1667        TGTAGGAATTTTAATTGGAGGAAGAGAGCCTTGAATGAGAATTACAAATCTTCTGAAACA    15240
PMP1663        TGTAGGAATTTTAATTGGAGGAAGAGAGCCTTGAATGAGAATTACAAATCTTCTGAAACA    15240
PMP1664        TGTAGGAATTTTAATTGGAGGAAGAGAGCCTTGAATGAGAATTACAAATCTTCTGAAACA    15240
PMP1662        TGTAGGAATTTTAATTGGAGGAAGAGAGCCTTGAATGAGAATTACAAATCTTCTGAAACA    15240
PMP1611        TGTAGGAATTTTAATTGGAGGAAGAGAGCCTTGAATGAGAATTACAAATCTTCTGAAACA    15240
               ************************************************************

PMP1610        ATTTTTAGGTGGGGAGTATAGTTATGAAATTGAAGTTTCTTATAACAAATTTGTTTCATG    15300
PMP1666        ATTTTTAGGTGGGGAGTATAGTTATGAAATTGAAGTTTCTTATAACAAATTTGTTTCATG    15300
PMP1665        ATTTTTAGGTGGGGAGTATAGTTATGAAATTGAAGTTTCTTATAACAAATTTGTTTCATG    15300
PMP1667        ATTTTTAGGTGGGGAGTATAGTTATGAAATTGAAGTTTCTTATAACAAATTTGTTTCATG    15300
PMP1663        ATTTTTAGGTGGGGAGTATAGTTATGAAATTGAAGTTTCTTATAACAAATTTGTTTCATG    15300
PMP1664        ATTTTTAGGTGGGGAGTATAGTTATGAAATTGAAGTTTCTTATAACAAATTTGTTTCATG    15300
PMP1662        ATTTTTAGGTGGGGAGTATAGTTATGAAATTGAAGTTTCTTATAACAAATTTGTTTCATG    15300
PMP1611        ATTTTTAGGTGGGGAGTATAGTTATGAAATTGAAGTTTCTTATAACAAATTTGTTTCATG    15300
               ************************************************************

PMP1610        TTCTTTTGTCTAATCTGATTACAATTCTTACATCAGTTATAGTTGTACTAATTTTACCAA    15360
PMP1666        TTCTTTTGTCTAATCTGATTACAATTCTTACATCAGTTATAGTTGTACTAATTTTACCAA    15360
PMP1665        TTCTTTTGTCTAATCTGATTACAATTCTTACATCAGTTATAGTTGTACTAATTTTACCAA    15360
PMP1667        TTCTTTTGTCTAATCTGATTACAATTCTTACATCAGTTATAGTTGTACTAATTTTACCAA    15360
PMP1663        TTCTTTTGTCTAATCTGATTACAATTCTTACATCAGTTATAGTTGTACTAATTTTACCAA    15360
PMP1664        TTCTTTTGTCTAATCTGATTACAATTCTTACATCAGTTATAGTTGTACTAATTTTACCAA    15360
```

```
PMP1662      TTCTTTTGTCTAATCTGATTACAATTCTTACATCAGTTATAGTTGTACTAATTTTACCAA   15360
PMP1611      TTCTTTTGTCTAATCTGATTACAATTCTTACATCAGTTATAGTTGTACTAATTTTACCAA   15360
             ************************************************************

PMP1610      AAATTATGGGAGTAACTGAGTATAGTTATTGGCAACTATATATTTTTTACCTAACATATA   15420
PMP1666      AAATTATGGGAGTAACTGAGTATAGTTATTGGCAACTATATATTTTTTACCTAACATATA   15420
PMP1665      AAATTATGGGAGTAACTGAGTATAGTTATTGGCAACTATATATTTTTTACCTAACATATA   15420
PMP1667      AAATTATGGGAGTAACTGAGTATAGTTATTGGCAACTATATATTTTTTACCTAACATATA   15420
PMP1663      AAATTATGGGAGTAACTGAGTATAGTTATTGGCAACTATATATTTTTTACCTAACATATA   15420
PMP1664      AAATTATGGGAGTAACTGAGTATAGTTATTGGCAACTATATATTTTTTACCTAACATATA   15420
PMP1662      AAATTATGGGAGTAACTGAGTATAGTTATTGGCAACTATATATTTTTTACCTAACATATA   15420
PMP1611      AAATTATGGGAGTAACTGAGTATAGTTATTGGCAACTATATATTTTTTACCTAACATATA   15420
             ************************************************************

PMP1610      TTGGTTTTTTTCATCTGGGATGGATTGATGGAATTTATCTTAAATATGGCGGATTAGAGT   15480
PMP1666      TTGGTTTTTTTCATCTGGGATGGATTGATGGAATTTATCTTAAATATGGCGGATTAGAGT   15480
PMP1665      TTGGTTTTTTTCATCTGGGATGGATTGATGGAATTTATCTTAAATATGGCGGATTAGAGT   15480
PMP1667      TTGGTTTTTTTCATCTGGGATGGATTGATGGAATTTATCTTAAATATGGCGGATTAGAGT   15480
PMP1663      TTGGTTTTTTTCATCTGGGATGGATTGATGGAATTTATCTTAAATATGGCGGATTAGAGT   15480
PMP1664      TTGGTTTTTTTCATCTGGGATGGATTGATGGAATTTATCTTAAATATGGCGGATTAGAGT   15480
PMP1662      TTGGTTTTTTTCATCTGGGATGGATTGATGGAATTTATCTTAAATATGGCGGATTAGAGT   15480
PMP1611      TTGGTTTTTTTCATCTGGGATGGATTGATGGAATTTATCTTAAATATGGCGGATTAGAGT   15480
             ************************************************************

PMP1610      ACCAGAACTTAGATAAGAAACAGTTTTATTCTCAAATACTTCAATTTTCCAGTTTTTTAA   15540
PMP1666      ACCAGAACTTAGATAAGAAACAGTTTTATTCTCAAATACTTCAATTTTCCAGTTTTTTAA   15540
PMP1665      ACCAGAACTTAGATAAGAAACAGTTTTATTCTCAAATACTTCAATTTTCCAGTTTTTTAA   15540
PMP1667      ACCAGAACTTAGATAAGAAACAGTTTTATTCTCAAATACTTCAATTTTCCAGTTTTTTAA   15540
PMP1663      ACCAGAACTTAGATAAGAAACAGTTTTATTCTCAAATACTTCAATTTTCCAGTTTTTTAA   15540
PMP1664      ACCAGAACTTAGATAAGAAACAGTTTTATTCTCAAATACTTCAATTTTCCAGTTTTTTAA   15540
PMP1662      ACCAGAACTTAGATAAGAAACAGTTTTATTCTCAAATACTTCAATTTTCCAGTTTTTTAA   15540
PMP1611      ACCAGAACTTAGATAAGAAACAGTTTTATTCTCAAATACTTCAATTTTCCAGTTTTTTAA   15540
             ************************************************************

PMP1610      TTTTAATTTCTTTTCTATTATTTGGTTTTAACTTATTGACTGTGACAGATCAAAATGCAA   15600
PMP1666      TTTTAATTTCTTTTCTATTATTTGGTTTTAACTTATTGACTGTGACAGATCAAAATGCAA   15600
```

```
PMP1665      TTTTAATTTCTTTTCTATTATTTGGTTTTAACTTATTGACTGTGACAGATCAAAATGCAA   15600
PMP1667      TTTTAATTTCTTTTCTATTATTTGGTTTTAACTTATTGACTGTGACAGATCAAAATGCAA   15600
PMP1663      TTTTAATTTCTTTTCTATTATTTGGTTTTAACTTATTGACTGTGACAGATCAAAATGCAA   15600
PMP1664      TTTTAATTTCTTTTCTATTATTTGGTTTTAACTTATTGACTGTGACAGATCAAAATGCAA   15600
PMP1662      TTTTAATTTCTTTTCTATTATTTGGTTTTAACTTATTGACTGTGACAGATCAAAATGCAA   15600
PMP1611      TTTTAATTTCTTTTCTATTATTTGGTTTTAACTTATTGACTGTGACAGATCAAAATGCAA   15600
             ************************************************************

PMP1610      AATATATTTATAACATGACTATTATTAGTATGATAGTTACAAATTTAAGAATGTTATTCG   15660
PMP1666      AATATATTTATAACATGACTATTATTAGTATGATAGTTACAAATTTAAGAATGTTATTCG   15660
PMP1665      AATATATTTATAACATGACTATTATTAGTATGATAGTTACAAATTTAAGAATGTTATTCG   15660
PMP1667      AATATATTTATAACATGACTATTATTAGTATGATAGTTACAAATTTAAGAATGTTATTCG   15660
PMP1663      AATATATTTATAACATGACTATTATTAGTATGATAGTTACAAATTTAAGAATGTTATTCG   15660
PMP1664      AATATATTTATAACATGACTATTATTAGTATGATAGTTACAAATTTAAGAATGTTATTCG   15660
PMP1662      AATATATTTATAACATGACTATTATTAGTATGATAGTTACAAATTTAAGAATGTTATTCG   15660
PMP1611      AATATATTTATAACATGACTATTATTAGTATGATAGTTACAAATTTAAGAATGTTATTCG   15660
             ************************************************************

PMP1610      TTTATATTTTGCAGATGACAAATCGATTAAAGGATAGTTCCATCATTCTAATCAGTGATC   15720
PMP1666      TTTATATTTTGCAGATGACAAATCGATTAAAGGATAGTTCCATCATTCTAATCAGTGATC   15720
PMP1665      TTTATATTTTGCAGATGACAAATCGATTAAAGGATAGTTCCATCATTCTAATCAGTGATC   15720
PMP1667      TTTATATTTTGCAGATGACAAATCGATTAAAGGATAGTTCCATCATTCTAATCAGTGATC   15720
PMP1663      TTTATATTTTGCAGATGACAAATCGATTAAAGGATAGTTCCATCATTCTAATCAGTGATC   15720
PMP1664      TTTATATTTTGCAGATGACAAATCGATTAAAGGATAGTTCCATCATTCTAATCAGTGATC   15720
PMP1662      TTTATATTTTGCAGATGACAAATCGATTAAAGGATAGTTCCATCATTCTAATCAGTGATC   15720
PMP1611      TTTATATTTTGCAGATGACAAATCGATTAAAGGATAGTTCCATCATTCTAATCAGTGATC   15720
             ************************************************************

PMP1610      GCGTTATATATGTTATTCTTTTATTCCTGTTTATTATATTTAAATGGCATGAATACAAGG   15780
PMP1666      GCGTTATATATGTTATTCTTTTATTCCTGTTTATTATATTTAAATGGCATGAATACAAGG   15780
PMP1665      GCGTTATATATGTTATTCTTTTATTCCTGTTTATTATATTTAAATGGCATGAATACAAGG   15780
PMP1667      GCGTTATATATGTTATTCTTTTATTCCTGTTTATTATATTTAAATGGCATGAATACAAGG   15780
PMP1663      GCGTTATATATGTTATTCTTTTATTCCTGTTTATTATATTTAAATGGCATGAATACAAGG   15780
PMP1664      GCGTTATATATGTTATTCTTTTATTCCTGTTTATTATATTTAAATGGCATGAATACAAGG   15780
PMP1662      GCGTTATATATGTTATTCTTTTATTCCTGTTTATTATATTTAAATGGCATGAATACAAGG   15780
PMP1611      GCGTTATATATGTTATTCTTTTATTCCTGTTTATTATATTTAAATGGCATGAATACAAGG   15780
```

```
                  ************************************************************

PMP1610           TAATGATTTGGGCAGATGTTTTGGGAAGGACATTTTCTCTCCTACTTTCTTTTTGGATTT    15840
PMP1666           TAATGATTTGGGCAGATGTTTTGGGAAGGACATTTTCTCTCCTACTTTCTTTTTGGATTT    15840
PMP1665           TAATGATTTGGGCAGATGTTTTGGGAAGGACATTTTCTCTCCTACTTTCTTTTTGGATTT    15840
PMP1667           TAATGATTTGGGCAGATGTTTTGGGAAGGACATTTTCTCTCCTACTTTCTTTTTGGATTT    15840
PMP1663           TAATGATTTGGGCAGATGTTTTGGGAAGGACATTTTCTCTCCTACTTTCTTTTTGGATTT    15840
PMP1664           TAATGATTTGGGCAGATGTTTTGGGAAGGACATTTTCTCTCCTACTTTCTTTTTGGATTT    15840
PMP1662           TAATGATTTGGGCAGATGTTTTGGGAAGGACATTTTCTCTCCTACTTTCTTTTTGGATTT    15840
PMP1611           TAATGATTTGGGCAGATGTTTTGGGAAGGACATTTTCTCTCCTACTTTCTTTTTGGATTT    15840
                  ************************************************************

PMP1610           GTAAAGATATTGTTTTTCAATCCTTATCCGAGTTCATATTGGATCTGAGAGAGTCTTTTG    15900
PMP1666           GTAAAGATATTGTTTTTCAATCCTTATCCGAGTTCATATTGGATCTGAGAGAGTCTTTTG    15900
PMP1665           GTAAAGATATTGTTTTTCAATCCTTATCCGAGTTCATATTGGATCTGAGAGAGTCTTTTG    15900
PMP1667           GTAAAGATATTGTTTTTCAATCCTTATCCGAGTTCATATTGGATCTGAGAGAGTCTTTTG    15900
PMP1663           GTAAAGATATTGTTTTTCAATCCTTATCCGAGTTCATATTGGATCTGAGAGAGTCTTTTG    15900
PMP1664           GTAAAGATATTGTTTTTCAATCCTTATCCGAGTTCATATTGGATCTGAGAGAGTCTTTTG    15900
PMP1662           GTAAAGATATTGTTTTTCAATCCTTATCCGAGTTCATATTGGATCTGAGAGAGTCTTTTG    15900
PMP1611           GTAAAGATATTGTTTTTCAATCCTTATCCGAGTTCATATTGGATCTGAGAGAGTCTTTTG    15900
                  ************************************************************

PMP1610           ACAATATCCGTGTTGGAATCAATTTAATGTTATCCAATATTGCAAGTAGTATGATTATTG    15960
PMP1666           ACAATATCCGTGTTGGAATCAATTTAATGTTATCCAATATTGCAAGTAGTATGATTATTG    15960
PMP1665           ACAATATCCGTGTTGGAATCAATTTAATGTTATCCAATATTGCAAGTAGTATGATTATTG    15960
PMP1667           ACAATATCCGTGTTGGAATCAATTTAATGTTATCCAATATTGCAAGTAGTATGATTATTG    15960
PMP1663           ACAATATCCGTGTTGGAATCAATTTAATGTTATCCAATATTGCAAGTAGTATGATTATTG    15960
PMP1664           ACAATATCCGTGTTGGAATCAATTTAATGTTATCCAATATTGCAAGTAGTATGATTATTG    15960
PMP1662           ACAATATCCGTGTTGGAATCAATTTAATGTTATCCAATATTGCAAGTAGTATGATTATTG    15960
PMP1611           ACAATATCCGTGTTGGAATCAATTTAATGTTATCCAATATTGCAAGTAGTATGATTATTG    15960
                  ************************************************************

PMP1610           GTATTGTTCGAATGGGAATTCAATGGAATTGGAATATCGAAACATTCGGGAAAGTATCAC    16020
PMP1666           GTATTGTTCGAATGGGAATTCAATGGAATTGGAATATCGAAACATTCGGGAAAGTATCAC    16020
PMP1665           GTATTGTTCGAATGGGAATTCAATGGAATTGGAATATCGAAACATTCGGGAAAGTATCAC    16020
PMP1667           GTATTGTTCGAATGGGAATTCAATGGAATTGGAATATCGAAACATTCGGGAAAGTATCAC    16020
```

```
PMP1663      GTATTGTTCGAATGGGAATTCAATGGAATTGGAATATCGAAACATTCGGGAAAGTATCAC    16020
PMP1664      GTATTGTTCGAATGGGAATTCAATGGAATTGGAATATCGAAACATTCGGGAAAGTATCAC    16020
PMP1662      GTATTGTTCGAATGGGAATTCAATGGAATTGGAATATCGAAACATTCGGGAAAGTATCAC    16020
PMP1611      GTATTGTTCGAATGGGAATTCAATGGAATTGGAATATCGAAACATTCGGGAAAGTATCAC    16020
             ************************************************************

PMP1610      TGACGCTAAGCATCTCTAATTTATTAATGACTTTTATTAATGCGATTGGTTTAGTTGTTT    16080
PMP1666      TGACGCTAAGCATCTCTAATTTATTAATGACTTTTATTAATGCGATTGGTTTAGTTGTTT    16080
PMP1665      TGACGCTAAGCATCTCTAATTTATTAATGACTTTTATTAATGCGATTGGTTTAGTTGTTT    16080
PMP1667      TGACGCTAAGCATCTCTAATTTATTAATGACTTTTATTAATGCGATTGGTTTAGTTGTTT    16080
PMP1663      TGACGCTAAGCATCTCTAATTTATTAATGACTTTTATTAATGCGATTGGTTTAGTTGTTT    16080
PMP1664      TGACGCTAAGCATCTCTAATTTATTAATGACTTTTATTAATGCGATTGGTTTAGTTGTTT    16080
PMP1662      TGACGCTAAGCATCTCTAATTTATTAATGACTTTTATTAATGCGATTGGTTTAGTTGTTT    16080
PMP1611      TGACGCTAAGCATCTCTAATTTATTAATGACTTTTATTAATGCGATTGGTTTAGTTGTTT    16080
             ************************************************************

PMP1610      TTCCTTTGTTAAAACGGACAAAAACGGAAAATTTATCTAAAATTTATTCCAACTTAAGAA    16140
PMP1666      TTCCTTTGTTAAAACGGACAAAAACGGAAAATTTATCTAAAATTTATTCCAACTTAAGAA    16140
PMP1665      TTCCTTTGTTAAAACGGACAAAAACGGAAAATTTATCTAAAATTTATTCCAACTTAAGAA    16140
PMP1667      TTCCTTTGTTAAAACGGACAAAAACGGAAAATTTATCTAAAATTTATTCCAACTTAAGAA    16140
PMP1663      TTCCTTTGTTAAAACGGACAAAAACGGAAAATTTATCTAAAATTTATTCCAACTTAAGAA    16140
PMP1664      TTCCTTTGTTAAAACGGACAAAAACGGAAAATTTATCTAAAATTTATTCCAACTTAAGAA    16140
PMP1662      TTCCTTTGTTAAAACGGACAAAAACGGAAAATTTATCTAAAATTTATTCCAACTTAAGAA    16140
PMP1611      TTCCTTTGTTAAAACGGACAAAAACGGAAAATTTATCTAAAATTTATTCCAACTTAAGAA    16140
             ************************************************************

PMP1610      ATGTTTTGATGCTTATCATGTTCGCGATTTTGCTCATTTACTATCCTTTAAAAATTGTAT    16200
PMP1666      ATGTTTTGATGCTTATCATGTTCGCGATTTTGCTCATTTACTATCCTTTAAAAATTGTAT    16200
PMP1665      ATGTTTTGATGCTTATCATGTTCGCGATTTTGCTCATTTACTATCCTTTAAAAATTGTAT    16200
PMP1667      ATGTTTTGATGCTTATCATGTTCGCGATTTTGCTCATTTACTATCCTTTAAAAATTGTAT    16200
PMP1663      ATGTTTTGATGCTTATCATGTTCGCGATTTTGCTCATTTACTATCCTTTAAAAATTGTAT    16200
PMP1664      ATGTTTTGATGCTTATCATGTTCGCGATTTTGCTCATTTACTATCCTTTAAAAATTGTAT    16200
PMP1662      ATGTTTTGATGCTTATCATGTTCGCGATTTTGCTCATTTACTATCCTTTAAAAATTGTAT    16200
PMP1611      ATGTTTTGATGCTTATCATGTTCGCGATTTTGCTCATTTACTATCCTTTAAAAATTGTAT    16200
             ************************************************************
```

```
PMP1610      TAGACCTCTGGTTGCCAGCCTATCAAGATGCCTTGATTTTCATGACCCTTATTTTCCCTA   16260
PMP1666      TAGACCTCTGGTTGCCAGCCTATCAAGATGCCTTGATTTTCATGACCCTTATTTTCCCTA   16260
PMP1665      TAGACCTCTGGTTGCCAGCCTATCAAGATGCCTTGATTTTCATGACCCTTATTTTCCCTA   16260
PMP1667      TAGACCTCTGGTTGCCAGCCTATCAAGATGCCTTGATTTTCATGACCCTTATTTTCCCTA   16260
PMP1663      TAGACCTCTGGTTGCCAGCCTATCAAGATGCCTTGATTTTCATGACCCTTATTTTCCCTA   16260
PMP1664      TAGACCTCTGGTTGCCAGCCTATCAAGATGCCTTGATTTTCATGACCCTTATTTTCCCTA   16260
PMP1662      TAGACCTCTGGTTGCCAGCCTATCAAGATGCCTTGATTTTCATGACCCTTATTTTCCCTA   16260
PMP1611      TAGACCTCTGGTTGCCAGCCTATCAAGATGCCTTGATTTTCATGACCCTTATTTTCCCTA   16260
             ************************************************************

PMP1610      TGTCAGTCTATGAAGGAAAAATGGCATTGGTCATTAATACTTACTTAAAGGCATTAAGAA   16320
PMP1666      TGTCAGTCTATGAAGGAAAAATGGCATTGGTCATTAATACTTACTTAAAGGCATTAAGAA   16320
PMP1665      TGTCAGTCTATGAAGGAAAAATGGCATTGGTCATTAATACTTACTTAAAGGCATTAAGAA   16320
PMP1667      TGTCAGTCTATGAAGGAAAAATGGCATTGGTCATTAATACTTACTTAAAGGCATTAAGAA   16320
PMP1663      TGTCAGTCTATGAAGGAAAAATGGCATTGGTCATTAATACTTACTTAAAGGCATTAAGAA   16320
PMP1664      TGTCAGTCTATGAAGGAAAAATGGCATTGGTCATTAATACTTACTTAAAGGCATTAAGAA   16320
PMP1662      TGTCAGTCTATGAAGGAAAAATGGCATTGGTCATTAATACTTACTTAAAGGCATTAAGAA   16320
PMP1611      TGTCAGTCTATGAAGGAAAAATGGCATTGGTCATTAATACTTACTTAAAGGCATTAAGAA   16320
             ************************************************************

PMP1610      TGGAGAGAGATATTCTCAAAATAAATACTTTGATTATGTTGTTCAGTATGTTAGTTACCC   16380
PMP1666      TGGAGAGAGATATTCTCAAAATAAATACTTTGATTATGTTGTTCAGTATGTTAGTTACCC   16380
PMP1665      TGGAGAGAGATATTCTCAAAATAAATACTTTGATTATGTTGTTCAGTATGTTAGTTACCC   16380
PMP1667      TGGAGAGAGATATTCTCAAAATAAATACTTTGATTATGTTGTTCAGTATGTTAGTTACCC   16380
PMP1663      TGGAGAGAGATATTCTCAAAATAAATACTTTGATTATGTTGTTCAGTATGTTAGTTACCC   16380
PMP1664      TGGAGAGAGATATTCTCAAAATAAATACTTTGATTATGTTGTTCAGTATGTTAGTTACCC   16380
PMP1662      TGGAGAGAGATATTCTCAAAATAAATACTTTGATTATGTTGTTCAGTATGTTAGTTACCC   16380
PMP1611      TGGAGAGAGATATTCTCAAAATAAATACTTTGATTATGTTGTTCAGTATGTTAGTTACCC   16380
             ************************************************************

PMP1610      TAATAACTACTCTATTATTAAATAGTTTAGAGCTGACTGTTGTATCGATAGTTGTTTTGC   16440
PMP1666      TAATAACTACTCTATTATTAAATAGTTTAGAGCTGACTGTTGTATCGATAGTTGTTTTGC   16440
PMP1665      TAATAACTACTCTATTATTAAATAGTTTAGAGCTGACTGTTGTATCGATAGTTGTTTTGC   16440
PMP1667      TAATAACTACTCTATTATTAAATAGTTTAGAGCTGACTGTTGTATCGATAGTTGTTTTGC   16440
PMP1663      TAATAACTACTCTATTATTAAATAGTTTAGAGCTGACTGTTGTATCGATAGTTGTTTTGC   16440
PMP1664      TAATAACTACTCTATTATTAAATAGTTTAGAGCTGACTGTTGTATCGATAGTTGTTTTGC   16440
```

```
PMP1662      TAATAACTACTCTATTATTAAATAGTTTAGAGCTGACTGTTGTATCGATAGTTGTTTTGC   16440
PMP1611      TAATAACTACTCTATTATTAAATAGTTTAGAGCTGACTGTTGTATCGATAGTTGTTTTGC   16440
             ************************************************************

PMP1610      TAGCTTTACGTAGCGTAATAGCAGAACTAATTCTATCTAAAAAACTGGATGTTTCGGTTA   16500
PMP1666      TAGCTTTACGTAGCGTAATAGCAGAACTAATTCTATCTAAAAAACTGGATGTTTCGGTTA   16500
PMP1665      TAGCTTTACGTAGCGTAATAGCAGAACTAATTCTATCTAAAAAACTGGATGTTTCGGTTA   16500
PMP1667      TAGCTTTACGTAGCGTAATAGCAGAACTAATTCTATCTAAAAAACTGGATGTTTCGGTTA   16500
PMP1663      TAGCTTTACGTAGCGTAATAGCAGAACTAATTCTATCTAAAAAACTGGATGTTTCGGTTA   16500
PMP1664      TAGCTTTACGTAGCGTAATAGCAGAACTAATTCTATCTAAAAAACTGGATGTTTCGGTTA   16500
PMP1662      TAGCTTTACGTAGCGTAATAGCAGAACTAATTCTATCTAAAAAACTGGATGTTTCGGTTA   16500
PMP1611      TAGCTTTACGTAGCGTAATAGCAGAACTAATTCTATCTAAAAAACTGGATGTTTCGGTTA   16500
             ************************************************************

PMP1610      AGAAAGATATTGTATTAGAATTTCTTTTGACGATTGTCTTTATTTCTTCAAGTTGGTACT   16560
PMP1666      AGAAAGATATTGTATTAGAATTTCTTTTGACGATTGTCTTTATTTCTTCAAGTTGGTACT   16560
PMP1665      AGAAAGATATTGTATTAGAATTTCTTTTGACGATTGTCTTTATTTCTTCAAGTTGGTACT   16560
PMP1667      AGAAAGATATTGTATTAGAATTTCTTTTGACGATTGTCTTTATTTCTTCAAGTTGGTACT   16560
PMP1663      AGAAAGATATTGTATTAGAATTTCTTTTGACGATTGTCTTTATTTCTTCAAGTTGGTACT   16560
PMP1664      AGAAAGATATTGTATTAGAATTTCTTTTGACGATTGTCTTTATTTCTTCAAGTTGGTACT   16560
PMP1662      AGAAAGATATTGTATTAGAATTTCTTTTGACGATTGTCTTTATTTCTTCAAGTTGGTACT   16560
PMP1611      AGAAAGATATTGTATTAGAATTTCTTTTGACGATTGTCTTTATTTCTTCAAGTTGGTACT   16560
             ************************************************************

PMP1610      TACCGATTTGGCCCGCAGTAATAGTTTATTTGTTAGCGTATACTTTATACTTGTATCTAA   16620
PMP1666      TACCGATTTGGCCCGCAGTAATAGTTTATTTGTTAGCGTATACTTTATACTTGTATCTAA   16620
PMP1665      TACCGATTTGGCCCGCAGTAATAGTTTATTTGTTAGCGTATACTTTATACTTGTATCTAA   16620
PMP1667      TACCGATTTGGCCCGCAGTAATAGTTTATTTGTTAGCGTATACTTTATACTTGTATCTAA   16620
PMP1663      TACCGATTTGGCCCGCAGTAATAGTTTATTTGTTAGCGTATACTTTATACTTGTATCTAA   16620
PMP1664      TACCGATTTGGCCCGCAGTAATAGTTTATTTGTTAGCGTATACTTTATACTTGTATCTAA   16620
PMP1662      TACCGATTTGGCCCGCAGTAATAGTTTATTTGTTAGCGTATACTTTATACTTGTATCTAA   16620
PMP1611      TACCGATTTGGCCCGCAGTAATAGTTTATTTGTTAGCGTATACTTTATACTTGTATCTAA   16620
             ************************************************************

PMP1610      AGCGTAAAGATATAAAAATGTATATAGAATATTTTAGAAAGAAAATATTTGAATAAAAAG   16680
PMP1666      AGCGTAAAGATATAAAAATGTATATAGAATATTTTAGAAAGAAAATATTTGAATAAAAAG   16680
```

```
PMP1665      AGCGTAAAGATATAAAAATGTATATAGAATATTTTAGAAAGAAAATATTTGAATAAAAAG   16680
PMP1667      AGCGTAAAGATATAAAAATGTATATAGAATATTTTAGAAAGAAAATATTTGAATAAAAAG   16680
PMP1663      AGCGTAAAGATATAAAAATGTATATAGAATATTTTAGAAAGAAAATATTTGAATAAAAAG   16680
PMP1664      AGCGTAAAGATATAAAAATGTATATAGAATATTTTAGAAAGAAAATATTTGAATAAAAAG   16680
PMP1662      AGCGTAAAGATATAAAAATGTATATAGAATATTTTAGAAAGAAAATATTTGAATAAAAAG   16680
PMP1611      AGCGTAAAGATATAAAAATGTATATAGAATATTTTAGAAAGAAAATATTTGAATAAAAAG   16680
             ************************************************************

PMP1610      AATTATATATCAGTTAGATGGCAAATTCTATTTTTACCTTTTTGTCGTTTAGTAGAAAAT   16740
PMP1666      AATTATATATCAGTTAGATGGCAAATTCTATTTTTACCTTTTTGTCGTTTAGTAGAAAAT   16740
PMP1665      AATTATATATCAGTTAGATGGCAAATTCTATTTTTACCTTTTTGTCGTTTAGTAGAAAAT   16740
PMP1667      AATTATATATCAGTTAGATGGCAAATTCTATTTTTACCTTTTTGTCGTTTAGTAGAAAAT   16740
PMP1663      AATTATATATCAGTTAGATGGCAAATTCTATTTTTACCTTTTTGTCGTTTAGTAGAAAAT   16740
PMP1664      AATTATATATCAGTTAGATGGCAAATTCTATTTTTACCTTTTTGTCGTTTAGTAGAAAAT   16740
PMP1662      AATTATATATCAGTTAGATGGCAAATTCTATTTTTACCTTTTTGTCGTTTAGTAGAAAAT   16740
PMP1611      AATTATATATCAGTTAGATGGCAAATTCTATTTTTACCTTTTTGTCGTTTAGTAGAAAAT   16740
             ************************************************************

PMP1610      GATAAAAAATATGATACTATTTTTTGCACATATCTAGAAGCAATTTAAATGTGTCAGGTG   16800
PMP1666      GATAAAAAATATGATACTATTTTTTGCACATATCTAGAAGCAATTTAAATGTGTCAGGTG   16800
PMP1665      GATAAAAAATATGATACTATTTTTTGCACATATCTAGAAGCAATTTAAATGTGTCAGGTG   16800
PMP1667      GATAAAAAATATGATACTATTTTTTGCACATATCTAGAAGCAATTTAAATGTGTCAGGTG   16800
PMP1663      GATAAAAAATATGATACTATTTTTTGCACATATCTAGAAGCAATTTAAATGTGTCAGGTG   16800
PMP1664      GATAAAAAATATGATACTATTTTTTGCACATATCTAGAAGCAATTTAAATGTGTCAGGTG   16800
PMP1662      GATAAAAAATATGATACTATTTTTTGCACATATCTAGAAGCAATTTAAATGTGTCAGGTG   16800
PMP1611      GATAAAAAATATGATACTATTTTTTGCACATATCTAGAAGCAATTTAAATGTGTCAGGTG   16800
             ************************************************************

PMP1610      ATAAATTAATTTAAACTAAGAATAGTTGCTGGAACATTGCTATTAGTGGGAACAAGTTAC   16860
PMP1666      ATAAATTAATTTAAACTAAGAATAGTTGCTGGAACATTGCTATTAGTGGGAACAAGTTAC   16860
PMP1665      ATAAATTAATTTAAACTAAGAATAGTTGCTGGAACATTGCTATTAGTGGGAACAAGTTAC   16860
PMP1667      ATAAATTAATTTAAACTAAGAATAGTTGCTGGAACATTGCTATTAGTGGGAACAAGTTAC   16860
PMP1663      ATAAATTAATTTAAACTAAGAATAGTTGCTGGAACATTGCTATTAGTGGGAACAAGTTAC   16860
PMP1664      ATAAATTAATTTAAACTAAGAATAGTTGCTGGAACATTGCTATTAGTGGGAACAAGTTAC   16860
PMP1662      ATAAATTAATTTAAACTAAGAATAGTTGCTGGAACATTGCTATTAGTGGGAACAAGTTAC   16860
PMP1611      ATAAATTAATTTAAACTAAGAATAGTTGCTGGAACATTGCTATTAGTGGGAACAAGTTAC   16860
```

```
                ************************************************************

PMP1610         GAAATAATTTATCAATTTTAAAGACGTTTTTTAGGAAATATAAAATAATGGATTCTATCA    16920
PMP1666         GAAATAATTTATCAATTTTAAAGACGTTTTTTAGGAAATATAAAATAATGGATTCTATCA    16920
PMP1665         GAAATAATTTATCAATTTTAAAGACGTTTTTTAGGAAATATAAAATAATGGATTCTATCA    16920
PMP1667         GAAATAATTTATCAATTTTAAAGACGTTTTTTAGGAAATATAAAATAATGGATTCTATCA    16920
PMP1663         GAAATAATTTATCAATTTTAAAGACGTTTTTTAGGAAATATAAAATAATGGATTCTATCA    16920
PMP1664         GAAATAATTTATCAATTTTAAAGACGTTTTTTAGGAAATATAAAATAATGGATTCTATCA    16920
PMP1662         GAAATAATTTATCAATTTTAAAGACGTTTTTTAGGAAATATAAAATAATGGATTCTATCA    16920
PMP1611         GAAATAATTTATCAATTTTAAAGACGTTTTTTAGGAAATATAAAATAATGGATTCTATCA    16920
                ************************************************************

PMP1610         ACAATTCTAATAGTAATGATAATGCTAGAAAATCAAACTGTTCATTTTCAAAGGAAGATT    16980
PMP1666         ACAATTCTAATAGTAATGATAATGCTAGAAAATCAAACTGTTCATTTTCAAAGGAAGATT    16980
PMP1665         ACAATTCTAATAGTAATGATAATGCTAGAAAATCAAACTGTTCATTTTCAAAGGAAGATT    16980
PMP1667         ACAATTCTAATAGTAATGATAATGCTAGAAAATCAAACTGTTCATTTTCAAAGGAAGATT    16980
PMP1663         ACAATTCTAATAGTAATGATAATGCTAGAAAATCAAACTGTTCATTTTCAAAGGAAGATT    16980
PMP1664         ACAATTCTAATAGTAATGATAATGCTAGAAAATCAAACTGTTCATTTTCAAAGGAAGATT    16980
PMP1662         ACAATTCTAATAGTAATGATAATGCTAGAAAATCAAACTGTTCATTTTCAAAGGAAGATT    16980
PMP1611         ACAATTCTAATAGTAATGATAATGCTAGAAAATCAAACTGTTCATTTTCAAAGGAAGATT    16980
                ************************************************************

PMP1610         ATTTTTAAAATAATATATCTT----TAAGTAAGAATCATTTTATTAGACTTAATCTAGCC    17036
PMP1666         ATTTTTAAAATAATATATCTTTAAGTAAGTAAGAATCATTTTATTAGACTTAATCTAGCC    17040
PMP1665         ATTTTTAAAATAATATATCTT----TAAGTAAGAATCATTTTATTAGACTTAATCTAGCC    17036
PMP1667         ATTTTTAAAATAATATATCTT----TAAGTAAGAATCATTTTATTAGACTTAATCTAGCC    17036
PMP1663         ATTTTTAAAATAATATATCTT----TAAGTAAGAATCATTTTATTAGACTTAATCTAGCC    17036
PMP1664         ATTTTTAAAATAATATATCTT----TAAGTAAGAATCATTTTATTAGACTTAATCTAGCC    17036
PMP1662         ATTTTTAAAATAATATATCTT----TAAGTAAGAATCATTTTATTAGACTTAATCTAGCC    17036
PMP1611         ATTTTTAAAATAATATATCTT----TAAGTAAGAATCATTTTATTAGACTTAATCTAGCC    17036
                *********************    ***********************************

PMP1610         TATCAGTTAAATTAAAATATCAACTTTGATTAATTAAAAATTAGCAAAATTTATTGACAT    17096
PMP1666         TATCAGTTAAATTAAAATATCAACTTTGATTAATTAAAAATTAGCAAAATTTATTGACAT    17100
PMP1665         TATCAGTTAAATTAAAATATCAACTTTGATTAATTAAAAATTAGCAAAATTTATTGACAT    17096
PMP1667         TATCAGTTAAATTAAAATATCAACTTTGATTAATTAAAAATTAGCAAAATTTATTGACAT    17096
```

```
PMP1663      TATCAGTTAAATTAAAATATCAACTTTGATTAATTAAAAATTAGCAAAATTTATTGACAT   17096
PMP1664      TATCAGTTAAATTAAAATATCAACTTTGATTAATTAAAAATTAGCAAAATTTATTGACAT   17096
PMP1662      TATCAGTTAAATTAAAATATCAACTTTGATTAATTAAAAATTAGCAAAATTTATTGACAT   17096
PMP1611      TATCAGTTAAATTAAAATATCAACTTTGATTAATTAAAAATTAGCAAAATTTATTGACAT   17096
             ************************************************************

PMP1610      TTTGCTTTGATAAATTGCAATAAAGGTCTAATTCTGAATTTCAGGGAATATAAGAAAGGT   17156
PMP1666      TTTGCTTTGATAAATTGCAATAAAGGTCTAATTCTGAATTTCAGGGAATATAAGAAAGGT   17160
PMP1665      TTTGCTTTGATAAATTGCAATAAAGGTCTAATTCTGAATTTCAGGGAATATAAGAAAGGT   17156
PMP1667      TTTGCTTTGATAAATTGCAATAAAGGTCTAATTCTGAATTTCAGGGAATATAAGAAAGGT   17156
PMP1663      TTTGCTTTGATAAATTGCAATAAAGGTCTAATTCTGAATTTCAGGGAATATAAGAAAGGT   17156
PMP1664      TTTGCTTTGATAAATTGCAATAAAGGTCTAATTCTGAATTTCAGGGAATATAAGAAAGGT   17156
PMP1662      TTTGCTTTGATAAATTGCAATAAAGGTCTAATTCTGAATTTCAGGGAATATAAGAAAGGT   17156
PMP1611      TTTGCTTTGATAAATTGCAATAAAGGTCTAATTCTGAATTTCAGGGAATATAAGAAAGGT   17156
             ************************************************************

PMP1610      ACACTATTATGAAAGGTATTATTCTTGCAGGCGGCTCAGGTACCCGCCTGTACCCACTTA   17216
PMP1666      ACACTATTATGAAAGGTATTATTCTTGCAGGCGGCTCAGGTACCCGCCTGTACCCACTTA   17220
PMP1665      ACACTATTATGAAAGGTATTATTCTTGCAGGCGGCTCAGGTACCCGCCTGTACCCACTTA   17216
PMP1667      ACACTATTATGAAAGGTATTATTCTTGCAGGCGGCTCAGGTACCCGCCTGTACCCACTTA   17216
PMP1663      ACACTATTATGAAAGGTATTATTCTTGCAGGCGGCTCAGGTACCCGCCTGTACCCACTTA   17216
PMP1664      ACACTATTATGAAAGGTATTATTCTTGCAGGCGGCTCAGGTACCCGCCTGTACCCACTTA   17216
PMP1662      ACACTATTATGAAAGGTATTATTCTTGCAGGCGGCTCAGGTACCCGCCTGTACCCACTTA   17216
PMP1611      ACACTATTATGAAAGGTATTATTCTTGCAGGCGGCTCAGGTACCCGCCTGTACCCACTTA   17216
             ************************************************************

PMP1610      CTCGGGCTGCGTCAAAACAGCTGATGCCGGTTTATGATAAACCTATGATTTATTATCCGT   17276
PMP1666      CTCGGGCTGCGTCAAAACAGCTGATGCCGGTTTATGATAAACCTATGATTTATTATCCGT   17280
PMP1665      CTCGGGCTGCGTCAAAACAGCTGATGCCGGTTTATGATAAACCTATGATTTATTATCCGT   17276
PMP1667      CTCGGGCTGCGTCAAAACAGCTGATGCCGGTTTATGATAAACCTATGATTTATTATCCGT   17276
PMP1663      CTCGGGCTGCGTCAAAACAGCTGATGCCGGTTTATGATAAACCTATGATTTATTATCCGT   17276
PMP1664      CTCGGGCTGCGTCAAAACAGCTGATGCCGGTTTATGATAAACCTATGATTTATTATCCGT   17276
PMP1662      CTCGGGCTGCGTCAAAACAGCTGATGCCGGTTTATGATAAACCTATGATTTATTATCCGT   17276
PMP1611      CTCGGGCTGCGTCAAAACAGCTGATGCCGGTTTATGATAAACCTATGATTTATTATCCGT   17276
             ************************************************************
```

```
PMP1610      TGTCGACATTAATGTTGGCTGGAATTAAAGATATTTTGATTATCTCAACTCCTCAAGATT   17336
PMP1666      TGTCGACATTAATGTTGGCTGGAATTAAAGATATTTTGATTATCTCAACTCCTCAAGATT   17340
PMP1665      TGTCGACATTAATGTTGGCTGGAATTAAAGATATTTTGATTATCTCAACTCCTCAAGATT   17336
PMP1667      TGTCGACATTAATGTTGGCTGGAATTAAAGATATTTTGATTATCTCAACTCCTCAAGATT   17336
PMP1663      TGTCGACATTAATGTTGGCTGGAATTAAAGATATTTTGATTATCTCAACTCCTCAAGATT   17336
PMP1664      TGTCGACATTAATGTTGGCTGGAATTAAAGATATTTTGATTATCTCAACTCCTCAAGATT   17336
PMP1662      TGTCGACATTAATGTTGGCTGGAATTAAAGATATTTTGATTATCTCAACTCCTCAAGATT   17336
PMP1611      TGTCGACATTAATGTTGGCTGGAATTAAAGATATTTTGATTATCTCAACTCCTCAAGATT   17336
             ************************************************************

PMP1610      TGCCCCGGTTTAAGGACTTGCTCTTGGATGGTTCCGAATTTGGGATCAAGCTTTCCTATG   17396
PMP1666      TGCCCCGGTTTAAGGACTTGCTCTTGGATGGTTCCGAATTTGGGATCAAGCTTTCCTATG   17400
PMP1665      TGCCCCGGTTTAAGGACTTGCTCTTGGATGGTTCCGAATTTGGGATCAAGCTTTCCTATG   17396
PMP1667      TGCCCCGGTTTAAGGACTTGCTCTTGGATGGTTCCGAATTTGGGATCAAGCTTTCCTATG   17396
PMP1663      TGCCCCGGTTTAAGGACTTGCTCTTGGATGGTTCCGAATTTGGGATCAAGCTTTCCTATG   17396
PMP1664      TGCCCCGGTTTAAGGACTTGCTCTTGGATGGTTCCGAATTTGGGATCAAGCTTTCCTATG   17396
PMP1662      TGCCCCGGTTTAAGGACTTGCTCTTGGATGGTTCCGAATTTGGGATCAAGCTTTCCTATG   17396
PMP1611      TGCCCCGGTTTAAGGACTTGCTCTTGGATGGTTCCGAATTTGGGATCAAGCTTTCCTATG   17396
             ************************************************************

PMP1610      CGGAACAACCTAGTCCCGATGGACTTGCTCAGGCTTTTCTTATCGGTGAAGAATTTATCG   17456
PMP1666      CGGAACAACCTAGTCCCGATGGACTTGCTCAGGCTTTTCTTATCGGTGAAGAATTTATCG   17460
PMP1665      CGGAACAACCTAGTCCCGATGGACTTGCTCAGGCTTTTCTTATCGGTGAAGAATTTATCG   17456
PMP1667      CGGAACAACCTAGTCCCGATGGACTTGCTCAGGCTTTTCTTATCGGTGAAGAATTTATCG   17456
PMP1663      CGGAACAACCTAGTCCCGATGGACTTGCTCAGGCTTTTCTTATCGGTGAAGAATTTATCG   17456
PMP1664      CGGAACAACCTAGTCCCGATGGACTTGCTCAGGCTTTTCTTATCGGTGAAGAATTTATCG   17456
PMP1662      CGGAACAACCTAGTCCCGATGGACTTGCTCAGGCTTTTCTTATCGGTGAAGAATTTATCG   17456
PMP1611      CGGAACAACCTAGTCCCGATGGACTTGCTCAGGCTTTTCTTATCGGTGAAGAATTTATCG   17456
             ************************************************************

PMP1610      GTGACGATAGTGTTGCCTTGATTTTGGGCGACAATATCTATCATGGACCTGGTTTGAGCA   17516
PMP1666      GTGACGATAGTGTTGCCTTGATTTTGGGCGACAATATCTATCATGGACCTGGTTTGAGCA   17520
PMP1665      GTGACGATAGTGTTGCCTTGATTTTGGGCGACAATATCTATCATGGACCTGGTTTGAGCA   17516
PMP1667      GTGACGATAGTGTTGCCTTGATTTTGGGCGACAATATCTATCATGGACCTGGTTTGAGCA   17516
PMP1663      GTGACGATAGTGTTGCCTTGATTTTGGGCGACAATATCTATCATGGACCTGGTTTGAGCA   17516
PMP1664      GTGACGATAGTGTTGCCTTGATTTTGGGCGACAATATCTATCATGGACCTGGTTTGAGCA   17516
```

```
PMP1662      GTGACGATAGTGTTGCCTTGATTTTGGGCGACAATATCTATCATGGACCTGGTTTGAGCA   17516
PMP1611      GTGACGATAGTGTTGCCTTGATTTTGGGCGACAATATCTATCATGGACCTGGTTTGAGCA   17516
             ************************************************************

PMP1610      AAATGCTTCAAAAGGCAGCCCAGAAAGAGAAAGGTGCGACTGTTTTTGGCTACCAAGTGA   17576
PMP1666      AAATGCTTCAAAAGGCAGCCCAGAAAGAGAAAGGTGCGACTGTTTTTGGCTACCAAGTGA   17580
PMP1665      AAATGCTTCAAAAGGCAGCCCAGAAAGAGAAAGGTGCGACTGTTTTTGGCTACCAAGTGA   17576
PMP1667      AAATGCTTCAAAAGGCAGCCCAGAAAGAGAAAGGTGCGACTGTTTTTGGCTACCAAGTGA   17576
PMP1663      AAATGCTTCAAAAGGCAGCCCAGAAAGAGAAAGGTGCGACTGTTTTTGGCTACCAAGTGA   17576
PMP1664      AAATGCTTCAAAAGGCAGCCCAGAAAGAGAAAGGTGCGACTGTTTTTGGCTACCAAGTGA   17576
PMP1662      AAATGCTTCAAAAGGCAGCCCAGAAAGAGAAAGGTGCGACTGTTTTTGGCTACCAAGTGA   17576
PMP1611      AAATGCTTCAAAAGGCAGCCCAGAAAGAGAAAGGTGCGACTGTTTTTGGCTACCAAGTGA   17576
             ************************************************************

PMP1610      AGGATCCAGAGCGTTTTGGTGTGGTCGAGTTTGATACAGACATGAATGCCATTTCCATAG   17636
PMP1666      AGGATCCAGAGCGTTTTGGTGTGGTCGAGTTTGATACAGACATGAATGCCATTTCCATAG   17640
PMP1665      AGGATCCAGAGCGTTTTGGTGTGGTCGAGTTTGATACAGACATGAATGCCATTTCCATAG   17636
PMP1667      AGGATCCAGAGCGTTTTGGTGTGGTCGAGTTTGATACAGACATGAATGCCATTTCCATAG   17636
PMP1663      AGGATCCAGAGCGTTTTGGTGTGGTCGAGTTTGATACAGACATGAATGCCATTTCCATAG   17636
PMP1664      AGGATCCAGAGCGTTTTGGTGTGGTCGAGTTTGATACAGACATGAATGCCATTTCCATAG   17636
PMP1662      AGGATCCAGAGCGTTTTGGTGTGGTCGAGTTTGATACAGACATGAATGCCATTTCCATAG   17636
PMP1611      AGGATCCAGAGCGTTTTGGTGTGGTCGAGTTTGATACAGACATGAATGCCATTTCCATAG   17636
             ************************************************************

PMP1610      AAGAAAAACCAGAGAATCCTCGCTCCAACTATGCCGTGACCGGTCTGTATTTCTATGATA   17696
PMP1666      AAGAAAAACCAGAGAATCCTCGCTCCAACTATGCCGTGACCGGTCTGTATTTCTATGATA   17700
PMP1665      AAGAAAAACCAGAGAATCCTCGCTCCAACTATGCCGTGACCGGTCTGTATTTCTATGATA   17696
PMP1667      AAGAAAAACCAGAGAATCCTCGCTCCAACTATGCCGTGACCGGTCTGTATTTCTATGATA   17696
PMP1663      AAGAAAAACCAGAGAATCCTCGCTCCAACTATGCCGTGACCGGTCTGTATTTCTATGATA   17696
PMP1664      AAGAAAAACCAGAGAATCCTCGCTCCAACTATGCCGTGACCGGTCTGTATTTCTATGATA   17696
PMP1662      AAGAAAAACCAGAGAATCCTCGCTCCAACTATGCCGTGACCGGTCTGTATTTCTATGATA   17696
PMP1611      AAGAAAAACCAGAGAATCCTCGCTCCAACTATGCCGTGACCGGTCTGTATTTCTATGATA   17696
             ************************************************************

PMP1610      ATGATGTTGTAGAAATTGCTAAAGGTATTAAACCAAGTGCACGTGGCGAGTTAGAAATTA   17756
PMP1666      ATGATGTTGTAGAAATTGCTAAAGGTATTAAACCAAGTGCACGTGGCGAGTTAGAAATTA   17760
```

```
PMP1665      ATGATGTTGTAGAAATTGCTAAAGGTATTAAACCAAGTGCACGTGGCGAGTTAGAAATTA   17756
PMP1667      ATGATGTTGTAGAAATTGCTAAAGGTATTAAACCAAGTGCACGTGGCGAGTTAGAAATTA   17756
PMP1663      ATGATGTTGTAGAAATTGCTAAAGGTATTAAACCAAGTGCACGTGGCGAGTTAGAAATTA   17756
PMP1664      ATGATGTTGTAGAAATTGCTAAAGGTATTAAACCAAGTGCACGTGGCGAGTTAGAAATTA   17756
PMP1662      ATGATGTTGTAGAAATTGCTAAAGGTATTAAACCAAGTGCACGTGGCGAGTTAGAAATTA   17756
PMP1611      ATGATGTTGTAGAAATTGCTAAAGGTATTAAACCAAGTGCACGTGGCGAGTTAGAAATTA   17756
             ************************************************************

PMP1610      CAGATATCAACAAGGCTTACCTAAATCGTGGTGACCTTTCTGTTGAGCTGATGGGGCGTG   17816
PMP1666      CAGATATCAACAAGGCTTACCTAAATCGTGGTGACCTTTCTGTTGAGCTGATGGGGCGTG   17820
PMP1665      CAGATATCAACAAGGCTTACCTAAATCGTGGTGACCTTTCTGTTGAGCTGATGGGGCGTG   17816
PMP1667      CAGATATCAACAAGGCTTACCTAAATCGTGGTGACCTTTCTGTTGAGCTGATGGGGCGTG   17816
PMP1663      CAGATATCAACAAGGCTTACCTAAATCGTGGTGACCTTTCTGTTGAGCTGATGGGGCGTG   17816
PMP1664      CAGATATCAACAAGGCTTACCTAAATCGTGGTGACCTTTCTGTTGAGCTGATGGGGCGTG   17816
PMP1662      CAGATATCAACAAGGCTTACCTAAATCGTGGTGACCTTTCTGTTGAGCTGATGGGGCGTG   17816
PMP1611      CAGATATCAACAAGGCTTACCTAAATCGTGGTGACCTTTCTGTTGAGCTGATGGGGCGTG   17816
             ************************************************************

PMP1610      GTTTTGCCTGGTTGGATACGGGAACCCATGAAAGCCTGCTAGAAGCTTCTCAGTATATCG   17876
PMP1666      GTTTTGCCTGGTTGGATACGGGAACCCATGAAAGCCTGCTAGAAGCTTCTCAGTATATCG   17880
PMP1665      GTTTTGCCTGGTTGGATACGGGAACCCATGAAAGCCTGCTAGAAGCTTCTCAGTATATCG   17876
PMP1667      GTTTTGCCTGGTTGGATACGGGAACCCATGAAAGCCTGCTAGAAGCTTCTCAGTATATCG   17876
PMP1663      GTTTTGCCTGGTTGGATACGGGAACCCATGAAAGCCTGCTAGAAGCTTCTCAGTATATCG   17876
PMP1664      GTTTTGCCTGGTTGGATACGGGAACCCATGAAAGCCTGCTAGAAGCTTCTCAGTATATCG   17876
PMP1662      GTTTTGCCTGGTTGGATACGGGAACCCATGAAAGCCTGCTAGAAGCTTCTCAGTATATCG   17876
PMP1611      GTTTTGCCTGGTTGGATACGGGAACCCATGAAAGCCTGCTAGAAGCTTCTCAGTATATCG   17876
             ************************************************************

PMP1610      AAACAGTTCAACGGATGCAGAATGTTCAAGTTGCAAACTTGGAAGAAATTGCCTATCGCA   17936
PMP1666      AAACAGTTCAACGGATGCAGAATGTTCAAGTTGCAAACTTGGAAGAAATTGCCTATCGCA   17940
PMP1665      AAACAGTTCAACGGATGCAGAATGTTCAAGTTGCAAACTTGGAAGAAATTGCCTATCGCA   17936
PMP1667      AAACAGTTCAACGGATGCAGAATGTTCAAGTTGCAAACTTGGAAGAAATTGCCTATCGCA   17936
PMP1663      AAACAGTTCAACGGATGCAGAATGTTCAAGTTGCAAACTTGGAAGAAATTGCCTATCGCA   17936
PMP1664      AAACAGTTCAACGGATGCAGAATGTTCAAGTTGCAAACTTGGAAGAAATTGCCTATCGCA   17936
PMP1662      AAACAGTTCAACGGATGCAGAATGTTCAAGTTGCAAACTTGGAAGAAATTGCCTATCGCA   17936
PMP1611      AAACAGTTCAACGGATGCAGAATGTTCAAGTTGCAAACTTGGAAGAAATTGCCTATCGCA   17936
```

```
             ************************************************************

PMP1610      TGGGCTATATCAGTTGTGAAGATGTGCTCGAGTTGGCGCAACCTCTGAAGAAGAATGAAT   17996
PMP1666      TGGGCTATATCAGTTGTGAAGATGTGCTCGAGTTGGCGCAACCTCTGAAGAAGAATGAAT   18000
PMP1665      TGGGCTATATCAGTTGTGAAGATGTGCTCGAGTTGGCGCAACCTCTGAAGAAGAATGAAT   17996
PMP1667      TGGGCTATATCAGTTGTGAAGATGTGCTCGAGTTGGCGCAACCTCTGAAGAAGAATGAAT   17996
PMP1663      TGGGCTATATCAGTTGTGAAGATGTGCTCGAGTTGGCGCAACCTCTGAAGAAGAATGAAT   17996
PMP1664      TGGGCTATATCAGTTGTGAAGATGTGCTCGAGTTGGCGCAACCTCTGAAGAAGAATGAAT   17996
PMP1662      TGGGCTATATCAGTTGTGAAGATGTGCTCGAGTTGGCGCAACCTCTGAAGAAGAATGAAT   17996
PMP1611      TGGGCTATATCAGTTGTGAAGATGTGCTCGAGTTGGCGCAACCTCTGAAGAAGAATGAAT   17996
             ************************************************************

PMP1610      ACGGACAATATTTGCTCCGTTTGATTGGAGAAGCATAGATGTCAGATAATTTTTTTGGAA   18056
PMP1666      ACGGACAATATTTGCTCCGTTTGATTGGAGAAGCATAGATGTCAGATAATTTTTTTGGAA   18060
PMP1665      ACGGACAATATTTGCTCCGTTTGATTGGAGAAGCATAGATGTCAGATAATTTTTTTGGAA   18056
PMP1667      ACGGACAATATTTGCTCCGTTTGATTGGAGAAGCATAGATGTCAGATAATTTTTTTGGAA   18056
PMP1663      ACGGACAATATTTGCTCCGTTTGATTGGAGAAGCATAGATGTCAGATAATTTTTTTGGAA   18056
PMP1664      ACGGACAATATTTGCTCCGTTTGATTGGAGAAGCATAGATGTCAGATAATTTTTTTGGAA   18056
PMP1662      ACGGACAATATTTGCTCCGTTTGATTGGAGAAGCATAGATGTCAGATAATTTTTTTGGAA   18056
PMP1611      ACGGACAATATTTGCTCCGTTTGATTGGAGAAGCATAGATGTCAGATAATTTTTTTGGAA   18056
             ************************************************************

PMP1610      AAACACTTGCAGTGCGTAAGATTGATGCTATACCAGGACTGCTAGAGTTTGACATTCCCG   18116
PMP1666      AAACACTTGCAGTGCGTAAGATTGATGCTATACCAGGACTGCTAGAGTTTGACATTCCCG   18120
PMP1665      AAACACTTGCAGTGCGTAAGATTGATGCTATACCAGGACTGCTAGAGTTTGACATTCCCG   18116
PMP1667      AAACACTTGCAGTGCGTAAGATTGATGCTATACCAGGACTGCTAGAGTTTGACATTCCCG   18116
PMP1663      AAACACTTGCAGTGCGTAAGATTGATGCTATACCAGGACTGCTAGAGTTTGACATTCCCG   18116
PMP1664      AAACACTTGCAGTGCGTAAGATTGATGCTATACCAGGACTGCTAGAGTTTGACATTCCCG   18116
PMP1662      AAACACTTGCAGTGCGTAAGATTGATGCTATACCAGGACTGCTAGAGTTTGACATTCCCG   18116
PMP1611      AAACACTTGCAGTGCGTAAGATTGATGCTATACCAGGACTGCTAGAGTTTGACATTCCCG   18116
             ************************************************************

PMP1610      TTCATGGAGACAATCGTGGTTGGTTTAAGGAAAACTTCCAGAAGGAAAAGATGGAGCCAC   18176
PMP1666      TTCATGGAGACAATCGTGGTTGGTTTAAGGAAAACTTCCAGAAGGAAAAGATGGAGCCAC   18180
PMP1665      TTCATGGAGACAATCGTGGTTGGTTTAAGGAAAACTTCCAGAAGGAAAAGATGGAGCCAC   18176
PMP1667      TTCATGGAGACAATCGTGGTTGGTTTAAGGAAAACTTCCAGAAGGAAAAGATGGAGCCAC   18176
```

```
PMP1663      TTCATGGAGACAATCGTGGTTGGTTTAAGGAAAACTTCCAGAAGGAAAAGATGGAGCCAC   18176
PMP1664      TTCATGGAGACAATCGTGGTTGGTTTAAGGAAAACTTCCAGAAGGAAAAGATGGAGCCAC   18176
PMP1662      TTCATGGAGACAATCGTGGTTGGTTTAAGGAAAACTTCCAGAAGGAAAAGATGGAGCCAC   18176
PMP1611      TTCATGGAGACAATCGTGGTTGGTTTAAGGAAAACTTCCAGAAGGAAAAGATGGAGCCAC   18176
             ************************************************************

PMP1610      TTGGCTTTCCTGAAAGCTTCTTTGCTGCAGGGAAATTGCAAAACAACGTCAGCTTTTCTC   18236
PMP1666      TTGGCTTTCCTGAAAGCTTCTTTGCTGCAGGGAAATTGCAAAACAACGTCAGCTTTTCTC   18240
PMP1665      TTGGCTTTCCTGAAAGCTTCTTTGCTGCAGGGAAATTGCAAAACAACGTCAGCTTTTCTC   18236
PMP1667      TTGGCTTTCCTGAAAGCTTCTTTGCTGCAGGGAAATTGCAAAACAACGTCAGCTTTTCTC   18236
PMP1663      TTGGCTTTCCTGAAAGCTTCTTTGCTGCAGGGAAATTGCAAAACAACGTCAGCTTTTCTC   18236
PMP1664      TTGGCTTTCCTGAAAGCTTCTTTGCTGCAGGGAAATTGCAAAACAACGTCAGCTTTTCTC   18236
PMP1662      TTGGCTTTCCTGAAAGCTTCTTTGCTGCAGGGAAATTGCAAAACAACGTCAGCTTTTCTC   18236
PMP1611      TTGGCTTTCCTGAAAGCTTCTTTGCTGCAGGGAAATTGCAAAACAACGTCAGCTTTTCTC   18236
             ************************************************************

PMP1610      GCAAAAATGTTCTTCGAGGATTGCATGCAGAACCTTGGGACAAGTATATCTCTGTTGCAG   18296
PMP1666      GCAAAAATGTTCTTCGAGGATTGCATGCAGAACCTTGGGACAAGTATATCTCTGTTGCAG   18300
PMP1665      GCAAAAATGTTCTTCGAGGATTGCATGCAGAACCTTGGGACAAGTATATCTCTGTTGCAG   18296
PMP1667      GCAAAAATGTTCTTCGAGGATTGCATGCAGAACCTTGGGACAAGTATATCTCTGTTGCAG   18296
PMP1663      GCAAAAATGTTCTTCGAGGATTGCATGCAGAACCTTGGGACAAGTATATCTCTGTTGCAG   18296
PMP1664      GCAAAAATGTTCTTCGAGGATTGCATGCAGAACCTTGGGACAAGTATATCTCTGTTGCAG   18296
PMP1662      GCAAAAATGTTCTTCGAGGATTGCATGCAGAACCTTGGGACAAGTATATCTCTGTTGCAG   18296
PMP1611      GCAAAAATGTTCTTCGAGGATTGCATGCAGAACCTTGGGACAAGTATATCTCTGTTGCAG   18296
             ************************************************************

PMP1610      ACGATGGGAAGGTTTTAGGATCTTGGGTTGATCTACGCGAGGGTGAAACCTTTGGGAATA   18356
PMP1666      ACGATGGGAAGGTTTTAGGATCTTGGGTTGATCTACGCGAGGGTGAAACCTTTGGGAATA   18360
PMP1665      ACGATGGGAAGGTTTTAGGATCTTGGGTTGATCTACGCGAGGGTGAAACCTTTGGGAATA   18356
PMP1667      ACGATGGGAAGGTTTTAGGATCTTGGGTTGATCTACGCGAGGGTGAAACCTTTGGGAATA   18356
PMP1663      ACGATGGGAAGGTTTTAGGATCTTGGGTTGATCTACGCGAGGGTGAAACCTTTGGGAATA   18356
PMP1664      ACGATGGGAAGGTTTTAGGATCTTGGGTTGATCTACGCGAGGGTGAAACCTTTGGGAATA   18356
PMP1662      ACGATGGGAAGGTTTTAGGATCTTGGGTTGATCTACGCGAGGGTGAAACCTTTGGGAATA   18356
PMP1611      ACGATGGGAAGGTTTTAGGATCTTGGGTTGATCTACGCGAGGGTGAAACCTTTGGGAATA   18356
             ************************************************************
```

```
PMP1610      CCTATCAGACAGTGATTGATGCGAGTAAGGGAATCTTTGTTCCTCGAGGCGTAGCTAATG   18416
PMP1666      CCTATCAGACAGTGATTGATGCGAGTAAGGGAATCTTTGTTCCTCGAGGCGTAGCTAATG   18420
PMP1665      CCTATCAGACAGTGATTGATGCGAGTAAGGGAATCTTTGTTCCTCGAGGCGTAGCTAATG   18416
PMP1667      CCTATCAGACAGTGATTGATGCGAGTAAGGGAATCTTTGTTCCTCGAGGCGTAGCTAATG   18416
PMP1663      CCTATCAGACAGTGATTGATGCGAGTAAGGGAATCTTTGTTCCTCGAGGCGTAGCTAATG   18416
PMP1664      CCTATCAGACAGTGATTGATGCGAGTAAGGGAATCTTTGTTCCTCGAGGCGTAGCTAATG   18416
PMP1662      CCTATCAGACAGTGATTGATGCGAGTAAGGGAATCTTTGTTCCTCGAGGCGTAGCTAATG   18416
PMP1611      CCTATCAGACAGTGATTGATGCGAGTAAGGGAATCTTTGTTCCTCGAGGCGTAGCTAATG   18416
             ************************************************************

PMP1610      GCTTCCAAGTTTTATCAGATACAGTGTCATATAGCTATCTGGTCAATGATTACTGGGCTC   18476
PMP1666      GCTTCCAAGTTTTATCAGATACAGTGTCATATAGCTATCTGGTCAATGATTACTGGGCTC   18480
PMP1665      GCTTCCAAGTTTTATCAGATACAGTGTCATATAGCTATCTGGTCAATGATTACTGGGCTC   18476
PMP1667      GCTTCCAAGTTTTATCAGATACAGTGTCATATAGCTATCTGGTCAATGATTACTGGGCTC   18476
PMP1663      GCTTCCAAGTTTTATCAGATACAGTGTCATATAGCTATCTGGTCAATGATTACTGGGCTC   18476
PMP1664      GCTTCCAAGTTTTATCAGATACAGTGTCATATAGCTATCTGGTCAATGATTACTGGGCTC   18476
PMP1662      GCTTCCAAGTTTTATCAGATACAGTGTCATATAGCTATCTGGTCAATGATTACTGGGCTC   18476
PMP1611      GCTTCCAAGTTTTATCAGATACAGTGTCATATAGCTATCTGGTCAATGATTACTGGGCTC   18476
             ************************************************************

PMP1610      TTGAACTCAAACCCAAGTATGCCTTTGTGAACTACGCTGATCCAAGCCTTGGTATTGAAT   18536
PMP1666      TTGAACTCAAACCCAAGTATGCCTTTGTGAACTACGCTGATCCAAGCCTTGGTATTGAAT   18540
PMP1665      TTGAACTCAAACCCAAGTATGCCTTTGTGAACTACGCTGATCCAAGCCTTGGTATTGAAT   18536
PMP1667      TTGAACTCAAACCCAAGTATGCCTTTGTGAACTACGCTGATCCAAGCCTTGGTATTGAAT   18536
PMP1663      TTGAACTCAAACCCAAGTATGCCTTTGTGAACTACGCTGATCCAAGCCTTGGTATTGAAT   18536
PMP1664      TTGAACTCAAACCCAAGTATGCCTTTGTGAACTACGCTGATCCAAGCCTTGGTATTGAAT   18536
PMP1662      TTGAACTCAAACCCAAGTATGCCTTTGTGAACTACGCTGATCCAAGCCTTGGTATTGAAT   18536
PMP1611      TTGAACTCAAACCCAAGTATGCCTTTGTGAACTACGCTGATCCAAGCCTTGGTATTGAAT   18536
             ************************************************************

PMP1610      GGGAAAATATTGCAGAAGCAGAGGTTTCAGAAGCAGATAAAAATCATCCACTACTTAAGG   18596
PMP1666      GGGAAAATATTGCAGAAGCAGAGGTTTCAGAAGCAGATAAAAATCATCCACTACTTAAGG   18600
PMP1665      GGGAAAATATTGCAGAAGCAGAGGTTTCAGAAGCAGATAAAAATCATCCACTACTTAAGG   18596
PMP1667      GGGAAAATATTGCAGAAGCAGAGGTTTCAGAAGCAGATAAAAATCATCCACTACTTAAGG   18596
PMP1663      GGGAAAATATTGCAGAAGCAGAGGTTTCAGAAGCAGATAAAAATCATCCACTACTTAAGG   18596
PMP1664      GGGAAAATATTGCAGAAGCAGAGGTTTCAGAAGCAGATAAAAATCATCCACTACTTAAGG   18596
```

```
PMP1662      GGGAAAATATTGCAGAAGCAGAGGTTTCAGAAGCAGATAAAAATCATCCACTACTTAAGG   18596
PMP1611      GGGAAAATATTGCAGAAGCAGAGGTTTCAGAAGCAGATAAAAATCATCCACTACTTAAGG   18596
             ************************************************************

PMP1610      ATGTAAAACCTTTGAAAAAAGAAGATTTGGAATAAGGAAAGAATATGACTGAATACAAAA   18656
PMP1666      ATGTAAAACCTTTGAAAAAAGAAGATTTGGAATAAGGAAAGAATATGACTGAATACAAAA   18660
PMP1665      ATGTAAAACCTTTGAAAAAAGAAGATTTGGAATAAGGAAAGAATATGACTGAATACAAAA   18656
PMP1667      ATGTAAAACCTTTGAAAAAAGAAGATTTGGAATAAGGAAAGAATATGACTGAATACAAAA   18656
PMP1663      ATGTAAAACCTTTGAAAAAAGAAGATTTGGAATAAGGAAAGAATATGACTGAATACAAAA   18656
PMP1664      ATGTAAAACCTTTGAAAAAAGAAGATTTGGAATAAGGAAAGAATATGACTGAATACAAAA   18656
PMP1662      ATGTAAAACCTTTGAAAAAAGAAGATTTGGAATAAGGAAAGAATATGACTGAATACAAAA   18656
PMP1611      ATGTAAAACCTTTGAAAAAAGAAGATTTGGAATAAGGAAAGAATATGACTGAATACAAAA   18656
             ************************************************************

PMP1610      ATATTATCGTGACAGGTGGAGCTGGCTTTATCGGTTCTAACTTTGTCCATTATGTTTACG   18716
PMP1666      ATATTATCGTGACAGGTGGAGCTGGCTTTATCGGTTCTAACTTTGTCCATTATGTTTACG   18720
PMP1665      ATATTATCGTGACAGGTGGAGCTGGCTTTATCGGTTCTAACTTTGTCCATTATGTTTACG   18716
PMP1667      ATATTATCGTGACAGGTGGAGCTGGCTTTATCGGTTCTAACTTTGTCCATTATGTTTACG   18716
PMP1663      ATATTATCGTGACAGGTGGAGCTGGCTTTATCGGTTCTAACTTTGTCCATTATGTTTACG   18716
PMP1664      ATATTATCGTGACAGGTGGAGCTGGCTTTATCGGTTCTAACTTTGTCCATTATGTTTACG   18716
PMP1662      ATATTATCGTGACAGGTGGAGCTGGCTTTATCGGTTCTAACTTTGTCCATTATGTTTACG   18716
PMP1611      ATATTATCGTGACAGGTGGAGCTGGCTTTATCGGTTCTAACTTTGTCCATTATGTTTACG   18716
             ************************************************************

PMP1610      AGAACTTTCCAGATGTTCACGTGACAGTCCTAGATAAGTTGACTTATGCTGGAAACCGCG   18776
PMP1666      AGAACTTTCCAGATGTTCACGTGACAGTCCTAGATAAGTTGACTTATGCTGGAAACCGCG   18780
PMP1665      AGAACTTTCCAGATGTTCACGTGACAGTCCTAGATAAGTTGACTTATGCTGGAAACCGCG   18776
PMP1667      AGAACTTTCCAGATGTTCACGTGACAGTCCTAGATAAGTTGACTTATGCTGGAAACCGCG   18776
PMP1663      AGAACTTTCCAGATGTTCACGTGACAGTCCTAGATAAGTTGACTTATGCTGGAAACCGCG   18776
PMP1664      AGAACTTTCCAGATGTTCACGTGACAGTCCTAGATAAGTTGACTTATGCTGGAAACCGCG   18776
PMP1662      AGAACTTTCCAGATGTTCACGTGACAGTCCTAGATAAGTTGACTTATGCTGGAAACCGCG   18776
PMP1611      AGAACTTTCCAGATGTTCACGTGACAGTCCTAGATAAGTTGACTTATGCTGGAAACCGCG   18776
             ************************************************************

PMP1610      CGAATATTGAGGAAATTTTAGGTAATCGTGTTGAGTTAGTTGTTGGTGACATTGCTGATG   18836
PMP1666      CGAATATTGAGGAAATTTTAGGTAATCGTGTTGAGTTAGTTGTTGGTGACATTGCTGATG   18840
```

```
PMP1665      CGAATATTGAGGAAATTTTAGGTAATCGTGTTGAGTTAGTTGTTGGTGACATTGCTGATG   18836
PMP1667      CGAATATTGAGGAAATTTTAGGTAATCGTGTTGAGTTAGTTGTTGGTGACATTGCTGATG   18836
PMP1663      CGAATATTGAGGAAATTTTAGGTAATCGTGTTGAGTTAGTTGTTGGTGACATTGCTGATG   18836
PMP1664      CGAATATTGAGGAAATTTTAGGTAATCGTGTTGAGTTAGTTGTTGGTGACATTGCTGATG   18836
PMP1662      CGAATATTGAGGAAATTTTAGGTAATCGTGTTGAGTTAGTTGTTGGTGACATTGCTGATG   18836
PMP1611      CGAATATTGAGGAAATTTTAGGTAATCGTGTTGAGTTAGTTGTTGGTGACATTGCTGATG   18836
             ************************************************************

PMP1610      CGGAGTTGGTAGACAAGTTGGCTGCTCAAGCAGATGCTATCGTTCATTATGCAGCGGAAA   18896
PMP1666      CGGAGTTGGTAGACAAGTTGGCTGCTCAAGCAGATGCTATCGTTCATTATGCAGCGGAAA   18900
PMP1665      CGGAGTTGGTAGACAAGTTGGCTGCTCAAGCAGATGCTATCGTTCATTATGCAGCGGAAA   18896
PMP1667      CGGAGTTGGTAGACAAGTTGGCTGCTCAAGCAGATGCTATCGTTCATTATGCAGCGGAAA   18896
PMP1663      CGGAGTTGGTAGACAAGTTGGCTGCTCAAGCAGATGCTATCGTTCATTATGCAGCGGAAA   18896
PMP1664      CGGAGTTGGTAGACAAGTTGGCTGCTCAAGCAGATGCTATCGTTCATTATGCAGCGGAAA   18896
PMP1662      CGGAGTTGGTAGACAAGTTGGCTGCTCAAGCAGATGCTATCGTTCATTATGCAGCGGAAA   18896
PMP1611      CGGAGTTGGTAGACAAGTTGGCTGCTCAAGCAGATGCTATCGTTCATTATGCAGCGGAAA   18896
             ************************************************************

PMP1610      GCCACAATGATAATTCGCTCAATGATCCATCGCCATTTATTCATACTAACTTCATTGGAA   18956
PMP1666      GCCACAATGATAATTCGCTCAATGATCCATCGCCATTTATTCATACTAACTTCATTGGAA   18960
PMP1665      GCCACAATGATAATTCGCTCAATGATCCATCGCCATTTATTCATACTAACTTCATTGGAA   18956
PMP1667      GCCACAATGATAATTCGCTCAATGATCCATCGCCATTTATTCATACTAACTTCATTGGAA   18956
PMP1663      GCCACAATGATAATTCGCTCAATGATCCATCGCCATTTATTCATACTAACTTCATTGGAA   18956
PMP1664      GCCACAATGATAATTCGCTCAATGATCCATCGCCATTTATTCATACTAACTTCATTGGAA   18956
PMP1662      GCCACAATGATAATTCGCTCAATGATCCATCGCCATTTATTCATACTAACTTCATTGGAA   18956
PMP1611      GCCACAATGATAATTCGCTCAATGATCCATCGCCATTTATTCATACTAACTTCATTGGAA   18956
             ************************************************************

PMP1610      CCTATACTCTTTTAGAAGCTGCTCGTAAGTATGATATTCGCTTCCACCATGTATCGACAG   19016
PMP1666      CCTATACTCTTTTAGAAGCTGCTCGTAAGTATGATATTCGCTTCCACCATGTATCGACAG   19020
PMP1665      CCTATACTCTTTTAGAAGCTGCTCGTAAGTATGATATTCGCTTCCACCATGTATCGACAG   19016
PMP1667      CCTATACTCTTTTAGAAGCTGCTCGTAAGTATGATATTCGCTTCCACCATGTATCGACAG   19016
PMP1663      CCTATACTCTTTTAGAAGCTGCTCGTAAGTATGATATTCGCTTCCACCATGTATCGACAG   19016
PMP1664      CCTATACTCTTTTAGAAGCTGCTCGTAAGTATGATATTCGCTTCCACCATGTATCGACAG   19016
PMP1662      CCTATACTCTTTTAGAAGCTGCTCGTAAGTATGATATTCGCTTCCACCATGTATCGACAG   19016
PMP1611      CCTATACTCTTTTAGAAGCTGCTCGTAAGTATGATATTCGCTTCCACCATGTATCGACAG   19016
```

```
             ************************************************************

PMP1610      ATGAAGTTTATGGGGATCTCCCTTTACGCGAAGATTTGCCAGGTCATGGAGAAGGGCCGG   19076
PMP1666      ATGAAGTTTATGGGGATCTCCCTTTACGCGAAGATTTGCCAGGTCATGGAGAAGGGCCGG   19080
PMP1665      ATGAAGTTTATGGGGATCTCCCTTTACGCGAAGATTTGCCAGGTCATGGAGAAGGGCCGG   19076
PMP1667      ATGAAGTTTATGGGGATCTCCCTTTACGCGAAGATTTGCCAGGTCATGGAGAAGGGCCGG   19076
PMP1663      ATGAAGTTTATGGGGATCTCCCTTTACGCGAAGATTTGCCAGGTCATGGAGAAGGGCCGG   19076
PMP1664      ATGAAGTTTATGGGGATCTCCCTTTACGCGAAGATTTGCCAGGTCATGGAGAAGGGCCGG   19076
PMP1662      ATGAAGTTTATGGGGATCTCCCTTTACGCGAAGATTTGCCAGGTCATGGAGAAGGGCCGG   19076
PMP1611      ATGAAGTTTATGGGGATCTCCCTTTACGCGAAGATTTGCCAGGTCATGGAGAAGGGCCGG   19076
             ************************************************************

PMP1610      GTGAGAAATTTACGGCTGAAACCAAGTACAATCCAAGCTCGCCTTACTCATCAACCAAGG   19136
PMP1666      GTGAGAAATTTACGGCTGAAACCAAGTACAATCCAAGCTCGCCTTACTCATCAACCAAGG   19140
PMP1665      GTGAGAAATTTACGGCTGAAACCAAGTACAATCCAAGCTCGCCTTACTCATCAACCAAGG   19136
PMP1667      GTGAGAAATTTACGGCTGAAACCAAGTACAATCCAAGCTCGCCTTACTCATCAACCAAGG   19136
PMP1663      GTGAGAAATTTACGGCTGAAACCAAGTACAATCCAAGCTCGCCTTACTCATCAACCAAGG   19136
PMP1664      GTGAGAAATTTACGGCTGAAACCAAGTACAATCCAAGCTCGCCTTACTCATCAACCAAGG   19136
PMP1662      GTGAGAAATTTACGGCTGAAACCAAGTACAATCCAAGCTCGCCTTACTCATCAACCAAGG   19136
PMP1611      GTGAGAAATTTACGGCTGAAACCAAGTACAATCCAAGCTCGCCTTACTCATCAACCAAGG   19136
             ************************************************************

PMP1610      CAGCCTCAGATTTGATTGTCAAAGCCTGGGTGCGTTCTTTTGGAGTCAAGGCAACGATTT   19196
PMP1666      CAGCCTCAGATTTGATTGTCAAAGCCTGGGTGCGTTCTTTTGGAGTCAAGGCAACGATTT   19200
PMP1665      CAGCCTCAGATTTGATTGTCAAAGCCTGGGTGCGTTCTTTTGGAGTCAAGGCAACGATTT   19196
PMP1667      CAGCCTCAGATTTGATTGTCAAAGCCTGGGTGCGTTCTTTTGGAGTCAAGGCAACGATTT   19196
PMP1663      CAGCCTCAGATTTGATTGTCAAAGCCTGGGTGCGTTCTTTTGGAGTCAAGGCAACGATTT   19196
PMP1664      CAGCCTCAGATTTGATTGTCAAAGCCTGGGTGCGTTCTTTTGGAGTCAAGGCAACGATTT   19196
PMP1662      CAGCCTCAGATTTGATTGTCAAAGCCTGGGTGCGTTCTTTTGGAGTCAAGGCAACGATTT   19196
PMP1611      CAGCCTCAGATTTGATTGTCAAAGCCTGGGTGCGTTCTTTTGGAGTCAAGGCAACGATTT   19196
             ************************************************************

PMP1610      CCAACTGTTCAAATAACTACGGTCCTTATCAACATATCGAAAAATTCATCCCACGTCAGA   19256
PMP1666      CCAACTGTTCAAATAACTACGGTCCTTATCAACATATCGAAAAATTCATCCCACGTCAGA   19260
PMP1665      CCAACTGTTCAAATAACTACGGTCCTTATCAACATATCGAAAAATTCATCCCACGTCAGA   19256
PMP1667      CCAACTGTTCAAATAACTACGGTCCTTATCAACATATCGAAAAATTCATCCCACGTCAGA   19256
```

```
PMP1663      CCAACTGTTCAAATAACTACGGTCCTTATCAACATATCGAAAAATTCATCCCACGTCAGA   19256
PMP1664      CCAACTGTTCAAATAACTACGGTCCTTATCAACATATCGAAAAATTCATCCCACGTCAGA   19256
PMP1662      CCAACTGTTCAAATAACTACGGTCCTTATCAACATATCGAAAAATTCATCCCACGTCAGA   19256
PMP1611      CCAACTGTTCAAATAACTACGGTCCTTATCAACATATCGAAAAATTCATCCCACGTCAGA   19256
             ************************************************************

PMP1610      TTACTAACATCCTAAGTGGTATCAAGCCAAAACTTTACGGTGAAGGTAAAAACGTTCGTG   19316
PMP1666      TTACTAACATCCTAAGTGGTATCAAGCCAAAACTTTACGGTGAAGGTAAAAACGTTCGTG   19320
PMP1665      TTACTAACATCCTAAGTGGTATCAAGCCAAAACTTTACGGTGAAGGTAAAAACGTTCGTG   19316
PMP1667      TTACTAACATCCTAAGTGGTATCAAGCCAAAACTTTACGGTGAAGGTAAAAACGTTCGTG   19316
PMP1663      TTACTAACATCCTAAGTGGTATCAAGCCAAAACTTTACGGTGAAGGTAAAAACGTTCGTG   19316
PMP1664      TTACTAACATCCTAAGTGGTATCAAGCCAAAACTTTACGGTGAAGGTAAAAACGTTCGTG   19316
PMP1662      TTACTAACATCCTAAGTGGTATCAAGCCAAAACTTTACGGTGAAGGTAAAAACGTTCGTG   19316
PMP1611      TTACTAACATCCTAAGTGGTATCAAGCCAAAACTTTACGGTGAAGGTAAAAACGTTCGTG   19316
             ************************************************************

PMP1610      ACTGGATTCATACCAATGACCATTCTTCAGGAGTTTGGACAATCTTGACAAAAGGGCAAA   19376
PMP1666      ACTGGATTCATACCAATGACCATTCTTCAGGAGTTTGGACAATCTTGACAAAAGGGCAAA   19380
PMP1665      ACTGGATTCATACCAATGACCATTCTTCAGGAGTTTGGACAATCTTGACAAAAGGGCAAA   19376
PMP1667      ACTGGATTCATACCAATGACCATTCTTCAGGAGTTTGGACAATCTTGACAAAAGGGCAAA   19376
PMP1663      ACTGGATTCATACCAATGACCATTCTTCAGGAGTTTGGACAATCTTGACAAAAGGGCAAA   19376
PMP1664      ACTGGATTCATACCAATGACCATTCTTCAGGAGTTTGGACAATCTTGACAAAAGGGCAAA   19376
PMP1662      ACTGGATTCATACCAATGACCATTCTTCAGGAGTTTGGACAATCTTGACAAAAGGGCAAA   19376
PMP1611      ACTGGATTCATACCAATGACCATTCTTCAGGAGTTTGGACAATCTTGACAAAAGGGCAAA   19376
             ************************************************************

PMP1610      TCGGTGAAACCTACTTGATTGGGGCTGATGGTGAGAAGAACAATAAGGAAGTTTTGGAAC   19436
PMP1666      TCGGTGAAACCTACTTGATTGGGGCTGATGGTGAGAAGAACAATAAGGAAGTTTTGGAAC   19440
PMP1665      TCGGTGAAACCTACTTGATTGGGGCTGATGGTGAGAAGAACAATAAGGAAGTTTTGGAAC   19436
PMP1667      TCGGTGAAACCTACTTGATTGGGGCTGATGGTGAGAAGAACAATAAGGAAGTTTTGGAAC   19436
PMP1663      TCGGTGAAACCTACTTGATTGGGGCTGATGGTGAGAAGAACAATAAGGAAGTTTTGGAAC   19436
PMP1664      TCGGTGAAACCTACTTGATTGGGGCTGATGGTGAGAAGAACAATAAGGAAGTTTTGGAAC   19436
PMP1662      TCGGTGAAACCTACTTGATTGGGGCTGATGGTGAGAAGAACAATAAGGAAGTTTTGGAAC   19436
PMP1611      TCGGTGAAACCTACTTGATTGGGGCTGATGGTGAGAAGAACAATAAGGAAGTTTTGGAAC   19436
             ************************************************************
```

```
PMP1610      TTATCCTTAAGGAAATGGGACAAGCTGCGGATGCCTATGATCATGTGACTGACCGTGCAG   19496
PMP1666      TTATCCTTAAGGAAATGGGACAAGCTGCGGATGCCTATGATCATGTGACTGACCGTGCAG   19500
PMP1665      TTATCCTTAAGGAAATGGGACAAGCTGCGGATGCCTATGATCATGTGACTGACCGTGCAG   19496
PMP1667      TTATCCTTAAGGAAATGGGACAAGCTGCGGATGCCTATGATCATGTGACTGACCGTGCAG   19496
PMP1663      TTATCCTTAAGGAAATGGGACAAGCTGCGGATGCCTATGATCATGTGACTGACCGTGCAG   19496
PMP1664      TTATCCTTAAGGAAATGGGACAAGCTGCGGATGCCTATGATCATGTGACTGACCGTGCAG   19496
PMP1662      TTATCCTTAAGGAAATGGGACAAGCTGCGGATGCCTATGATCATGTGACTGACCGTGCAG   19496
PMP1611      TTATCCTTAAGGAAATGGGACAAGCTGCGGATGCCTATGATCATGTGACTGACCGTGCAG   19496
             ************************************************************

PMP1610      GACATGACCTTCGCTATGCGATTGATGCTAGCAAGCTCCGTGATGAGTTGGGGTGGAAAC   19556
PMP1666      GACATGACCTTCGCTATGCGATTGATGCTAGCAAGCTCCGTGATGAGTTGGGGTGGAAAC   19560
PMP1665      GACATGACCTTCGCTATGCGATTGATGCTAGCAAGCTCCGTGATGAGTTGGGGTGGAAAC   19556
PMP1667      GACATGACCTTCGCTATGCGATTGATGCTAGCAAGCTCCGTGATGAGTTGGGGTGGAAAC   19556
PMP1663      GACATGACCTTCGCTATGCGATTGATGCTAGCAAGCTCCGTGATGAGTTGGGGTGGAAAC   19556
PMP1664      GACATGACCTTCGCTATGCGATTGATGCTAGCAAGCTCCGTGATGAGTTGGGGTGGAAAC   19556
PMP1662      GACATGACCTTCGCTATGCGATTGATGCTAGCAAGCTCCGTGATGAGTTGGGGTGGAAAC   19556
PMP1611      GACATGACCTTCGCTATGCGATTGATGCTAGCAAGCTCCGTGATGAGTTGGGGTGGAAAC   19556
             ************************************************************

PMP1610      CTGAATTTACCAACTTTGAAGCTGGGCTCAAGGCAACAATCAAGTGGTATACAGATAACC   19616
PMP1666      CTGAATTTACCAACTTTGAAGCTGGGCTCAAGGCAACAATCAAGTGGTATACAGATAACC   19620
PMP1665      CTGAATTTACCAACTTTGAAGCTGGGCTCAAGGCAACAATCAAGTGGTATACAGATAACC   19616
PMP1667      CTGAATTTACCAACTTTGAAGCTGGGCTCAAGGCAACAATCAAGTGGTATACAGATAACC   19616
PMP1663      CTGAATTTACCAACTTTGAAGCTGGGCTCAAGGCAACAATCAAGTGGTATACAGATAACC   19616
PMP1664      CTGAATTTACCAACTTTGAAGCTGGGCTCAAGGCAACAATCAAGTGGTATACAGATAACC   19616
PMP1662      CTGAATTTACCAACTTTGAAGCTGGGCTCAAGGCAACAATCAAGTGGTATACAGATAACC   19616
PMP1611      CTGAATTTACCAACTTTGAAGCTGGGCTCAAGGCAACAATCAAGTGGTATACAGATAACC   19616
             ************************************************************

PMP1610      AAGAATGGTGGAAAGCAGAAAAAGAAGCTGTTGAAGCCAATTATGCTAAGACTCAGGAGA   19676
PMP1666      AAGAATGGTGGAAAGCAGAAAAAGAAGCTGTTGAAGCCAATTATGCTAAGACTCAGGAGA   19680
PMP1665      AAGAATGGTGGAAAGCAGAAAAAGAAGCTGTTGAAGCCAATTATGCTAAGACTCAGGAGA   19676
PMP1667      AAGAATGGTGGAAAGCAGAAAAAGAAGCTGTTGAAGCCAATTATGCTAAGACTCAGGAGA   19676
PMP1663      AAGAATGGTGGAAAGCAGAAAAAGAAGCTGTTGAAGCCAATTATGCTAAGACTCAGGAGA   19676
PMP1664      AAGAATGGTGGAAAGCAGAAAAAGAAGCTGTTGAAGCCAATTATGCTAAGACTCAGGAGA   19676
```

```
PMP1662      AAGAATGGTGGAAAGCAGAAAAAGAAGCTGTTGAAGCCAATTATGCTAAGACTCAGGAGA   19676
PMP1611      AAGAATGGTGGAAAGCAGAAAAAGAAGCTGTTGAAGCCAATTATGCTAAGACTCAGGAGA   19676
             ************************************************************

PMP1610      TTATTACAGTATAAAAAGCAGGAAATAGCTGCTTTTTATTGCTATATTGGGAAGAGTTAC   19736
PMP1666      TTATTACAGTATAAAAAGCAGGAAATAGCTGCTTTTTATTGCTATATTGGGAAGAGTTAC   19740
PMP1665      TTATTACAGTATAAAAAGCAGGAAATAGCTGCTTTTTATTGCTATATTGGGAAGAGTTAC   19736
PMP1667      TTATTACAGTATAAAAAGCAGGAAATAGCTGCTTTTTATTGCTATATTGGGAAGAGTTAC   19736
PMP1663      TTATTACAGTATAAAAAGCAGGAAATAGCTGCTTTTTATTGCTATATTGGGAAGAGTTAC   19736
PMP1664      TTATTACAGTATAAAAAGCAGGAAATAGCTGCTTTTTATTGCTATATTGGGAAGAGTTAC   19736
PMP1662      TTATTACAGTATAAAAAGCAGGAAATAGCTGCTTTTTATTGCTATATTGGGAAGAGTTAC   19736
PMP1611      TTATTACAGTATAAAAAGCAGGAAATAGCTGCTTTTTATTGCTATATTGGGAAGAGTTAC   19736
             ************************************************************

PMP1610      ATATTAGAAAGGTCTAGAGATGATTTTAATTACAGGGGCAAATGGCCAATTAGGAACGGA   19796
PMP1666      ATATTAGAAAGGTCTAGAGATGATTTTAATTACAGGGGCAAATGGCCAATTAGGAACGGA   19800
PMP1665      ATATTAGAAAGGTCTAGAGATGATTTTAATTACAGGGGCAAATGGCCAATTAGGAACGGA   19796
PMP1667      ATATTAGAAAGGTCTAGAGATGATTTTAATTACAGGGGCAAATGGCCAATTAGGAACGGA   19796
PMP1663      ATATTAGAAAGGTCTAGAGATGATTTTAATTACAGGGGCAAATGGCCAATTAGGAACGGA   19796
PMP1664      ATATTAGAAAGGTCTAGAGATGATTTTAATTACAGGGGCAAATGGCCAATTAGGAACGGA   19796
PMP1662      ATATTAGAAAGGTCTAGAGATGATTTTAATTACAGGGGCAAATGGCCAATTAGGAACGGA   19796
PMP1611      ATATTAGAAAGGTCTAGAGATGATTTTAATTACAGGGGCAAATGGCCAATTAGGAACGGA   19796
             ************************************************************

PMP1610      ACTTCGCTATTTATTGGATGAACGTAATGAAGAATACGTGGCAGTAGATGTGGCTAAGAT   19856
PMP1666      ACTTCGCTATTTATTGGATGAACGTAATGAAGAATACGTGGCAGTAGATGTGGCTAAGAT   19860
PMP1665      ACTTCGCTATTTATTGGATGAACGTAATGAAGAATACGTGGCAGTAGATGTGGCTAAGAT   19856
PMP1667      ACTTCGCTATTTATTGGATGAACGTAATGAAGAATACGTGGCAGTAGATGTGGCTAAGAT   19856
PMP1663      ACTTCGCTATTTATTGGATGAACGTAATGAAGAATACGTGGCAGTAGATGTGGCTAAGAT   19856
PMP1664      ACTTCGCTATTTATTGGATGAACGTAATGAAGAATACGTGGCAGTAGATGTGGCTAAGAT   19856
PMP1662      ACTTCGCTATTTATTGGATGAACGTAATGAAGAATACGTGGCAGTAGATGTGGCTAAGAT   19856
PMP1611      ACTTCGCTATTTATTGGATGAACGTAATGAAGAATACGTGGCAGTAGATGTGGCTAAGAT   19856
             ************************************************************

PMP1610      GGACATTACCAATGAAGAAATGGTTGAGAAAGTTTTTGAAGAGGTGAAACCGACTTTAGT   19916
PMP1666      GGACATTACCAATGAAGAAATGGTTGAGAAAGTTTTTGAAGAGGTGAAACCGACTTTAGT   19920
```

```
PMP1665      GGACATTACCAATGAAGAAATGGTTGAGAAAGTTTTTGAAGAGGTGAAACCGACTTTAGT   19916
PMP1667      GGACATTACCAATGAAGAAATGGTTGAGAAAGTTTTTGAAGAGGTGAAACCGACTTTAGT   19916
PMP1663      GGACATTACCAATGAAGAAATGGTTGAGAAAGTTTTTGAAGAGGTGAAACCGACTTTAGT   19916
PMP1664      GGACATTACCAATGAAGAAATGGTTGAGAAAGTTTTTGAAGAGGTGAAACCGACTTTAGT   19916
PMP1662      GGACATTACCAATGAAGAAATGGTTGAGAAAGTTTTTGAAGAGGTGAAACCGACTTTAGT   19916
PMP1611      GGACATTACCAATGAAGAAATGGTTGAGAAAGTTTTTGAAGAGGTGAAACCGACTTTAGT   19916
             ************************************************************

PMP1610      CTACCATTGTGCAGCCTACACCGCTGTTGATGCAGCAGAGGATGAAGGAAAAGAGTTGGA   19976
PMP1666      CTACCATTGTGCAGCCTACACCGCTGTTGATGCAGCAGAGGATGAAGGAAAAGAGTTGGA   19980
PMP1665      CTACCATTGTGCAGCCTACACCGCTGTTGATGCAGCAGAGGATGAAGGAAAAGAGTTGGA   19976
PMP1667      CTACCATTGTGCAGCCTACACCGCTGTTGATGCAGCAGAGGATGAAGGAAAAGAGTTGGA   19976
PMP1663      CTACCATTGTGCAGCCTACACCGCTGTTGATGCAGCAGAGGATGAAGGAAAAGAGTTGGA   19976
PMP1664      CTACCATTGTGCAGCCTACACCGCTGTTGATGCAGCAGAGGATGAAGGAAAAGAGTTGGA   19976
PMP1662      CTACCATTGTGCAGCCTACACCGCTGTTGATGCAGCAGAGGATGAAGGAAAAGAGTTGGA   19976
PMP1611      CTACCATTGTGCAGCCTACACCGCTGTTGATGCAGCAGAGGATGAAGGAAAAGAGTTGGA   19976
             ************************************************************

PMP1610      CTTCGCCATCAATGTGACGGGGACAAAAAATGTCGCAAAAGCATCTGAAAAGCATGGTGC   20036
PMP1666      CTTCGCCATCAATGTGACGGGGACAAAAAATGTCGCAAAAGCATCTGAAAAGCATGGTGC   20040
PMP1665      CTTCGCCATCAATGTGACGGGGACAAAAAATGTCGCAAAAGCATCTGAAAAGCATGGTGC   20036
PMP1667      CTTCGCCATCAATGTGACGGGGACAAAAAATGTCGCAAAAGCATCTGAAAAGCATGGTGC   20036
PMP1663      CTTCGCCATCAATGTGACGGGGACAAAAAATGTCGCAAAAGCATCTGAAAAGCATGGTGC   20036
PMP1664      CTTCGCCATCAATGTGACGGGGACAAAAAATGTCGCAAAAGCATCTGAAAAGCATGGTGC   20036
PMP1662      CTTCGCCATCAATGTGACGGGGACAAAAAATGTCGCAAAAGCATCTGAAAAGCATGGTGC   20036
PMP1611      CTTCGCCATCAATGTGACGGGGACAAAAAATGTCGCAAAAGCATCTGAAAAGCATGGTGC   20036
             ************************************************************

PMP1610      AACTCTAGTTTATATTTCTACGGACTATGTCTTTGACGGTAAGAAACCAGTTGGACAAGA   20096
PMP1666      AACTCTAGTTTATATTTCTACGGACTATGTCTTTGACGGTAAGAAACCAGTTGGACAAGA   20100
PMP1665      AACTCTAGTTTATATTTCTACGGACTATGTCTTTGACGGTAAGAAACCAGTTGGACAAGA   20096
PMP1667      AACTCTAGTTTATATTTCTACGGACTATGTCTTTGACGGTAAGAAACCAGTTGGACAAGA   20096
PMP1663      AACTCTAGTTTATATTTCTACGGACTATGTCTTTGACGGTAAGAAACCAGTTGGACAAGA   20096
PMP1664      AACTCTAGTTTATATTTCTACGGACTATGTCTTTGACGGTAAGAAACCAGTTGGACAAGA   20096
PMP1662      AACTCTAGTTTATATTTCTACGGACTATGTCTTTGACGGTAAGAAACCAGTTGGACAAGA   20096
PMP1611      AACTCTAGTTTATATTTCTACGGACTATGTCTTTGACGGTAAGAAACCAGTTGGACAAGA   20096
```

```
                   ************************************************************

PMP1610      GTGGGAAGTTGATGACCGACCAGATCCACAGACAGAATATGGACGCACTAAGCGTATGGG   20156
PMP1666      GTGGGAAGTTGATGACCGACCAGATCCACAGACAGAATATGGACGCACTAAGCGTATGGG   20160
PMP1665      GTGGGAAGTTGATGACCGACCAGATCCACAGACAGAATATGGACGCACTAAGCGTATGGG   20156
PMP1667      GTGGGAAGTTGATGACCGACCAGATCCACAGACAGAATATGGACGCACTAAGCGTATGGG   20156
PMP1663      GTGGGAAGTTGATGACCGACCAGATCCACAGACAGAATATGGACGCACTAAGCGTATGGG   20156
PMP1664      GTGGGAAGTTGATGACCGACCAGATCCACAGACAGAATATGGACGCACTAAGCGTATGGG   20156
PMP1662      GTGGGAAGTTGATGACCGACCAGATCCACAGACAGAATATGGACGCACTAAGCGTATGGG   20156
PMP1611      GTGGGAAGTTGATGACCGACCAGATCCACAGACAGAATATGGACGCACTAAGCGTATGGG   20156
             ************************************************************

PMP1610      GGAAGAGTTAGTTGAGAAGCATGTGTCTAATTTCTATATTATCCGTACTGCCTGGGTATT   20216
PMP1666      GGAAGAGTTAGTTGAGAAGCATGTGTCTAATTTCTATATTATCCGTACTGCCTGGGTATT   20220
PMP1665      GGAAGAGTTAGTTGAGAAGCATGTGTCTAATTTCTATATTATCCGTACTGCCTGGGTATT   20216
PMP1667      GGAAGAGTTAGTTGAGAAGCATGTGTCTAATTTCTATATTATCCGTACTGCCTGGGTATT   20216
PMP1663      GGAAGAGTTAGTTGAGAAGCATGTGTCTAATTTCTATATTATCCGTACTGCCTGGGTATT   20216
PMP1664      GGAAGAGTTAGTTGAGAAGCATGTGTCTAATTTCTATATTATCCGTACTGCCTGGGTATT   20216
PMP1662      GGAAGAGTTAGTTGAGAAGCATGTGTCTAATTTCTATATTATCCGTACTGCCTGGGTATT   20216
PMP1611      GGAAGAGTTAGTTGAGAAGCATGTGTCTAATTTCTATATTATCCGTACTGCCTGGGTATT   20216
             ************************************************************

PMP1610      TGGAAATTATGGCAAAAACTTCGTTTTTACCATGCAAAATCTTGCGAAAACTCATAAGAC   20276
PMP1666      TGGAAATTATGGCAAAAACTTCGTTTTTACCATGCAAAATCTTGCGAAAACTCATAAGAC   20280
PMP1665      TGGAAATTATGGCAAAAACTTCGTTTTTACCATGCAAAATCTTGCGAAAACTCATAAGAC   20276
PMP1667      TGGAAATTATGGCAAAAACTTCGTTTTTACCATGCAAAATCTTGCGAAAACTCATAAGAC   20276
PMP1663      TGGAAATTATGGCAAAAACTTCGTTTTTACCATGCAAAATCTTGCGAAAACTCATAAGAC   20276
PMP1664      TGGAAATTATGGCAAAAACTTCGTTTTTACCATGCAAAATCTTGCGAAAACTCATAAGAC   20276
PMP1662      TGGAAATTATGGCAAAAACTTCGTTTTTACCATGCAAAATCTTGCGAAAACTCATAAGAC   20276
PMP1611      TGGAAATTATGGCAAAAACTTCGTTTTTACCATGCAAAATCTTGCGAAAACTCATAAGAC   20276
             ************************************************************

PMP1610      TTTAACAGTTGTAAATGATCAGTACGGTCGTCCGACTTGGACTCGTACCTTGGCTGAGTT   20336
PMP1666      TTTAACAGTTGTAAATGATCAGTACGGTCGTCCGACTTGGACTCGTACCTTGGCTGAGTT   20340
PMP1665      TTTAACAGTTGTAAATGATCAGTACGGTCGTCCGACTTGGACTCGTACCTTGGCTGAGTT   20336
PMP1667      TTTAACAGTTGTAAATGATCAGTACGGTCGTCCGACTTGGACTCGTACCTTGGCTGAGTT   20336
```

```
PMP1663      TTTAACAGTTGTAAATGATCAGTACGGTCGTCCGACTTGGACTCGTACCTTGGCTGAGTT   20336
PMP1664      TTTAACAGTTGTAAATGATCAGTACGGTCGTCCGACTTGGACTCGTACCTTGGCTGAGTT   20336
PMP1662      TTTAACAGTTGTAAATGATCAGTACGGTCGTCCGACTTGGACTCGTACCTTGGCTGAGTT   20336
PMP1611      TTTAACAGTTGTAAATGATCAGTACGGTCGTCCGACTTGGACTCGTACCTTGGCTGAGTT   20336
             ************************************************************

PMP1610      CATGACCTACCTAGCTGAAAATCGTAAGGAATTTGGTTATTATCATTTGTCAAATGATGC   20396
PMP1666      CATGACCTACCTAGCTGAAAATCGTAAGGAATTTGGTTATTATCATTTGTCAAATGATGC   20400
PMP1665      CATGACCTACCTAGCTGAAAATCGTAAGGAATTTGGTTATTATCATTTGTCAAATGATGC   20396
PMP1667      CATGACCTACCTAGCTGAAAATCGTAAGGAATTTGGTTATTATCATTTGTCAAATGATGC   20396
PMP1663      CATGACCTACCTAGCTGAAAATCGTAAGGAATTTGGTTATTATCATTTGTCAAATGATGC   20396
PMP1664      CATGACCTACCTAGCTGAAAATCGTAAGGAATTTGGTTATTATCATTTGTCAAATGATGC   20396
PMP1662      CATGACCTACCTAGCTGAAAATCGTAAGGAATTTGGTTATTATCATTTGTCAAATGATGC   20396
PMP1611      CATGACCTACCTAGCTGAAAATCGTAAGGAATTTGGTTATTATCATTTGTCAAATGATGC   20396
             ************************************************************

PMP1610      GACAGAAGACACAACATGGTATGATTTTGCAGTTGAAATTTTGAAAGATACAGATGTCGA   20456
PMP1666      GACAGAAGACACAACATGGTATGATTTTGCAGTTGAAATTTTGAAAGATACAGATGTCGA   20460
PMP1665      GACAGAAGACACAACATGGTATGATTTTGCAGTTGAAATTTTGAAAGATACAGATGTCGA   20456
PMP1667      GACAGAAGACACAACATGGTATGATTTTGCAGTTGAAATTTTGAAAGATACAGATGTCGA   20456
PMP1663      GACAGAAGACACAACATGGTATGATTTTGCAGTTGAAATTTTGAAAGATACAGATGTCGA   20456
PMP1664      GACAGAAGACACAACATGGTATGATTTTGCAGTTGAAATTTTGAAAGATACAGATGTCGA   20456
PMP1662      GACAGAAGACACAACATGGTATGATTTTGCAGTTGAAATTTTGAAAGATACAGATGTCGA   20456
PMP1611      GACAGAAGACACAACATGGTATGATTTTGCAGTTGAAATTTTGAAAGATACAGATGTCGA   20456
             ************************************************************

PMP1610      AGTCAAGCCAGTAGATTCCAGTCAATTTCCAGCCAAAGCTAAACGTCCGCTAAACTCAAC   20516
PMP1666      AGTCAAGCCAGTAGATTCCAGTCAATTTCCAGCCAAAGCTAAACGTCCGCTAAACTCAAC   20520
PMP1665      AGTCAAGCCAGTAGATTCCAGTCAATTTCCAGCCAAAGCTAAACGTCCGCTAAACTCAAC   20516
PMP1667      AGTCAAGCCAGTAGATTCCAGTCAATTTCCAGCCAAAGCTAAACGTCCGCTAAACTCAAC   20516
PMP1663      AGTCAAGCCAGTAGATTCCAGTCAATTTCCAGCCAAAGCTAAACGTCCGCTAAACTCAAC   20516
PMP1664      AGTCAAGCCAGTAGATTCCAGTCAATTTCCAGCCAAAGCTAAACGTCCGCTAAACTCAAC   20516
PMP1662      AGTCAAGCCAGTAGATTCCAGTCAATTTCCAGCCAAAGCTAAACGTCCGCTAAACTCAAC   20516
PMP1611      AGTCAAGCCAGTAGATTCCAGTCAATTTCCAGCCAAAGCTAAACGTCCGCTAAACTCAAC   20516
             ************************************************************
```

```
PMP1610     GATGAGCCTGGCCAAAGCCAAAGCTACTGGATTTGTTATTCCAACTTGGCAAGATGCATT   20576
PMP1666     GATGAGCCTGGCCAAAGCCAAAGCTACTGGATTTGTTATTCCAACTTGGCAAGATGCATT   20580
PMP1665     GATGAGCCTGGCCAAAGCCAAAGCTACTGGATTTGTTATTCCAACTTGGCAAGATGCATT   20576
PMP1667     GATGAGCCTGGCCAAAGCCAAAGCTACTGGATTTGTTATTCCAACTTGGCAAGATGCATT   20576
PMP1663     GATGAGCCTGGCCAAAGCCAAAGCTACTGGATTTGTTATTCCAACTTGGCAAGATGCATT   20576
PMP1664     GATGAGCCTGGCCAAAGCCAAAGCTACTGGATTTGTTATTCCAACTTGGCAAGATGCATT   20576
PMP1662     GATGAGCCTGGCCAAAGCCAAAGCTACTGGATTTGTTATTCCAACTTGGCAAGATGCATT   20576
PMP1611     GATGAGCCTGGCCAAAGCCAAAGCTACTGGATTTGTTATTCCAACTTGGCAAGATGCATT   20576
            ************************************************************

PMP1610     GCAAGAATTTTACAAACAAGAAGTGAGATAAGTAGTAGAATGATTTTCTAGTCTAATAAA   20636
PMP1666     GCAAGAATTTTACAAACAAGAAGTGAGATAAGTAGTAGAATGATTTTCTAGTCTAATAAA   20640
PMP1665     GCAAGAATTTTACAAACAAGAAGTGAGATAAGTAGTAGAATGATTTTCTAGTCTAATAAA   20636
PMP1667     GCAAGAATTTTACAAACAAGAAGTGAGATAAGTAGTAGAATGATTTTCTAGTCTAATAAA   20636
PMP1663     GCAAGAATTTTACAAACAAGAAGTGAGATAAGTAGTAGAATGATTTTCTAGTCTAATAAA   20636
PMP1664     GCAAGAATTTTACAAACAAGAAGTGAGATAAGTAGTAGAATGATTTTCTAGTCTAATAAA   20636
PMP1662     GCAAGAATTTTACAAACAAGAAGTGAGATAAGTAGTAGAATGATTTTCTAGTCTAATAAA   20636
PMP1611     GCAAGAATTTTACAAACAAGAAGTGAGATAAGTAGTAGAATGATTTTCTAGTCTAATAAA   20636
            ************************************************************

PMP1610     AGAGGCAGAGAATGAACTCCAAAGGAGCATAAGATGTACGATTATCTTATCGTTGGTGCT   20696
PMP1666     AGAGGCAGAGAATGAACTCCAAAGGAGCATAAGATGTACGATTATCTTATCGTTGGTGCT   20700
PMP1665     AGAGGCAGAGAATGAACTCCAAAGGAGCATAAGATGTACGATTATCTTATCGTTGGTGCT   20696
PMP1667     AGAGGCAGAGAATGAACTCCAAAGGAGCATAAGATGTACGATTATCTTATCGTTGGTGCT   20696
PMP1663     AGAGGCAGAGAATGAACTCCAAAGGAGCATAAGATGTACGATTATCTTATCGTTGGTGCT   20696
PMP1664     AGAGGCAGAGAATGAACTCCAAAGGAGCATAAGATGTACGATTATCTTATCGTTGGTGCT   20696
PMP1662     AGAGGCAGAGAATGAACTCCAAAGGAGCATAAGATGTACGATTATCTTATCGTTGGTGCT   20696
PMP1611     AGAGGCAGAGAATGAACTCCAAAGGAGCATAAGATGTACGATTATCTTATCGTTGGTGCT   20696
            ************************************************************

PMP1610     GATCTCTTTGGCGCATAGCTTTGGCTCAGTTTCTATTATCGCTCACACCATCCATCAGAA   20756
PMP1666     GATCTCTTTGGCGCATAGCTTTGGCTCAGTTTCTATTATCGCTCACACCATCCATCAGAA   20760
PMP1665     GATCTCTTTGGCGCATAGCTTTGGCTCAGTTTCTATTATCGCTCACACCATCCATCAGAA   20756
PMP1667     GATCTCTTTGGCGCATAGCTTTGGCTCAGTTTCTATTATCGCTCACACCATCCATCAGAA   20756
PMP1663     GATCTCTTTGGCGCATAGCTTTGGCTCAGTTTCTATTATCGCTCACACCATCCATCAGAA   20756
PMP1664     GATCTCTTTGGCGCATAGCTTTGGCTCAGTTTCTATTATCGCTCACACCATCCATCAGAA   20756
```

```
PMP1662      GATCTCTTTGGCGCATAGCTTTGGCTCAGTTTCTATTATCGCTCACACCATCCATCAGAA    20756
PMP1611      GATCTCTTTGGCGCATAGCTTTGGCTCAGTTTCTATTATCGCTCACACCATCCATCAGAA    20756
             ************************************************************

PMP1610      GTTTAATTTGAAGGTACCCAATTATCGCCAAGAAGAAGATTGGGCTAGGATGGGTTTACC    20816
PMP1666      GTTTAATTTGAAGGTACCCAATTATCGCCAAGAAGAAGATTGGGCTAGGATGGGTTTACC    20820
PMP1665      GTTTAATTTGAAGGTACCCAATTATCGCCAAGAAGAAGATTGGGCTAGGATGGGTTTACC    20816
PMP1667      GTTTAATTTGAAGGTACCCAATTATCGCCAAGAAGAAGATTGGGCTAGGATGGGTTTACC    20816
PMP1663      GTTTAATTTGAAGGTACCCAATTATCGCCAAGAAGAAGATTGGGCTAGGATGGGTTTACC    20816
PMP1664      GTTTAATTTGAAGGTACCCAATTATCGCCAAGAAGAAGATTGGGCTAGGATGGGTTTACC    20816
PMP1662      GTTTAATTTGAAGGTACCCAATTATCGCCAAGAAGAAGATTGGGCTAGGATGGGTTTACC    20816
PMP1611      GTTTAATTTGAAGGTACCCAATTATCGCCAAGAAGAAGATTGGGCTAGGATGGGTTTACC    20816
             ************************************************************

PMP1610      AATCACACGTAAGGAAATCTCTAATTGGCATATCAAGGCAAGTCAATACTATTTAGAGTC    20876
PMP1666      AATCACACGTAAGGAAATCTCTAATTGGCATATCAAGGCAAGTCAATACTATTTAGAGTC    20880
PMP1665      AATCACACGTAAGGAAATCTCTAATTGGCATATCAAGGCAAGTCAATACTATTTAGAGTC    20876
PMP1667      AATCACACGTAAGGAAATCTCTAATTGGCATATCAAGGCAAGTCAATACTATTTAGAGTC    20876
PMP1663      AATCACACGTAAGGAAATCTCTAATTGGCATATCAAGGCAAGTCAATACTATTTAGAGTC    20876
PMP1664      AATCACACGTAAGGAAATCTCTAATTGGCATATCAAGGCAAGTCAATACTATTTAGAGTC    20876
PMP1662      AATCACACGTAAGGAAATCTCTAATTGGCATATCAAGGCAAGTCAATACTATTTAGAGTC    20876
PMP1611      AATCACACGTAAGGAAATCTCTAATTGGCATATCAAGGCAAGTCAATACTATTTAGAGTC    20876
             ************************************************************

PMP1610      CCTTTATAACCTTTTACGAGAAAAGTTGTTAGAACAACCTCTTCTTCATGCGGATGAAAC    20936
PMP1666      CCTTTATAACCTTTTACGAGAAAAGTTGTTAGAACAACCTCTTCTTCATGCGGATGAAAC    20940
PMP1665      CCTTTATAACCTTTTACGAGAAAAGTTGTTAGAACAACCTCTTCTTCATGCGGATGAAAC    20936
PMP1667      CCTTTATAACCTTTTACGAGAAAAGTTGTTAGAACAACCTCTTCTTCATGCGGATGAAAC    20936
PMP1663      CCTTTATAACCTTTTACGAGAAAAGTTGTTAGAACAACCTCTTCTTCATGCGGATGAAAC    20936
PMP1664      CCTTTATAACCTTTTACGAGAAAAGTTGTTAGAACAACCTCTTCTTCATGCGGATGAAAC    20936
PMP1662      CCTTTATAACCTTTTACGAGAAAAGTTGTTAGAACAACCTCTTCTTCATGCGGATGAAAC    20936
PMP1611      CCTTTATAACCTTTTACGAGAAAAGTTGTTAGAACAACCTCTTCTTCATGCGGATGAAAC    20936
             ************************************************************

PMP1610      CTCTTATCGGGTGCTAGAGAGTGATAGCCATCTGACCTACTATTGGACCTTTTTGTCTGG    20996
PMP1666      CTCTTATCGGGTGCTAGAGAGTGATAGCCATCTGACCTACTATTGGACCTTTTTGTCTGG    21000
```

```
PMP1665      CTCTTATCGGGTGCTAGAGAGTGATAGCCATCTGACCTACTATTGGACCTTTTTGTCTGG    20996
PMP1667      CTCTTATCGGGTGCTAGAGAGTGATAGCCATCTGACCTACTATTGGACCTTTTTGTCTGG    20996
PMP1663      CTCTTATCGGGTGCTAGAGAGTGATAGCCATCTGACCTACTATTGGACCTTTTTGTCTGG    20996
PMP1664      CTCTTATCGGGTGCTAGAGAGTGATAGCCATCTGACCTACTATTGGACCTTTTTGTCTGG    20996
PMP1662      CTCTTATCGGGTGCTAGAGAGTGATAGCCATCTGACCTACTATTGGACCTTTTTGTCTGG    20996
PMP1611      CTCTTATCGGGTGCTAGAGAGTGATAGCCATCTGACCTACTATTGGACCTTTTTGTCTGG    20996
             ************************************************************

PMP1610      GAAAGCTGAGAATCAAGCAATCACGCTGTACCATCATGATCAGCGTCGGAGTGGTTTAGT    21056
PMP1666      GAAAGCTGAGAATCAAGCAATCACGCTGTACCATCATGATCAGCGTCGGAGTGGTTTAGT    21060
PMP1665      GAAAGCTGAGAATCAAGCAATCACGCTGTACCATCATGATCAGCGTCGGAGTGGTTTAGT    21056
PMP1667      GAAAGCTGAGAATCAAGCAATCACGCTGTACCATCATGATCAGCGTCGGAGTGGTTTAGT    21056
PMP1663      GAAAGCTGAGAATCAAGCAATCACGCTGTACCATCATGATCAGCGTCGGAGTGGTTTAGT    21056
PMP1664      GAAAGCTGAGAATCAAGCAATCACGCTGTACCATCATGATCAGCGTCGGAGTGGTTTAGT    21056
PMP1662      GAAAGCTGAGAATCAAGCAATCACGCTGTACCATCATGATCAGCGTCGGAGTGGTTTAGT    21056
PMP1611      GAAAGCTGAGAATCAAGCAATCACGCTGTACCATCATGATCAGCGTCGGAGTGGTTTAGT    21056
             ************************************************************

PMP1610      AGTACAAGAATTCCTAGGAGATTATTCTGGCTATGTGCATTGTGATATGTTGCGGCAGTA    21116
PMP1666      AGTACAAGAATTCCTAGGAGATTATTCTGGCTATGTGCATTGTGATATGTTGCGGCAGTA    21120
PMP1665      AGTACAAGAATTCCTAGGAGATTATTCTGGCTATGTGCATTGTGATATGTTGCGGCAGTA    21116
PMP1667      AGTACAAGAATTCCTAGGAGATTATTCTGGCTATGTGCATTGTGATATGTTGCGGCAGTA    21116
PMP1663      AGTACAAGAATTCCTAGGAGATTATTCTGGCTATGTGCATTGTGATATGTTGCGGCAGTA    21116
PMP1664      AGTACAAGAATTCCTAGGAGATTATTCTGGCTATGTGCATTGTGATATGTTGCGGCAGTA    21116
PMP1662      AGTACAAGAATTCCTAGGAGATTATTCTGGCTATGTGCATTGTGATATGTTGCGGCAGTA    21116
PMP1611      AGTACAAGAATTCCTAGGAGATTATTCTGGCTATGTGCATTGTGATATGTTGCGGCAGTA    21116
             ************************************************************

PMP1610      ACTTAGGACTTTAGTCCTCTAGTTCTGCCTATGCGATAGCAGTCCAAGGTTTAGGAGCAA    21176
PMP1666      ACTTAGGACTTTAGTCCTCTAGTTCTGCCTATGCGATAGCAGTCCAAGGTTTAGGAGCAA    21180
PMP1665      ACTTAGGACTTTAGTCCTCTAGTTCTGCCTATGCGATAGCAGTCCAAGGTTTAGGAGCAA    21176
PMP1667      ACTTAGGACTTTAGTCCTCTAGTTCTGCCTATGCGATAGCAGTCCAAGGTTTAGGAGCAA    21176
PMP1663      ACTTAGGACTTTAGTCCTCTAGTTCTGCCTATGCGATAGCAGTCCAAGGTTTAGGAGCAA    21176
PMP1664      ACTTAGGACTTTAGTCCTCTAGTTCTGCCTATGCGATAGCAGTCCAAGGTTTAGGAGCAA    21176
PMP1662      ACTTAGGACTTTAGTCCTCTAGTTCTGCCTATGCGATAGCAGTCCAAGGTTTAGGAGCAA    21176
PMP1611      ACTTAGGACTTTAGTCCTCTAGTTCTGCCTATGCGATAGCAGTCCAAGGTTTAGGAGCAA    21176
```

```
               ************************************************************

PMP1610        GGCGACGCTAAGCTTGGTAAACTGCGAACCGCTAGAAGCTTATCGTCAACTGGAAGAAGC    21236
PMP1666        GGCGACGCTAAGCTTGGTAAACTGCGAACCGCTAGAAGCTTATCGTCAACTGGAAGAAGC    21240
PMP1665        GGCGACGCTAAGCTTGGTAAACTGCGAACCGCTAGAAGCTTATCGTCAACTGGAAGAAGC    21236
PMP1667        GGCGACGCTAAGCTTGGTAAACTGCGAACCGCTAGAAGCTTATCGTCAACTGGAAGAAGC    21236
PMP1663        GGCGACGCTAAGCTTGGTAAACTGCGAACCGCTAGAAGCTTATCGTCAACTGGAAGAAGC    21236
PMP1664        GGCGACGCTAAGCTTGGTAAACTGCGAACCGCTAGAAGCTTATCGTCAACTGGAAGAAGC    21236
PMP1662        GGCGACGCTAAGCTTGGTAAACTGCGAACCGCTAGAAGCTTATCGTCAACTGGAAGAAGC    21236
PMP1611        GGCGACGCTAAGCTTGGTAAACTGCGAACCGCTAGAAGCTTATCGTCAACTGGAAGAAGC    21236
               ************************************************************

PMP1610        TGAACTTGTTGGATGTTGGGCACATGTGAGAAGGAAGTTTTTTGAAGCGACCCCCAAGCA    21296
PMP1666        TGAACTTGTTGGATGTTGGGCACATGTGAGAAGGAAGTTTTTTGAAGCGACCCCCAAGCA    21300
PMP1665        TGAACTTGTTGGATGTTGGGCACATGTGAGAAGGAAGTTTTTTGAAGCGACCCCCAAGCA    21296
PMP1667        TGAACTTGTTGGATGTTGGGCACATGTGAGAAGGAAGTTTTTTGAAGCGACCCCCAAGCA    21296
PMP1663        TGAACTTGTTGGATGTTGGGCACATGTGAGAAGGAAGTTTTTTGAAGCGACCCCCAAGCA    21296
PMP1664        TGAACTTGTTGGATGTTGGGCACATGTGAGAAGGAAGTTTTTTGAAGCGACCCCCAAGCA    21296
PMP1662        TGAACTTGTTGGATGTTGGGCACATGTGAGAAGGAAGTTTTTTGAAGCGACCCCCAAGCA    21296
PMP1611        TGAACTTGTTGGATGTTGGGCACATGTGAGAAGGAAGTTTTTTGAAGCGACCCCCAAGCA    21296
               ************************************************************

PMP1610        AGCAGATAAATCATCCTTAGGAGCTAAAGGTTTAGCTTATTGTGATCAGTTATTTTCCCT    21356
PMP1666        AGCAGATAAATCATCCTTAGGAGCTAAAGGTTTAGCTTATTGTGATCAGTTATTTTCCCT    21360
PMP1665        AGCAGATAAATCATCCTTAGGAGCTAAAGGTTTAGCTTATTGTGATCAGTTATTTTCCCT    21356
PMP1667        AGCAGATAAATCATCCTTAGGAGCTAAAGGTTTAGCTTATTGTGATCAGTTATTTTCCCT    21356
PMP1663        AGCAGATAAATCATCCTTAGGAGCTAAAGGTTTAGCTTATTGTGATCAGTTATTTTCCCT    21356
PMP1664        AGCAGATAAATCATCCTTAGGAGCTAAAGGTTTAGCTTATTGTGATCAGTTATTTTCCCT    21356
PMP1662        AGCAGATAAATCATCCTTAGGAGCTAAAGGTTTAGCTTATTGTGATCAGTTATTTTCCCT    21356
PMP1611        AGCAGATAAATCATCCTTAGGAGCTAAAGGTTTAGCTTATTGTGATCAGTTATTTTCCCT    21356
               ************************************************************

PMP1610        GGAAAGAGACTGGGAGGCTTTGCCAGCTGATGAACGACTACAGAAACGTCAAGAACATCT    21416
PMP1666        GGAAAGAGACTGGGAGGCTTTGCCAGCTGATGAACGACTACAGAAACGTCAAGAACATCT    21420
PMP1665        GGAAAGAGACTGGGAGGCTTTGCCAGCTGATGAACGACTACAGAAACGTCAAGAACATCT    21416
PMP1667        GGAAAGAGACTGGGAGGCTTTGCCAGCTGATGAACGACTACAGAAACGTCAAGAACATCT    21416
```

```
PMP1663     GGAAAGAGACTGGGAGGCTTTGCCAGCTGATGAACGACTACAGAAACGTCAAGAACATCT   21416
PMP1664     GGAAAGAGACTGGGAGGCTTTGCCAGCTGATGAACGACTACAGAAACGTCAAGAACATCT   21416
PMP1662     GGAAAGAGACTGGGAGGCTTTGCCAGCTGATGAACGACTACAGAAACGTCAAGAACATCT   21416
PMP1611     GGAAAGAGACTGGGAGGCTTTGCCAGCTGATGAACGACTACAGAAACGTCAAGAACATCT   21416
            ************************************************************

PMP1610     CCAGCCCTTAATGGAAGACTTCTTTGCTTAGTGCCGGCGTCAGTCAGTTTTAGCAGGTTC   21476
PMP1666     CCAGCCCTTAATGGAAGACTTCTTTGCTTAGTGCCGGCGTCAGTCAGTTTTAGCAGGTTC   21480
PMP1665     CCAGCCCTTAATGGAAGACTTCTTTGCTTAGTGCCGGCGTCAGTCAGTTTTAGCAGGTTC   21476
PMP1667     CCAGCCCTTAATGGAAGACTTCTTTGCTTAGTGCCGGCGTCAGTCAGTTTTAGCAGGTTC   21476
PMP1663     CCAGCCCTTAATGGAAGACTTCTTTGCTTAGTGCCGGCGTCAGTCAGTTTTAGCAGGTTC   21476
PMP1664     CCAGCCCTTAATGGAAGACTTCTTTGCTTAGTGCCGGCGTCAGTCAGTTTTAGCAGGTTC   21476
PMP1662     CCAGCCCTTAATGGAAGACTTCTTTGCTTAGTGCCGGCGTCAGTCAGTTTTAGCAGGTTC   21476
PMP1611     CCAGCCCTTAATGGAAGACTTCTTTGCTTAGTGCCGGCGTCAGTCAGTTTTAGCAGGTTC   21476
            ************************************************************

PMP1610     AAAACTAGGAAGGGCAATTGAATACAGCCTCAAGTATGAAGAAACCTTTAAGACTATTTT   21536
PMP1666     AAAACTAGGAAGGGCAATTGAATACAGCCTCAAGTATGAAGAAACCTTTAAGACTATTTT   21540
PMP1665     AAAACTAGGAAGGGCAATTGAATACAGCCTCAAGTATGAAGAAACCTTTAAGACTATTTT   21536
PMP1667     AAAACTAGGAAGGGCAATTGAATACAGCCTCAAGTATGAAGAAACCTTTAAGACTATTTT   21536
PMP1663     AAAACTAGGAAGGGCAATTGAATACAGCCTCAAGTATGAAGAAACCTTTAAGACTATTTT   21536
PMP1664     AAAACTAGGAAGGGCAATTGAATACAGCCTCAAGTATGAAGAAACCTTTAAGACTATTTT   21536
PMP1662     AAAACTAGGAAGGGCAATTGAATACAGCCTCAAGTATGAAGAAACCTTTAAGACTATTTT   21536
PMP1611     AAAACTAGGAAGGGCAATTGAATACAGCCTCAAGTATGAAGAAACCTTTAAGACTATTTT   21536
            ************************************************************

PMP1610     GAAAGACGGACATCTGGTCCTTTCCAATAATCTAGCTGAACGCGCCATTAAATCATTGGT   21596
PMP1666     GAAAGACGGACATCTGGTCCTTTCCAATAATCTAGCTGAACGCGCCATTAAATCATTGGT   21600
PMP1665     GAAAGACGGACATCTGGTCCTTTCCAATAATCTAGCTGAACGCGCCATTAAATCATTGGT   21596
PMP1667     GAAAGACGGACATCTGGTCCTTTCCAATAATCTAGCTGAACGCGCCATTAAATCATTGGT   21596
PMP1663     GAAAGACGGACATCTGGTCCTTTCCAATAATCTAGCTGAACGCGCCATTAAATCATTGGT   21596
PMP1664     GAAAGACGGACATCTGGTCCTTTCCAATAATCTAGCTGAACGCGCCATTAAATCATTGGT   21596
PMP1662     GAAAGACGGACATCTGGTCCTTTCCAATAATCTAGCTGAACGCGCCATTAAATCATTGGT   21596
PMP1611     GAAAGACGGACATCTGGTCCTTTCCAATAATCTAGCTGAACGCGCCATTAAATCATTGGT   21596
            ************************************************************
```

```
PMP1610      TATGGGACGGAGTAAAAGAATTCAGTGGACTCTTTTAGCCTAAGCTAAATTTTAAAAAGC   21656
PMP1666      TATGGGACGGAGTAAAAGAATTCAGTGGACTCTTTTAGCCTAAGCTAAATTTTAAAAAGC   21660
PMP1665      TATGGGACGGAGTAAAAGAATTCAGTGGACTCTTTTAGCCTAAGCTAAATTTTAAAAAGC   21656
PMP1667      TATGGGACGGAGTAAAAGAATTCAGTGGACTCTTTTAGCCTAAGCTAAATTTTAAAAAGC   21656
PMP1663      TATGGGACGGAGTAAAAGAATTCAGTGGACTCTTTTAGCCTAAGCTAAATTTTAAAAAGC   21656
PMP1664      TATGGGACGGAGTAAAAGAATTCAGTGGACTCTTTTAGCCTAAGCTAAATTTTAAAAAGC   21656
PMP1662      TATGGGACGGAGTAAAAGAATTCAGTGGACTCTTTTAGCCTAAGCTAAATTTTAAAAAGC   21656
PMP1611      TATGGGACGGAGTAAAAGAATTCAGTGGACTCTTTTAGCCTAAGCTAAATTTTAAAAAGC   21656
             ************************************************************

PMP1610      GAGGGTGGTTATTTTCTCAAAGTTTTGAAGGAGCTAAAGCAACAGCTATTATTATGAGTT   21716
PMP1666      GAGGGTGGTTATTTTCTCAAAGTTTTGAAGGAGCTAAAGCAACAGCTATTATTATGAGTT   21720
PMP1665      GAGGGTGGTTATTTTCTCAAAGTTTTGAAGGAGCTAAAGCAACAGCTATTATTATGAGTT   21716
PMP1667      GAGGGTGGTTATTTTCTCAAAGTTTTGAAGGAGCTAAAGCAACAGCTATTATTATGAGTT   21716
PMP1663      GAGGGTGGTTATTTTCTCAAAGTTTTGAAGGAGCTAAAGCAACAGCTATTATTATGAGTT   21716
PMP1664      GAGGGTGGTTATTTTCTCAAAGTTTTGAAGGAGCTAAAGCAACAGCTATTATTATGAGTT   21716
PMP1662      GAGGGTGGTTATTTTCTCAAAGTTTTGAAGGAGCTAAAGCAACAGCTATTATTATGAGTT   21716
PMP1611      GAGGGTGGTTATTTTCTCAAAGTTTTGAAGGAGCTAAAGCAACAGCTATTATTATGAGTT   21716
             ************************************************************

PMP1610      TGTTGGAAACAGCTAAACGTCATCAATTAAATAGCGAGAAATATCTATTCTATCTTCTAG   21776
PMP1666      TGTTGGAAACAGCTAAACGTCATCAATTAAATAGCGAGAAATATCTATTCTATCTTCTAG   21780
PMP1665      TGTTGGAAACAGCTAAACGTCATCAATTAAATAGCGAGAAATATCTATTCTATCTTCTAG   21776
PMP1667      TGTTGGAAACAGCTAAACGTCATCAATTAAATAGCGAGAAATATCTATTCTATCTTCTAG   21776
PMP1663      TGTTGGAAACAGCTAAACGTCATCAATTAAATAGCGAGAAATATCTATTCTATCTTCTAG   21776
PMP1664      TGTTGGAAACAGCTAAACGTCATCAATTAAATAGCGAGAAATATCTATTCTATCTTCTAG   21776
PMP1662      TGTTGGAAACAGCTAAACGTCATCAATTAAATAGCGAGAAATATCTATTCTATCTTCTAG   21776
PMP1611      TGTTGGAAACAGCTAAACGTCATCAATTAAATAGCGAGAAATATCTATTCTATCTTCTAG   21776
             ************************************************************

PMP1610      AATGTCTTCCAAACGAGGAAACTCTCGTAAACAAAGAGGTTTTAGAGGCTTATTTACCAT   21836
PMP1666      AATGTCTTCCAAACGAGGAAACTCTCGTAAACAAAGAGGTTTTAGAGGCTTATTTACCAT   21840
PMP1665      AATGTCTTCCAAACGAGGAAACTCTCGTAAACAAAGAGGTTTTAGAGGCTTATTTACCAT   21836
PMP1667      AATGTCTTCCAAACGAGGAAACTCTCGTAAACAAAGAGGTTTTAGAGGCTTATTTACCAT   21836
PMP1663      AATGTCTTCCAAACGAGGAAACTCTCGTAAACAAAGAGGTTTTAGAGGCTTATTTACCAT   21836
PMP1664      AATGTCTTCCAAACGAGGAAACTCTCGTAAACAAAGAGGTTTTAGAGGCTTATTTACCAT   21836
```

```
PMP1662       AATGTCTTCCAAACGAGGAAACTCTCGTAAACAAAGAGGTTTTAGAGGCTTATTTACCAT    21836
PMP1611       AATGTCTTCCAAACGAGGAAACTCTCGTAAACAAAGAGGTTTTAGAGGCTTATTTACCAT    21836
              ************************************************************

PMP1610       GGACTAAAGTTGTACAAGAAAAGTGCAAATAAGAAATCTCCAGATTAGGAACTATCCGTG    21896
PMP1666       GGACTAAAGTTGTACAAGAAAAGTGCAAATAAGAAATCTCCAGATTAGGAACTATCCGTG    21900
PMP1665       GGACTAAAGTTGTACAAGAAAAGTGCAAATAAGAAATCTCCAGATTAGGAACTATCCGTG    21896
PMP1667       GGACTAAAGTTGTACAAGAAAAGTGCAAATAAGAAATCTCCAGATTAGGAACTATCCGTG    21896
PMP1663       GGACTAAAGTTGTACAAGAAAAGTGCAAATAAGAAATCTCCAGATTAGGAACTATCCGTG    21896
PMP1664       GGACTAAAGTTGTACAAGAAAAGTGCAAATAAGAAATCTCCAGATTAGGAACTATCCGTG    21896
PMP1662       GGACTAAAGTTGTACAAGAAAAGTGCAAATAAGAAATCTCCAGATTAGGAACTATCCGTG    21896
PMP1611       GGACTAAAGTTGTACAAGAAAAGTGCAAATAAGAAATCTCCAGATTAGGAACTATCCGTG    21896
              ************************************************************

PMP1610       AGTTCTCTAGTCTGGAGATTTTTCAATATACTTCGTTATTGGACGGTTACGATATTCATA    21956
PMP1666       AGTTCTCTAGTCTGGAGATTTTTCAATATACTTCGTTATTGGACGGTTACGATATTCATA    21960
PMP1665       AGTTCTCTAGTCTGGAGATTTTTCAATATACTTCGTTATTGGACGGTTACGATATTCATA    21956
PMP1667       AGTTCTCTAGTCTGGAGATTTTTCAATATACTTCGTTATTGGACGGTTACGATATTCATA    21956
PMP1663       AGTTCTCTAGTCTGGAGATTTTTCAATATACTTCGTTATTGGACGGTTACGATATTCATA    21956
PMP1664       AGTTCTCTAGTCTGGAGATTTTTCAATATACTTCGTTATTGGACGGTTACGATATTCATA    21956
PMP1662       AGTTCTCTAGTCTGGAGATTTTTCAATATACTTCGTTATTGGACGGTTACGATATTCATA    21956
PMP1611       AGTTCTCTAGTCTGGAGATTTTTCAATATACTTCGTTATTGGACGGTTACGATATTCATA    21956
              ************************************************************

PMP1610       TTTTTTGCAAAGATGTTGTTTGAAAAATAATTTTCAAAAATTCTGAAAATTCTGTTGACA    22016
PMP1666       TTTTTTGCAAAGATGTTGTTTGAAAAATAATTTTCAAAAATTCTGAAAATTCTGTTGACA    22020
PMP1665       TTTTTTGCAAAGATGTTGTTTGAAAAATAATTTTCAAAAATTCTGAAAATTCTGTTGACA    22016
PMP1667       TTTTTTGCAAAGATGTTGTTTGAAAAATAATTTTCAAAAATTCTGAAAATTCTGTTGACA    22016
PMP1663       TTTTTTGCAAAGATGTTGTTTGAAAAATAATTTTCAAAAATTCTGAAAATTCTGTTGACA    22016
PMP1664       TTTTTTGCAAAGATGTTGTTTGAAAAATAATTTTCAAAAATTCTGAAAATTCTGTTGACA    22016
PMP1662       TTTTTTGCAAAGATGTTGTTTGAAAAATAATTTTCAAAAATTCTGAAAATTCTGTTGACA    22016
PMP1611       TTTTTTGCAAAGATGTTGTTTGAAAAATAATTTTCAAAAATTCTGAAAATTCTGTTGACA    22016
              ************************************************************

PMP1610       ACTTTCTGAAAAGAGTCTATAATGGAGAGAAAGTTTTAAAGGAGAAAATGATGAAAAGTT    22076
PMP1666       ACTTTCTGAAAAGAGTCTATAATGGAGAGAAAGTTTTAAAGGAGAAAATGATGAAAAGTT    22080
```

```
PMP1665      ACTTTCTGAAAAGAGTCTATAATGGAGAGAAAGTTTTAAAGGAGAAAATGATGAAAAGTT    22076
PMP1667      ACTTTCTGAAAAGAGTCTATAATGGAGAGAAAGTTTTAAAGGAGAAAATGATGAAAAGTT    22076
PMP1663      ACTTTCTGAAAAGAGTCTATAATGGAGAGAAAGTTTTAAAGGAGAAAATGATGAAAAGTT    22076
PMP1664      ACTTTCTGAAAAGAGTCTATAATGGAGAGAAAGTTTTAAAGGAGAAAATGATGAAAAGTT    22076
PMP1662      ACTTTCTGAAAAGAGTCTATAATGGAGAGAAAGTTTTAAAGGAGAAAATGATGAAAAGTT    22076
PMP1611      ACTTTCTGAAAAGAGTCTATAATGGAGAGAAAGTTTTAAAGGAGAAAATGATGAAAAGTT    22076
             ************************************************************

PMP1610      CAAAACTATTTGCCCTTGCGGGCGTGACATTATTGGCGGCGACTACTTTAGCTGCATGCT    22136
PMP1666      CAAAACTATTTGCCCTTGCGGGCGTGACATTATTGGCGGCGACTACTTTAGCTGCATGCT    22140
PMP1665      CAAAACTATTTGCCCTTGCGGGCGTGACATTATTGGCGGCGACTACTTTAGCTGCATGCT    22136
PMP1667      CAAAACTATTTGCCCTTGCGGGCGTGACATTATTGGCGGCGACTACTTTAGCTGCATGCT    22136
PMP1663      CAAAACTATTTGCCCTTGCGGGCGTGACATTATTGGCGGCGACTACTTTAGCTGCATGCT    22136
PMP1664      CAAAACTATTTGCCCTTGCGGGCGTGACATTATTGGCGGCGACTACTTTAGCTGCATGCT    22136
PMP1662      CAAAACTATTTGCCCTTGCGGGCGTGACATTATTGGCGGCGACTACTTTAGCTGCATGCT    22136
PMP1611      CAAAACTATTTGCCCTTGCGGGCGTGACATTATTGGCGGCGACTACTTTAGCTGCATGCT    22136
             ************************************************************

PMP1610      CTGGATCAGGTTCAAGCACTAAAGGTGAGAAGACATTCTCATACATTTATGAGACAGACC    22196
PMP1666      CTGGATCAGGTTCAAGCACTAAAGGTGAGAAGACATTCTCATACATTTATGAGACAGACC    22200
PMP1665      CTGGATCAGGTTCAAGCACTAAAGGTGAGAAGACATTCTCATACATTTATGAGACAGACC    22196
PMP1667      CTGGATCAGGTTCAAGCACTAAAGGTGAGAAGACATTCTCATACATTTATGAGACAGACC    22196
PMP1663      CTGGATCAGGTTCAAGCACTAAAGGTGAGAAGACATTCTCATACATTTATGAGACAGACC    22196
PMP1664      CTGGATCAGGTTCAAGCACTAAAGGTGAGAAGACATTCTCATACATTTATGAGACAGACC    22196
PMP1662      CTGGATCAGGTTCAAGCACTAAAGGTGAGAAGACATTCTCATACATTTATGAGACAGACC    22196
PMP1611      CTGGATCAGGTTCAAGCACTAAAGGTGAGAAGACATTCTCATACATTTATGAGACAGACC    22196
             ************************************************************

PMP1610      CTGATAACCTCAACTATTTGACAACTGCTAAGGCTGCGACAGCAAATATTACCAGTAACG    22256
PMP1666      CTGATAACCTCAACTATTTGACAACTGCTAAGGCTGCGACAGCAAATATTACCAGTAACG    22260
PMP1665      CTGATAACCTCAACTATTTGACAACTGCTAAGGCTGCGACAGCAAATATTACCAGTAACG    22256
PMP1667      CTGATAACCTCAACTATTTGACAACTGCTAAGGCTGCGACAGCAAATATTACCAGTAACG    22256
PMP1663      CTGATAACCTCAACTATTTGACAACTGCTAAGGCTGCGACAGCAAATATTACCAGTAACG    22256
PMP1664      CTGATAACCTCAACTATTTGACAACTGCTAAGGCTGCGACAGCAAATATTACCAGTAACG    22256
PMP1662      CTGATAACCTCAACTATTTGACAACTGCTAAGGCTGCGACAGCAAATATTACCAGTAACG    22256
PMP1611      CTGATAACCTCAACTATTTGACAACTGCTAAGGCTGCGACAGCAAATATTACCAGTAACG    22256
```

```
            ************************************************************

PMP1610     TGGTTGATGGTTTGCTAGAAAATGATCGCTACGGGAACTTTGTGCCGTCTATGGCTGAGG   22316
PMP1666     TGGTTGATGGTTTGCTAGAAAATGATCGCTACGGGAACTTTGTGCCGTCTATGGCTGAGG   22320
PMP1665     TGGTTGATGGTTTGCTAGAAAATGATCGCTACGGGAACTTTGTGCCGTCTATGGCTGAGG   22316
PMP1667     TGGTTGATGGTTTGCTAGAAAATGATCGCTACGGGAACTTTGTGCCGTCTATGGCTGAGG   22316
PMP1663     TGGTTGATGGTTTGCTAGAAAATGATCGCTACGGGAACTTTGTGCCGTCTATGGCTGAGG   22316
PMP1664     TGGTTGATGGTTTGCTAGAAAATGATCGCTACGGGAACTTTGTGCCGTCTATGGCTGAGG   22316
PMP1662     TGGTTGATGGTTTGCTAGAAAATGATCGCTACGGGAACTTTGTGCCGTCTATGGCTGAGG   22316
PMP1611     TGGTTGATGGTTTGCTAGAAAATGATCGCTACGGGAACTTTGTGCCGTCTATGGCTGAGG   22316
            ************************************************************

PMP1610     ATTGGTCTGTATCCAAGGATGGATTGACTTACACTTATACTATCCGTAAGGATGCAAAAT   22376
PMP1666     ATTGGTCTGTATCCAAGGATGGATTGACTTACACTTATACTATCCGTAAGGATGCAAAAT   22380
PMP1665     ATTGGTCTGTATCCAAGGATGGATTGACTTACACTTATACTATCCGTAAGGATGCAAAAT   22376
PMP1667     ATTGGTCTGTATCCAAGGATGGATTGACTTACACTTATACTATCCGTAAGGATGCAAAAT   22376
PMP1663     ATTGGTCTGTATCCAAGGATGGATTGACTTACACTTATACTATCCGTAAGGATGCAAAAT   22376
PMP1664     ATTGGTCTGTATCCAAGGATGGATTGACTTACACTTATACTATCCGTAAGGATGCAAAAT   22376
PMP1662     ATTGGTCTGTATCCAAGGATGGATTGACTTACACTTATACTATCCGTAAGGATGCAAAAT   22376
PMP1611     ATTGGTCTGTATCCAAGGATGGATTGACTTACACTTATACTATCCGTAAGGATGCAAAAT   22376
            ************************************************************

PMP1610     GGTATACTTCTGAAGGTGAAGAATACGCGGCAGTCAAAGCTCAAGACTTTGTAACAGGAC   22436
PMP1666     GGTATACTTCTGAAGGTGAAGAATACGCGGCAGTCAAAGCTCAAGACTTTGTAACAGGAC   22440
PMP1665     GGTATACTTCTGAAGGTGAAGAATACGCGGCAGTCAAAGCTCAAGACTTTGTAACAGGAC   22436
PMP1667     GGTATACTTCTGAAGGTGAAGAATACGCGGCAGTCAAAGCTCAAGACTTTGTAACAGGAC   22436
PMP1663     GGTATACTTCTGAAGGTGAAGAATACGCGGCAGTCAAAGCTCAAGACTTTGTAACAGGAC   22436
PMP1664     GGTATACTTCTGAAGGTGAAGAATACGCGGCAGTCAAAGCTCAAGACTTTGTAACAGGAC   22436
PMP1662     GGTATACTTCTGAAGGTGAAGAATACGCGGCAGTCAAAGCTCAAGACTTTGTAACAGGAC   22436
PMP1611     GGTATACTTCTGAAGGTGAAGAATACGCGGCAGTCAAAGCTCAAGACTTTGTAACAGGAC   22436
            ************************************************************

PMP1610     TAAAATATGCTGCTGATAAAAAATCAGATGCTCTTTACCTTGTTCAAGAATCAATCAAAG   22496
PMP1666     TAAAATATGCTGCTGATAAAAAATCAGATGCTCTTTACCTTGTTCAAGAATCAATCAAAG   22500
PMP1665     TAAAATATGCTGCTGATAAAAAATCAGATGCTCTTTACCTTGTTCAAGAATCAATCAAAG   22496
PMP1667     TAAAATATGCTGCTGATAAAAAATCAGATGCTCTTTACCTTGTTCAAGAATCAATCAAAG   22496
```

```
PMP1663        TAAAATATGCTGCTGATAAAAAATCAGATGCTCTTTACCTTGTTCAAGAATCAATCAAAG    22496
PMP1664        TAAAATATGCTGCTGATAAAAAATCAGATGCTCTTTACCTTGTTCAAGAATCAATCAAAG    22496
PMP1662        TAAAATATGCTGCTGATAAAAAATCAGATGCTCTTTACCTTGTTCAAGAATCAATCAAAG    22496
PMP1611        TAAAATATGCTGCTGATAAAAAATCAGATGCTCTTTACCTTGTTCAAGAATCAATCAAAG    22496
               ************************************************************

PMP1610        GGTTGGATGCCTATGTAAAAGGGGAAATCAAAGATTTCTCACAAGTAGGAATTAAGGCTT    22556
PMP1666        GGTTGGATGCCTATGTAAAAGGGGAAATCAAAGATTTCTCACAAGTAGGAATTAAGGCTT    22560
PMP1665        GGTTGGATGCCTATGTAAAAGGGGAAATCAAAGATTTCTCACAAGTAGGAATTAAGGCTT    22556
PMP1667        GGTTGGATGCCTATGTAAAAGGGGAAATCAAAGATTTCTCACAAGTAGGAATTAAGGCTT    22556
PMP1663        GGTTGGATGCCTATGTAAAAGGGGAAATCAAAGATTTCTCACAAGTAGGAATTAAGGCTT    22556
PMP1664        GGTTGGATGCCTATGTAAAAGGGGAAATCAAAGATTTCTCACAAGTAGGAATTAAGGCTT    22556
PMP1662        GGTTGGATGCCTATGTAAAAGGGGAAATCAAAGATTTCTCACAAGTAGGAATTAAGGCTT    22556
PMP1611        GGTTGGATGCCTATGTAAAAGGGGAAATCAAAGATTTCTCACAAGTAGGAATTAAGGCTT    22556
               ************************************************************

PMP1610        TGGATGATCAGACAGTTCAGTACACTTTGAACAAACCAGAAAGCTTCTGGAATTCTAAGA    22616
PMP1666        TGGATGATCAGACAGTTCAGTACACTTTGAACAAACCAGAAAGCTTCTGGAATTCTAAGA    22620
PMP1665        TGGATGATCAGACAGTTCAGTACACTTTGAACAAACCAGAAAGCTTCTGGAATTCTAAGA    22616
PMP1667        TGGATGATCAGACAGTTCAGTACACTTTGAACAAACCAGAAAGCTTCTGGAATTCTAAGA    22616
PMP1663        TGGATGATCAGACAGTTCAGTACACTTTGAACAAACCAGAAAGCTTCTGGAATTCTAAGA    22616
PMP1664        TGGATGATCAGACAGTTCAGTACACTTTGAACAAACCAGAAAGCTTCTGGAATTCTAAGA    22616
PMP1662        TGGATGATCAGACAGTTCAGTACACTTTGAACAAACCAGAAAGCTTCTGGAATTCTAAGA    22616
PMP1611        TGGATGATCAGACAGTTCAGTACACTTTGAACAAACCAGAAAGCTTCTGGAATTCTAAGA    22616
               ************************************************************

PMP1610        CAACCATGGGTGTGCTTGCGCCAGTTAATGAAGAGTTTTTGAATTCAAAAGGAGATGATT    22676
PMP1666        CAACCATGGGTGTGCTTGCGCCAGTTAATGAAGAGTTTTTGAATTCAAAAGGAGATGATT    22680
PMP1665        CAACCATGGGTGTGCTTGCGCCAGTTAATGAAGAGTTTTTGAATTCAAAAGGAGATGATT    22676
PMP1667        CAACCATGGGTGTGCTTGCGCCAGTTAATGAAGAGTTTTTGAATTCAAAAGGAGATGATT    22676
PMP1663        CAACCATGGGTGTGCTTGCGCCAGTTAATGAAGAGTTTTTGAATTCAAAAGGAGATGATT    22676
PMP1664        CAACCATGGGTGTGCTTGCGCCAGTTAATGAAGAGTTTTTGAATTCAAAAGGAGATGATT    22676
PMP1662        CAACCATGGGTGTGCTTGCGCCAGTTAATGAAGAGTTTTTGAATTCAAAAGGAGATGATT    22676
PMP1611        CAACCATGGGTGTGCTTGCGCCAGTTAATGAAGAGTTTTTGAATTCAAAAGGAGATGATT    22676
               ************************************************************
```

```
PMP1610     TTGCCAAAGCTACGGATCCAAGTAGTCTCTTGTATAATGGACCTTATTTGTTGAAATCCA   22736
PMP1666     TTGCCAAAGCTACGGATCCAAGTAGTCTCTTGTATAATGGACCTTATTTGTTGAAATCCA   22740
PMP1665     TTGCCAAAGCTACGGATCCAAGTAGTCTCTTGTATAATGGACCTTATTTGTTGAAATCCA   22736
PMP1667     TTGCCAAAGCTACGGATCCAAGTAGTCTCTTGTATAATGGACCTTATTTGTTGAAATCCA   22736
PMP1663     TTGCCAAAGCTACGGATCCAAGTAGTCTCTTGTATAATGGACCTTATTTGTTGAAATCCA   22736
PMP1664     TTGCCAAAGCTACGGATCCAAGTAGTCTCTTGTATAATGGACCTTATTTGTTGAAATCCA   22736
PMP1662     TTGCCAAAGCTACGGATCCAAGTAGTCTCTTGTATAATGGACCTTATTTGTTGAAATCCA   22736
PMP1611     TTGCCAAAGCTACGGATCCAAGTAGTCTCTTGTATAATGGACCTTATTTGTTGAAATCCA   22736
            ************************************************************

PMP1610     TTGTGACCAAATCTTCTGTTGAATTTGCGAAAAATCCGAACTACTGGGATAAGGACAATG   22796
PMP1666     TTGTGACCAAATCTTCTGTTGAATTTGCGAAAAATCCGAACTACTGGGATAAGGACAATG   22800
PMP1665     TTGTGACCAAATCTTCTGTTGAATTTGCGAAAAATCCGAACTACTGGGATAAGGACAATG   22796
PMP1667     TTGTGACCAAATCTTCTGTTGAATTTGCGAAAAATCCGAACTACTGGGATAAGGACAATG   22796
PMP1663     TTGTGACCAAATCTTCTGTTGAATTTGCGAAAAATCCGAACTACTGGGATAAGGACAATG   22796
PMP1664     TTGTGACCAAATCTTCTGTTGAATTTGCGAAAAATCCGAACTACTGGGATAAGGACAATG   22796
PMP1662     TTGTGACCAAATCTTCTGTTGAATTTGCGAAAAATCCGAACTACTGGGATAAGGACAATG   22796
PMP1611     TTGTGACCAAATCTTCTGTTGAATTTGCGAAAAATCCGAACTACTGGGATAAGGACAATG   22796
            ************************************************************

PMP1610     TGCATGTTGACAAAGTTAAATTGTCATTCTGGGATGGTCAAGATACCAGCAAACCTGCAG   22856
PMP1666     TGCATATTGACAAAGTTAAATTGTCATTCTGGGATGGTCAAGATACCAGCAAACCTGCAG   22860
PMP1665     TGCATATTGACAAAGTTAAATTGTCATTCTGGGATGGTCAAGATACCAGCAAACCTGCAG   22856
PMP1667     TGCATATTGACAAAGTTAAATTGTCATTCTGGGATGGTCAAGATACCAGCAAACCTGCAG   22856
PMP1663     TGCATATTGACAAAGTTAAATTGTCATTCTGGGATGGTCAAGATACCAGCAAACCTGCAG   22856
PMP1664     TGCATATTGACAAAGTTAAATTGTCATTCTGGGATGGTCAAGATACCAGCAAACCTGCAG   22856
PMP1662     TGCATATTGACAAAGTTAAATTGTCATTCTGGGATGGTCAAGATACCAGCAAACCTGCAG   22856
PMP1611     TGCATATTGACAAAGTTAAATTGTCATTCTGGGATGGTCAAGATACCAGCAAACCTGCAG   22856
            ***** ******************************************************

PMP1610     AAAACTTTAAAGATGGTAGCCTTACAGCAGCTCGTCTCTATCCAACAAGTGCAAGTTTCG   22916
PMP1666     AAAACTTTAAAGATGGTAGCCTTACAGCAGCTCGTCTCTATCCAACAAGTGCAAGTTTCG   22920
PMP1665     AAAACTTTAAAGATGGTAGCCTTACAGCAGCTCGTCTCTATCCAACAAGTGCAAGTTTCG   22916
PMP1667     AAAACTTTAAAGATGGTAGCCTTACAGCAGCTCGTCTCTATCCAACAAGTGCAAGTTTCG   22916
PMP1663     AAAACTTTAAAGATGGTAGCCTTACAGCAGCTCGTCTCTATCCAACAAGTGCAAGTTTCG   22916
PMP1664     AAAACTTTAAAGATGGTAGCCTTACAGCAGCTCGTCTCTATCCAACAAGTGCAAGTTTCG   22916
```

```
PMP1662      AAAACTTTAAAGATGGTAGCCTTACAGCAGCTCGTCTCTATCCAACAAGTGCAAGTTTCG   22916
PMP1611      AAAACTTTAAAGATGGTAGCCTTACAGCAGCTCGTCTCTATCCAACAAGTGCAAGTTTCG   22916
             ************************************************************

PMP1610      CAGAGCTTGAGAAGAGTATGAAGGACAATATTGTCTATACTCAACAAGACTCTATTACGT   22976
PMP1666      CAGAGCTTGAGAAGAGTATGAAGGACAATATTGTCTATACTCAACAAGACTCTATTACGT   22980
PMP1665      CAGAGCTTGAGAAGAGTATGAAGGACAATATTGTCTATACTCAACAAGACTCTATTACGT   22976
PMP1667      CAGAGCTTGAGAAGAGTATGAAGGACAATATTGTCTATACTCAACAAGACTCTATTACGT   22976
PMP1663      CAGAGCTTGAGAAGAGTATGAAGGACAATATTGTCTATACTCAACAAGACTCTATTACGT   22976
PMP1664      CAGAGCTTGAGAAGAGTATGAAGGACAATATTGTCTATACTCAACAAGACTCTATTACGT   22976
PMP1662      CAGAGCTTGAGAAGAGTATGAAGGACAATATTGTCTATACTCAACAAGACTCTATTACGT   22976
PMP1611      CAGAGCTTGAGAAGAGTATGAAGGACAATATTGTCTATACTCAACAAGACTCTATTACGT   22976
             ************************************************************

PMP1610      ATCTAGTTGGTACAAATATTGACCGTCAGTCCTATAAATACACATCTAAGACCAGCGACG   23036
PMP1666      ATCTAGTTGGTACAAATATTGACCGTCAGTCCTATAAATACACATCTAAGACCAGCGACG   23040
PMP1665      ATCTAGTTGGTACAAATATTGACCGTCAGTCCTATAAATACACATCTAAGACCAGCGACG   23036
PMP1667      ATCTAGTTGGTACAAATATTGACCGTCAGTCCTATAAATACACATCTAAGACCAGCGACG   23036
PMP1663      ATCTAGTTGGTACAAATATTGACCGTCAGTCCTATAAATACACATCTAAGACCAGCGACG   23036
PMP1664      ATCTAGTTGGTACAAATATTGACCGTCAGTCCTATAAATACACATCTAAGACCAGCGACG   23036
PMP1662      ATCTAGTTGGTACAAATATTGACCGTCAGTCCTATAAATACACATCTAAGACCAGCGACG   23036
PMP1611      ATCTAGTTGGTACAAATATTGACCGTCAGTCCTATAAATACACATCTAAGACCAGCGACG   23036
             ************************************************************

PMP1610      AACAAAAGGCATCGACTAAAAAGGCTCTCTTAAACAAGGATTTCCGTCAGGCTATTGCCT   23096
PMP1666      AACAAAAGGCATCGACTAAAAAGGCTCTCTTAAACAAGGATTTCCGTCAGGCTATTGCCT   23100
PMP1665      AACAAAAGGCATCGACTAAAAAGGCTCTCTTAAACAAGGATTTCCGTCAGGCTATTGCCT   23096
PMP1667      AACAAAAGGCATCGACTAAAAAGGCTCTCTTAAACAAGGATTTCCGTCAGGCTATTGCCT   23096
PMP1663      AACAAAAGGCATCGACTAAAAAGGCTCTCTTAAACAAGGATTTCCGTCAGGCTATTGCCT   23096
PMP1664      AACAAAAGGCATCGACTAAAAAGGCTCTCTTAAACAAGGATTTCCGTCAGGCTATTGCCT   23096
PMP1662      AACAAAAGGCATCGACTAAAAAGGCTCTCTTAAACAAGGATTTCCGTCAGGCTATTGCCT   23096
PMP1611      AACAAAAGGCATCGACTAAAAAGGCTCTCTTAAACAAGGATTTCCGTCAGGCTATTGCCT   23096
             ************************************************************

PMP1610      TTGGATTTGACCGTACAGCCTATGCCTCTCAGTTGAATGGACAAACTGGAGCAAGCAAAA   23156
PMP1666      TTGGTTTTGATCGTACAGCCTATGCCTCTCAGTTGAATGGACAAACTGGAGCAAGTAAAA   23160
```

```
PMP1665      TTGGTTTTGATCGTACAGCCTATGCCTCTCAGTTGAATGGACAAACTGGAGCAAGTAAAA   23156
PMP1667      TTGGTTTTGATCGTACAGCCTATGCCTCTCAGTTGAATGGACAAACTGGAGCAAGTAAAA   23156
PMP1663      TTGGTTTTGATCGTACAGCCTATGCCTCTCAGTTGAATGGACAAACTGGAGCAAGTAAAA   23156
PMP1664      TTGGTTTTGATCGTACAGCCTATGCCTCTCAGTTGAATGGACAAACTGGAGCAAGTAAAA   23156
PMP1662      TTGGTTTTGATCGTACAGCCTATGCCTCTCAGTTGAATGGACAAACTGGAGCAAGTAAAA   23156
PMP1611      TTGGTTTTGATCGTACAGCCTATGCCTCTCAGTTGAATGGACAAACTGGAGCAAGTAAAA   23156
             **** ***** ******************************************** ****

PMP1610      TCTTACGTAATATCTTTGTTCCACCAACATTTGTTCAAGCAGATGGTAAAAACTTTGGCG   23216
PMP1666      TCTTGCGTAATCTCTTTGTGCCACCAACATTTGTTCAAGCAGATGGTAAAAACTTTGGCG   23220
PMP1665      TCTTGCGTAATCTCTTTGTGCCACCAACATTTGTTCAAGCAGATGGTAAAAACTTTGGCG   23216
PMP1667      TCTTGCGTAATCTCTTTGTGCCACCAACATTTGTTCAAGCAGATGGTAAAAACTTTGGCG   23216
PMP1663      TCTTGCGTAATCTCTTTGTGCCACCAACATTTGTTCAAGCAGATGGTAAAAACTTTGGCG   23216
PMP1664      TCTTGCGTAATCTCTTTGTGCCACCAACATTTGTTCAAGCAGATGGTAAAAACTTTGGCG   23216
PMP1662      TCTTGCGTAATCTCTTTGTGCCACCAACATTTGTTCAAGCAGATGGTAAAAACTTTGGCG   23216
PMP1611      TCTTGCGTAATCTCTTTGTGCCACCAACATTTGTTCAAGCAGATGGTAAAAACTTTGGCG   23216
             **** ****** ******* ****************************************

PMP1610      ATATGGTCAAAGAGAAATTGGTTACTTATGGGGATGAATGGAAGGATGTTAATCTTGCAG   23276
PMP1666      ATATGGTCAAAGAGAAATTGGTCACTTATGGGGATGAATGGAAGGATGTTAATCTTGCAG   23280
PMP1665      ATATGGTCAAAGAGAAATTGGTCACTTATGGGGATGAATGGAAGGATGTTAATCTTGCAG   23276
PMP1667      ATATGGTCAAAGAGAAATTGGTCACTTATGGGGATGAATGGAAGGATGTTAATCTTGCAG   23276
PMP1663      ATATGGTCAAAGAGAAATTGGTCACTTATGGGGATGAATGGAAGGATGTTAATCTTGCAG   23276
PMP1664      ATATGGTCAAAGAGAAATTGGTCACTTATGGGGATGAATGGAAGGATGTTAATCTTGCAG   23276
PMP1662      ATATGGTCAAAGAGAAATTGGTCACTTATGGGGATGAATGGAAGGATGTTAATCTTGCAG   23276
PMP1611      ATATGGTCAAAGAGAAATTGGTCACTTATGGGGATGAATGGAAGGATGTTAATCTTGCAG   23276
             ********************** *************************************

PMP1610      ATTCTCAGGATGGTCTTTACAATCCAGAAAAAGCCAAGGCTGAATTTGCTAAAGCTAAAT   23336
PMP1666      ATTCTCAGGATGGTCTTTACAATCCAGAAAAAGCCAAGGCTGAATTTGCTAAAGCTAAAT   23340
PMP1665      ATTCTCAGGATGGTCTTTACAATCCAGAAAAAGCCAAGGCTGAATTTGCTAAAGCTAAAT   23336
PMP1667      ATTCTCAGGATGGTCTTTACAATCCAGAAAAAGCCAAGGCTGAATTTGCTAAAGCTAAAT   23336
PMP1663      ATTCTCAGGATGGTCTTTACAATCCAGAAAAAGCCAAGGCTGAATTTGCTAAAGCTAAAT   23336
PMP1664      ATTCTCAGGATGGTCTTTACAATCCAGAAAAAGCCAAGGCTGAATTTGCTAAAGCTAAAT   23336
PMP1662      ATTCTCAGGATGGTCTTTACAATCCAGAAAAAGCCAAGGCTGAATTTGCTAAAGCTAAAT   23336
PMP1611      ATTCTCAGGATGGTCTTTACAATCCAGAAAAAGCCAAGGCTGAATTTGCTAAAGCTAAAT   23336
```

```
                ************************************************************

PMP1610         CAGCCTTACAAGCAGAAGGTGTGACATTCCCAATTCATTTGGATATGCCAGTTGACCAAA    23396
PMP1666         CAGCCTTACAAGCAGAAGGAGTCCAATTCCCAATTCATTTGGATATGCCAGTTGACCAAA    23400
PMP1665         CAGCCTTACAAGCAGAAGGAGTCCAATTCCCAATTCATTTGGATATGCCAGTTGACCAAA    23396
PMP1667         CAGCCTTACAAGCAGAAGGAGTCCAATTCCCAATTCATTTGGATATGCCAGTTGACCAAA    23396
PMP1663         CAGCCTTACAAGCAGAAGGAGTCCAATTCCCAATTCATTTGGATATGCCAGTTGACCAAA    23396
PMP1664         CAGCCTTACAAGCAGAAGGAGTCCAATTCCCAATTCATTTGGATATGCCAGTTGACCAAA    23396
PMP1662         CAGCCTTACAAGCAGAAGGAGTCCAATTCCCAATTCATTTGGATATGCCAGTTGACCAAA    23396
PMP1611         CAGCCTTACAAGCAGAAGGAGTCCAATTCCCAATTCATTTGGATATGCCAGTTGACCAAA    23396
                ******************* **   ***********************************

PMP1610         CAGCAACTACAAAAGTTCAGCGCGTCCAATCTATGAAACAATCCTTGGAAGCAACTTTAG    23456
PMP1666         CAGCAACTACAAAAGTTCAGCGCGTCCAATCTATGAAACAATCCTTGGAAGCAACTTTAG    23460
PMP1665         CAGCAACTACAAAAGTTCAGCGCGTCCAATCTATGAAACAATCCTTGGAAGCAACTTTAG    23456
PMP1667         CAGCAACTACAAAAGTTCAGCGCGTCCAATCTATGAAACAATCCTTGGAAGCAACTTTAG    23456
PMP1663         CAGCAACTACAAAAGTTCAGCGCGTCCAATCTATGAAACAATCCTTGGAAGCAACTTTAG    23456
PMP1664         CAGCAACTACAAAAGTTCAGCGCGTCCAATCTATGAAACAATCCTTGGAAGCAACTTTAG    23456
PMP1662         CAGCAACTACAAAAGTTCAGCGCGTCCAATCTATGAAACAATCCTTGGAAGCAACTTTAG    23456
PMP1611         CAGCAACTACAAAAGTTCAGCGCGTCCAATCTATGAAACAATCCTTGGAAGCAACTTTAG    23456
                ************************************************************

PMP1610         GAGCGGATAATGTAGTCATTGATATTCAACAACTACAAAAAGACGAAGTAAACAATATTA    23516
PMP1666         GAGCTGATAATGTCATTATTGATATTCAACAACTACAAAAAGACGAAGTAAACAATATTA    23520
PMP1665         GAGCTGATAATGTCATTATTGATATTCAACAACTACAAAAAGACGAAGTAAACAATATTA    23516
PMP1667         GAGCTGATAATGTCATTATTGATATTCAACAACTACAAAAAGACGAAGTAAACAATATTA    23516
PMP1663         GAGCTGATAATGTCATTATTGATATTCAACAACTACAAAAAGACGAAGTAAACAATATTA    23516
PMP1664         GAGCTGATAATGTCATTATTGATATTCAACAACTACAAAAAGACGAAGTAAACAATATTA    23516
PMP1662         GAGCTGATAATGTCATTATTGATATTCAACAACTACAAAAAGACGAAGTAAACAATATTA    23516
PMP1611         GAGCTGATAATGTCATTATTGATATTCAACAACTACAAAAAGACGAAGTAAACAATATTA    23516
                **** ********  * *******************************************

PMP1610         CATATTTTGCTGAAAATGCTGCTGGCGAAGACTGGGATTTATCAGATAATGTCGGTTGGG    23576
PMP1666         CATATTTTGCTGAAAATGCTGCTGGCGAAGACTGGGATTTATCAGATAATGTCGGTTGGG    23580
PMP1665         CATATTTTGCTGAAAATGCTGCTGGCGAAGACTGGGATTTATCAGATAATGTCGGTTGGG    23576
PMP1667         CATATTTTGCTGAAAATGCTGCTGGCGAAGACTGGGATTTATCAGATAATGTCGGTTGGG    23576
```

```
PMP1663      CATATTTTGCTGAAAATGCTGCTGGCGAAGACTGGGATTTATCAGATAATGTCGGTTGGG   23576
PMP1664      CATATTTTGCTGAAAATGCTGCTGGCGAAGACTGGGATTTATCAGATAATGTCGGTTGGG   23576
PMP1662      CATATTTTGCTGAAAATGCTGCTGGCGAAGACTGGGATTTATCAGATAATGTCGGTTGGG   23576
PMP1611      CATATTTTGCTGAAAATGCTGCTGGCGAAGACTGGGATTTATCAGATAATGTCGGTTGGG   23576
             ************************************************************

PMP1610      GTCCAGACTTTGCCGATCCATCAACCTACCTTGATATCATCAAACCATCTGTAGGAGAAA   23636
PMP1666      GTCCAGACTTTGCCGATCCATCAACCTACCTTGATATTATCAAACCTTCTGTAGGAGAAA   23640
PMP1665      GTCCAGACTTTGCCGATCCATCAACCTACCTTGATATTATCAAACCTTCTGTAGGAGAAA   23636
PMP1667      GTCCAGACTTTGCCGATCCATCAACCTACCTTGATATTATCAAACCTTCTGTAGGAGAAA   23636
PMP1663      GTCCAGACTTTGCCGATCCATCAACCTACCTTGATATTATCAAACCTTCTGTAGGAGAAA   23636
PMP1664      GTCCAGACTTTGCCGATCCATCAACCTACCTTGATATTATCAAACCTTCTGTAGGAGAAA   23636
PMP1662      GTCCAGACTTTGCCGATCCATCAACCTACCTTGATATTATCAAACCTTCTGTAGGAGAAA   23636
PMP1611      GTCCAGACTTTGCCGATCCATCAACCTACCTTGATATTATCAAACCTTCTGTAGGAGAAA   23636
             ************************************* ******** *************

PMP1610      GTACTAAAACATATTTAGGGTTTGACTCAGGGGAAGATAATGTAGCTGCTAAAAAAGTAG   23696
PMP1666      GTACTAAAACATATTTAGGGTTTGACTCAGGGGAAGATAATGTAGCTGCTAAAAAAGTAG   23700
PMP1665      GTACTAAAACATATTTAGGGTTTGACTCAGGGGAAGATAATGTAGCTGCTAAAAAAGTAG   23696
PMP1667      GTACTAAAACATATTTAGGGTTTGACTCAGGGGAAGATAATGTAGCTGCTAAAAAAGTAG   23696
PMP1663      GTACTAAAACATATTTAGGGTTTGACTCAGGGGAAGATAATGTAGCTGCTAAAAAAGTAG   23696
PMP1664      GTACTAAAACATATTTAGGGTTTGACTCAGGGGAAGATAATGTAGCTGCTAAAAAAGTAG   23696
PMP1662      GTACTAAAACATATTTAGGGTTTGACTCAGGGGAAGATAATGTAGCTGCTAAAAAAGTAG   23696
PMP1611      GTACTAAAACATATTTAGGGTTTGACTCAGGGGAAGATAATGTAGCTGCTAAAAAAGTAG   23696
             ************************************************************

PMP1610      GTCTATATGACTACGAAAAAATTGGTTACTGAGGCTGGTGATGAGACTACAGATGTTGCTA   23756
PMP1666      GTCTATATGACTACGAAAAAATTGGTTACTGAGGCTGGTGATGAGACTACAGATGTTGCTA   23760
PMP1665      GTCTATATGACTACGAAAAAATTGGTTACTGAGGCTGGTGATGAGACTACAGATGTTGCTA   23756
PMP1667      GTCTATATGACTACGAAAAAATTGGTTACTGAGGCTGGTGATGAGACTACAGATGTTGCTA   23756
PMP1663      GTCTATATGACTACGAAAAAATTGGTTACTGAGGCTGGTGATGAGACTACAGATGTTGCTA   23756
PMP1664      GTCTATATGACTACGAAAAAATTGGTTACTGAGGCTGGTGATGAGACTACAGATGTTGCTA   23756
PMP1662      GTCTATATGACTACGAAAAAATTGGTTACTGAGGCTGGTGATGAGACTACAGATGTTGCTA   23756
PMP1611      GTCTATATGACTACGAAAAAATTGGTTACTGAGGCTGGTGATGAGACTACAGATGTTGCTA   23756
             ************************************************************
```

```
PMP1610      AACGCTATGATAAATACGCTGCAGCCCAAGCTTGGTTGACAGATAGTGCTTTGATTATTC   23816
PMP1666      AACGCTATGATAAATACGCTGCAGCCCAAGCTTGGTTGACAGATAGTGCTTTGATTATTC   23820
PMP1665      AACGCTATGATAAATACGCTGCAGCCCAAGCTTGGTTGACAGATAGTGCTTTGATTATTC   23816
PMP1667      AACGCTATGATAAATACGCTGCAGCCCAAGCTTGGTTGACAGATAGTGCTTTGATTATTC   23816
PMP1663      AACGCTATGATAAATACGCTGCAGCCCAAGCTTGGTTGACAGATAGTGCTTTGATTATTC   23816
PMP1664      AACGCTATGATAAATACGCTGCAGCCCAAGCTTGGTTGACAGATAGTGCTTTGATTATTC   23816
PMP1662      AACGCTATGATAAATACGCTGCAGCCCAAGCTTGGTTGACAGATAGTGCTTTGATTATTC   23816
PMP1611      AACGCTATGATAAATACGCTGCAGCCCAAGCTTGGTTGACAGATAGTGCTTTGATTATTC   23816
             ************************************************************

PMP1610      CAACTACATCTCGTACAGGGCGTCCAATCTTGTCTAAGATGGTACCATTTACAATACCAT   23876
PMP1666      CAACTACATCTCGTACAGGGCGTCCAATCTTGTCTAAGATGGTACCATTTACAATACCAT   23880
PMP1665      CAACTACATCTCGTACAGGGCGTCCAATCTTGTCTAAGATGGTACCATTTACAATACCAT   23876
PMP1667      CAACTACATCTCGTACAGGGCGTCCAATCTTGTCTAAGATGGTACCATTTACAATACCAT   23876
PMP1663      CAACTACATCTCGTACAGGGCGTCCAATCTTGTCTAAGATGGTACCATTTACAATACCAT   23876
PMP1664      CAACTACATCTCGTACAGGGCGTCCAATCTTGTCTAAGATGGTACCATTTACAATACCAT   23876
PMP1662      CAACTACATCTCGTACAGGGCGTCCAATCTTGTCTAAGATGGTACCATTTACAATACCAT   23876
PMP1611      CAACTACATCTCGTACAGGGCGTCCAATCTTGTCTAAGATGGTACCATTTACAATACCAT   23876
             ************************************************************

PMP1610      TTGCATTGTCAGGAAATAAAGGTACAAGTGAACCAGTCTTGTATAAATACTTGGAACTTC   23936
PMP1666      TTGCATTGTCAGGAAATAAAGGTACAAGTGAACCAGTCTTGTATAAATACTTGGAACTTC   23940
PMP1665      TTGCATTGTCAGGAAATAAAGGTACAAGTGAACCAGTCTTGTATAAATACTTGGAACTTC   23936
PMP1667      TTGCATTGTCAGGAAATAAAGGTACAAGTGAACCAGTCTTGTATAAATACTTGGAACTTC   23936
PMP1663      TTGCATTGTCAGGAAATAAAGGTACAAGTGAACCAGTCTTGTATAAATACTTGGAACTTC   23936
PMP1664      TTGCATTGTCAGGAAATAAAGGTACAAGTGAACCAGTCTTGTATAAATACTTGGAACTTC   23936
PMP1662      TTGCATTGTCAGGAAATAAAGGTACAAGTGAACCAGTCTTGTATAAATACTTGGAACTTC   23936
PMP1611      TTGCATTGTCAGGAAATAAAGGTACAAGTGAACCAGTCTTGTATAAATACTTGGAACTTC   23936
             ************************************************************

PMP1610      AAGACAAGGCAGTCACTGTAGATGAATATCAAAAAGCTCAGGAAAAATGGATGAAAGAAA   23996
PMP1666      AAGACAAGGCAGTCACTGTAGATGAATACCAAAAAGCTCAGGAAAAATGGATGAAAGAAA   24000
PMP1665      AAGACAAGGCAGTCACTGTAGATGAATACCAAAAAGCTCAGGAAAAATGGATGAAAGAAA   23996
PMP1667      AAGACAAGGCAGTCACTGTAGATGAATACCAAAAAGCTCAGGAAAAATGGATGAAAGAAA   23996
PMP1663      AAGACAAGGCAGTCACTGTAGATGAATACCAAAAAGCTCAGGAAAAATGGATGAAAGAAA   23996
PMP1664      AAGACAAGGCAGTCACTGTAGATGAATACCAAAAAGCTCAGGAAAAATGGATGAAAGAAA   23996
```

```
PMP1662      AAGACAAGGCAGTCACTGTAGATGAATACCAAAAAGCTCAGGAAAAATGGATGAAAGAAA    23996
PMP1611      AAGACAAGGCAGTCACTGTAGATGAATACCAAAAAGCTCAGGAAAAATGGATGAAAGAAA    23996
             **************************** *******************************

PMP1610      AAGAAGAGTCTAATAAAAAGGCTCAAGAAGATCTCGCAAAACATGTGAAATAA    24049
PMP1666      AAGAAGAGTCTAATAAAAAGGCTCAAGAAGATCTCGCAAAACATGTGAAATAA    24053
PMP1665      AAGAAGAGTCTAATAAAAAGGCTCAAGAAGATCTCGCAAAACATGTGAAATAA    24049
PMP1667      AAGAAGAGTCTAATAAAAAGGCTCAAGAAGATCTCGCAAAACATGTGAAATAA    24049
PMP1663      AAGAAGAGTCTAATAAAAAGGCTCAAGAAGATCTCGCAAAACATGTGAAATAA    24049
PMP1664      AAGAAGAGTCTAATAAAAAGGCTCAAGAAGATCTCGCAAAACATGTGAAATAA    24049
PMP1662      AAGAAGAGTCTAATAAAAAGGCTCAAGAAGATCTCGCAAAACATGTGAAATAA    24049
PMP1611      AAGAAGAGTCTAATAAAAAGGCTCAAGAAGATCTCGCAAAACATGTGAAATAA    24049
             *****************************************************
```

**Supplementary Figure S5. DNA sequence alignment of capsule loci from serotype 6B pre-vaccine (GPSC23) and post-vaccine (GPSC6).** Capsule locus and genes upstream and downstream of the capsule locus upstream regulatory sequences (*dexB* to *aliA*) of serotype 6B genomes were examined using Clustal Omega (version 1.2.2) to calculate percent identity matrix and align sequences. Highlighted positions denote differences.

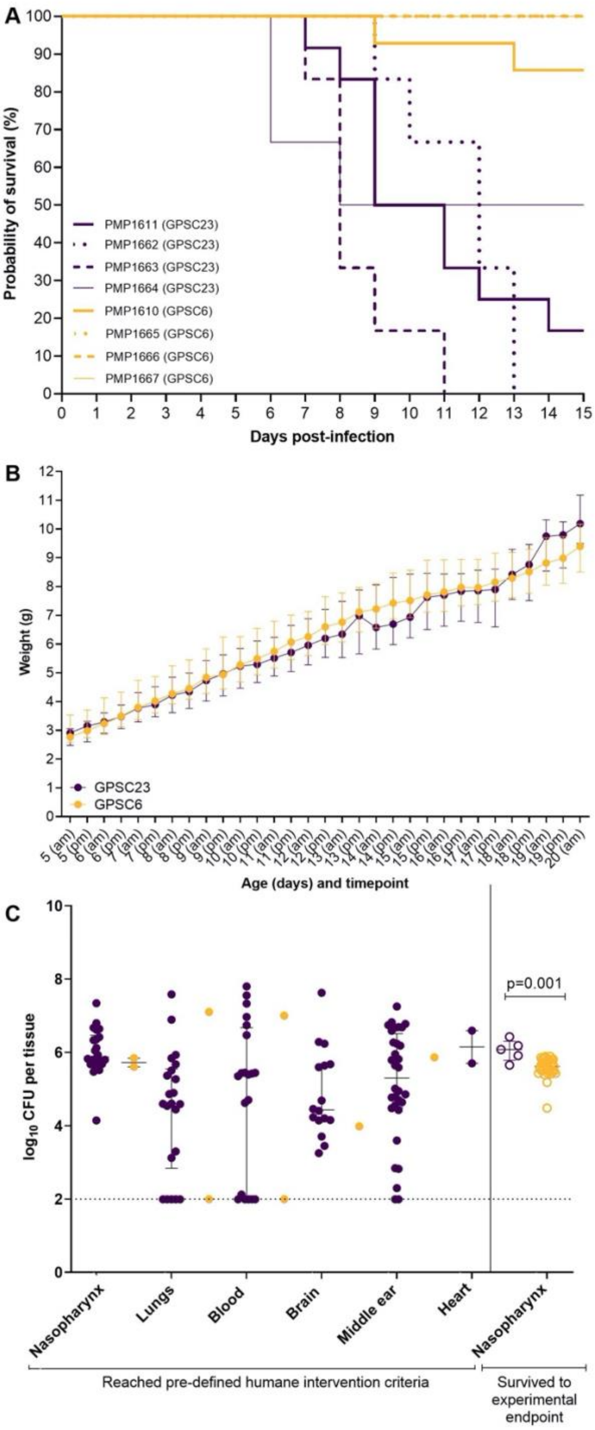

**Supplementary Figure S6. Examination of isolates representing pre- (GPSC23) and post- (GPSC6) vaccine lineages of serotype 6B in a murine model of invasive pneumococcal disease.** Survival curve (A), weight data in grams (B) and pneumococcal loads in the nasopharynx, lungs, blood, brain, middle ear recovered from mice (C) over a 15 day period. At five days old, C57BL/6 mice were infected intranasally with $2x10^3$ CFU in 3 µl without anaesthesia. Mice were monitored for symptoms and euthanised when they met a pre-defined endpoint criterion. Survival was compared using Kaplan-Meier survival curve. Weight data were collected twice daily for the duration of the experiment. Each data point for weight represents the median and error bars represent the IQR. Data on pneumococcal loads in the nasopharynx, lungs, blood, brain and middle ear were recovered from euthanised mice for the duration of the experiment. Individual dots represent an individual mouse, closed circles are mice that reached the pre-defined humane intervention criteria, open circles are mice that surived to experimental endpoint, error bars represent the IQR and the dotted line representing the limit of detection.
